# Introducing RELAX (the Reduction of Electroencephalographic Artifacts): A fully automated pre-processing pipeline for cleaning EEG data - Part 1: Algorithm and Application to Oscillations

**DOI:** 10.1101/2022.03.08.483548

**Authors:** NW Bailey, M Biabani, AT Hill, A Miljevic, NC Rogasch, B McQueen, OW Murphy, PB Fitzgerald

## Abstract

Electroencephalographic (EEG) data is typically contaminated with non-neural artifacts which can confound the results of experiments. Artifact cleaning approaches are available, but often require time-consuming manual input and significant expertise. Advancements in artifact cleaning often only address a single artifact, are only compared against a small selection of pre-existing methods, and seldom assess whether a proposed advancement improves experimental outcomes. To address these issues, we developed RELAX (the Reduction of Electroencephalographic Artifacts), an automated EEG cleaning pipeline implemented within EEGLAB that reduces all artifact types. RELAX cleans continuous data using Multiple Wiener filtering [MWF] and/or wavelet enhanced independent component analysis [wICA] applied to artifacts identified by ICLabel [wICA_ICLabel]). Several versions of RELAX were tested using three datasets containing a mix of cognitive and resting recordings (N = 213, 60 and 23 respectively). RELAX was compared against six commonly used EEG cleaning approaches across a wide range of artifact cleaning quality metrics, including signal-to-error and artifact-to-residue ratios, measures of remaining blink and muscle activity, and the amount of variance explained by experimental manipulations after cleaning. RELAX with MWF and wICA_ICLabel showed amongst the best performance for cleaning blink and muscle artifacts while still preserving neural signal. RELAX with wICA_ICLabel (and no MWF) may perform better at detecting the effect of experimental manipulations on alpha oscillations in working memory tasks. The pipeline is easy to implement in MATLAB and freely available on GitHub. Given its high cleaning performance, objectivity, and ease of use, we recommend RELAX for data cleaning across EEG studies.

## Introduction

Electroencephalography (EEG) allows investigators to non-invasively measure voltage fluctuations generated by the brain. As such, EEG is useful for uncovering functional relationships between neural activity and cognition, as well as for assessing how neural activity differs between healthy and clinical populations. However, the voltage fluctuations detected by EEG electrodes are produced not only by neural activity, but also by non-brain related “artifacts”. These artifacts can be biological in origin, for example eye blinks, movements, and muscle activity. Biological artifacts generate electrical potentials that show stereotypical characteristics, making them reasonably easy to identify (Fitzgibbon et al., 2016; Kleifges et al., 2017; Muthukumaraswamy, 2013). Non-biological artifacts are also often present. For example, voltage drift, which can be caused by changes to scalp electrical impedances due to participant sweat or electrical interference; 50Hz or 60Hz line noise generated by the alternating current of electrical grids; and channel noise due to disrupted electrical connection between electrodes and the scalp (Pion-Tonachini et al., 2019). Because each EEG electrode records a mixture of neural activity and artifacts with an unknown ‘ground-truth’, it is difficult to precisely disentangle the neural activity from artifacts when analysing EEG signals. To address this issue, a multitude of methods for pre-processing EEG data have been developed (see Islam et al. (2016) and Ranjan et al. (2021) for reviews, and Barban et al. (2021) and Robbins et al. (2020) for comparisons across a range of cleaning methods). The fundamental aim of artifact cleaning is to reduce the influence of artifacts on the EEG data while leaving signals from neural activity unaltered.

Although existing EEG pre-processing methods seem adequate, there are still several outstanding issues that may adversely affect the results of EEG research. These limitations include:

1. The lack of a single gold-standard method or even best single approach (Barban et al., 2021). A large part of the reason for the lack of a gold-standard method is that, unfortunately, the ‘ground-truth’ of EEG recordings (real brain signal without any artifacts) is typically unknown. As such, while measures can be used to infer the performance of a cleaning pipeline, researchers cannot *prove* a pipeline leads to perfect signal extraction. As a result, certainty regarding *the* optimal pipeline is elusive. Additionally, newly-developed pipelines are also typically compared against only a small number of existing methods, which further limits the field’s ability to determine an optimal EEG pre-processing pipeline. Similarly, few studies have examined a large range of data cleaning quality metrics simultaneously (Robbins et al., 2020). Demonstrating the superior cleaning of a single artifact type (for example muscle activity) does not necessarily mean the effect of other artifacts (for example blinks) are completely mitigated, so evaluating cleaning efficacy for all artifact types simultaneously is important. Additionally, very few EEG cleaning studies have included an assessment of the most practically important metric - whether cleaning increases the signal-to-noise ratio so that experiments are more likely to detect real effects of interest (Clayson, Baldwin, et al., 2021). While these reasons for the lack of a gold-standard EEG cleaning pipeline are reasonable, the absence of consistent cleaning methods across studies limits our ability to compare findings across different studies, as cleaning methods can vary considerably from study to study. This inconsistency has practical consequences, with research demonstrating variability in experimental results when different cleaning pipelines have been applied even to the same dataset (Robbins et al., 2020).
2. The majority of currently available EEG pre-processing pipelines are only partially automated, leaving results vulnerable to potential bias or experimenter error/subjectivity, as well as being time intensive (and often boring!). The lack of full automation also means that significant technical expertise is required to pre-process the EEG data, increasing the barrier for entry to EEG research and decreasing the potential for student projects to provide meaningful contributions.
3. While fully automated pre-processing methods are available (for example, The Harvard Automated Processing Pipeline for Electroencephalography [HAPPE]; Gabard-Durnam et al. (2018)), in our experience, these methods often do not match our expert judgements, either over or under-rejecting data, leading to reductions in signal (an issue that is demonstrated in the results of the current study).
4. While current EEG pre-processing methods seem to be effective for most data, we have found that some data are not effectively cleaned by existing approaches. In fact, the inspiration for the current work was our observation that using a traditional independent component analysis (ICA) subtraction approach, some data had blink artifacts remaining after cleaning, while other data were over-cleaned, leaving an inverted blink signal in the EEG trace. This effect has also been demonstrated in previous research (Dimigen, 2020).
5. Within cleaning pipelines, there are often many parameters available for adjustment and no clear indication of which parameter settings are optimal. This can lead to confusion and wasted exploratory time for researchers. It also poses the issue that multiple analyses could be undertaken to obtain the desired result. If a well-intentioned researcher assumes an effect is likely, it might seem sensible to apply multiple EEG cleaning approaches to ensure null results were not produced by inferior data cleaning, and to only report the expected positive results. Researchers may do this without realising their results might simply be dependent on the cleaning approach used, reducing the replicability of their results and potential producing false positive results.

To address these issues, we have developed a new EEG pre-processing pipeline by combining and adapting pre-existing approaches and iteratively testing all cleaning parameters in an attempt to find the optimal artifact reduction approach which concurrently preserves neural signal. We have called this pipeline RELAX (short for “Reduction of Electroencephalographic Artifacts”). The pipeline removes extreme outlying electrodes and periods using objective methods derived from previous research. The methods were also subjected to extensive informal testing to maximise outcome metrics, and to ensure a match with our expert judgement. The key artifact reduction components of RELAX are Multi-channel Wiener Filters (MWF) (Borowicz, 2018; Somers et al., 2018) and/or wavelet enhanced ICA (wICA) (Castellanos & Makarov, 2006) applied to artifacts detected by the machine learning algorithm ‘ICLabel’ (Pion-Tonachini et al., 2019). These two approaches are implemented to reduce blinks, muscle activity, horizontal eye movement, voltage drift, and atypical artifacts as effectively as possible while preserving neural signal. The pipeline is implemented in EEGLAB, and is fully automated, while also being modular so that the pipeline (which is freely available on GitHub at https://github.com/NeilwBailey/RELAX/releases) can be easily adapted for optimal cleaning for a user’s intended application. In the following sections we present the results of the formal comparisons across multiple versions of the RELAX pipeline and six commonly used alternative pipelines, focused on data that is typically examined using the power of neural oscillations as an outcome measure. We tested each pipeline in its ability to clean multiple artifact types, as well as its ability to provide data that maximised the detection of experimental effects of interest in a large heterogenous EEG dataset. We also undertook tests of cleaning efficacy in two independent datasets with different recording parameters to ensure generalisability. Additional comparisons between different versions of the RELAX pipeline (with specific parameters varied) are provided in the Supplementary Materials (sections 5-7), which at this stage can be found at the RELAX GitHub repository. Additionally, our companion article explores the application of RELAX to data analysed using event related potentials (ERPs) as an outcome measure (Bailey et al., 2022).

## Methods

### RELAX Pipeline

The RELAX pipeline is implemented on raw continuous multi-channel EEG data in EEGLAB format, and uses EEGLAB (Delorme & Makeig, 2004) and fieldtrip (Oostenveld et al., 2011) functions, implemented within MATLAB. The pipeline outputs cleaned continuous data, with extreme outlying periods and channels removed, referenced to the robust average reference (Bigdely-Shamlo et al., 2015) (note that other re-referencing options are possible from the cleaned data). Both MWF and wICA cleaning methods are used in the RELAX pipeline (with the ability to use only one of these methods available for selection, and recommendations for specific use cases provided in the discussion). In the following, we provide a narrative description of the RELAX pipeline, with a fully detailed explanation in the Supplementary Materials (section 2, pages 5-10).

MWF has recently been proposed as an effective method for cleaning EEG data (Borowicz, 2018; Somers et al., 2018). The MWF toolbox uses a template of time windows of the continuous EEG data that are identified as showing artifacts to determine artifact patterns in the data that should be reduced, and periods identified as clean data to determine the patterns which should be preserved (Somers et al., 2018). These templates of clean data periods and artifact periods are then transformed into covariance matrices (Borowicz, 2018; Somers et al., 2018). Using the template of clean and artifact periods, a low-rank approximation of the artifact is constructed based on the generalised eigenvalue decomposition (Somers et al., 2018). This approximation contains both spatial and spectral information. The multi-channel artifact signal estimate is produced by obtaining a solution to the equation for the mean squared error between the artifact and the clean data (which are both included in the equation), with the optimal solution reflecting the minimum mean squared error. The calculation of this minimum mean squared error provides the artifact signal estimate value. This artifact signal estimate produces a spatial filter, which is extended into the temporal dimension by applying the spatial filter to each channel with a ‘delay period’ – a positive and negative time lag from each timepoint (typically 5-10 samples in each direction), which turns the MWF into a finite impulse response filter as well as a temporal filter, which cleans the data based on both spatial and temporal information. For a more detailed explanation, see Somers et al. (2018). Because the MWF captures both spatial and temporal information in its estimate of the artifact signal, the approach is highly effective at removing artifacts even if they are non-stationary (unlike ICA which only models stationary information) (Borowicz, 2018; Somers et al., 2018). Previously, MWF cleaning has been implemented within the MWF toolbox by manual identification of artifacts (Somers et al., 2019). This is time consuming and vulnerable to inconsistency, mistakes, and subjective decisions, so we have automated the template construction within the RELAX pipeline.

Although MWF cleaning has the aforementioned advantages, it requires a template of both artifact and clean data periods. As such, it cannot address artifacts that are consistent through an entire EEG recording (for example, persistent muscle activity in temporal electrodes or line noise). As such, we tested whether combining wICA and MWF together would address the limitations of each technique when applied separately. wICA has been suggested to address some of the limitations of traditional ICA. Traditional ICA iteratively models signals of putative source components underlying EEG data to derive source signals that are maximally independent (Delorme & Makeig, 2004). Artifactual components are (mostly) independent of neural activity, so by using ICA, researchers can subtract components identified as artifacts, leaving behind only components that reflect neural activity (Delorme et al., 2007). Scalp level EEG data is then reconstructed, resulting in a signal reflective of neural activity with artifact influences minimised (Delorme et al., 2007). However, ICA decomposition is adversely affected by large amplitude artifacts, the presence of which leads to components with a mixture of neural activity and artifact (Akhtar et al., 2012; Anders et al., 2020). Additionally, ICA is never a perfect model, so even without large amplitude artifacts ICA components typically contain a mix of neural activity and artifact (Castellanos & Makarov, 2006; Inuso et al., 2007). Due to this mixing, subtraction of putative artifact components also removes neural activity, while keeping putative artifact components fails to clean the artifact (Akhtar et al., 2012). To remedy these limitations, wICA uses wavelet thresholding of ICA components to detect the oscillatory frequencies with the largest influence on the time course of an ICA component (Castellanos & Makarov, 2006). wICA then subtracts wavelet coefficients above a certain threshold (assumed to reflect artifacts) from the component, leaving behind a component with only the smaller amplitude (presumably neural) activity (Castellanos & Makarov, 2006). As with the ICA subtraction method, this artifact reduced component data is then reconstructed into scalp level EEG data. Previous research has commonly applied wICA to all components (Castellanos & Makarov, 2006). However, our experience indicated that applying wICA to all components reduced considerable amounts of the neural signal as well as the artifacts (which can be seen in our results). To prevent this over-cleaning, RELAX applies wICA cleaning only to components that are identified as artifacts by the machine learning algorithm ICLabel (Pion-Tonachini et al., 2019).

Now that we have introduced the techniques underpinning RELAX, we provide below an abbreviated description of the steps undertaken by the RELAX pipeline, with the full details provided in the Supplementary Materials (section 2, pages 5-10). RELAX implements the following:

1. removes extreme outlying electrodes and time periods using methods adapted from previous research which exactly matched our expert judgement (after extensive informal testing), and which were based on the median and Median Absolute Deviation (MAD) where possible, as this approach produces thresholds that are robust against the effect of more extreme outliers (Alday & van Paridon, 2021);
2. performs three separate sequential MWF cleaning of: i) muscle activity, ii) blinks, and iii) horizontal eye movements and drift;
3. reduces remaining artifacts using wICA, applied only to independent components identified as artifacts by the machine learning algorithm, ICLabel, which has been trained on large datasets to match expert identification of artifact components (Pion-Tonachini et al., 2019). An overview of the specific steps included in our recommended version of the RELAX pipeline is provided in Figure 1, and an abbreviated explanation of the full methods are provided in the following section.

**Figure 1.**
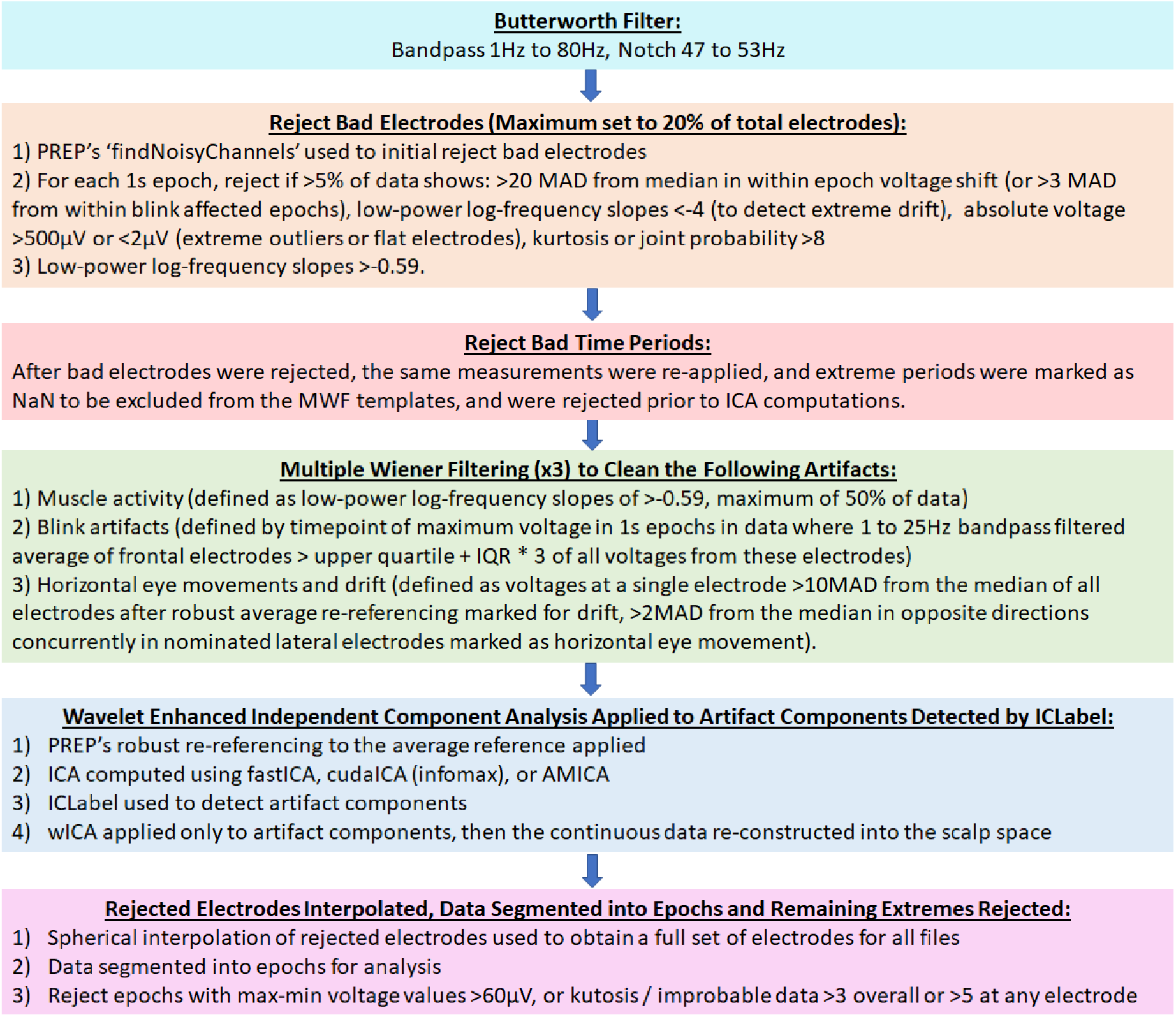
Steps involved in the full RELAX pipeline. Abbreviations: Hz = hertz; s = seconds; PREP = EEG Preprocessing Pipeline; MAD = median absolute deviation; MWF = multiple Wiener filters; ICA = independent component analysis; IQR = inter-quartile range; AMICA adaptive mixture ICA; cudaICA = ICA performed using the cuda cores of a graphic card; wICA = wavelet enhanced ICA.

#### Filtering and Rejection of Extreme Outliers

Firstly, a fourth order acausal Butterworth bandpass filter was applied from 1 to 80Hz and a second order acausal Butterworth notch filter was applied from 47 to 53Hz (this can be easily adapted for 60Hz line noise). Note that ICA performs best with 1Hz high-pass filters (Winkler et al., 2015), and filtering out <1Hz data typically does not affect oscillation measures of interest. However, high-pass filter settings for ERPs likely need to be <0.3Hz, with the exact optimal high-pass filter setting still under debate. We address the issue of using RELAX with high-pass filtering for ERPs in our companion article (Bailey et al., 2022). After filtering, a multi-step process was used to identify and remove bad electrodes. Firstly, the “findNoisyChannels” function from the ‘EEG Preprocessing Pipeline’ (PREP) was applied to provide an initial rejection of bad electrodes (Bigdely-Shamlo et al., 2015). Then data were split into 1 second epochs with a 0.5 second overlap, and within these epochs, RELAX identified: 1) extreme outlying amplitudes unlikely to reflect data periods with retrievable brain activity, which were identified using MAD from the median, providing robust estimates even with extreme outliers (Alday & van Paridon, 2021); 2) extreme drift; 3) extreme kurtosis; 4) extremely improbable voltage distributions; and finally 5) log-power log-frequency slopes indicative of muscle activity (Fitzgibbon et al., 2016) were used to mark electrodes for rejection if >5% of an electrode’s data showed contamination with muscle activity. A limit of removing 20% of electrodes was imposed (a setting which can be adjusted by the user), and if >20% of electrodes were marked for rejection, electrodes were ranked in order of the total number of epochs showing extreme artifacts and only the worst 20% were rejected. Following the electrode rejection step, the same extreme outlier detection approaches were used to mark extreme periods in the EEG data for exclusion. These periods were marked with “not a number” (NaN) within the MWF template. The MWF toolbox ignores these periods when constructing cleaning templates, so that the MWF cleaning focuses on artifacts that are likely to contain retrievable neural activity, rather than periods that may only reflect extreme artifacts without the potential to retrieve underlying neural activity (Somers et al., 2019). These extreme outlier periods were also rejected from the data completely prior to the ICA decomposition for the same reason. Both electrode rejection and extreme outlying period detections were transferred from the epoched data back to the continuous data, which was cleaned in the following steps.

#### Multiple Wiener Filtering

Following the extreme outlier rejection steps, three sequential MWF cleaning steps were implemented to address (in chronological order): 1) muscle activity, then 2) blink activity, then 3) both horizontal eye movement and drift activity together. Note that for some applications, skipping straight to wICA might be preferred, and is easily implemented in RELAX (these applications are explored in the discussion). Firstly, to create a template for the detection of muscle activity for MWF cleaning, the data separated into 1s epochs with a 500ms overlap. For each channel, epochs affected by muscle activity were detected using a log-power log-frequency slope threshold of >-0.59, a threshold chosen because very little data within EEG recordings taken from people who have had their muscles pharmacologically paralysed show slopes above this threshold (Fitzgibbon et al., 2016). A full EEG data length 1D template of muscle artifact periods (marked as 1) and clean periods (marked as 0) was constructed, and the MWF cleaning was applied using this template.

Secondly, to detect blinks, a copy of the data were bandpass filtered using a fourth order Butterworth filter from 1-25Hz (which focused on this data on blink relevant frequencies). We then averaged pre-specified blink affected electrodes (’FP1’; ‘FPZ’; ‘FP2’; ‘AF3’; ‘AF4’; ‘F3’; ‘F1’; ‘FZ’; ‘F2’; ‘F4’ – this list can be easily adapted to match a researcher’s EEG files). Blinks were marked as the maximum point within each time-period that exceeded the value of the upper quartile of all voltages + the interquartile range (IQR) * 3. A blink artifact mask for MWF cleaning was created by marking the 800ms surrounding all blink maximums as artifacts.

Third, drift and horizontal eye movements were identified and cleaned with the MWF. For the drift identification step, a copy of the data were re-referenced using PREP’s robust average re-referencing approach as we found drift to be more accurately identified in average re-referenced data. However, the re-referencing was only used at this stage to identify drift, and re-referenced data were not used in the MWF cleaning as our tests showed MWF performed better when implemented prior to average re-referencing. Epochs that showed an amplitude at any electrode >10MAD from the median of all electrodes were deemed to be affected by drift and marked as artifact periods in the template for MWF cleaning (this approach was adapted from Nolan et al. (2010), who used the standard deviation (SD) instead of MAD to set the threshold). For detection of horizontal eye movements, a list of the lateral electrodes affected by horizontal eye movements is provided by the user. The RELAX pipeline selected a single electrode from both the left and right side of the head from this list, selecting the first electrode in the list if available, then moving to the second electrode in the list if the first electrode had been rejected, and so on until a horizontal eye movement affected electrode from the list was available. Electrodes used in the current study were “F7”, “FT7”, “F5”, “T7”, “FC5”, “C5”, “TP7”, “AF3” for the left side, and “F8”,”FT8”,”F6”,”T8”, “FC6”, “C6”, “TP8”, “AF4” for the right side. Periods where the selected electrodes showed voltages >2MAD from the median of their overall amplitude, with an opposite voltage movement >2MAD from the median polarity shift in the electrode on the opposite side of the head were assumed to reflect horizontal eye movements (Rogasch et al., 2017). These periods were marked as artifact periods in the MWF template. A delay period of 8 samples was used for all MWF cleaning. This delay period implemented a positive and negative time lag for applying the MWF filter, which turned the spatial MWF into a spatio-temporal finite impulse response filter. RELAX was also set to detect generalised eigenvector deficiency, which sometimes occurs with longer delay periods and can impair the ability of the MWF to clean the data. If generalised eigenvector deficiencies were detected, the algorithm reduced the delay period by a value of 1 sample and ran the MWF again. This approach was repeated up to three times, resulting in a minimum delay period of 5 samples. The full explanation of MWF template construction and cleaning process is provided in the Supplementary Materials (section 2, pages 6-9).

#### Wavelet-enhanced Independent Component Analysis

Following the three sequential MWF steps, wICA was used to reduce artifactual components identified by the machine learning algorithm “ICLabel”, then electrode space data was reconstructed. Firstly, the MWF cleaned data were average re-referenced using PREP’s robust re-referencing method, which interpolates missing channels before average re-referencing to prevent reference asymmetries, following the re-removal of the bad channels to prevent rank issues in ICA decompositions (Bigdely-Shamlo et al., 2015). The periods that were marked as extreme outliers (which had previously been marked as NaN for the MWF cleaning) were rejected at this stage. ICA was then computed using either cudaICA (Raimondo et al., 2012), AMICA (Palmer et al., 2012) or fastICA (Hyvarinen, 1999) with the fastICA deflation setting applied to avoid non-convergence issues. ICLabel was used to detect artifactual components (defined by ICLabel as “more likely to reflect any artifact category than to reflect a brain component”), and only these artifactual components were cleaned with wICA before the continuous data were reconstructed back into the scalp space. The above steps left cleaned and robust average re-referenced continuous data which can then be epoched as desired for different types of analyses.

### Comparison Pipelines

To optimize the RELAX pipeline, we tested six potential versions of RELAX with various parameter variations applied. For brevity, we present a summary of the variations of the RELAX pipeline approaches in Table 1. To provide a robust assessment of the performance of our RELAX pipeline, we also compared our pipeline to six of the most commonly reported cleaning approaches. A full explanation of each pipeline is provided in the Supplementary Materials (section 3, page 10), and in the references cited in the following summary: Firstly, we tested six potential versions of our RELAX pipeline. These included using different ICA algorithms, namely, 1) MWF_wICA_infomax, which used extended-infomax via cudaICA (Raimondo et al., 2012), 2) MWF_wICA_fastICA, which used fastICA (Hyvarinen, 1999), or 3) MWF_wICA_AMICA, which used adaptive mixture ICA (AMICA) (Palmer et al., 2012). For brevity, we only report the results of MWF_wICA_infomax in the main manuscript (described simply as MWF_wICA hereafter), since infomax was used as the ICA method for all other pipelines, and the MWF_wICA versions differed only minimally depending on ICA method. We also tested: 4) MWF_ICA_subtract, which subtracted artifactual ICA components instead of using wICA. Further, we tested: 5) MWF_wICA_CCA, which excluded the cleaning of artifactual components identified as muscle by ICLabel from the wICA step in MWF_wICA, and cleaned muscle activity with the extended canonical correlation analysis (CCA) instead (Janani et al., 2018), and finally: 6) wICA_ICLabel, which excluded MWF cleaning completely and only used wICA applied to artifactual components identified with ICLabel (Pion-Tonachini et al., 2019). This pipeline is similar to those used by Issa and Juhasz (2019) and Mammone et al. (2011), except that our approach identified all artifacts instead of only identifying eye movements (Issa & Juhasz, 2019), and used ICLabel to identify artifacts instead of using entropy and kurtosis measures (Mammone et al., 2011). To help the reader assess the potential of the RELAX pipeline for application to their data, we note here that our recommendation is to use either “**<u>MWF_wICA</u>**” or “**<u>wICA_ICLabel</u>**”.

The comparison pipelines we tested were: 7) MWF_only, which was identical to the MWF cleaning steps in our RELAX pipeline but did not apply any additional cleaning after the MWF stage (Somers et al. 2018). 8) MWF_CCA, which applied the sequential MWF cleaned data, but instead of applying wICA to this data as per the RELAX methods, it used the extended CCA to further clean any remaining muscle artifacts (Janani et al., 2018). 9) ICA_subtract, which is probably the most commonly used approach in EEG research – this pipeline computed ICA, then subtracted the components identified as artifacts from the ICA unmixing matrix using ICLabel, then reconstructed the electrode space data (without any MWF cleaning applied) (Pion-Tonachini et al., 2019). 10) ASR applied the Artifact Subspace Reconstruction (ASR) approach followed by ICA subtraction of artifacts identified by ICLabel (Chang et al., 2019). 11) HAPPE applied the Harvard Automated Processing Pipeline for EEG (Gabard-Durnam et al., 2018). Finally, 12) wICA_all, which cleaned data by applying wICA to all components after the ICA decomposition. For brevity, we present a summary of the comparison pipeline approaches in Table 2, and the full description in the Supplementary Materials.

**Table 1.**
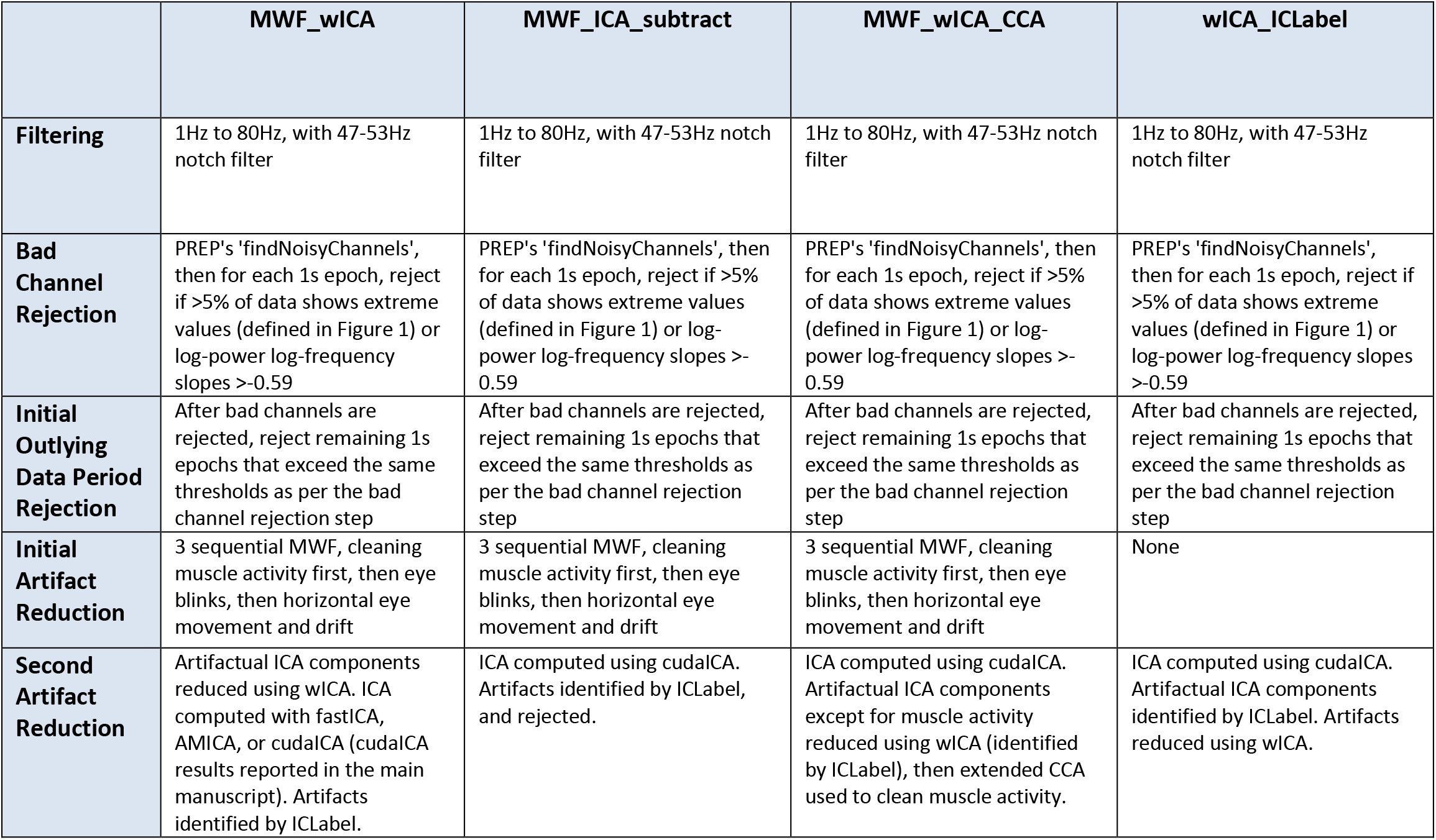
A summary of the steps involved in each variation of the RELAX pipeline. Abbreviations: Hz = hertz; s = seconds; PREP = EEG Preprocessing Pipeline; MAD = median absolute deviation; MWF = multiple Wiener filters; ICA = independent component analysis; IQR = inter-quartile range; AMICA adaptive mixture ICA; cudaICA = ICA performed using the cuda cores of a graphic card; wICA = wavelet enhanced ICA.

**Table 2.** A summary of the steps involved in each of the comparison cleaning pipelines. Abbreviations: Hz = hertz; s = seconds; PREP = EEG Preprocessing Pipeline; MAD = median absolute deviation; MWF = multiple Wiener filters; ICA = independent component analysis; IQR = inter-quartile range; AMICA adaptive mixture ICA; cudaICA = ICA performed using the cuda cores of a graphic card; wICA = wavelet enhanced ICA, ASR = Artifact Subspace Reconstruction; HAPPE = Harvard Automated Processing Pipeline.

|  | ICA_subtract | MWF_only | MWF_CCA | wICA_all | ASR | HAPPE |
| --- | --- | --- | --- | --- | --- | --- |
| **Filtering** | 1Hz to 80Hz, with 47-53Hz notch filter | 1Hz to 80Hz, with 47-53Hz notch filter | 1Hz to 80Hz, with 47-53Hz notch filter | 1Hz to 80Hz, with 47-53Hz notch filter | 1Hz to 80Hz, with 47-53Hz notch filter | 1Hz high-pass, CleanLine to remove 50Hz |
| **Bad Channel Rejection** | PREP's 'findNoisyChannels', then for each 1s epoch, reject if >5% of data shows extreme values (defined in Figure 1) or log-power log-frequency slopes >-0.59 | PREP's 'findNoisyChannels', then for each 1s epoch, reject if >5% of data shows extreme values (defined in Figure 1) or log-power log-frequency slopes >-0.59 | PREP's 'findNoisyChannels', then for each 1s epoch, reject if >5% of data shows extreme values (defined in Figure 1) or log-power log-frequency slopes >-0.59 | PREP's 'findNoisyChannels', then for each 1s epoch, reject if >5% of data shows extreme values (defined in Figure 1) or log-power log-frequency slopes >-0.59 | PREP's 'findNoisyChannels', then for each 1s epoch, reject if >5% of data shows extreme values (defined in Figure 1) or log-power log-frequency slopes >-0.59 | Rejects channels with joint probability >3SD from mean of average log power from 1-125Hz. Performed twice. |
| **Initial Outlying Data Period Rejection** | After bad channels are rejected, reject remaining 1s epochs that exceed the same thresholds as per the bad channel rejection step | After bad channels are rejected, reject remaining 1s epochs that exceed the same thresholds as per the bad channel rejection step | After bad channels are rejected, reject remaining 1s epochs that exceed the same thresholds as per the bad channel rejection step | After bad channels are rejected, reject remaining 1s epochs that exceed the same thresholds as per the bad channel rejection step | After bad channels are rejected, reject remaining 1s epochs that exceed the same thresholds as per the bad channel rejection step | None |
| **Initial Artifact Reduction** | None | 3 sequential MWF, cleaning muscle activity first, then eye blinks, then horizontal eye movement and drift | 3 sequential MWF, cleaning muscle activity first, then eye blinks, then horizontal eye movement and drift | None | ASR: automatic detection of clean data periods to determine thresholds, followed by rejection of components with large variance (similar to principal component analysis), then reconstruction of the original electrode space data. | wICA applied to all components |
| **Second Artifact Reduction** | Artifactual ICA components subtracted (identified by ICLabel) | None | Extended canonical correlation analysis to clean muscle activity (Janani et al., 2018). | All ICA components reduced using wICA | Artifactual ICA components rejected (identified by ICLabel) | Artifactual ICA components rejected (identified by MARA) |

### Data

We examined the effectiveness of each cleaning pipeline primarily on a large combination dataset (N = 213). The data we used were specifically selected to be challenging to clean, as is often the case for typical EEG data. Data were collected mostly by students, with many data files collected while participants performed cognitive tasks, which produced concentration related forehead and temporal muscle activity and blinks. The data also commonly contained bad electrodes, temporarily disconnected electrodes, and other atypical artifacts. As such, the data provided a good “real world” test case for fully automated pre-processing pipelines. In addition, we tested the approach on two additional smaller datasets collected from separate laboratories, with the total data analysed comprising two laboratories, three different EEG systems, and three different tasks (see Supplementary Materials section 5 and 6, pages 45-71 for the results in these additional datasets, as well as our companion article for analysis of a fourth task with RELAX applied to event-related potential analyses: Bailey et al. 2022). All data were recorded from healthy adults with all participants providing informed consent prior to participation. The study was approved by the Ethics Committee of the Alfred Hospital and Monash University. Tasks included the Sternberg task, 2-back task, and Colour-Wheel Recall task (all of which measure working memory), and recordings also included both eyes open (EO) and eyes closed (EC) resting-state. For our primary analyses we combined the datasets that used identical recording parameters and would be analysed in similar ways, hence oscillatory analysis approaches were tested using a combined Sternberg and EO/EC resting-state dataset (N = 213). This combination approach was used to provide robust tests of cleaning quality with large datasets.

The combination dataset comprising our primary analyses were recorded using a Neuroscan Synamps2 amplifier with the SCAN 4.3 software interface (Compumedics, Melbourne, Australia) with a 64-channel Quickcap (excluding CB1 and CB2 electrodes) and a sampling rate of 1000Hz with a 0.01Hz high-pass and 200Hz low pass filter. The ground electrode was located at AFz, and the reference was located between Cz and CPz. The first combined dataset we report included EEG recordings from participants completing a Sternberg task (Bailey et al., 2020), and both EO and EC resting-state recordings (total N = 213 after excluding participants who showed very little blink or muscle artifact even prior to cleaning, EO resting-state N = 93, EC resting-state N = 62, and Sternberg N = 58). Blink metrics were not assessed for the EC resting-state recordings, and 10 files were excluded from blink metric analyses due to not enough blink-locked epochs remaining after the exclusion of epochs that contained multiple blinks (N = 140).

An additional combination of 2-back task related data, as well as EO and EC resting-state recordings, were assessed, reported in the Supplementary Materials (section 5, pages 54-64) (N = 20 healthy adults providing one recording from each type of EEG recording, for a total of 60 EEG files). In this dataset, data were recorded using a Synamps2 amplifier running through the SCAN 4.3 software interface (Compumedics, Melbourne, Australia) with 44 Ag/AgCl electrodes embedded within an EasyCap (Herrsching, Germany). The ground electrode was placed at AFz, and the reference was placed at CPz. A sampling rate of 1000Hz was used for these recordings, with an online bandpass filter between 0.1 to 200 Hz. Finally, one further dataset of Colour-Wheel Recall task data was tested (N = 23) reported in the Supplementary Materials (section 6, pages 65-71). These data were recorded using a Neuroscan amplifier (Compumedics, Melbourne, Australia) with a 62-channel EasyCap (Herrsching, Germany) and a sampling rate of 10kHz, downsampled to 1000Hz prior to cleaning. To reduce computation time, MWF_ICA_subtract, MWF_CCA and MWF_wICA_CCA were not tested on these two datasets, as our primary analysis did not indicate these approaches were superior to MWF_wICA.

Various cleaning quality metrics were calculated, using either continuous or epoched data. Continuous data were required to compute Signal-to-Error Ratio (SER) and Artifact-to-Residue Ratio (ARR) values. However, other metrics required epoched data (the total proportion of the epochs rejected by cleaning, and the variance explained by the experimental manipulation). To obtain the epoched data, we interpolated rejected electrodes back into the data (with EEGLAB’s ‘pop_interp’ function and the spherical setting) and applied a typical rejection of epochs based on max-min voltage values >60 microvolts, or kurtosis / improbable data for all channels >3 or any channel >5 to remove any remaining artifacts (since no EEG cleaning pipeline completely addresses all artifacts for all files). This enabled an analysis of differences between the pipelines in the variance explained by differences between commonly analysed experimental conditions (such as between working memory delay and probe periods of the Sternberg task), as well as a measure of the proportion of epochs that were removed by the overall cleaning process, including extreme period rejection and outlier epoch rejection. Less effective cleaning pipelines left more artifacts remaining after cleaning, which were rejected by the final epoch rejection step, providing less data available for analysis after cleaning.

### Cleaning Quality Evaluation Metrics

When assessing the performance of an EEG cleaning method, there are three major challenges. Firstly, the ‘ground-truth’ of the brain activity signal in isolation is impossible to ascertain in real data, so assessments of cleaning efficacy and signal preservation can only be estimates. Simulated data has been used to address this issue, but it also difficult to ascertain how accurately simulations reflect real data and artifacts, so effective cleaning in simulated data may not translate to effective cleaning in real data (Kumaravel et al., 2022; Mumtaz et al., 2021; Rošťáková & Rosipal, 2021). As such, we have limited our analysis to real data, and used a combination of many artifact cleaning metrics and signal preservation metrics. We deliberately selected a complimentary combination of metrics, so some metrics address the potential limitations of other included metrics. We have also included metrics that simply aim to detect whether a well-characterised artifact is still present after cleaning (rather than trying to estimate the artifact signal and how much cleaning reduces that signal). For example, one metric tested whether blink periods are still associated with increased frontal voltage amplitudes after cleaning, and another whether epochs still show log-power log-frequency slopes that indicate muscle activity remains after cleaning. This approach avoids the ‘ground-truth’ issue by simply asking “are the well-known artifact characteristics still present in the data after cleaning?”. Secondly, EEG data is contaminated by many different artifact types, all of which should be cleaned effectively for optimal data analysis. To address this, we have included artifact reduction metrics that address all common artifacts. Thirdly, perhaps the most important metrics should assess the impact of EEG data cleaning on the practical outcomes of the research (Clayson, Baldwin, et al., 2021). As such, we have included multiple metrics that assess whether cleaning by the different pipelines leads to more variance explained by different experimental manipulations.

To ensure we assessed the different cleaning pipelines fully for effectiveness at cleaning both the full range of potential artifacts, and for preserving the neural signal (not over-cleaning the data), we tested the pipelines across six different types of cleaning quality metrics, for a total of 13 different metrics. All metrics are explained in detail in the Supplementary Materials (section 3, pages 13-18) and summarised in Table 3. Briefly, these metrics included: the SER: a measure of the amount of signal left unaffected during the clean EEG periods after cleaning. The SER is obtained by dividing the expected value operator of the squared signal amplitude in periods marked as clean in each electrode in the raw EEG by the squared signal of the signal that was removed by cleaning during the periods marked as clean the MWF templates. The individual electrode values are then averaged across electrodes with weighting applied by the artifact signal in each electrode proportional to the artifact signal in all electrodes, so electrodes with larger artifact contribute more to the SER measure. Secondly, we included a measure of the extent to which all artifacts identified by our MWF template were reduced – the ARR. The ARR was calculated by obtaining the expected value operator of the square of the removed artifact, divided by the expected value operator of the square of the raw data from the artifact periods minus the removed artifact signal from the artifact periods. Effective cleaning reflects reductions in large artifact signals, such that the denominator is small and high ARR values result from the division. As such, high values for the SER and ARR indicate good performance (Bertrand, 2015; Somers & Bertrand, 2016; Somers et al., 2018). It is important to note that SER and ARR are complementary metrics. To achieve the highest cleaning quality a pipeline should produce both high SER and high ARR values; it would be easy to obtain very high ARR values and very low SER values by being excessively stringent in artifact reduction and not concerned with preserving the neural signal, but this would reduce the neural activity of interest.

**Table 3.**
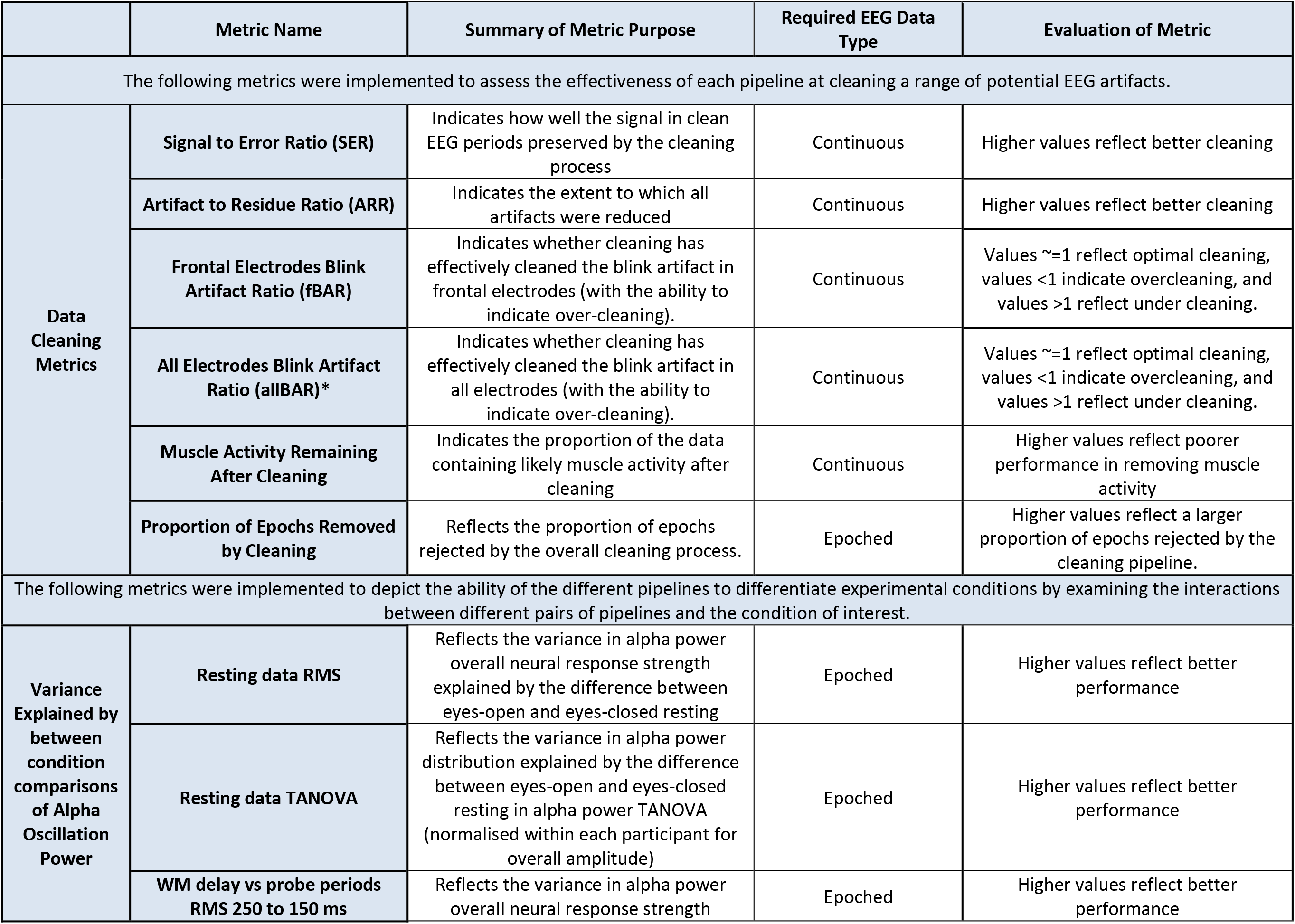
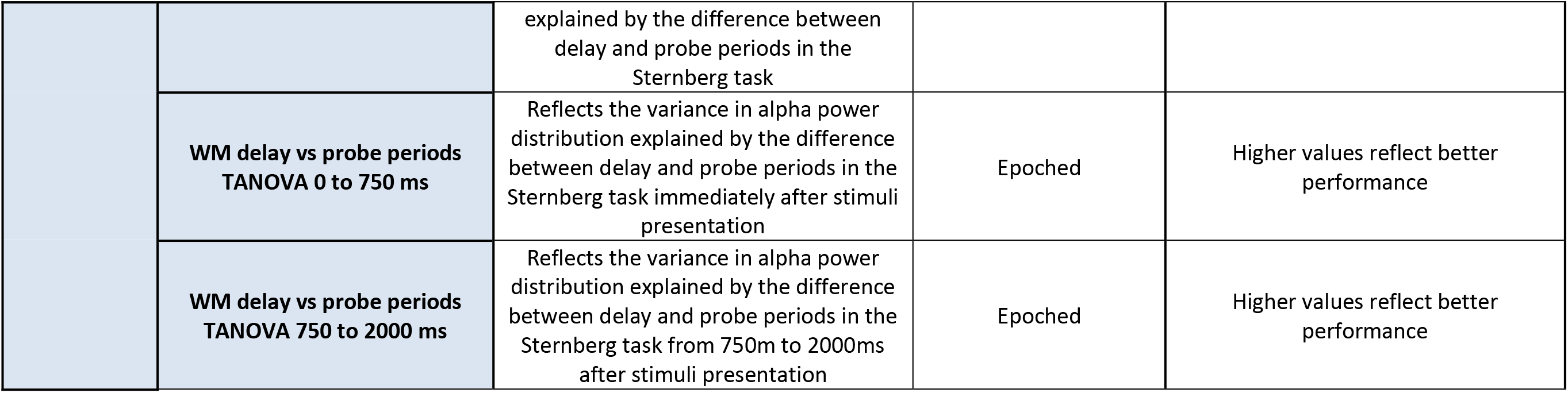
Description of cleaning quality evaluation metrics used for comparing pipelines. Not all results from the pipelines are included in the main manuscript but can be viewed in the Supplementary Materials. Metrics denoted with an asterisk are only presented in the Supplementary Materials. RMS: Root Mean Square, TANOVA: topographical analysis of variance.

|  | Metric Name | Summary of Metric Purpose | Required EEG Data Type | Evaluation of Metric |
| --- | --- | --- | --- | --- |
| The following metrics were implemented to assess the effectiveness of each pipeline at cleaning a range of potential EEG artifacts. |  |  |  |  |
| Data Cleaning Metrics | Signal to Error Ratio (SER) | Indicates how well the signal in clean EEG periods preserved by the cleaning process | Continuous | Higher values reflect better cleaning |
|  | Artifact to Residue Ratio (ARR) | Indicates the extent to which all artifacts were reduced | Continuous | Higher values reflect better cleaning |
| | Frontal Electrodes Blink Artifact Ratio (fBAR) | Indicates whether cleaning has effectively cleaned the blink artifact in frontal electrodes (with the ability to indicate over-cleaning). | Continuous | Values $\sim 1$ reflect optimal cleaning, values $< 1$ indicate overcleaning, and values $> 1$ reflect under cleaning. |
| | All Electrodes Blink Artifact Ratio (allBAR)* | Indicates whether cleaning has effectively cleaned the blink artifact in all electrodes (with the ability to indicate over-cleaning). | Continuous | Values $\sim 1$ reflect optimal cleaning, values $< 1$ indicate overcleaning, and values $> 1$ reflect under cleaning. |
|  | Muscle Activity Remaining After Cleaning | Indicates the proportion of the data containing likely muscle activity after cleaning | Continuous | Higher values reflect poorer performance in removing muscle activity |
|  | Proportion of Epochs Removed by Cleaning | Reflects the proportion of epochs rejected by the overall cleaning process. | Epoched | Higher values reflect a larger proportion of epochs rejected by the cleaning pipeline. |
| The following metrics were implemented to depict the ability of the different pipelines to differentiate experimental conditions by examining the interactions between different pairs of pipelines and the condition of interest. |  |  |  |  |
| Variance Explained by between condition comparisons of Alpha Oscillation Power | Resting data RMS | Reflects the variance in alpha power overall neural response strength explained by the difference between eyes-open and eyes-closed resting | Epoched | Higher values reflect better performance |
|  | Resting data TANOVA | Reflects the variance in alpha power distribution explained by the difference between eyes-open and eyes-closed resting in alpha power TANOVA (normalised within each participant for overall amplitude) | Epoched | Higher values reflect better performance |
|  | WM delay vs probe periods RMS 250 to 150 ms | Reflects the variance in alpha power overall neural response strength explained by the difference between delay and probe periods in the Sternberg task | Epoched | Higher values reflect better performance |
|  | **WM delay vs probe periods**<br>**TANOVA 0 to 750 ms** | Reflects the variance in alpha power distribution explained by the difference between delay and probe periods in the Sternberg task immediately after stimuli presentation | Epoch | Higher values reflect better performance |
|  | **WM delay vs probe periods**<br>**TANOVA 750 to 2000 ms** | Reflects the variance in alpha power distribution explained by the difference between delay and probe periods in the Sternberg task from 750m to 2000ms after stimuli presentation | Epoch | Higher values reflect better performance |

Additionally, we measured the ratio of blink amplitude to non-blink periods after cleaning, referred to as the Blink Amplitude Ratio (BAR). We examined BAR averaged across the frontal electrodes affected by blinks (fBAR), and across all electrodes (allBAR, reported in the Supplementary Materials, section 4 page 25). For fBAR and allBAR, values near to 1 indicate optimal performance, with activity that is time-locked to blinks showing the same amplitude as non-blink activity after cleaning, while values <1 reflect overcleaning, and values >1 reflect under cleaning (Robbins et al., 2020). Next, we measured the number of epochs showing log-power log-frequency slopes >-0.59 that indicated the epoch was still contaminated by muscle activity (Fitzgibbon et al., 2016), and the severity by which these slopes exceeded the muscle threshold within epochs showing muscle activity remaining after cleaning (these results are reported in the Supplementary Materials, section 4, page 30). Higher values reflect poorer performance for these metrics. The final cleaning metric we assessed was the proportion of epochs that were rejected through the cleaning process against the initial number of epochs in the raw data. While this measure can be affected by the raw data quality (with lower raw data quality requiring more data rejection), comparisons of the number of epochs rejected by different cleaning pipelines (applied to the same datasets) indicate which pipelines are better at cleaning data (such that fewer bad epochs need to be removed, and more epochs are available for analysis). Providing higher quality data with more epochs for analysis can be more important for statistical power than recruiting more participants (Kolossa & Kopp, 2018). As such, lower values for the proportion of epochs rejected reflect better performance.

Next, and perhaps most importantly, we assessed the amount of variance explained by a variety of experimental manipulations after cleaning by each pipeline. We chose experimental manipulations that are well established to provide differentiation of neural activity in the comparison of two conditions, which assesses the real-world applicability of the cleaning pipelines (Clayson, Baldwin, et al., 2021). We assessed the variance explained by a comparison of alpha oscillatory power between the working memory delay and probe periods of the Sternberg task (Bailey et al., 2020) and alpha power between EO and EC resting-state data (computation of which is described in the Supplementary Materials, section 4, pages 36-53). Within these metrics, higher values are likely to reflect better performance, with cleaning pipelines that provided higher values providing a better chance to detect statistically significant differences within an experiment.

### Statistics

In order to compare the effectiveness of artifact cleaning between the cleaning pipelines across each of the metrics, we used the robust repeated measures ANOVA function “rmanova” from the WRS2 package in R (R Core Team v4.0.4) (Mair & Wilcox, 2020). These tests are robust against the normality and homoscedasticity assumptions of traditional parametric statistical tests, while still providing equivalent power (Mair & Wilcox, 2020). When the overall ANOVA was significant, pairwise comparisons of the cleaning pipelines were performed using the robust post-hoc t-test function “rmmcp”. This applies multiple comparison controls across the post-hoc tests within each omnibus ANOVA using Hochberg’s approach to control for the family-wise error (Mair & Wilcox, 2020). However, experiment-wise multiple comparison controls were not implemented as we deemed it was more important to provide sensitivity to differences in cleaning outcomes than to protect against false positive results when exploring a wide range of EEG cleaning methods for the optimal approach (Bender & Lange, 2001). Additionally, we tested a large number of metrics to demonstrate which pipelines provide the best overall performance (rather than for a single artifact type). We also used multiple datasets to provide an indication of how replicable our results are across datasets. Experiment-wise multiple comparison controls in this large number of comparisons across this wide range of metrics and across multiple datasets would severely reduce the sensitivity of our study to detect important differences between cleaning approaches. It would have also adversely encouraged us to reduce the number of metrics we assessed, leading to a less well tested final pipeline. As such, we emphasised sensitivity to detect meaningful differences between pipelines over the potential for false positives. To visualise the cleaning efficacy data, we provide raincloud plots with boxplot medians and IQR marked for the reader to see the full data with minimal distortion (Allen et al., 2019). This includes outliers, which are important to note, as outliers indicate individual files in which a cleaning method might not have adequately cleaned the artifact.

In addition to the robust statistics used to compare artifact cleaning metrics, we have provided graphs of the amount of variance explained (np^2^) by the comparison between the two experimental conditions for each pipeline for the comparisons of the amount of variance explained by the experimental manipulation (as well as the absolute values for each condition from each pipeline in the Supplementary Materials, section 4, pages 36-55). To test for differences between pipelines in np^2^ for the experimental manipulation, we tested the interaction between each pair of pipelines and the two experimental conditions using the Randomisation Graphical User Interface (RAGU) Global Field Potential (GFP) test which measures overall neural response amplitudes. We also used the Topographical Analysis of Variance (TANOVA) with L2 norm tests, which measures the distribution of neural activity after normalisation for amplitude (Habermann et al., 2018; Koenig et al., 2011). We have provided heat maps displaying np^2^ values for each interaction in the Supplementary Materials (section 4, pages 36-55), with the multiple comparison controls applied across the post-hoc tests within each omnibus ANOVA for each measure using the false discovery rate (FDR-p) (Benjamini & Hochberg, 1995).

We also provide a rank order of the means from each pipeline for each cleaning efficacy and variance explained metric, with significant differences noted (Table 4). We have also provided means and SD tables, and specific details of post-hoc tests within a heat map, including confidence intervals for t-test comparisons with significant tests highlighted for all metrics in the Supplementary Materials (section 4). Finally, we have provided scatter plots of mean SER x ARR values for each pipeline across the different datasets, so that these two metrics can be considered together for each cleaning pipeline.

**Table 4.**
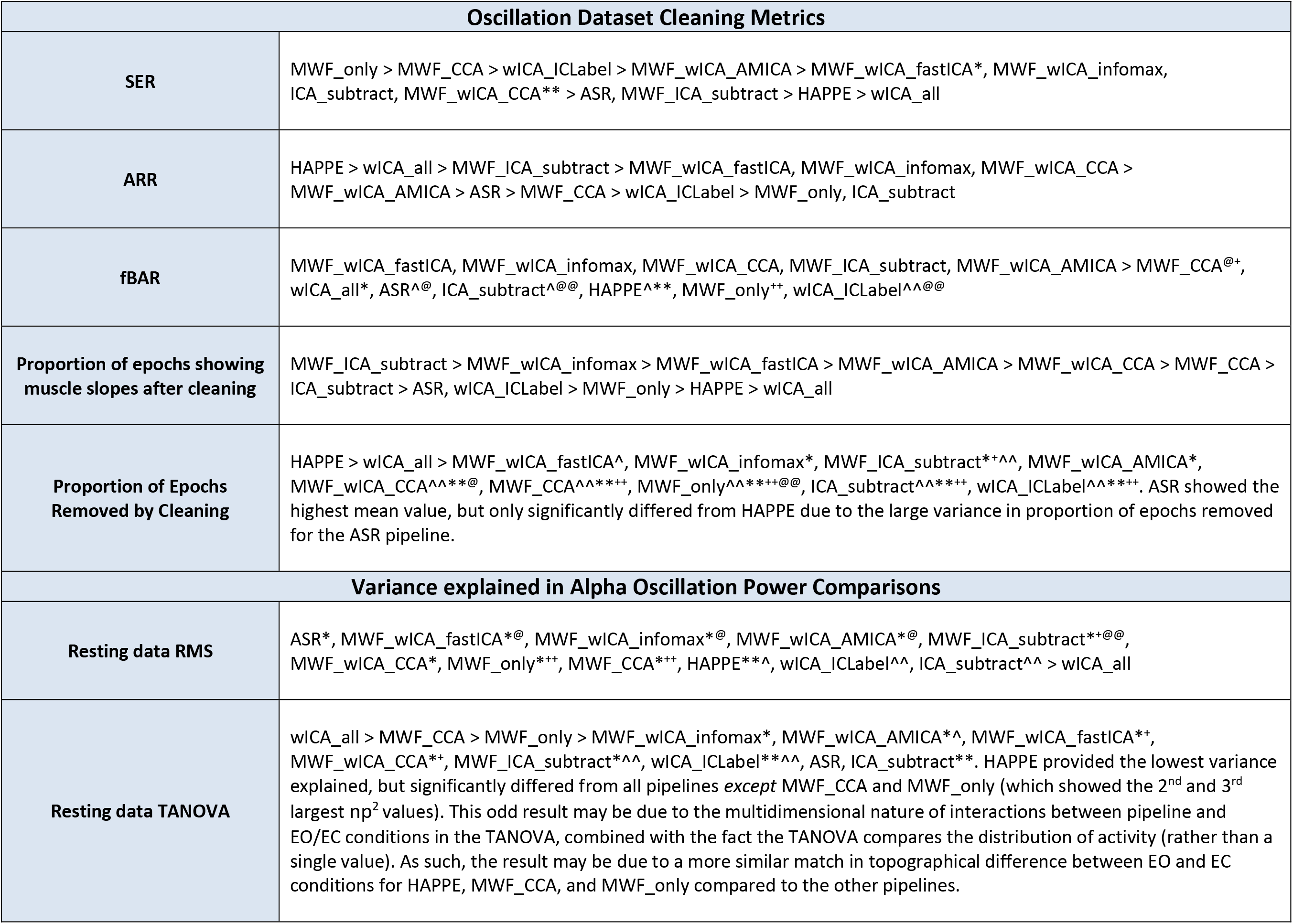
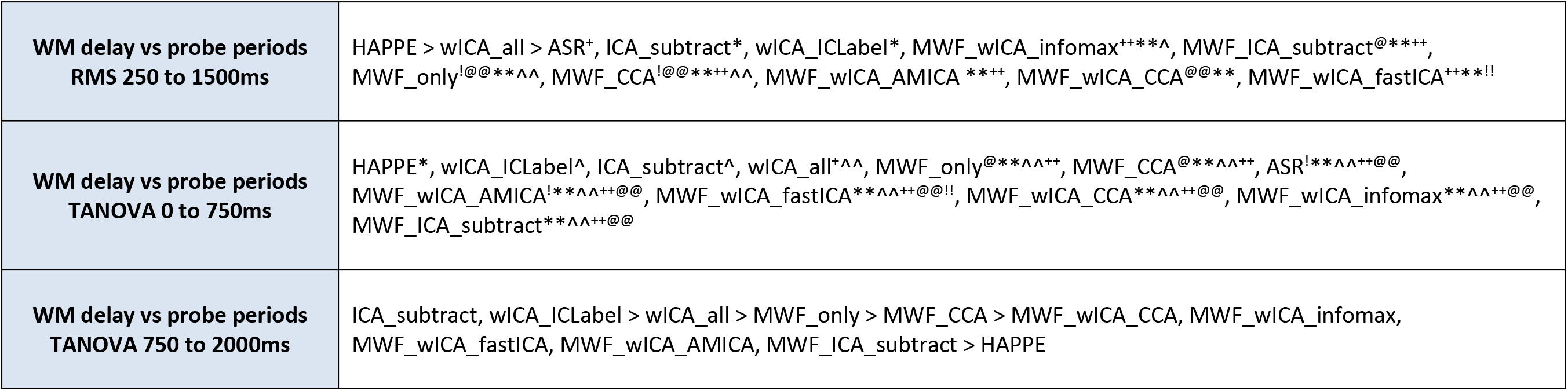
Rank order from best performance (on the left) to worst performance (on the right) (by mean) for the oscillatory analysis datasets. Note that for some metrics, better performance is reflected by lower values, for example the proportion of epochs showing muscle activity remaining after cleaning, and these metrics are ordered from best performance (on the right) to worst performance (on the left) rather than from larger to smaller values. Significant differences are highlighted for which pipelines performed significantly better than other pipelines using the following notation for ease of understanding: better performance > worse performance (rather than higher values > lower values). Because sometimes pipeline 1 differed from pipeline 2, but pipeline 3 did not differ from either 1 or 2, we have used the following notation: ^ = significantly higher than the pipeline marked with a ^^ within the same section (while the others in the category are not significantly different from each other). * = significantly higher than the pipeline marked with a ** in the same category, and so on for the following symbols: ^+@$!+^.

## Results

All metrics we tested showed a significant difference in the omnibus ANOVA (all p < 0.01). A rank order of the means from each pipeline (with significant differences as detected by the post-hoc t-tests noted) can be viewed in Table 4. Due to the high number of comparisons, for readability, we provide here a summary of only the results that were most relevant to the evaluation of the pipelines for cleaning efficacy and the detection of differences between well-established experimental conditions. Full details are provided in the Supplementary Materials (section 4, pages 20-53). In addition to the results reported below, we also note that RELAX is quick to run - performing MWF_wICA using fastICA without computing metrics took 225 seconds, on a 60 electrode 7-minute EEG file, recorded at 1000Hz using an Intel Core i7-10875H CPU @ 2.3GHz, 32GB RAM. Performing wICA_ICLabel without metrics only took 140 seconds. Computing metrics only added 25 seconds to the computation time.

### Artifact Cleaning Metrics

#### Signal-to-Error Ratio and Artifact-to-Residue Ratio

SER and ARR values can be viewed in Figure 2. When SER and ARR values were viewed together (Figure 3), it was apparent that MWF_wICA and MWF_wICA_CCA performed better than ASR in both metrics. MWF_wICA and MWF_wICA_CCA also performed equally to ICA_subtract in SER, while at the same time performing better in the ARR metric. MWF_ICA_subtract provided higher ARR values but at the expense of lower SER values than MWF_wICA. MWF_ICA_subtract also provided similar SER and higher ARR values compared to ASR suggesting that MWF cleaning is superior to ASR as an initial data cleaning step prior to using ICA to subtract artifacts. HAPPE and wICA_all showed very high ARR values, but at the expense of very low SER values. Inversely, MWF_only, MWF_CCA, and wICA_ICLabel showed high SER values but lower ARR values (MWF_only performed the best of these three pipelines, providing high SER and a similar value for ARR). Finally, note that wICA_ICLabel outperformed ICA_subtract when viewing SER and ARR concurrently. While these initial results might suggest MWF_only is the best pipeline, the SER and ARR values should be viewed in the context of the other metrics we have included, which indicate how much blink and muscle artifacts still affect the data after cleaning, as well as the variance explained by the experimental manipulations.

**Figure 2.**
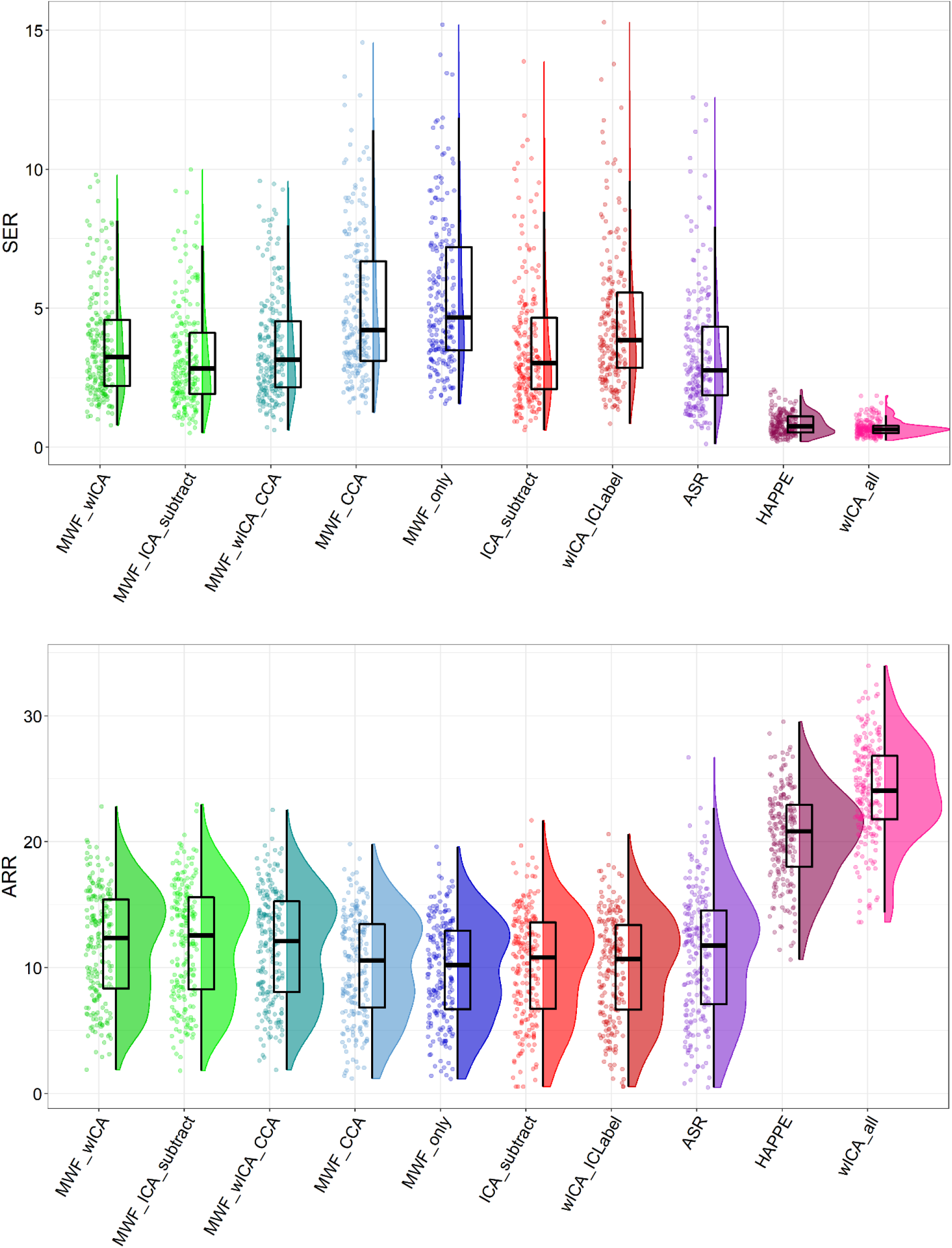
Raincloud plots depicting Signal-to-Error (SER) and Artifact-to-Residue (ARR) values from the combined EO, EC, and Sternberg data (N = 213) for each of the cleaning pipelines.

**Figure 3.**
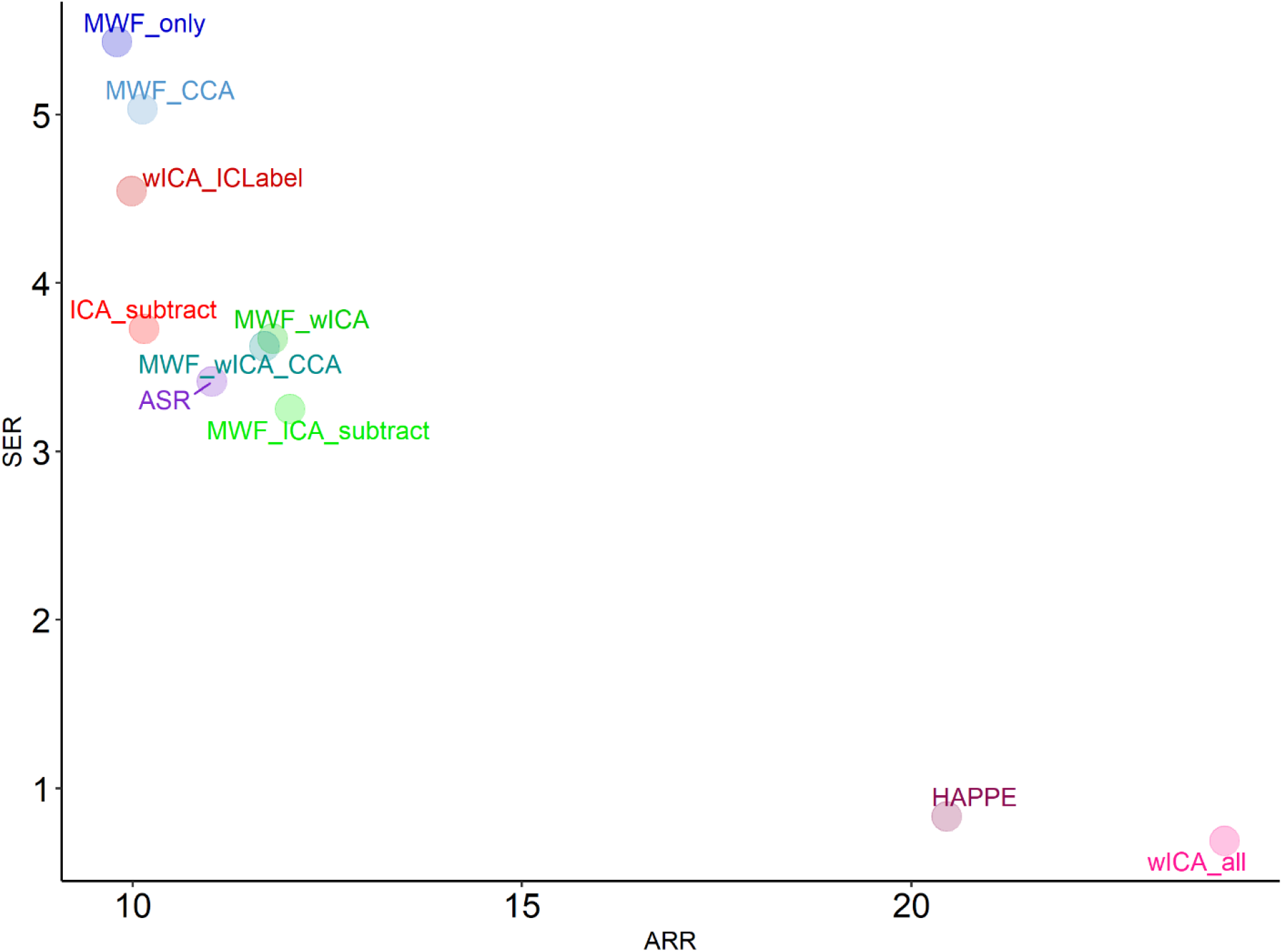
A scatterplot depicting both mean Signal-to-Error (SER) and mean Artifact-to-Residue (ARR) values for the resting-state EO, EC, and Sternberg dataset from each cleaning pipeline.

#### Blink Amplitude Ratios

MWF_wICA and MWF_wICA_CCA performed significantly better than all other pipelines (while not differing from each other), providing values closest to 1 for both fBAR (Figure 4) and allBAR (reported in the Supplementary Materials, section 3, page 29). This was with the exception of MWF_ICA_subtract and MWF_CCA, which only performed significantly worse than MWF_wICA and MWF_wICA_CCA for allBAR values (and did not significantly differ in fBAR values). HAPPE, MWF_only, and wICA_ICLabel were the worst performers for fBAR, and MWF_only, ASR, ICA_subtract and wICA_ICLabel were the worst performers for all_BAR. However, all pipelines performed reasonably well with mean fBAR and allBAR values ranging from 1.015 to 1.077. Having said this, it is important to note the presence of outliers, which indicated where a pipeline did not clean blinks effectively for a specific EEG file. Visual inspection of the raincloud plot indicated MWF_wICA and MWF_ICA_subtract showed the fewest outliers.

**Figure 4.**
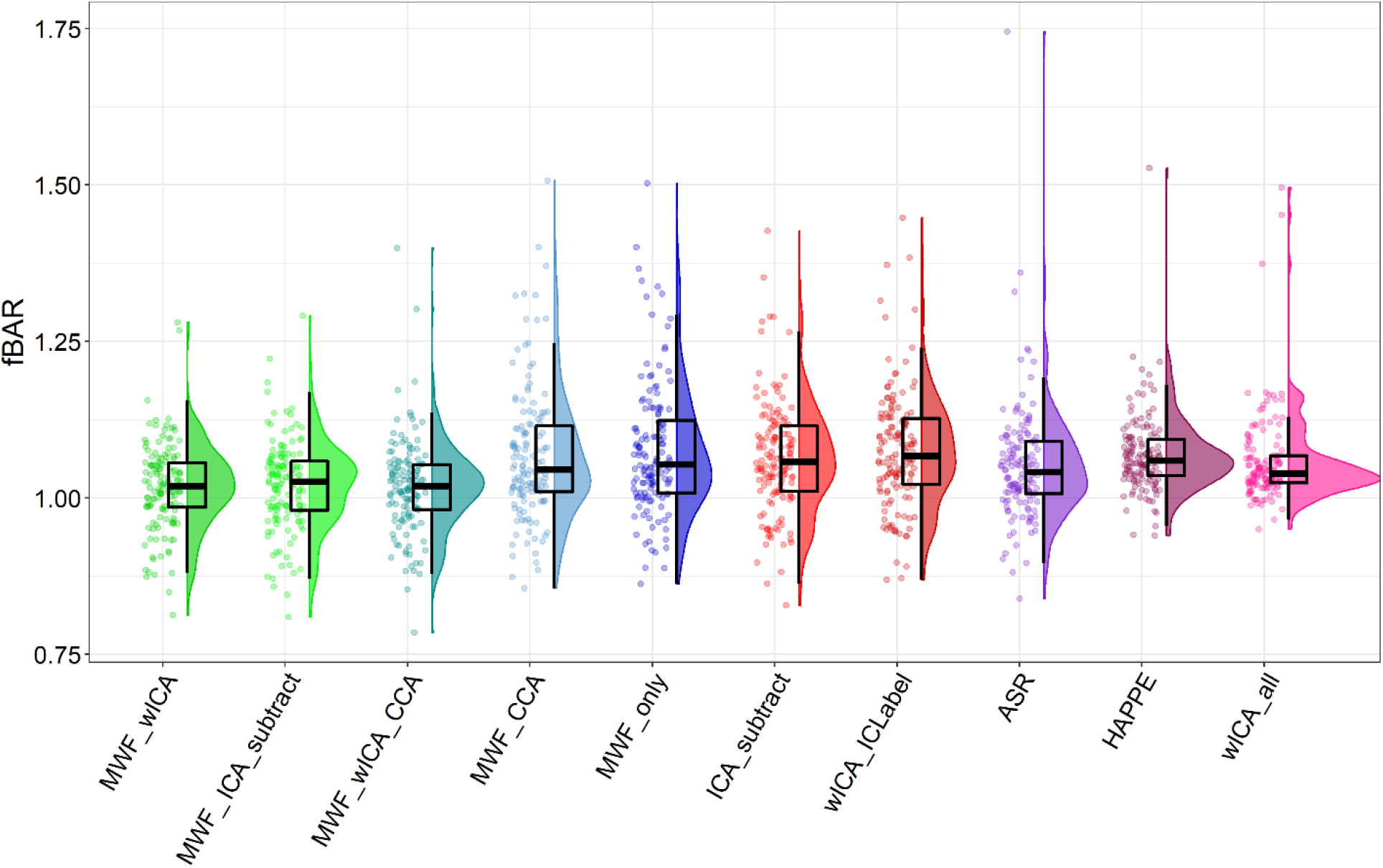
Raincloud plot depicting fBAR values from the combined EO, and Sternberg data (N = 140) for each of the cleaning pipelines.

#### Muscle Activity Remaining After Cleaning

MWF_ICA_subtract outperformed all other pipelines in terms of the proportion of epochs showing muscle activity remaining after cleaning (essentially 0 for almost all EEG files, Figure 5). MWF_wICA performed the next best (with values of almost 0 for most EEG files, but some outlying files showed more epochs contaminated by muscle activity), followed by MWF_wICA_CCA, then MWF_CCA then ICA_subtract. ASR and wICA_ICLabel performed worse, followed by MWF_only. This overall rank order was similar for the severity of muscle slopes that exceeded the threshold after cleaning, with MWF_ICA_subtract performing the best, followed by MWF_wICA (reported in the Supplementary Materials, section 4, page 31). Of note, both the HAPPE and wICA_all pipelines showed considerably more epochs identified as containing residual muscle activity after cleaning than other pipelines (>75% of epochs compared to <10% for other pipelines). Instead of reflecting genuine residual muscle activity, this may reflect an overall flattening of the spectra slope due to considerable removal of power in the lower frequencies by these pipelines (producing slope values more like muscle affected log-power log-frequency slopes). However, note that these pipelines showed relatively more beta range power in the power-frequency plot, a feature which is not present in the other pipelines (Figures 7 and 9).

**Figure 5.**
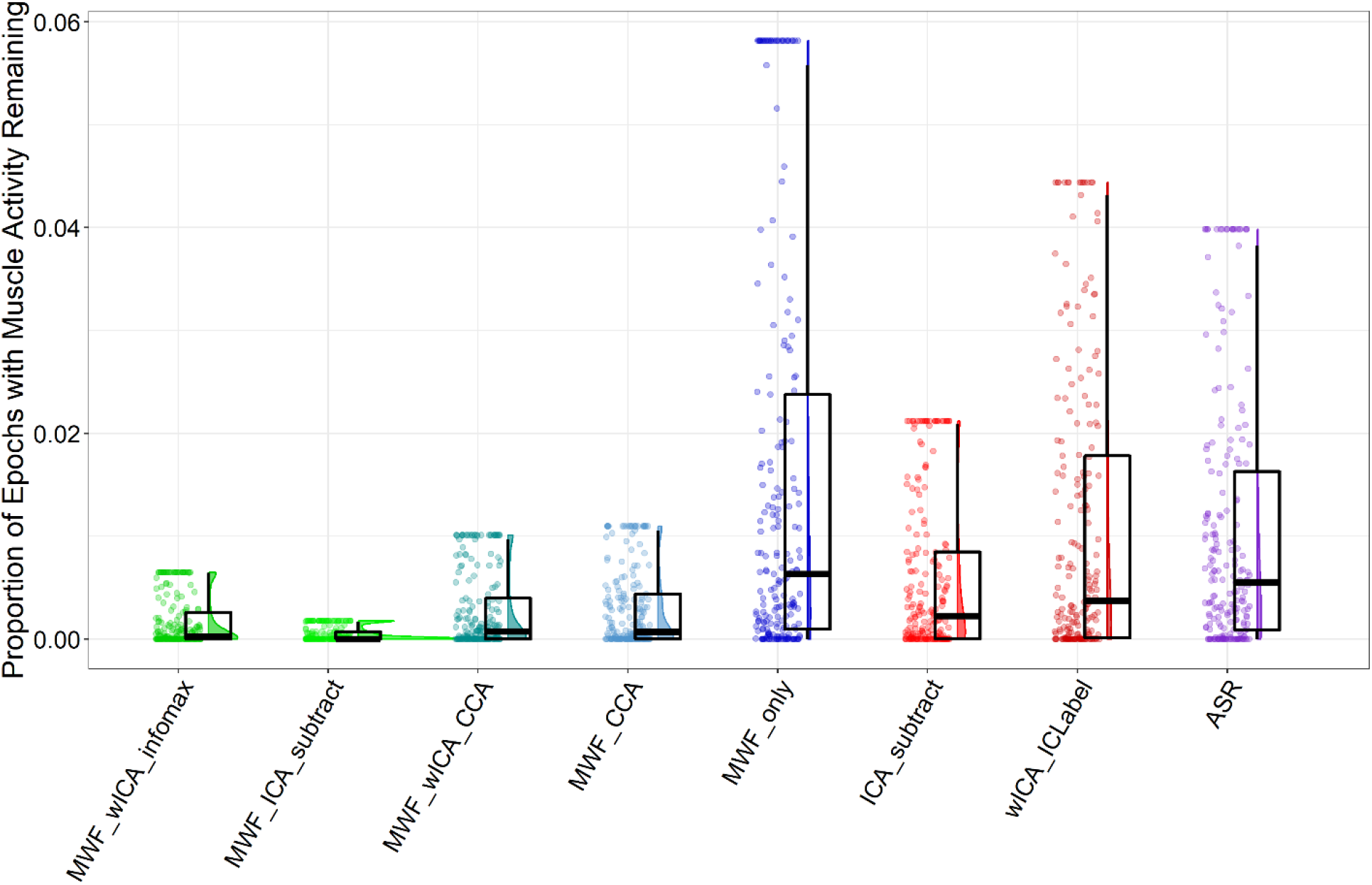
Raincloud plot depicting the proportion of epochs showing log-power log-frequency values above the -0.59 threshold from the combined EO, EC, and Sternberg data (N = 213) for each of the cleaning pipelines. Note that this figure excludes HAPPE and wICA_all, as these pipelines showed median values > 0.75 and made the scale of the graph such that it was difficult to visualise differences in the other pipelines. Note also that we have winsorized the data in the figure, as the outliers also made the scale such that it was difficult to visualise differences in the other pipelines. The full data can be viewed in the Supplementary Materials Figure S11 (page 30).

**Figure 6.**
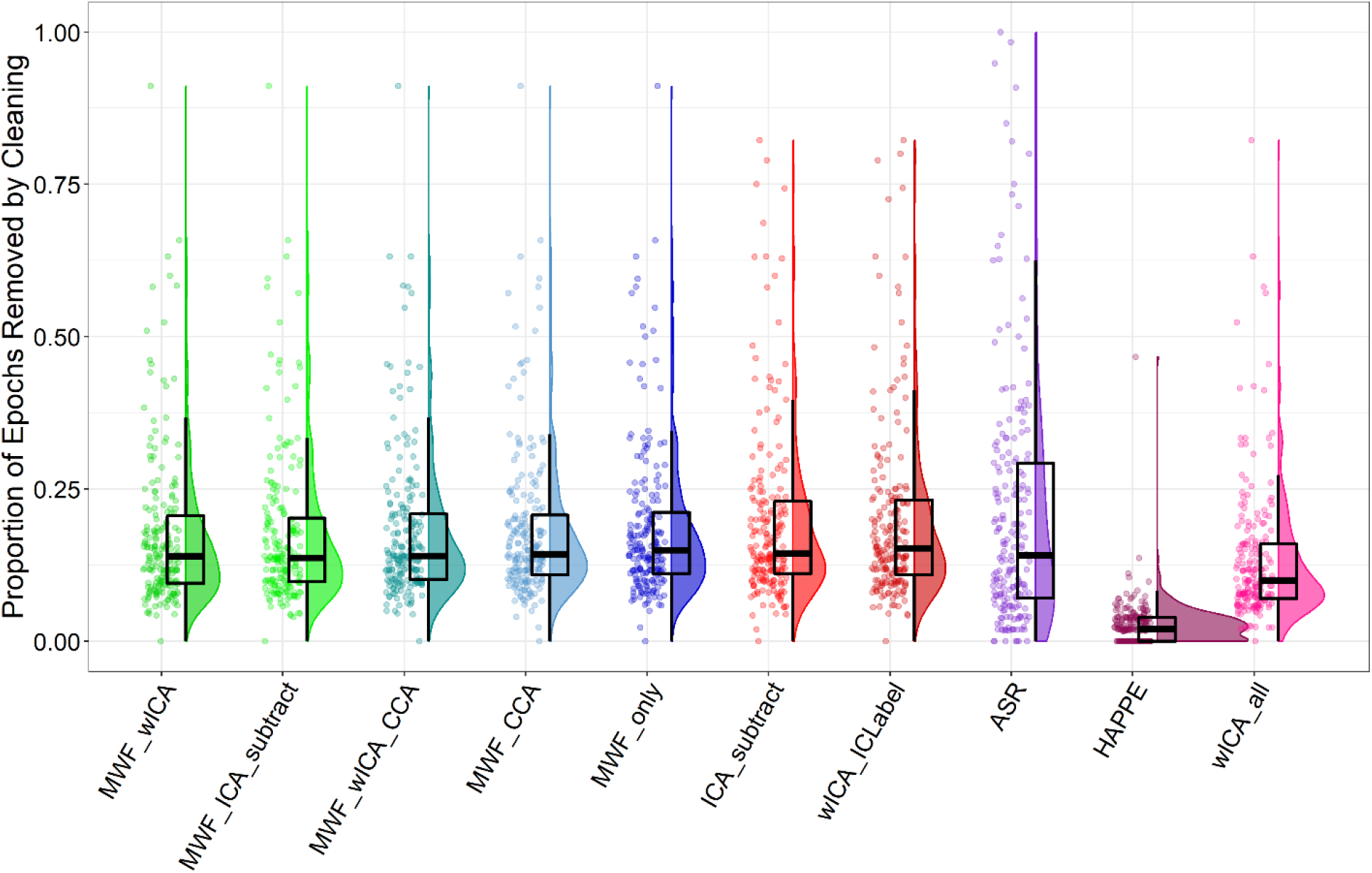
Raincloud plot depicting the proportion of epochs in the data rejected from the combined EO, EC, and Sternberg data (N = 213) for each of the cleaning pipelines.

**Figure 7.**
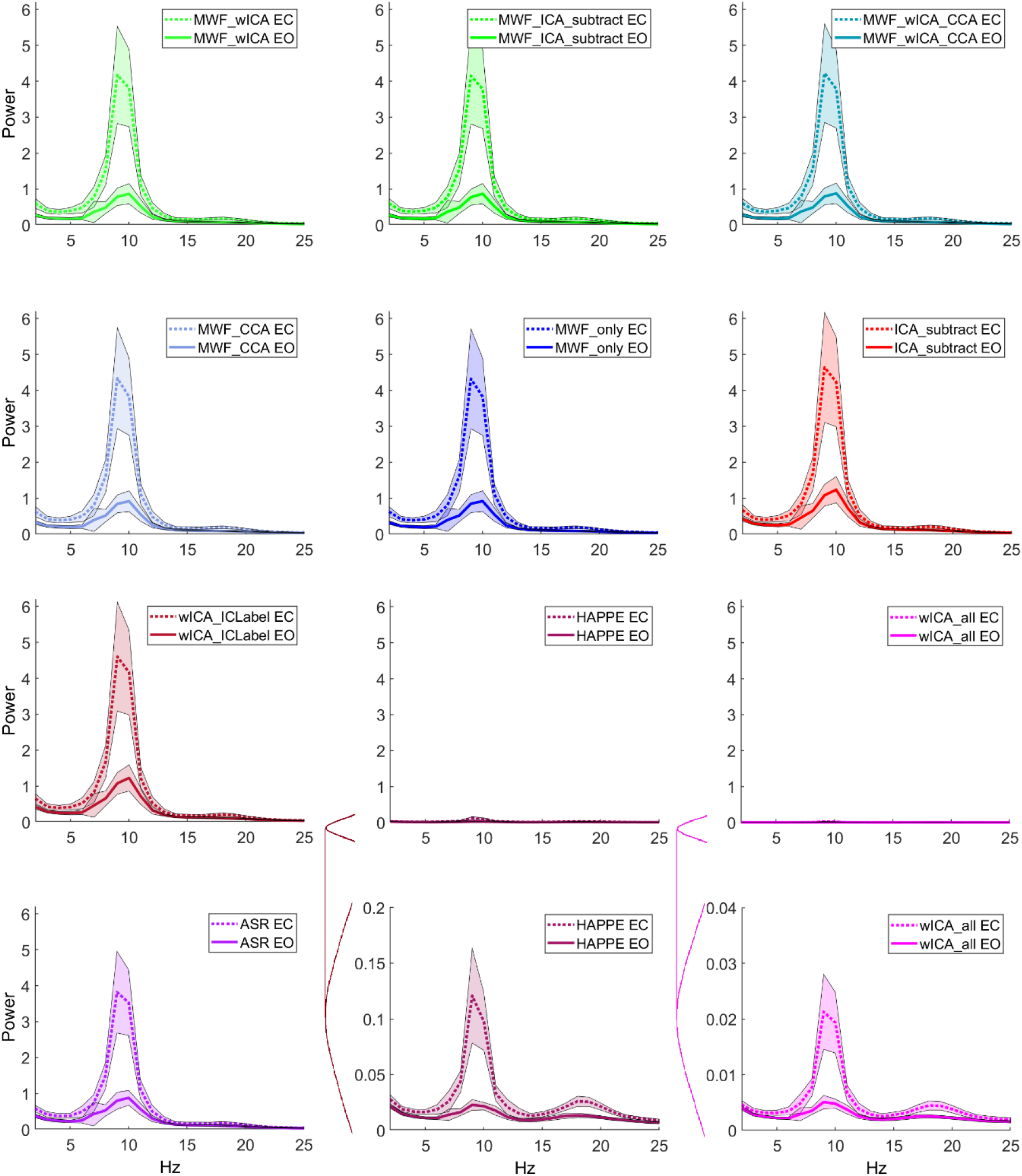
Frequency power plots depicting fast-Fourier transformed power averaged across PO7 and PO8 at each frequency from the EO and EC resting-state EEG recordings for each cleaning pipeline (shaded errors reflect 95% confidence intervals). Note the different scale required to see the power spectrum for HAPPE and wICA_all.

**Figure 8.**
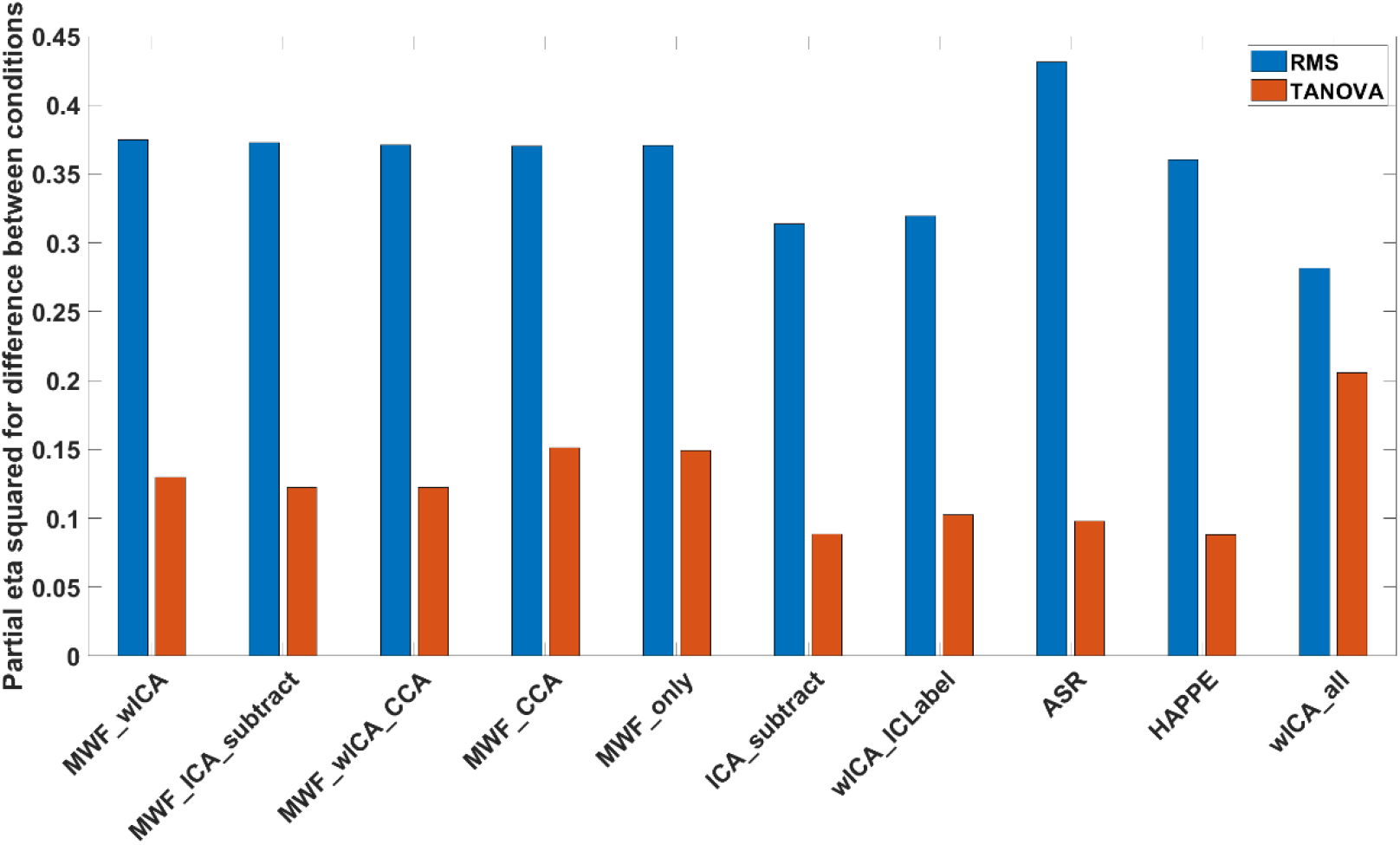
The variance explained (np^2^) by differences in averaged alpha power between eyes open and eyes closed for RMS and TANOVA tests for each of the cleaning pipelines.

**Figure 9.**
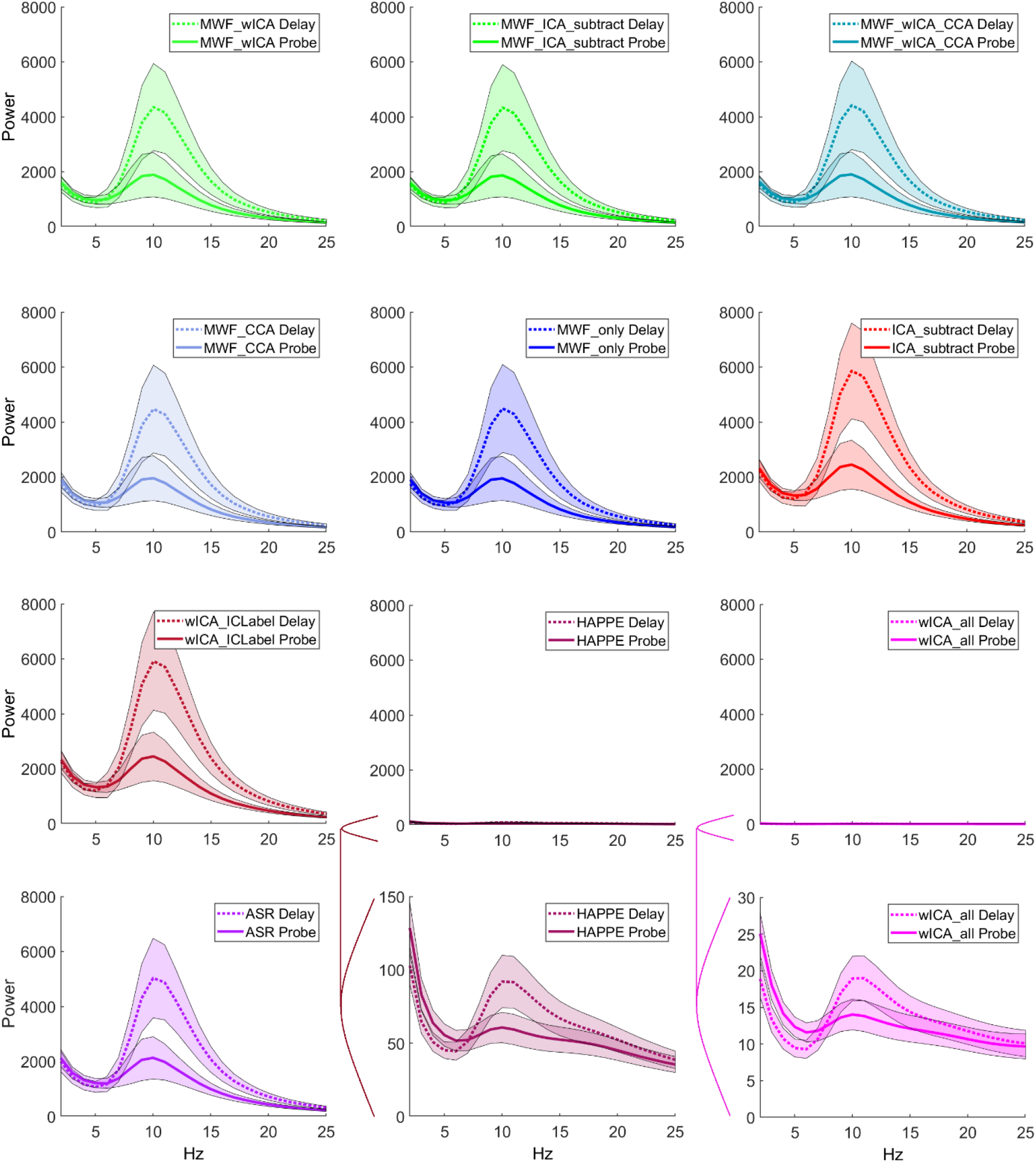
Frequency power plots depicting Morlet wavelet transformed power averaged across PO7 and PO8 at each frequency from the WM delay and probe periods for each cleaning pipeline (shaded errors reflect 95% confidence intervals). Note the different scale required to see the power spectrum for HAPPE and wICA_all.

#### Proportion of Epochs Removed by All Cleaning Steps

HAPPE and wICA produced data that had the fewest epochs removed by cleaning (Figure 6). These pipelines also produced very low amplitude cleaned data, so we suspect their high performance in this metric is the result of producing cleaned data with no epochs that exceeded the epoch rejection criteria in our final epoch rejection cleaning step. MWF_wICA then MWF_ICA_subtract were the next best performers, followed by MWF_wICA_CCA, MWF_CCA, MWF_only, ICA_subtract and wICA_ICLabel. ASR showed the highest mean value (worst performance), but only significantly differed from HAPPE (we suspect this was due to its high variance which can be seen in Figure 6). When we tested for post-hoc differences between individual pipelines using a bootstrap pairwise t-test (pairdepb in the WRS2 package), ASR showed a higher proportion of epochs rejected than all other pipelines.

### Variance Explained by Experimental Manipulations

#### Variance Explained by the Difference Between Eyes-Open and Eyes-Closed Resting

Figure 7 depicts frequency-power spectrum plots for EO and EC resting, and Figure 8 depicts the amount of variance explained by the difference between EO and EC resting-state in alpha power root mean square (RMS; reflecting overall neural response strength) and TANOVA (distribution of neural activity) tests across the different cleaning pipelines. When comparing the difference in EEG signals between resting-state conditions, all pipelines showed the expected pattern of higher alpha-band power during EC compared to EO. For measures of the variance explained by the difference between EO and EC resting-states in the global strength of alpha power across all electrodes (RMS), ASR showed the highest np^2^ value, but did not significantly differ from MWF_wICA, MWF_ICA_subtract, MWF_wICA_CCA, wICA_ICLabel, or ICA_subtract. MWF_wICA also outperformed MWF_ICA_subtract, HAPPE and wICA_all.

With regards to the distribution of activity (TANOVA), wICA_all showed the highest np^2^ value, followed by MWF_CCA, MWF_only then MWF_wICA. ASR, wICA_ICLabel, ICA_subtract, and HAPPE showed the lowest np^2^ values, with the later three of these pipelines showing significant differences compared to all other pipelines, with the unusual exception of HAPPE, which significantly differed from all pipelines except MWF_CCA and MWF_only (pipelines that showed the 2nd and 3rd largest np^2^ values). Interpreting interactions in the normalised distribution of activity is complicated, but we suspect this finding might be due to a more similar pattern of differences in the distribution of activity between the EO and EC activity for HAPPE, MWF_CCA and MWF_only (while other pipelines may have showed a different pattern to HAPPE, and more variance explained than HAPPE but less than MWF_only and MWF_CCA). The distribution of activity from each pipeline can be viewed in the Supplementary Materials (Figure S22-23, pages 41-42).

#### Variance Explained by the Experimental Manipulation in the Sternberg Task

Figure 9 depicts the power-frequency spectrum plots for the Sternberg delay and probe periods. Figure 10 depicts the amount of variance explained by the difference between the Sternberg delay and probe periods in alpha power RMS test across the different cleaning pipelines (which compared overall neural response strength from 250-1500ms after the stimuli) and the TANOVA from 0 to 750ms and 750 to 2000ms after the stimuli. All pipelines showed higher alpha power during the Sternberg delay period than the probe period, as expected. With regards to the alpha power RMS, HAPPE showed the highest np^2^ value for the difference between the two conditions, followed by wICA_all. ASR, ICA_subtract, and wICA_ICLabel showed the next highest, significantly larger than MWF_wICA, MWF_ICA_subtract, MWF_only, MWF_CCA, and MWF_wICA_CCA. While the HAPPE and wICA methods provided the largest np^2^ values, these two pipelines resulted in alpha power values >2 orders of magnitude lower than the other pipelines. This aligns with HAPPE and wICA_all showing the lowest SER values, suggesting much of the neural activity signal was eliminated by these methods. wICA_all and HAPPE also produced a different topography of alpha activity to the other pipelines, with smaller differences in alpha power between Sternberg retention and probe conditions at PO7/8 electrodes, where differences are usually the largest (see Figures 9, 11 and 12 and Supplementary Materials Figures S31-32, pages 41-52). As such, we suspect the application of wICA to all components in these methods may remove characteristics of the EEG activity, so that although these methods might provide more power to detect differences between experimental conditions in some circumstances, they may do so by only providing a partial characterization of the neural activity recorded to the EEG data.

**Figure 10.**
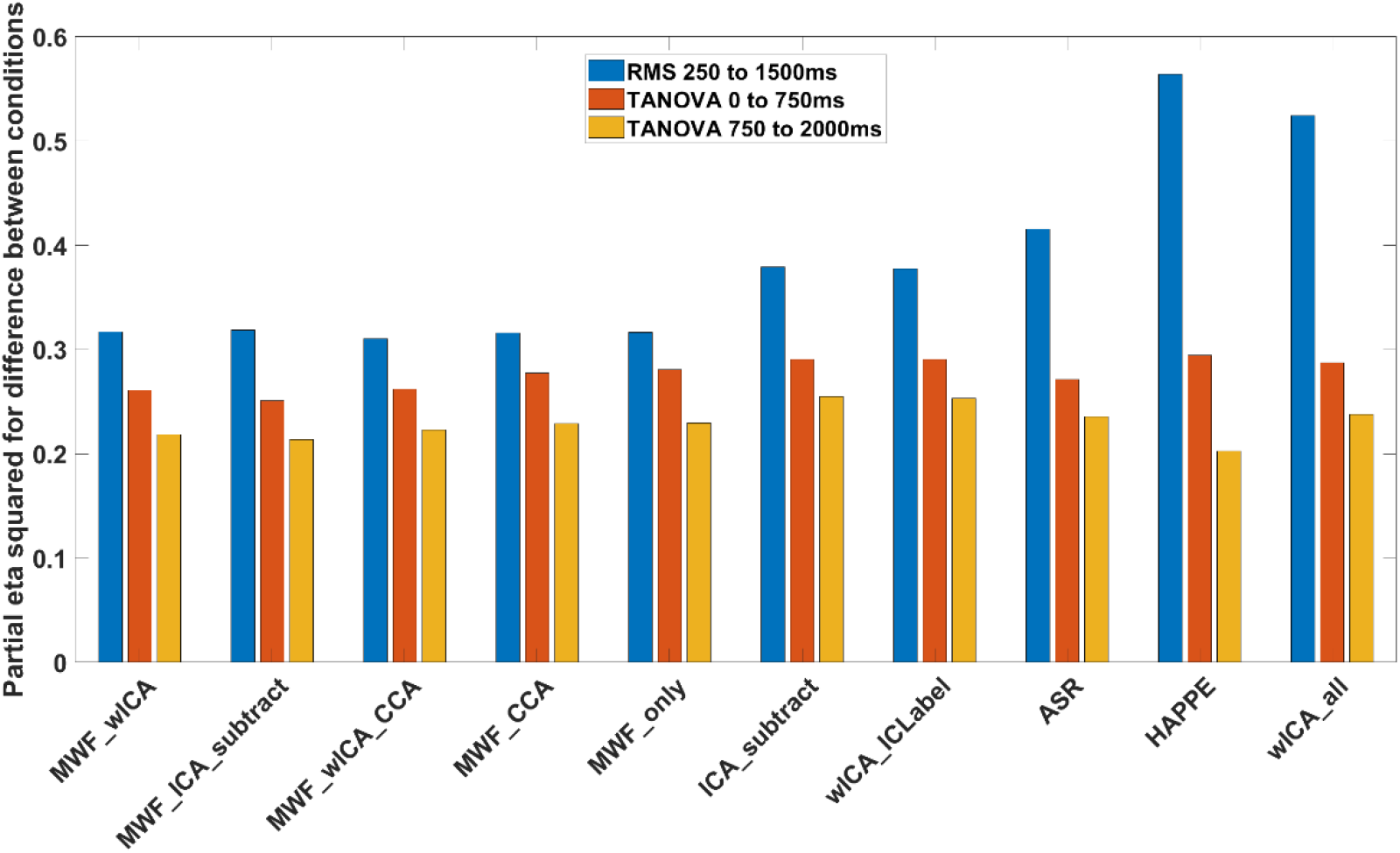
The variance explained (np^2^) by the difference between alpha activity during the delay and probe periods of the Sternberg task after cleaning by each pipeline.

**Figure 11.**
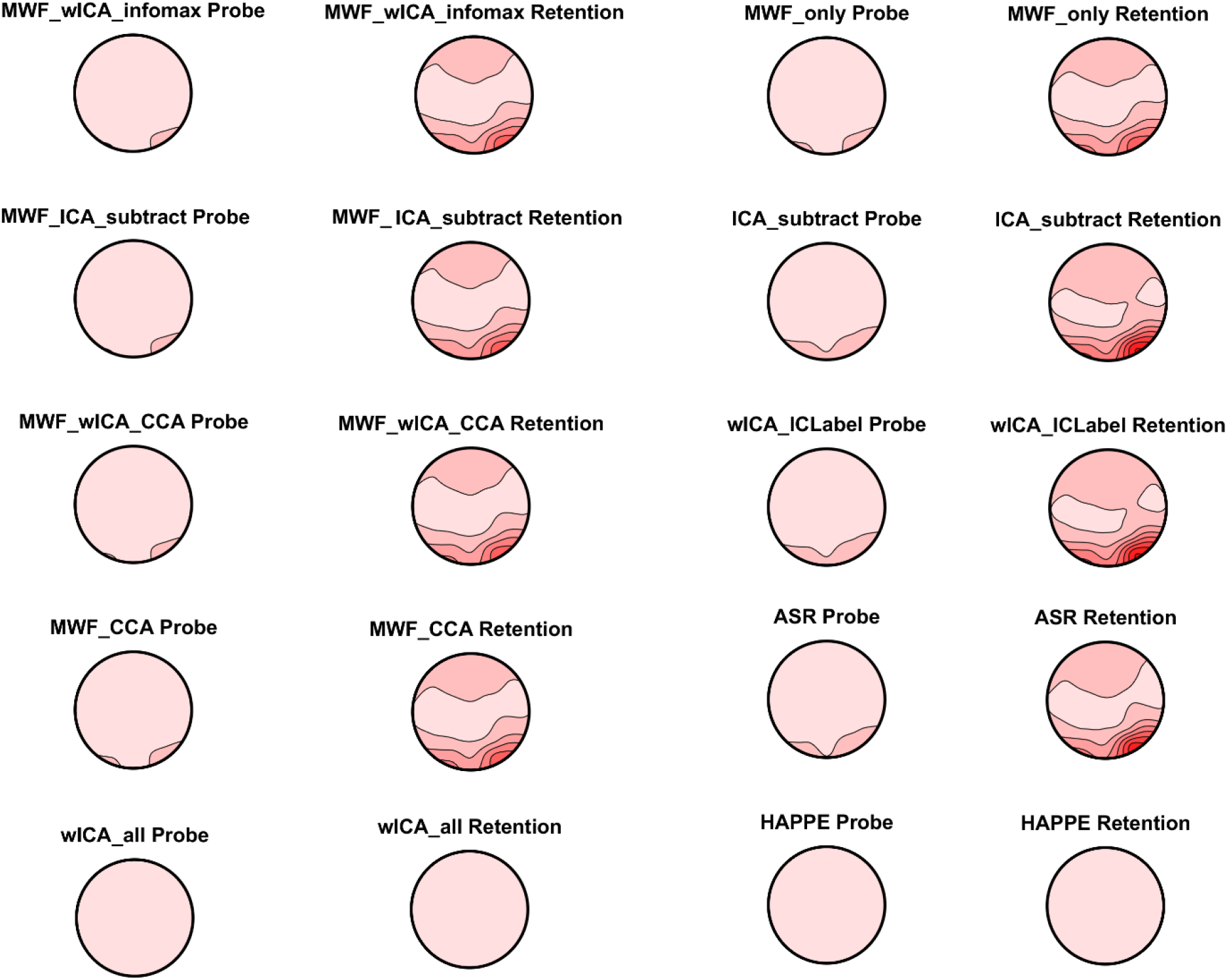
Alpha power distribution during the early period (0 to 750ms) after the stimuli of the working memory delay (retention) and working probe periods from each of the cleaning pipelines. All plots are on the same scale so they can all be compared to all other pipelines. Note that wICA_all and HAPPE have removed almost all alpha activity compared to the other pipelines.

**Figure 12.**
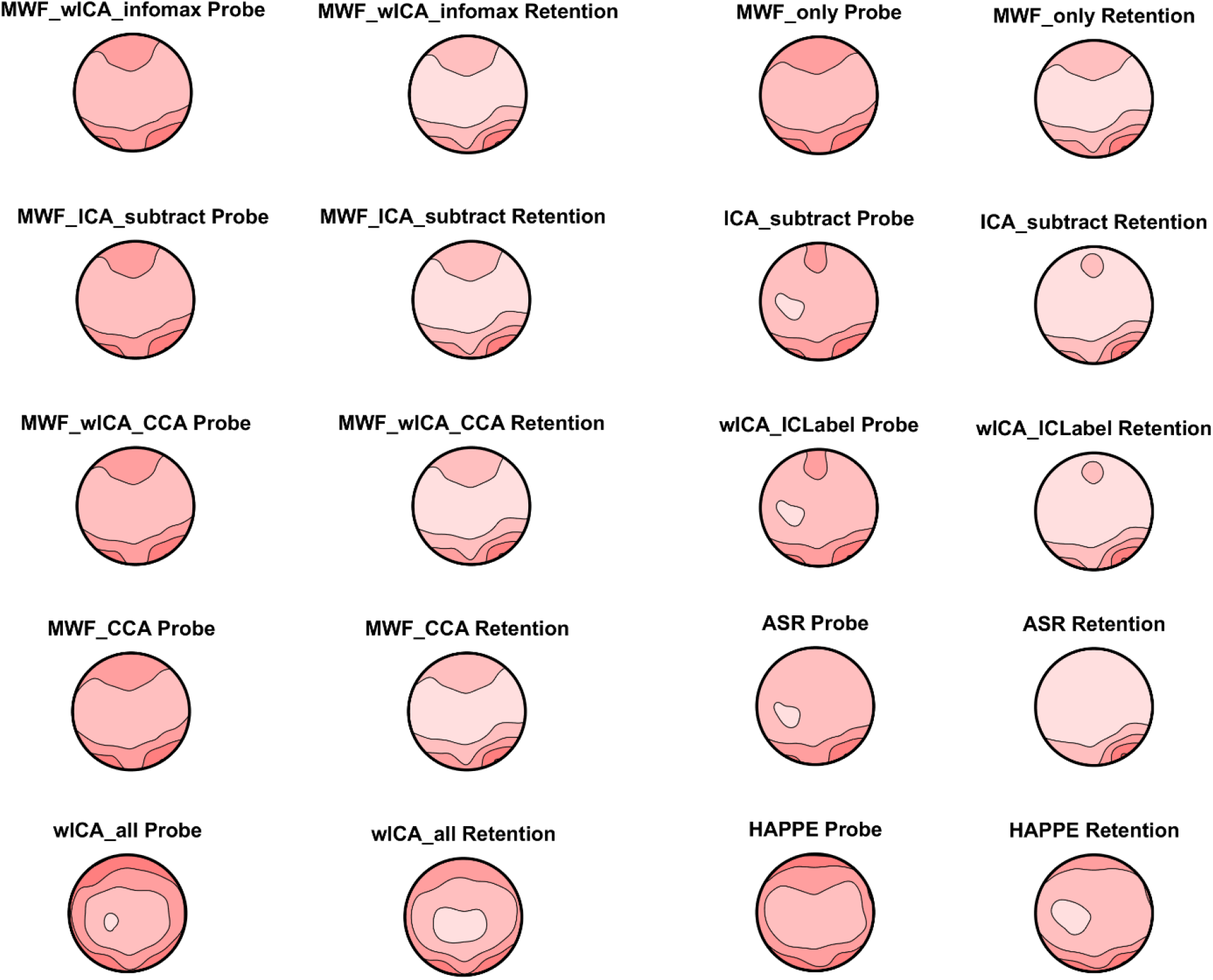
Alpha power distribution during the early period (0 to 750ms) after the stimuli of the working memory delay (retention) and working probe periods from each of the cleaning pipelines. All plots are on their own individual scale so the pattern of alpha activity distributions can be viewed within each plot (but comparisons of amplitudes between pipelines are not useful). As expected, the retention period showed more occipital / parietal alpha maximums, while the probe period showed more widespread alpha. Note the similarity in pattern across most pipelines, including ICA only and MWF only methods, implying different cleaning approaches still lead to similar patterns. ASR, HAPPE, and wICA_all were the most different to the other cleaning pipelines, with ASR removing most of the frontal alpha, and HAPPE and wICA showing less qualitative differentiation between the probe and retention period alpha, despite these pipelines showing the highest variance explained for the comparison between the two conditions. We suspect this may be the result of removal of significant amount of the variance by these pipelines, leading to highly precise estimates of the retention and probe alpha activity, which allow strong inferences about the differences in these periods (which no longer overlap after the removal of so much variance). However, we suspect this comes at the cost of distorting the distribution/characterization of the neural activity.

With regards to comparisons of the distribution of activity (assessed by the TANOVA), two separate time periods showed a difference in the distribution of alpha activity between the Sternberg delay and probe period in all pipelines – first a large difference from 0 to 750ms, then a smaller difference from 750 to 2000ms. For the first time-period (0 to 750ms), HAPPE, wICA_ICLabel and ICA_subtract showed the best performance, significantly better than wICA_all, which outperformed MWF_only, and MWF_CCA, which outperformed ASR, MWF_wICA, MWF_wICA_CCA, and MWF_ICA_subtract. However, it is worth noting that for this metric, all pipelines produced np^2^ values between 0.24 and 0.28, so the differences were not large. For the second time-period examined with the TANOVA test (750 to 2000ms), a different pattern was apparent, with ICA_subtract and wICA_ICLabel performing the best, follow ed by wICA_all, which performed better than MWF_only, MWF_CCA, MWF_wICA_CCA, MWF_wICA, and MWF_ICA_subtract. HAPPE showed the worst performance. However, similar to the earlier time-period, all pipelines produced np^2^ values between 0.18 and 0.25, so the differences were not large.

Together, these findings demonstrate that different cleaning pipelines can alter the EEG signal in differing ways. While the pattern of change during different resting-state and task conditions is broadly similar across pipelines, the choice of pipeline can alter the overall effect size between conditions. Additionally, while most pipelines showed a very similar distribution of alpha activity, HAPPE and wICA_all showed a different distribution of activity to all other pipelines, suggesting (alongside the alpha RMS, SER, and ARR values) that these two pipelines may be overcleaning the data, removing alpha power from specific electrodes such that the distribution of activity after cleaning is significantly altered (see Figures 11 and 12, and Supplementary Materials Figure S31-32, pages 51-52).

## Discussion

This study reports the results of comparisons between multiple variations of our newly developed RELAX EEG cleaning pipeline and six other commonly used pipelines. These comparisons were made across a wide range of cleaning and signal preservation metrics. Considering the results of the comparisons across the cleaning pipelines, multiple datasets with different recording parameters, and taking account of the overall performance on both a range of cleaning quality and signal preservation metrics, we conclude that the RELAX MWF_wICA pipeline provided the best performance at cleaning the data (without “overcleaning” the data), while also performing well at detecting experimental effects. Additionally, either the RELAX MWF_wICA or wICA_ICLabel methods were amongst the highest performing pipelines in metrics assessing the ability to detect experimental effects after cleaning with the pipeline. MWF_wICA also provided high values for both signal preservation (SER) and overall artifact reduction (ARR), with simultaneously higher values for both SER and ARR than ASR or ICA_subtract across all datasets, indicating MWF_wICA was better at both removing artifacts and preserving signal. While MWF_only or wICA_ICLabel approaches maintained more of the signal as indicated by higher SER values, these methods were also less effective at reducing artifacts, as indicated by lower ARR values and poorer performance in the blink and muscle metrics. Blink and muscle metrics also showed that MWF_wICA was better at cleaning these artifacts than MWF_only, wICA_ICLabel, ICA_subtract, and ASR. Additionally, the RELAX MWF_wICA methods resulted in the least amount of data having to be removed by cleaning and outlying epoch exclusion, except for HAPPE and wICA_all (which our results indicate excessively reduce the neural signal as well as the artifacts). This means that the RELAX MWF_wICA method makes more data available for inclusion in analyses than most pipelines, reducing the risk that participant data needs to be excluded for having too few epochs for inclusion, or that results might be biased by exclusion of epochs containing neural activity due to inferior cleaning. Our companion article additionally indicated that MWF_wICA produced data that showed amongst the highest scores for reliability of ERPs across trials and participants, suggesting cleaning with MWF_wICA provided amongst the most consistency in detecting experimental effects (Bailey et al., 2022). The inclusion of more high-quality epochs for analysis and more reliable data has been suggested by simulations to improve study power even more than increasing sample size (Clayson, Carbine, et al., 2021; Kolossa & Kopp, 2018; Luck et al., 2021). For these reasons, we recommend the use of the RELAX pipeline with MWF_wICA as a default effective EEG cleaning pipeline when no rationale exists to prefer another variation of RELAX. However, in specific circumstances, the RELAX pipeline with the wICA_ICLabel setting might be preferred (discussed in more detail below). Our results also recommend MWF_wICA or wICA_ICLabel should be preferred over the approach that is most used in the literature currently - ICA_subtract.

Our results indicate the value of combining multiple artifact reduction approaches in a single EEG cleaning pipeline, which is likely to address artifacts that a single method in isolation might not clean completely. For example, muscle activity often appeared to be inadequately addressed by MWF alone, and blinks were not as effectively addressed by wICA_ICLabel alone, while the combination of both methods did effectively clean both artifacts (note that it is easy to perform only MWF or only wICA_ICLabel cleaning with RELAX, so users can test this on their own data). This is in alignment with some other research, where combination cleaning approaches that performed CCA, ICA and independent vector analysis have also been demonstrated to be more effective than single approaches (Barban et al., 2021). We suspect a benefit of the combined MWF and wICA approach is that the combination provided a useful backup (in case one step inadequately addressed an artifact, the other step was still likely to reduce that artifact) while not over-cleaning the data (if the MWF cleaning completely addressed the blink artifact, ICLabel would not detect any blink artifact and thus the wICA step would not further attempt to reduce blink artifacts leading to over cleaning). However, while adding CCA to the MWF pipeline did improve the cleaning of muscle activity compared to MWF_only, it also resulted in inferior blink correction compared to MWF_wICA, suggesting that wICA applied to artifacts detected by ICLabel is an optimal second step as it addresses all artifacts rather than just muscle artifacts. Additionally, using CCA in addition to MWF_wICA (MWF_wICA_CCA) did not enhance performance compared to MWF_wICA. As such, while the application of specific multiple cleaning methods does seem to improve cleaning performance, it is not the case that adding more cleaning methods is always better. Additionally, our results suggested that both when applied in isolation and after MWF cleaning, wICA_ICLabel was better at preserving signal (higher SER values) than ICA_subtract, while still providing similar artifact reduction (ARR values). Since the SER reflects the ability of a pipeline to preserve signal in periods that are not affected by artifacts, we suspect this highlights the manner in which subtracting an ICA component means removing not only artifact, but neural activity that is mixed into the component due to imperfect ICA decomposition. In contrast, because of the wavelet thresholding used within wICA, only the largest amplitude signal is removed (which likely represents artifact), while the clean data periods are left unaffected (producing higher SER values). Similarly, while subtraction of ICA after MWF reduced the metrics assessing remaining muscle activity to approximately zero, the method never led to higher explained variance from the experimental tests and resulted in lower variance explained in most metrics. As such, while subtracting artifactual components with ICA is probably the most common cleaning approach, we recommend the use of wICA to reduce artifactual components instead. Given these two points, our recommended RELAX pipeline applies MWF cleaning then wICA cleaning (but not CCA or ICA subtraction).

While the preceding paragraphs suggest that the cleaning metrics we tested strongly recommend the use of RELAX with the MWF_wICA setting, our perspective is that the most important metric for cleaning efficacy is whether a cleaning approach enhances a researcher’s ability to detect experimental outcomes (a view shared by other researchers) (Clayson, Baldwin, et al., 2021). Here, the results do not indicate a single best cleaning approach for all applications. This is consistent with previous research, which has also been unable to recommend a single best pipeline for all situations (Barban et al., 2021). Encouragingly, except for the wICA_all and HAPPE pipelines, all the cleaning approaches we tested led to broadly similar amounts of variance explained by the experimental manipulation and similar patterns for outcome measures. This suggests that as long as the established confounding influences of artifacts are sufficiently addressed and signal is sufficiently preserved, EEG cleaning choices may not severely confound results across different studies, at least providing effects in the same direction and of similar magnitudes. However, we deliberately chose tasks that provided well established reasonably sized differences between the experimental conditions. Studies examining differences between clinical groups or treatment conditions often show smaller effect sizes than between condition comparisons, in which case optimal pipeline selection might be more likely to influence results. Indeed, some research has suggested that subtle effects do not always replicate between different data cleaning approaches, and that effect sizes vary across EEG cleaning pipelines (Clayson, Baldwin, et al., 2021; Robbins et al., 2020; Rogasch et al., 2020). As such, we recommend the use of RELAX MWF_wICA as a default due to its superior artifact cleaning and signal preservation attributes. However, for specific uses, the wICA_ICLabel setting might at times be preferred.

In particular, the amount of variance explained by the experimental manipulations indicated that the MWF_wICA pipeline did not significantly differ from the most effective pipelines when examining resting-state alpha power. However, the wICA_ICLabel setting of RELAX performed more highly in alpha power measures from the Sternberg task, and as such might be preferred for studies focused on similar oscillatory outcome measures. Having said that, because wICA_ICLabel did not clean blinks or muscle as effectively, we recommend wICA_ICLabel only be used where enough trials are collected (or enough participants) that lower reliability could be acceptable, as research has suggested data quality is more important for power than data quantity (Kolossa & Kopp, 2018). As such, wICA_ICLabel might be preferred when using robust statistics and methods that include all single trials in the statistical inferences, where reduced cleaning can avoid the potential negative effects of arbitrary cleaning thresholds, and robust statistics can account for the noisier data (Alday & van Paridon, 2021). However, note that if an inadequately cleaned artifact is time-locked to a period of interest, even robust statistics might not address its potential to confound results.

wICA_ICLabel might additionally be recommended for studies examining connectivity between electrodes, as the effect of MWF on connectivity measures is currently unknown. For researchers interested in connectivity measures, it may be useful to note that cleaning EEG data using wavelet enhanced ICA (wICA) to reduce artifacts does not reduce the rank of the data, so might allow for higher resolution of nodes when using connectivity analysis in source space (in contrast to subtracting ICA components, which does reduce the rank) (Castellanos & Makarov, 2006). Additionally, for research examining gamma oscillations, it might be worth considering that MWF_ICA_subtract removed essentially all muscle activity, which is a potential confound in studies examining gamma oscillations due to the overlap in frequencies between muscle activity and gamma neural oscillations. However, note that the ‘ground-truth’ of gamma oscillations was not determined, so we cannot eliminate the possibility that MWF_ICA_subtract removed neural oscillations as well as muscle.

In contrast to the preceding recommendations, while ASR provided high values for the variance explained between our experimental conditions, there were no cases where it provided higher performance than both wICA_ICLabel and MWF_wICA, and ASR performed worse than MWF_wICA in all artifact reduction metrics. Our results also suggested potentially high variability in some measures after ASR cleaning, and our experience using ASR has suggested it produce inconsistent cleaning outcomes even when applied to the same file (including a high number of periods removed as reflecting extreme artifacts when using the default settings). Unfortunately, we were not able to discern whether this variability reflects ‘ground-truth’ differences (in which case the variability is valuable), or variability produced by the ASR cleaning (in which case the variability is artifactual), and our objective measures do not address the cause of the variability.

Our results also indicated that HAPPE and wICA_all were associated with very low SER values, and very low amplitude EEG activity after cleaning (including a reduction by >2 orders of magnitude for alpha power), indicating considerable reduction of the neural activity signal. After cleaning with these pipelines, the majority of epochs also showed log-power log-frequency slopes that were indicative of residual muscle activity. It may be that this was not because these pipelines inadequately cleaned muscle activity, but rather they over-cleaned the neural activity signal in the lower frequency ranges, so the cleaned slopes were flattened and did not reflect the 1/f power-frequency distributions of typical neural activity (Donoghue et al., 2020). However, our power frequency plots for wICA_all and HAPPE pipelines did show more power in the beta frequency range relative to lower frequencies compared to other pipelines (Figures 7 and 9), so it may be that these methods simply do not adequately clean muscle activity. The combination of low SER values, reduced power frequency amplitudes, and altered activity distributions compared to the other pipelines suggests that the wICA_all and HAPPE pipelines overclean the data, removing elements of the neural signal. It is worth noting that HAPPE was designed to address noisy data with large artifacts, so our result perhaps highlights that it is not appropriate for typical cognitive data without excessive noise (Gabard-Durnam et al., 2018). As such, while the HAPPE and wICA_all methods did seem to produce higher explained variance for several experimental outcomes, we do not recommend their use, except perhaps in the case of extremely noisy data, or where the outcome of interest is not concerned with characterizing the data, but simply differentiating two conditions (for example, brain-computer interface applications might usefully apply the wICA_all approach in real time using very rapid ICA computation approaches).

Since it seems there is not a single best cleaning pipeline that can be recommended, we have made RELAX modular so that different approaches can easily be implemented. Some default settings can be recommended. For example, the results reported in our companion paper indicated that the ICA methods infomax or fastICA with the symmetric setting were the best performers within RELAX (Bailey et al., 2022). If users can install cudaICA, this setting is likely to be optimal, providing both speed and the best performance (Raimondo et al., 2012). Traditional extended infomax can also be used, but is slower (Lee et al., 1999). Otherwise, we have left the default setting as fastICA with the symmetric setting (Hyvarinen, 1999), which our companion paper has indicate is (just slightly) better than the deflation setting (Bailey et al., 2022). We have set this fastICA to repeat the ICA up to three times in the case of a non-convergence issue which can adversely affect ICA decompositions, then to switch to the ‘defl’ setting if non-convergence still occurs. Other defaults have been set after an extensive process of informal testing to determine optimal cleaning and values for the variance explained by different experimental manipulations. Specific RELAX settings or parameter variations used in future research should be justified, primarily with reference to a previous demonstration of maximal explained variance in a similar experimental design. Ideally, the parameters selected should also be pre-registered to prevent “fishing” for positive results. However, while we recommend the use of methods that ensure “fishing” does not inflate the false positive rate, it may also be useful to other researchers to report analyses using multiple parameter selections. This would help guide future research towards the optimal methods for particular use cases. It may have the additional benefit of demonstrating robustness of results to variations in cleaning methods, eliminating the possibility that a specific result is produced by or dependent on a specific cleaning method. If researchers implement this approach, the a priori selected, primary, and ideally pre-registered cleaning method should be specified so readers can be aware of which analysis method was pre-planned, and understand the risk that positive results were produced by multiple re-analyses. RELAX provides an output of the cleaning efficacy metrics used in the current study, which should be reported so reviewers and readers can determine the risk that remaining artifacts could confound conclusions (perhaps in a Supplementary Materials section for the sake of brevity). We have provided further discussion points in our Supplementary Materials, section 7, pages 75-76).

### Limitations

Despite our extensive testing, there are many limitations to the conclusions we can draw from the current study. Firstly, while our results are relatively comprehensive, we have only scratched the surface of the full potential parameter space of cleaning options. We have also only tested the cleaning pipelines on a small number of datasets (although certainly with large sample sizes), and only tested the cleaning pipelines on a minority of experimental tasks and experimental outcome measures (although more than most cleaning studies - see also our companion article for examination of ERP related data, (Bailey et al., 2022)). In particular, we did not test the effect of data cleaning approaches on connectivity analyses or metrics that assess the 1/f aperiodic / non-oscillatory component of the data.

Secondly, we have not tested RELAX on extremely noisy data contaminated with large movement artifacts, nor scans of children/infants, nor the elderly who often produce EEG data with more artifacts, and which may contain different characteristics to the datasets we tested, potentially requiring alternative artifact/signal discrimination approaches. We note that ICLabel was built sampling from adult data (Pion-Tonachini et al., 2019), and the log-power log-frequency muscle slope thresholds were derived from adult data (Fitzgibbon et al., 2016). Similarly, ICLabel was built from data containing 32 or more electrodes, so we suspect data with less than 32 electrodes might not provide ICLabel with sufficient data for optimal artifact detection, leading to reduced cleaning performance (Pion-Tonachini et al., 2019). It is also worth noting that short EEG data periods may not have enough samples for effective ICA computation, nor for identification of artifacts for MWF cleaning. At least [Number of channels ^ 2 x (∼30)] data points are recommended (∼2.5 minutes for 64 channel data recorded at 1000Hz) (Miyakoshi, 2018).

Third, while automation of EEG data pre-processing prevents the risk that subjectivity would affect the data compared to manual pre-processing, there are also potential pitfalls. One particularly risky pitfall is that some files may be corrupted during the recording process and contain no neural related data whatsoever, an issue that automated cleaning methods may not identify. This issue occurred for one file we submitted to our pre-processing pipelines. As such, we recommend that EEG recording technicians exclude data files that appear to contain no usable signal before submitting files to pre-processing. As a double-check, we have also included in the pipeline a visual test of outlying median voltages from each electrode in the cleaned data, which indicates if a particular file contains atypical data to be manually checked (a method that effectively identified our problem file).

Additionally, there is no way to be sure of the ‘ground-truth’ of our tests of the amount of variance explained by different experimental manipulations. As such, it may be possible that the higher levels of explained variance are due to artifacts rather than actual differences in neural activity. However, this seems an unlikely explanation for our results for a number of reasons. First, the artifacts would have to be both time-locked to the stimuli in the case of the Sternberg task (and the Go Nogo data reported in our companion manuscript) and appear more strongly or more frequently in one condition than the other. The only artifact that is likely to fit this requirement is eye movement. However, all pipelines produce blink amplitude ratios ranging from 1.015 to 1.077, values that are likely too small to produce much influence on differences in variance explained between pipelines. Second, we think it is unlikely that any artifact would increase the explained variance by the experimental manipulation across multiple different tasks and types of brain activity (especially considering the different topographical patterns and underpinning mechanisms for each measure). Particularly since the measures used were also selected because they have been robustly characterised by previous research, which has demonstrated that the experimental manipulations lead to the patterns of brain activity that we detected. Finally, if the explained variance was due to the influence of artifacts, we would instead see an artifact pattern in the TANOVA for one of the conditions (the most obvious of which would have been a blink topography). No such artifact topography was present in any of our pipelines or comparisons for the tests of experimental effects. While it may be that previous research that has demonstrated the expected effect of the different experimental manipulations might have also been driven by artifacts in the data, these artifacts would have to be consistent across the range of cleaning pipelines that have replicated the effect for the analyses we included (including the range of pipelines tested in our study, which almost all showed the expected effects). In some cases, previous research has used electrocorticography or fMRI in conjunction with EEG to demonstrate that EEG measures relate to brain activity detected with these other methods, providing confidence that artifacts are not producing the result (Ahmad et al., 2016; Baumeister et al., 2014; Iannaccone et al., 2015; Meltzer et al., 2007; Smith et al., 2013; Zhang et al., 2018). The putative brain activity detected for many of the metrics we included has also been shown to predict behavioural performance, suggesting functional relationships that are unlikely to be present for artifacts (Bashivan et al., 2014; Clark et al., 2004; Karamacoska et al., 2018). As such, we suggest that our results indicate that RELAX cleans artifacts in such a way that the signal (brain) to noise (muscle, eye movement etc.) ratio is maximised, and the higher explained variance values reflect better detection of the effects of the experimental manipulation on the brain activity rather than artifacts remaining after cleaning, providing a strong recommendation for its use when cleaning EEG data.

#### Potential improvements and future research

We tested many potential versions of RELAX, and informally explored most parameter settings for optimal performance prior to formal testing. However, there are potential improvements that may be possible which are worth exploring in future EEG cleaning pipeline development. These are discussed in full in the Supplementary Materials (section 7, page 76), but include: 1) using an adaptive threshold for the wICA cleaning (which we briefly tested using an array of the level dependent settings within the MATLAB function ‘wdenoise’, but found this approach to be less effective at reducing blink artifacts); 2) using a less stringent amplitude threshold for outlying epoch rejection after cleaning, or alternatively, no epoch rejection and robust statistics for comparisons (Alday & van Paridon, 2021); 3) taking temporal information into account in the ICA computation (with Independent Vector Analysis for example; (Barban et al., 2021), although we note that this is slow and potentially complicated to implement); and 4) adapting ICLabel to identify artifactual components based on empirically established objective methods, rather than its current design which used expert consensus (for example, applying the objective approach to identifying muscle components based on comparison of paralysed vs non-paralysed scalp recordings provided by Fitzgibbon et al. [2016]).

The current study also required a very large amount of computation and researcher hours, involving massive repeated iterative testing of different cleaning parameters by multiple researchers across multiple datasets and multiple outcome metrics. As such, the comprehensiveness of a project like this is difficult to achieve. Despite this difficulty, we would recommend examining an even higher number of metrics and varying an even broader parameter space in future pipeline development. To achieve this, we suggest to the field that an online resource containing multiple large EEG datasets could be collated for the purposes of testing EEG cleaning pipelines (note that small scale examples set up that enabling testing of cleaning approaches already exist (Barban et al., 2021; Kappenman et al., 2021; Zhang et al., 2020), and large-scale open access datasets are also available (van Dijk et al., in submission). The cleaning efficacy of newly proposed pipelines could be compared via these datasets across multiple metrics, and after cleaning, multiple neural activity measures of explained variance could be assessed (for example, ERPs, oscillations, 1/f metrics, and connectivity measures). A ‘leader board’ of the current optimal methods could be established, so that new cleaning approaches would not have to replicate pipelines that have already been tested to make comparisons. An automated algorithm could iteratively test parameter details that may affect cleaning quality (such as the wICA threshold or outlier identification thresholds). It is also possible for different settings of different parameters to interact with each other to influence cleaning performance. For example, lower extreme outlier exclusion thresholds may improve wICA performance more for higher wICA thresholds than lower thresholds. As such, the potential parameter space might be practically infinite. This being the case, automated EEG cleaning pipeline development is much better performed by an automated system, perhaps with many pipelines being tested concurrently to optimise pipeline development as quickly as possible. If this were achieved, the most optimal current pipeline could be automatically uploaded to GitHub each time the testing system discovered an improvement. Different outcome measures or experimental designs could even have their own version of the most optimal pipeline (Clayson, Baldwin, et al., 2021). As such, the system could reflect a “living” gold-standard in EEG pre-processing. We unfortunately do not have the resources to implement this approach, and we are not able to share our data due to ethical approval criteria. We mention it here in the hope that the field may be inspired to achieve this, perhaps via large scale collaboration (we note that #EEGManyLabs could be a good platform for this discussion) (Pavlov et al., 2021).

## Conclusion

To conclude, we recommend future researchers use the RELAX approach with the default MWF_wICA setting for cleaning EEG when no clear rationale exists to use another setting, as it provides the following benefits:

1. RELAX MWF_wICA provided the best or equivalent to the best cleaning of artifacts.
2. RELAX is fully automatic and quick to apply, saving considerable time, and eliminating subjectivity throughout the cleaning process.
3. The pipeline contains embedded cleaning quality metrics, which can also be easily reported in publications. This will enable researchers to evaluate the likelihood that the specific cleaning parameters or remaining artifacts might be influencing conclusions drawn from the data.
4. The RELAX MWF_wICA approach maximises the number of epochs included for analysis, thus reducing the possibility of biasing the data based on epoch exclusion and increasing the statistical power of the study.
5. RELAX MWF_wICA provided high levels of variance explained by the experimental manipulation. We have provided the source code and made the pipeline straightforward to use and publicly accessible, with the only technical skills required being the installation of the MATLAB specific and external dependencies. RELAX also does not require EOG electrodes (unlike regression approaches), so is applicable to a wider range of datasets and robust against participant discomfort with eye electrodes (Gómez-Herrero et al., 2006), as well as the somewhat common (in our experience) “bad eye electrode” issue. RELAX is available for download from GitHub (https://github.com/NeilwBailey/RELAX/releases). The pipeline is designed to run within the commonly used EEGLAB software (implemented in MATLAB). Finally, we note the acronym “RELAX” was also chosen as a prompt for researchers - while technological advances have the potential to reduce our workload, they have typically made us simply more productive instead. Given the stressors researchers are exposed to (Bowen et al., 2016), and the reductions in health associated with that stress (Faragher et al., 2013; Nixon et al., 2011), we recommend that at least part of the time saved by using our pipeline could be spent on researcher well-being, rather than simply further increasing productivity.

### Recommendations for using RELAX when examining oscillations

1. Use RELAX_MWF_wICA as the default cleaning pipeline (with fastICA on the symm setting, or cudaICA if it can be installed)
2. Use RELAX_wICA_ICLabel if analysing task related oscillatory power or connectivity, but only if data are relatively clean, if using analysis methods that prevent remaining artifacts confounding conclusions (for example single trial analyses with robust statistics) and if a large number of trials is available for analysis.
3. Consider using RELAX_MWF_ICA_subtract if analysing gamma power (or if removing all muscle activity is required)

See our companion article for recommendations related to ERP analyses (Bailey et al., 2022).

## Notes

https://github.com/NeilwBailey/RELAX/releases/

### Competing Interest Statement

PBF has received equipment for research from MagVenture A/S, Nexstim, Neuronetics and Brainsway Ltd and funding for research from Neuronetics. He is a founder of TMS Clinics Australia and Resonance Therapeutics. MB, ATH, NWB, AM, MB, NCR and OWM reported no biomedical financial interests or potential conflicts of interest.

## References

Ahmad, R. F., Malik, A. S., Kamel, N., Reza, F., & Abdullah, J. M. (2016). Simultaneous EEG-fMRI for working memory of the human brain. Australasian physical & engineering sciences in medicine, 39(2), 363–378.

Akhtar, M. T., Mitsuhashi, W., & James, C. J. (2012). Employing spatially constrained ICA and wavelet denoising, for automatic removal of artifacts from multichannel EEG data. Signal processing, 92(2), 401–416.

Alday, P. M., & van Paridon, J. (2021). Away from arbitrary thresholds: using robust statistics to improve artifact rejection in ERP.

Allen, M., Poggiali, D., Whitaker, K., Marshall, T. R., & Kievit, R. A. (2019). Raincloud plots: a multi-platform tool for robust data visualization. Wellcome open research, 4.

Anders, M., Anders, B., Kreuzer, M., Zinn, S., & Walter, C. (2020). Application of referencing techniques in EEG-based Recordings of Contact Heat Evoked Potentials (CHEPS). Frontiers in human neuroscience, 14, 527.

Bailey, N., Hill, A., Biabani, M., Murphy, O., Rogasch, N., McQueen, B., Miljevic, A., & Fitzgerald, P. (2022). Introducing RELAX (the Reduction of Electroencephalographic Artifacts): A fully automated pre-processing pipeline for cleaning EEG data – Part 2: Application to Event-Related Potentials. bioRxiv.

Bailey, N. W., Freedman, G., Raj, K., Spierings, K. N., Piccoli, L. R., Sullivan, C. M., Chung, S. W., Hill, A. T., Rogasch, N. C., & Fitzgerald, P. B. (2020). Mindfulness meditators show enhanced accuracy and different neural activity during working memory. Mindfulness, 11, 1762–1781.

Barban, F., Chiappalone, M., Bonassi, G., Mantini, D., & Semprini, M. (2021). Yet another artefact rejection study: an exploration of cleaning methods for biological and neuromodulatory noise. Journal of Neural Engineering.

Bashivan, P., Bidelman, G. M., & Yeasin, M. (2014). Spectrotemporal dynamics of the EEG during working memory encoding and maintenance predicts individual behavioral capacity. European Journal of Neuroscience, 40(12), 3774–3784.

Baumeister, S., Hohmann, S., Wolf, I., Plichta, M. M., Rechtsteiner, S., Zangl, M., Ruf, M., Holz, N., Boecker, R., & Meyer-Lindenberg, A. (2014). Sequential inhibitory control processes assessed through simultaneous EEG–fMRI. NeuroImage, 94, 349–359.

Bender, R., & Lange, S. (2001). Adjusting for multiple testing—when and how? Journal of clinical epidemiology, 54(4), 343–349.

Benjamini, Y., & Hochberg, Y. (1995). Controlling the false discovery rate: a practical and powerful approach to multiple testing. Journal of the Royal statistical society: series B (Methodological), 57(1), 289–300.

Bertrand, A. (2015). Distributed signal processing for wireless EEG sensor networks. IEEE transactions on neural systems and rehabilitation engineering, 23(6), 923–935.

Bigdely-Shamlo, N., Mullen, T., Kothe, C., Su, K.-M., & Robbins, K. A. (2015). The PREP pipeline: standardized preprocessing for large-scale EEG analysis. Frontiers in neuroinformatics, 9, 16.

Borowicz, A. (2018). Using a multichannel Wiener filter to remove eye-blink artifacts from EEG data. Biomedical Signal Processing and Control, 45, 246–255.

Bowen, P., Rose, R., & Pilkington, A. (2016). Perceived stress amongst university academics. American International Journal of Contemporary Research, 6(1), 22–28.

Castellanos, N. P., & Makarov, V. A. (2006). Recovering EEG brain signals: Artifact suppression with wavelet enhanced independent component analysis. Journal of neuroscience methods, 158(2), 300–312.

Chang, C.-Y., Hsu, S.-H., Pion-Tonachini, L., & Jung, T.-P. (2019). Evaluation of artifact subspace reconstruction for automatic artifact components removal in multi-channel EEG recordings. IEEE Transactions on Biomedical Engineering, 67(4), 1114–1121.

Clark, C. R., Veltmeyer, M. D., Hamilton, R. J., Simms, E., Paul, R., Hermens, D., & Gordon, E. (2004). Spontaneous alpha peak frequency predicts working memory performance across the age span. International Journal of Psychophysiology, 53(1), 1–9.

Clayson, P. E., Baldwin, S., Rocha, H., & Larson, M. J. (2021). The Data-Processing Multiverse of Event-Related Potentials (ERPs): A Roadmap for the Optimization and Standardization of ERP Processing and Reduction Pipelines.

Clayson, P. E., Carbine, K. A., Baldwin, S. A., Olsen, J. A., & Larson, M. J. (2021). Using generalizability theory and the ERP Reliability Analysis (ERA) Toolbox for assessing test-retest reliability of ERP scores Part 1: Algorithms, framework, and implementation. International Journal of Psychophysiology.

Delorme, A., & Makeig, S. (2004). EEGLAB: an open source toolbox for analysis of single-trial EEG dynamics including independent component analysis. Journal of neuroscience methods, 134(1), 9–21.

Delorme, A., Sejnowski, T., & Makeig, S. (2007). Enhanced detection of artifacts in EEG data using higher-order statistics and independent component analysis. NeuroImage, 34(4), 1443–1449.

Dimigen, O. (2020). Optimizing the ICA-based removal of ocular EEG artifacts from free viewing experiments. NeuroImage, 207, 116117.

Donoghue, T., Haller, M., Peterson, E. J., Varma, P., Sebastian, P., Gao, R., Noto, T., Lara, A. H., Wallis, J. D., & Knight, R. T. (2020). Parameterizing neural power spectra into periodic and aperiodic components. Nature neuroscience, 23(12), 1655–1665.

Faragher, E. B., Cass, M., & Cooper, C. L. (2013). The relationship between job satisfaction and health: a meta-analysis. From stress to wellbeing Volume 1, 254–271.

Fitzgibbon, S., DeLosAngeles, D., Lewis, T., Powers, D., Grummett, T., Whitham, E., Ward, L., Willoughby, J., & Pope, K. (2016). Automatic determination of EMG-contaminated components and validation of independent component analysis using EEG during pharmacologic paralysis. Clinical neurophysiology, 127(3), 1781–1793.

Gabard-Durnam, L. J., Mendez Leal, A. S., Wilkinson, C. L., & Levin, A. R. (2018). The Harvard Automated Processing Pipeline for Electroencephalography (HAPPE): standardized processing software for developmental and high-artifact data. Frontiers in neuroscience, 12, 97.

Gómez-Herrero, G., De Clercq, W., Anwar, H., Kara, O., Egiazarian, K., Van Huffel, S., & Van Paesschen, W. (2006). Automatic removal of ocular artifacts in the EEG without an EOG reference channel. Proceedings of the 7th Nordic Signal Processing Symposium-NORSIG 2006,

Habermann, M., Weusmann, D., Stein, M., & Koenig, T. (2018). A student’s guide to randomization statistics for multichannel event-related potentials using ragu. Frontiers in neuroscience, 12, 355.

Hyvarinen, A. (1999). Fast ICA for noisy data using Gaussian moments. 1999 IEEE international symposium on circuits and systems (ISCAS),

Iannaccone, R., Hauser, T. U., Staempfli, P., Walitza, S., Brandeis, D., & Brem, S. (2015). Conflict monitoring and error processing: new insights from simultaneous EEG–fMRI. NeuroImage, 105, 395–407.

Inuso, G., La Foresta, F., Mammone, N., & Morabito, F. C. (2007). Wavelet-ICA methodology for efficient artifact removal from Electroencephalographic recordings. 2007 international joint conference on neural networks,

Islam, M. K., Rastegarnia, A., & Yang, Z. (2016). Methods for artifact detection and removal from scalp EEG: A review. Neurophysiologie Clinique/Clinical Neurophysiology, 46(4-5), 287–305.

Issa, M. F., & Juhasz, Z. (2019). Improved EOG artifact removal using wavelet enhanced independent component analysis. Brain sciences, 9(12), 355.

Janani, A. S., Grummett, T. S., Lewis, T. W., Fitzgibbon, S. P., Whitham, E. M., DelosAngeles, D., Bakhshayesh, H., Willoughby, J. O., & Pope, K. J. (2018). Improved artefact removal from EEG using Canonical Correlation Analysis and spectral slope. Journal of neuroscience methods, 298, 1–15.

Kappenman, E. S., Farrens, J. L., Zhang, W., Stewart, A. X., & Luck, S. J. (2021). ERP CORE: An open resource for human event-related potential research. NeuroImage, 225, 117465.

Karamacoska, D., Barry, R. J., & Steiner, G. Z. (2018). Electrophysiological underpinnings of response variability in the Go/NoGo task. International Journal of Psychophysiology, 134, 159–167.

Kleifges, K., Bigdely-Shamlo, N., Kerick, S. E., & Robbins, K. A. (2017). BLINKER: Automated extraction of ocular indices from EEG enabling large-scale analysis. Frontiers in neuroscience, 11, 12.

Koenig, T., Kottlow, M., Stein, M., & Melie-García, L. (2011). Ragu: a free tool for the analysis of EEG and MEG event-related scalp field data using global randomization statistics. Computational intelligence and neuroscience, 2011.

Kolossa, A., & Kopp, B. (2018). Data quality over data quantity in computational cognitive neuroscience. NeuroImage, 172, 775–785.

Kumaravel, V. P., Farella, E., Parise, E., & Buiatti, M. (2022). Near: An artifact removal pipeline for human newborn EEG data. Developmental Cognitive Neuroscience, 101068.

Lee, T.-W., Girolami, M., & Sejnowski, T. J. (1999). Independent component analysis using an extended infomax algorithm for mixed subgaussian and supergaussian sources. Neural computation, 11(2), 417–441.

Luck, S. J., Stewart, A. X., Simmons, A. M., & Rhemtulla, M. (2021). Standardized measurement error: A universal metric of data quality for averaged event-related potentials. *Psychophysiology*, e13793.

Mair, P., & Wilcox, R. (2020). Robust statistical methods in R using the WRS2 package. Behavior research methods, 52(2), 464–488.

Mammone, N., La Foresta, F., & Morabito, F. C. (2011). Automatic artifact rejection from multichannel scalp EEG by wavelet ICA. IEEE Sensors Journal, 12(3), 533–542.

Meltzer, J. A., Negishi, M., Mayes, L. C., & Constable, R. T. (2007). Individual differences in EEG theta and alpha dynamics during working memory correlate with fMRI responses across subjects. Clinical neurophysiology, 118(11), 2419–2436.

Miyakoshi, M. (2018). Makoto’s preprocessing pipeline. https://sccn.ucsd.edu/wiki/Makotos_preprocessing_pipeline. Accessed February, 1, 2019.

Mumtaz, W., Rasheed, S., & Irfan, A. (2021). Review of challenges associated with the EEG artifact removal methods. Biomedical Signal Processing and Control, 68, 102741.

Muthukumaraswamy, S. (2013). High-frequency brain activity and muscle artifacts in MEG/EEG: a review and recommendations. Frontiers in human neuroscience, 7, 138.

Nixon, A. E., Mazzola, J. J., Bauer, J., Krueger, J. R., & Spector, P. E. (2011). Can work make you sick? A meta-analysis of the relationships between job stressors and physical symptoms. Work & Stress, 25(1), 1–22.

Nolan, H., Whelan, R., & Reilly, R. B. (2010). FASTER: fully automated statistical thresholding for EEG artifact rejection. Journal of neuroscience methods, 192(1), 152–162.

Oostenveld, R., Fries, P., Maris, E., & Schoffelen, J.-M. (2011). FieldTrip: open source software for advanced analysis of MEG, EEG, and invasive electrophysiological data. Computational intelligence and neuroscience, 2011.

Palmer, J. A., Kreutz-Delgado, K., & Makeig, S. (2012). AMICA: An adaptive mixture of independent component analyzers with shared components. Swartz Center for Computatonal Neursoscience, University of California San Diego, Tech. Rep.

Pavlov, Y. G., Adamian, N., Appelhoff, S., Arvaneh, M., Benwell, C. S., Beste, C., Bland, A. R., Bradford, D. E., Bublatzky, F., & Busch, N. A. (2021). # EEGManyLabs: Investigating the replicability of influential EEG experiments. Cortex.

Pion-Tonachini, L., Kreutz-Delgado, K., & Makeig, S. (2019). ICLabel: An automated electroencephalographic independent component classifier, dataset, and website. NeuroImage, 198, 181–197.

Raimondo, F., Kamienkowski, J. E., Sigman, M., & Fernandez Slezak, D. (2012). CUDAICA: GPU optimization of infomax-ICA EEG analysis. Computational intelligence and neuroscience, 2012.

Ranjan, R., Sahana, B. C., & Bhandari, A. K. (2021). Ocular artifact elimination from electroencephalography signals: A systematic review. Biocybernetics and Biomedical Engineering.

Robbins, K. A., Touryan, J., Mullen, T., Kothe, C., & Bigdely-Shamlo, N. (2020). How sensitive are EEG results to preprocessing methods: a benchmarking study. IEEE transactions on neural systems and rehabilitation engineering, 28(5), 1081–1090.

Rogasch, N. C., Sullivan, C., Thomson, R. H., Rose, N. S., Bailey, N. W., Fitzgerald, P. B., Farzan, F., & Hernandez-Pavon, J. C. (2017). Analysing concurrent transcranial magnetic stimulation and electroencephalographic data: A review and introduction to the open-source TESA software. NeuroImage, 147, 934–951.

Rogasch, N. C., Zipser, C., Darmani, G., Mutanen, T. P., Biabani, M., Zrenner, C., Desideri, D., Belardinelli, P., Müller-Dahlhaus, F., & Ziemann, U. (2020). The effects of NMDA receptor blockade on TMS-evoked EEG potentials from prefrontal and parietal cortex. Scientific reports, 10(1), 1–12.

Rošťáková, Z., & Rosipal, R. (2021). Determining the number of components in the PARAFAC model with a nonnegative tensor structure: A simulated EEG data study.

Smith, J. L., Jamadar, S., Provost, A. L., & Michie, P. T. (2013). Motor and non-motor inhibition in the Go/NoGo task: an ERP and fMRI study. International Journal of Psychophysiology, 87(3), 244–253.

Somers, B., & Bertrand, A. (2016). Removal of eye blink artifacts in wireless EEG sensor networks using reduced-bandwidth canonical correlation analysis. Journal of Neural Engineering, 13(6), 066008.

Somers, B., Francart, T., & Bertrand, A. (2018). A generic EEG artifact removal algorithm based on the multi-channel Wiener filter. Journal of Neural Engineering, 15(3), 036007.

Somers, B., Francart, T., & Bertrand, A. (2019). MWF toolbox for EEG artifact removal.

van Dijk, H., van Wingen, G., Denys, D., Olbrich, S., van Ruth, R., & Arns, M. (in submission). The two decades - Brainclinics research archive for insights in neurophysiology (TD-BRAIN) database.

Winkler, I., Debener, S., Müller, K.-R., & Tangermann, M. (2015). On the influence of high-pass filtering on ICA-based artifact reduction in EEG-ERP. 2015 37th Annual International Conference of the IEEE Engineering in Medicine and Biology Society (EMBC),

Zhang, H., Zhao, M., Wei, C., Mantini, D., Li, Z., & Liu, Q. (2020). Eegdenoisenet: A benchmark dataset for deep learning solutions of eeg denoising. arXiv preprint arXiv:2009.11662.

Zhang, Q., van Vugt, M., Borst, J. P., & Anderson, J. R. (2018). Mapping working memory retrieval in space and in time: A combined electroencephalography and electrocorticography approach. NeuroImage, 174, 472–484.

