## Supplementary Materials for "Introducing RELAX (the Reduction of Electroencephalographic Artifacts): A fully automated pre-processing pipeline for cleaning EEG data - Part 1: Algorithm and Application to Oscillations"

Results - Combined Sternberg, EO and EC resting data ... 20
Signal-to-Error Ratio ... 20
Artifact-to-Residue Ratio ... 21
Frontal Electrode Blink Amplitude Ratio ... 24
Blink Amplitude Ratio for All Electrodes ... 25
Proportion of Epochs Showing Muscle Activity After Cleaning ... 27
Severity by which the Log-Power Log-Frequency Slopes Exceed the Muscle Threshold ... 30
ICA Variance Explained by Brain Components ... 32
Proportion of Epochs Removed by Cleaning ... 34
Variance Explained by the Difference Between Eyes Open and Eyes Closed Resting. 36
Variance Explained by the Difference in Alpha Power Between WM Periods ... 42
SECTION FIVE ... 54
Analysis of a Combined EO, EC resting and 2back dataset ... 54
Signal-to-Error-Ratio ... 54
Artifact-to-Residue-Ratio ... 55
Blink Amplitude Ratio in Frontal Electrodes for the Combined 2back EO and EC Dataset ... 58
Blink Amplitude Ratio Across All Electrodes for the Combined 2back EO and EC Dataset ... 59
Proportion of Epochs Showing Muscle Activity Remaining After Cleaning from the Combined 2back, EO and EC Dataset ... 61
Severity by which Log-Power Log-Frequency Slopes Exceeded the Threshold from the combined 2back, EO and EC dataset ... 62
SECTION SIX ... 65
Analysis of a Colour-Wheel Recall task dataset ... 65
Signal-to-Error-Ratio in the Colour Wheel Task dataset ... 65
Artifact-to-Residue-Ratio in the Colour Wheel Task dataset ... 66
Blink Amplitude Ratio in the Colour Wheel Task dataset ... 69
Muscle Activity Statistics for the Colour Wheel Task dataset ... 71
SECTION SEVEN ... 75
Supplementary discussion points ... 75
Limitations and Potential improvements ... 76
SECTION EIGHT ... 79
Cleaned dataset examples ... 79
Supplementary Materials References ... 85

### LIST OF ABBREVIATIONS

| Acronym | Description |
|---|---|
| allBAR | All electrodes Blink Amplitude Ratios |
| ANOVA | Analysis of Variance |
| ARR | Artifact to Residue Ratio |
| ASR | Artifact Subspace Reconstruction |
| BAR | Blink Amplitude Ratios |
| CCA | Canonical Correlation Analysis |
| EC | Eyes Closed |
| EEG | Electroencephalography |
| EO | Eyes Open |
| ERP | Event Related Potentials |
| fBAR | Frontal Electrodes Blink Amplitude Ratios |
| FDR | False Discovery Rate |
| GFP | Global Field Potential |
| HAPPE | Harvard Automated Processing Pipeline for Electroencephalography |
| ICA | Independent Component Analysis |
| IQR | Interquartile Range |
| MAD | Maximum Absolute Deviation |
| MWF | Multiple Wiener Filtering |
| RAGU | Randomised Graphical User Interface |
| RELAX | Reduction of Electroencephalographic Artifacts |
| RMS | Root Mean Square |
| SER | Signal to Error Ratio |
| wICA | Wavelet Enhanced Independent Component Analysis |
| WM | Working memory |

#### SECTION ONE

##### Supplementary Background Points

Note that while eye blinks, eye movements, and muscle activity show stereotypical characteristics (making them reasonably easy to identify) small amplitude examples of these artifacts can be more difficult to distinguish from ongoing EEG activity, and the full extent of their influence is difficult to know [1-3]. Additionally, non-stationary electrooculogram (EOG) artifacts have been suggested to be not fully addressed by ICA, which does not include temporal information in its modelling [4]. In contrast, wavelet ICA (wICA) has the advantage of not requiring artifacts to be stationary [5].

As mentioned in the main text, the multi-channel Weiner filter (MWF) approach performs well at reducing temporary artifacts that can be identified in limited time windows, such as muscle activity, eye movement / blinks, and electrode drift. After cleaning with the MWF, the data primarily contains only smaller artifacts and most of the brain activity is preserved [6], allowing for optimal application of the ICA algorithm. The use of wICA instead of the typical approach of subtracting independent components means that reducing artifact components with wICA has a reduced chance of removing probable brain activity as well as the artifact.

Each step in our cleaning pipeline allows for the selection of multiple parameters, which can affect cleaning outcomes. During the design stage of our pipeline, we varied the selection of each of the parameters across the spectrum of potential values via considerable informal testing, to narrow down to the optimal outcomes in terms of metrics showing artifact reduction, the signal of identified non-artifact periods being as minimally altered as possible, and the variance explained by the experimental design being optimized across multiple large datasets and experimental designs. As such, we recommend use of the default parameters, but if future research demonstrates other parameters are superior, it is simple to adjust the selected parameters.

#### SECTION TWO

##### RELAX pipeline methods

###### *Filtering*

Firstly, data were loaded from a ".set" file in EEGLAB, and the electrode locations were specified (these are set by the user). Unused electrodes were deleted (these are also set by the user). Following the deletion of unused electrodes, a record was taken of the labels of the included electrodes, to enable interpolation of bad electrodes at a later stage. Data were 2nd order butterworth notch filtered from 47-53Hz (to address 50Hz line noise), and 4th order bandpass filtered from 0.25Hz to 80Hz (both are adjustable by the user).

###### *Bad Electrode Rejection*

Since both MWF and ICA are adversely affected by extreme outlying data, an initial step was undertaken to exclude extreme data, firstly by electrode, then by time period. For the bad electrode rejection step, first, PREP's "`findNoisyChannels`" function was used in an initial approach to remove bad electrodes [7].

A secondary bad electrode rejection was then implemented, which involved first epoching the data into 1 second epochs with a 500ms overlap. Additionally, eye blinks were marked in the continuous data through a multi-step process (detailed below). Epochs including blinks were excluded from the initial epoching and collected as a separate list of epochs for 1 second around the blink maximum (this ensured blinks were not excluded as extreme outliers). To detect blinks, firstly data were bandpass filtered using a fourth order butterworth filter from 1-25Hz. Then we averaged the pre-specified blink affected electrodes ('FP1'; 'FPZ'; 'FP2'; 'AF3'; 'AF4'; 'F3'; 'F1'; 'FZ'; 'F2'; 'F4'). Blinks were marked as the maximum point within each time period that exceeded a value of the upper quartile + interquartile range (IQR) * 3 when all voltages were included.

A matrix of electrodes x epochs was created for: i) the total voltage shift within the epoch, ii) for the maximum absolute voltage, (with each of these performed separately for both the typical epochs and the blink affected epochs) and iii) for the log-power log-frequency slope for each electrode. Cells in this electrode x epoch matrix that exceed 20 median absolute deviations (MAD) from the median in max - min voltage shift within the epoch, or 3 MAD from the median in max - min voltage within the epoch from the epochs contaminated by blinks were marked as extreme outliers. The log-power log-frequency slopes were computed for each epoch using fieldtrip's ft_freqanalysis (set to mtmfft with hanning tapers for a frequency range of 1-75Hz with netpow2 padding for all time periods with a 0.05s resolution) then MATLAB's polyfit function. Log-power log-frequency slopes steeper than -4 were marked as extreme outliers (this measure indicated epochs showing no brain activity, only voltage drift). Epochs with absolute voltages of more than 500 microvolts, or less than 2 microvolts in max - min (suggesting a dead electrode) were also marked as extreme outliers. EEGLAB's `pop_rejkurt` function was used to detect kurtosis values > 8, and `pop_jointprob` function was used to detect improbable voltage distributions >8 (with the same threshold for both single electrodes and all electrodes), and epochs with values exceeding these thresholds were marked as extreme outliers.

Electrodes that showed extreme outlier epochs affecting more than 5% of the data were removed. However, we imposed a limit of rejecting 20% or fewer of the total number of electrodes. If data still contained more than 80% of the original electrodes after PREP's noisy electrode rejection, then electrodes showing >5% of epochs contaminated by these extreme artifacts were rejected. If more electrodes than 20% were contaminated by extreme artifacts in this manner, electrodes were ranked according to the number of epochs showing values that exceed the extreme thresholds, and the 20% of electrodes that showed the highest number of epochs marked as extreme outliers for the specific file were rejected.

Next, within the remaining electrodes, a log-power log-frequency slope was computed from 7 to 75Hz. Slopes >-0.59 are suggestive of muscle activity, as slopes from epochs within typical EEG recordings show values above this threshold, while almost no epochs within EEG recordings taken from people who have had their muscles pharmacologically paralysed show slopes above this threshold [8]. As such, using this threshold obtained from EEG recordings taken from individuals with paralysed scalps means that very little of the distribution of EMG free EEG activity would be marked as containing muscle activity. If fewer than 20% of electrodes had been rejected for being extreme outliers at this stage, electrodes that show more than 5% of epochs contaminated by muscle (log-power log-frequency slopes >-0.59) were marked for rejection. As with the extreme outlier detection steps, if the muscle activity electrode rejection step recommended electrode rejections such that more than 20% of total electrodes would end up being rejected, the electrodes were automatically ranked by the proportion of epochs showing muscle activity above the threshold, and only the most severely affected electrodes were rejected, within the limit of a total of 20% of all electrodes being rejected (including all three electrode rejection steps: PREP's “`findNoisyChannels`“ function, the extreme period rejection methods, and the muscle activity rejection method).

##### *Extreme Outlier Marking*

After these bad electrode rejection steps, the extreme outlier rejection approaches were re-computed (without the influence of bad electrodes, and not including the muscle activity detection method) and these extreme outlier detections were used to mark extreme epochs in the EEG data, using the same criteria as explained above (muscle activity was not marked at this stage, and was left to be cleaned by the MWF and wICA steps). The time periods for each of these extreme outlier epoch periods were marked with NaNs in the MWF cleaning template, which is used by MWF to identify artifact and clean periods, as well as periods to ignore in its model (marked with NaNs) (the MWF template is described further below). These extreme outlier periods were also rejected from the data completely prior to the ICA computation step (explained below).

After the extreme outlier rejection steps, three steps of MWF cleaning were implemented to address different artifacts, firstly muscle activity, secondly blinks, and thirdly, horizontal eye movements and single electrode drift.

##### *MWF1 – Muscle Activity*

Firstly, a 1D matrix that represented a template of artifact periods (marked as 1), clean periods (marked as 0), and extreme periods (for the MWF cleaning to ignore, marked as NaN) was constructed, which matched the length of the continuous data file. NaNs were also applied into the first and last 5.5 seconds of the data, as these periods often contained artifacts, or were affected by filtering edge effects, so were ignored in the MWF templates and removed from the data prior to the wICA. The MWF template was constructed using the following approach:

To begin with, for each electrode, epochs that were affected by muscle activity were detected (defined by log-power log-frequency slopes >-0.59). This was achieved by performing a fast Fourier transform (ft_freqanalysis using the mtmfft setting, with a hanning taper) on each 1 second epoch, then computing the log-power log-frequency slope (using the MATLAB function "`polyfit`") of data from 7 to 75Hz for each epoch separately [8]. In this electrode x epoch matrix, NaNs replaced all values <-0.59 (which reflected slopes not indicative of muscle activity, so these values were ignored by the algorithm), and the -0.59 threshold was subtracted from all remaining values (so all values are made positive, with 0 as the putative threshold value). The values from these computations for each electrode were summed within each epoch, to provide a value reflecting the amount that each epoch showed muscle activity exceeding the -0.59 threshold, cumulated across channels. From these values, to increase the resolution of the artifact template, we took advantage of the 500ms overlap included in each epoch, which meant that the first half of the second epoch overlapped with the second half of the first epoch. Odd and even numbered epochs from the matrix of epoch x electrode slope data were separated. Then a template of data with the same duration as the continuous data was constructed from the single log-power log-frequency slope values from each individual epoch for the odd and even numbered epochs separately. This was done by taking the single value by which each epoch exceeded the slope threshold and extrapolating this value across the 1000ms of timepoints that the epoch represented in the length of the continuous data (so epoch 1 lasted 1-1000ms, epoch 2 lasted 500-1500ms, epoch 3 lasted 1000-2000ms etc.). The last 500ms of the matrix from the odd epochs, and the first 500ms of the matrix from the even epochs were marked as NaN, as no values were available in these periods from the odd and even epochs respectively. These two 1D matrices (which were of the same length as the continuous data and containing muscle slopes for the odd and even numbered epochs) were then averaged, creating a full-length mask of the cumulative amount by which each epoch exceeded the slope threshold, with 500ms resolution of the cumulative slope exceeding values within each time period (since the epochs overlapped by 500ms). The periods marked as extreme outliers in the previous section were marked as NaNs in this template. After this process, all time periods where an electrode showed a muscle slope that exceeded the -0.59 threshold were marked as part of the artifact template for cleaning by the first MWF step. If more than 50% of the total time-period was marked as artifact, then only the 50% most severely affected time periods were marked as artifact, which was calculated from this 1D continuous data matrix.

Our initial testing indicated that very brief artifact or clean template periods impaired the ability of MWF to clean the data. To ensure clean and artifact masked periods were of sufficient length, artifact periods shorter than 1200ms were padded with artifact marks (1's) equally on each side to reach 1200ms in length. It has been recommend that it is better to be liberal in marking around artifacts, as the MWF cleaning is more effective when potentially clean periods are included as artifacts compared to when artifacts are marked as clean periods [6]. Clean periods that lasted less than 1200ms and had artifact markings on each side were also marked as artifacts. Finally, any periods remaining that were shorter than 1200ms after these two steps were marked with NaNs so they would not be included as artifacts or clean periods in the MWF template (but would still be cleaned by the MWF cleaning).

To ensure the MWF approach had sufficient artifact example data to work with, we required a minimum of 5% of the data to be marked as artifact before MWF cleaning was applied. If less than 5% of the data was marked as artifact, the MWF cleaning step was ignored, and we left the artifacts for the later wICA cleaning step (note that if the wICA was not effective at cleaning these periods, that are typically excluded in an epoch rejection step). If more than

5% of the data was marked as artifact, the clean and artifact template was then submitted to the MWF cleaning, with a delay period set to 8. This delay period was a positive and negative time lag for 8 samples from each timepoint, which turns the MWF into a finite impulse response filter, allowing the MWF to clean the data based on both spatial and temporal information. The RELAX script was set to detect generalised eigenvector deficiency. This sometimes occurs when MWF cleaning is applied, particularly for longer delay periods when data has been filtered (which creates a degree of temporal dependence in the EEG data, reducing the rank of the data obtained when covariance is assessed across timepoints). Generalised eigenvector deficiency can impair the ability of the MWF to clean the data, so the reduction in delay period then repetition of the MWF cleaning ensures this does not lead to inadequate data cleaning. If generalised eigenvector deficiencies were detected, the algorithm went back to the data prior to the initial MWF application, reduced the delay period by a value of 1, then ran the MWF cleaning again. This approach was repeated up to 3 times, resulting in a minimum delay period of 5. The algorithm records the filename of all files that still show generalised eigenvector deficiency at the minimum delay period of 5. This enables the user to inspect these files and determine the reason for the issue or exclude the file as bad data if visual inspection indicates no usable EEG activity. None of the files included in our analyses showed this issue. Note also that if low frequency data is removed by using robust detrending [9] instead of filtering, no temporal dependence is created, and higher MWF delay periods can be used. However, the optimal parameter settings for robust detrending are not yet established, can vary considerably, and may be data dependent. As such, we have not implemented the method in our pipeline, but this is something we intend to explore further.

***MWF2 – Blink Activity***

Secondly, a blink artifact mask was created by marking the 800ms surrounding all blink maximums as artifacts (recall that the continuous data were marked for blink maximums in the initial extreme outlier detection step, so these marks were used). Similar to the approach used for the muscle cleaning step, we ensured a minimum clean period between blinks of 800ms and marked the clean periods that did not last this length as artifacts. This is because blinks are often brief, and unlikely to have lagging undetected edge effects, in contrast to the muscle detection, where muscle activity may extend for brief periods outside of the 1 second epochs used, yet not be detected due to the 1 second epoch used to calculate the log-power log-frequency slopes. Similar to the muscle artifact template, the NaNs from the extreme artifact step were added to the blink artifact template. As per the muscle activity cleaning, the MWF template was used to clean the data with a delay period of 8 (which was reduced up to three times in the case of generalised eigenvector deficiency).

If less than 5% of data was marked as artifact in the blink cleaning step, the second MWF cleaning step was not performed, and instead the blink artifact mask was added to the third MWF cleaning round, which cleaned horizontal eye movements and drift. Blinks are characteristically more similar to horizontal eye movement and drift than muscle activity, so blinks could be included with the third MWF cleaning step, while muscle activity was not included as our initial testing suggested doing so decreased the efficacy of the third MWF cleaning step. The addition of muscle templates that were not included in the initial MWF cleaning step because less than <5% was marked as artifact was not implemented for muscle activity, as we deemed it to be acceptable to miss even two rounds of MWF cleaning, since wICA-ICLabel cleaning alone also cleaned the data reasonably well (described later). As such, the RELAX pipeline has a useful redundancy in the cleaning process, with wICA cleaning remaining artifacts that MWF cleaning might have missed, while also not reducing the signal if no artifacts are detected.

##### *MWF3 – Horizontal Eye Movements and Single Electrode Drift*

Third, horizontal eye movements and single electrode drift were identified and cleaned with the MWF. In order to identify single electrode drift that was not resolved by initial filtering, data were low pass filtered using a fourth order acausal Butterworth filter at 5Hz (to ensure high power alpha oscillations were not inadvertently marked as drift) and re-referenced using PREP's robust average re-referencing approach, which was only used for the detection of single electrode drift and construction of the artifact template at this stage, and not applied to the data prior to the 3$^{rd}$ MWF cleaning step. Epochs showing an amplitude at any electrode that was more than 10MAD from the median of all electrodes after this average re-referencing were deemed to be affected by drift and marked as artifact periods (this was adapted from [10] who used SD instead of MAD to set the threshold). Periods where the pre-specified horizontal electrooculogram (HEOG) affected lateral electrodes (defined shortly) showed more than 2MAD from the median of their typical amplitude, with opposite voltage movement polarity on the opposite sides of the head (reflecting horizontal eye movements) were assumed to reflect horizontal eye movements and were also marked as artifact periods [11]. The HEOG affected electrode can be set by the user, with an order of preference of electrode to be used by the algorithm specified by the user. A list of electrodes was used because the lateral electrodes that are often affected by HEOG are often also excluded as bad due to their proximity to temporal muscles, which generate large artifacts in these electrodes. In the current study, the electrodes were listed in order as: "AF7", "F7", "FT7", "F5", "T7", "FC5", "C5", "TP7", "AF3" for the left side, and "AF8", "F8","FT8","F6","T8", "FC6", "C6", "TP8", "AF4" for right side electrodes. As with the muscle cleaning template, artifact periods were padded if shorter than 1200ms. Clean periods that lasted less than 1200ms and had artifact markings on each side were also marked as artifacts. Finally, any periods remaining that were shorter than 1200ms after these two steps were marked with NaNs (so they would not be included as artifacts or clean periods in the MWF template). Similar to the previous two MWF steps, the NaNs from the extreme artifact step were added to the horizontal eye movement and drift artifact template. Data were then cleaned with the MWF algorithm using this artifact and clean period template (and a delay period of 8, reduced 1 value at a time if necessary due to generalised eigenvector deficiency to a minimum of 5 as per the preceding MWF cleaning steps).

##### *wICA Applied to Artifact Components Identified by ICLabel*

After the data had been cleaned by the three sequential MWF steps, data were first average re-referenced using PREP's robust re-referencing, which avoided asymmetry as a result of rejected electrodes affecting the average re-referencing, but still excluded the previously rejected electrodes from the data [7]. This approach also added the online reference back into the data prior to average re-referencing which prevents rank issues, then removed that online reference from the data [7]. As mentioned earlier, the periods that were marked as extreme outliers in the initial cleaning steps were rejected at this stage, along with the first and last 5.5 seconds of the data. ICA was then computed using one of three ICA algorithms, which each use different approaches to separating the scalp level signal into its putative underlying source signals, and have shown differences in cleaning outcomes [12]. We tested the application of fastICA (with the deflation setting implemented to avoid non-convergence issues), cudaICA, or AMICA (all three methods were used, cudaICA results are reported in the main manuscript, and all results including fastICA and AMICA are reported in these supplementary materials). ICLabel was used to identify artifactual components (defined as components that ICLabel marked as more likely to be any of the artifact categories than to be produced by the brain). These artifactual components were then reduced with wavelet enhanced ICA (wICA) with the default settings (mult = 1, L=5, wave='coif5') [5]. The non-artifactual components were left as they were, with no modification by wICA. Finally, after the wICA reduction of artifactual components, the continuous data was reconstructed back into the scalp space.

The above steps left cleaned, continuous data, which could be epoched for different types of analyses. Some of our metrics required continuous data, so for these metrics we used this continuous data (for example, SER and ARR values, and the proportion and strength above threshold of 1 second epochs across the whole data that showed log-power log-frequency slopes reflective of muscle activity). Other metrics required epoched data (the total proportion of data rejected by cleaning, the variance explained by the experimental manipulation). In order to obtain the epoched data, we used a typical approach of interpolating the rejected electrodes back into the data (using EEGLAB's "`pop_interp`" function with a spherical approach), and rejected epochs based on max-min voltage values >60 microvolts, or kurtosis / improbable data for all electrodes >3 or any electrode >5.

##### *Parameter Selection Notes*

The 2nd order Butterworth notch filtered from 47-53Hz, and 4th order bandpass filtered from 0.25Hz to 80Hz filter settings can be adjusted by the user. We found that EEGLAB's default filter seemed incompatible with the MWF cleaning (leading to common eigenvector deficiencies and cleaning artifacts being introduced into the data), so we do not recommend the use of EEGLAB's default filter. It may be that a more sophisticated filtering approach such as robust detrending or the trial masked robust detrending could be superior (particularly as these methods would avoid eigenvector deficiencies as they do not create any temporal dependencies) [9, 13]. Unfortunately, our preliminary tests indicated robust detrending with 3 second windows and an order of 5 produced worse artifact cleaning and less variance explained by experimental manipulations than high pass filtering at 0.25Hz, even when we took advantage of the lack of temporal dependencies and used higher delay periods for the MWF cleaning. As such, we have not applied robust detrending in the current version of RELAX.

With regards to the selection of a log-power log-frequency slope threshold used to detect muscle activity, the -0.59 threshold performed best in our piloting of the pipeline. This -0.59 slope threshold reflects the point where the histogram of slopes from the paralysed scalp EEG recordings crosses the histogram of non-paralysed scalp recordings, such that only a minimum of non-muscle affected epochs will be included in the muscle cleaning template, while all muscle contaminated epochs would be included in the cleaning template [8]. However, the value can be adjusted to be more or less stringent. A -0.72 threshold would only accept log-power log-frequency slopes similar to the EEG data obtained from individuals who have their scalp muscles pharmacologically paralysed (and reject slopes outside of this paralysed scalp range), while -0.31 would avoid marking any data showing a similar slope to the paralysed dataset as muscle activity [8]. Also, note that during the construction of RELAX we informally but extensively tested whether BLINKER [14] would be an effective objective tool for detecting eye blinks in our blink cleaning step of the MWF. Unfortunately, we found that BLINKER missed a considerable proportion of blinks, or alternatively sometimes marked large alpha oscillations as blinks. As such, we found that the IQR method to detect blinks performed better, so implemented the IQR method in our pipeline.

In addition to the cleaning method explained above, we have provided a quick visual check of potential outlying data across all participants cleaned by RELAX. The code to produce the OutlierParticipantsToManuallyCheck values after the epoch rejection script takes the maximum value minus the minimum value from each epoch within each participant, then calculates the median amount of voltage shift within epochs for each participant at each electrode. This distribution is usually skewed (with some participants showing very large values due to large alpha activity, but most participants showing a smaller value), so the data is log transformed. The upper threshold for detecting outliers was set as 'the 75% point in the distribution + 2 * IQR' and the lower threshold was set as 'the 25% point in the distribution - 2 * IQR'. A line graph of these values for each electrode and each participant is output after the epoching script to prompt users to visually inspect participant data that is specified as an outlier. We found the approach to provide a quick and simple prompt for the user to check whether outlying participant data might be just irretrievably corrupted (the code was written because one of the authors had one participant file with no neural activity in their data due to technical faults in the EEG session, but this issue was not initially detected by the cleaning pipeline, and the outlier check effectively highlighted this file for further inspection).

Lastly, note that the canonical correlation analysis (CCA) method described for the comparison methods in the main text can be applied after MWF or wICA, but not after ICA subtraction, as CCA requires full rank data, and ICA subtraction reduces the rank of the data.

#### SECTION THREE

##### Comparison Pipeline Description

For the primary analyses (of the combined Sternberg and resting data), cudaICA was the ICA method used within the comparison pipelines, and for the other datasets (reported only in these supplementary materials), fastICA was used (as these data were processed on Apple Mac computers without cudaICA compatible graphics cards).

The first comparison pipeline we included was the Harvard Automated Processing Pipeline for EEG (HAPPE [15]). This pipeline filters the data with a 1Hz high pass filter and uses CleanLine to remove 50 or 60Hz line noise [16]. It then uses a joint probability of the average log-power from 1 to 125 Hz to reject bad electrodes (with probabilities of >3SD from the mean), which is performed twice. wICA is then applied to all components. This wICA was proposed by the authors to reduce initial high amplitude artifacts, prior to another ICA run being conducted, upon which the Multiple Artifact Rejection Algorithm (MARA) is used to identify and reject remaining artifacts [17]. After both the wICA and ICA cleaning have been applied, bad electrodes are interpolated back into the dataset, and the data are re-referenced to the average reference. A full description of the algorithm can be found in [15].

The second comparison pipeline we included was the artifact subspace reconstruction (ASR) method [18], followed by ICA subtraction using ICLabel. ASR is an automatic approach similar to principal component analysis-based methods where components with large variance are rejected, but with an additional automatic identification of clean data to determine thresholds prior to this subtraction, followed by reconstruction of the original electrode space data. ASR is additionally non-stationary (i.e., it takes temporal information into account) in contrast to the stationary approach implemented with ICA. Further details of the ASR approach can be found in [18]. Since the ASR approach does not specify particular extreme outlier rejection steps prior to ASR cleaning, we applied our initial RELAX extreme outlying electrode and period rejection, prior to cleaning the data with ASR in order to maximise comparability. ASR also enables a number of parameter selections. Of particular importance is the “burst criterion”, which was set at SD = 20 as per the suggested optimum [18]. We set the “WindowCriterion” to 0.25, and the “WindowCriterionTolerances” to [-Inf 7] as per previous recommendations [18]. Additionally, we turned off the “BurstRejection” setting, as extreme outliers had already been rejected, and during initial piloting we found that leaving this parameter on lead to a very high number of data periods rejected following the ASR. Similarly, we turned off the “FlatlineCriterion”, “ChannelCriterion”, “LineNoiseCriterion”, and “HighPass” criterion, as all these had already been addressed by our earlier processing steps. After the ASR was implemented, ICA was computed, and components were rejected with ICLabel to maximize comparability with the RELAX pipelines.

The third comparison approach we used is perhaps one of the most commonly implemented: simply rejecting outlying data first (as per the approach used in the initial steps of our RELAX pipeline), computing ICA, and subtracting the components identified as artifacts, then reconstructing the electrode space data [19]. We implemented this using ICLabel to identify artifactual components [19]. We refer to this pipeline throughout as ICA_subtract.

The fourth comparison method was identical to ICA_subtract, but instead of simple ICA subtraction on artifact components, it applied the wICA approach to all components (as per [5]). We refer to this pipeline throughout as wICA_all.

A similar approach was used in our fifth comparison pipeline, but instead of applying wICA to all components, wICA was applied only to components identified as artifacts by ICLabel. Although a similar approach of applying wICA to only artifact components has been previously implemented [20, 21], as far as we are aware, this is the first time it has been tested by selecting components with ICLabel. We refer to this pipeline throughout as wICA_ICLabel.

Our sixth comparison pipeline implemented only a sequential MWF cleaning identical to the MWF cleaning steps in our RELAX pipeline but did not apply any additional cleaning after the MWF stage (no wICA unlike the RELAX methods). This is similar to the approach used by [6], with the extension of applying their suggested sequential MWF cleaning to clean multiple different categories of artifacts. We refer to this pipeline throughout as MWF_only.

Our seventh pipeline used the sequential MWF cleaned data, but instead of applying wICA to this data as per the RELAX methods, it used the extended canonical correlation analysis (CCA) to further clean any remaining muscle artifacts [22]. CCA separates the EEG data into components that are not correlated with each other but are maximally autocorrelated at a lag of one datapoint. Muscle activity is characterised by a similar pattern to white noise, with a low autocorrelation (in contrast to neural activity, which shows voltage fluctuations at a slower rate with higher autocorrelation). As such, CCA is an effective method for identifying and removing muscle activity from EEG and has been suggested to be superior to ICA methods [23, 24]. Recently CCA has been improved through the use of the log-power log-frequency slope thresholds identified by the comparison of paralysed and non-paralysed scalp EEG recordings to detect probable muscle contaminated components for removal [22]. This approach removes components with a one timepoint-lag autocorrelation of less than 0.19 and log-power log-frequency slopes of more than -0.48 [22]. We use this extended CCA here and refer to this method throughout as MWF_CCA.

Similarly, our eighth comparison pipeline was identical to MWF_wICA, except that muscle components were not cleaned in the wICA cleaning step. Instead, CCA was implemented after the wICA step in order to address any remaining muscle components. This method was referred to throughout as MWF_wICA_CCA.

Lastly, we tested a few modifications of our RELAX pipeline. These included using different ICA algorithms, namely, infomax [25] (MWF_wICA_infomax), fastica [26] (MWF_wICA_fastICA), or AMICA [27] (MWF_wICA_AMICA), subtracting artifactual ICA components instead of using wICA (MWF_ICA_subtract), and low pass filtering at 45Hz prior to implementation of the ICA algorithm (MWF_wICA_45Hz) (which has been suggested to improve ICA decomposition [28]).

**Cleaning Quality Evaluation Metrics**

In order to ensure we assessed the different cleaning pipelines fully for effectiveness at cleaning both the range of potential artifacts, and the ability for the pipelines to not over-clean the data, we tested the pipelines across six different types of metrics. These metrics have all been used by previous research.

##### *The Signal-to-Error Ratio*

The Signal-to-Error Ratio (SER) and Artifact-to-Residue Ratio (ARR) are complimentary metrics that were used to assess any potential distortion of the clean EEG periods by the cleaning pipelines (SER), and the amount of artifact that was reduced by the cleaning approach (ARR). These measures have been used by previous research to compare cleaning effectiveness across MWF and ICA approaches [6, 29, 30].

The SER was calculated from segments of the data marked as free of artifacts by the automatic artifact detection approaches implemented in the RELAX pipeline. The SER is calculated on each electrode (*i*) first by obtaining the expected value operator (which is analogous to the weighted average, where more probable values are given stronger weights when computing the average) of the square of the signal in the “raw” (not yet cleaned) data across all periods marked as clean ($y_i$), then dividing this value by the expected value operator of the square of the signal that was removed by the cleaning pipeline across the periods marked as clean, then multiplying this value by the log10 of 10 ($\hat{d}_i$) (see Equation S1) [6, 29, 30]. Note that the “raw” data used in the calculation of the SER was obtained after data had been filtered and extreme outlying electrodes and periods had been rejected (and before any of the MWF cleaning steps were applied). In order to obtain a single measure for each cleaned dataset, the SER from each electrode is combined by weighted averaging over all electrodes (Equation S2) [6, 29, 30], with the weighting performed by the proportion of artifact power an electrode produces relative to the artifact power from all other electrodes ($p_i$) (estimated by subtracting the power in the clean segments from the power in the artifact segments, Equation S3) [6, 29, 30]. This has the effect that electrodes containing the most artifact contribute the most to the final SER value for that dataset. This approach ensures SER values appropriately reflect the contribution of noisier electrodes and protects the measure against high SER values being produced by mostly clean data with a single electrode which is very noisy in artifact periods and inadvertently distorted by the cleaning process in the clean periods.

Equation S1:

$$SER_i = 10log_{10} \frac{E\{(y_i)^2\}Ho}{E\{(\hat{d}_i)^2\}} \ (for\ clean\ segments)$$

Equation S2:

$$SER = \sum_{i=1}^{M} \quad p_i \ SER_i$$

Equation S3:

$$p_i = \frac{E\{(y_i)^2\}\ (artifact\ segments) - \ E\{(y_i)^2\}\ (clean\ segments)}{\sum_{i=1}^{M} \quad (E\{(y_i)^2\}\ (artifact\ segments) - \ E\{(y_i)^2\}\ (clean\ segments))}$$

Since the EEG periods that are marked as clean by the automated approach do not include blinks, muscle activity, horizontal eye movements and drift (nor do they include extreme artifacts, which were marked as NaNs in the clean/artifact template) these segments should be minimally modified by the cleaning pipelines. As such, high SER values are expected if cleaning has left the non-artifact periods undistorted, so high values indicate good performance [6, 29, 30].

***The Artifact-to-Residue Ratio***

The ARR was calculated from the periods of the data marked as artifact by the automatic artifact detection approaches implemented in the RELAX pipeline. As with the SER, the calculation of this measure was performed first on individual electrodes by obtaining the expected value operator of the square of the removed artifact ($d_i$), divided by the expected value of the square of the total signal from the artifact periods ($\hat{y}_i$) from the "raw" (not cleaned) data ($y_i$) (when ARR is calculated on real data where the true artifact signal is not known), then multiplying this total by the log10 of 10 (Equation S4) [6, 29, 30]. To obtain a single value for each dataset, the individual electrode values were then combined via weighting in the same manner as the SER (weighting via $p_i$). As such, the ARR provides large values when more artifact is removed relative to the original data (as denominator of the equation: the "raw" data minus the artifact: $[y_i - \hat{d}_i]$ will be as small as possible) and is valid when artifacts are high in amplitude relative to the clean data (as per the artifacts selected by the MWF artifact template in the current study, which are mostly based on outlying amplitudes or artifacts that are typically large in amplitude, and are comprised of eye movements, muscle activity, and drift). Note that the "raw" data used in the calculation of the ARR was obtained after data had been filtered and extreme outlying electrodes and periods had been rejected (and before any of the MWF cleaning steps were applied).

Equation S4:

$$ARR_i = 10log_{10} \frac{E\{(d_i)^2\}}{E\{(y_i - \hat{d}_i)^2\}} \ (for\ artifact\ segments)$$

The units for the SER and ARR measures are decibels (dB) and should be evaluated together - higher values for both simultaneously reflects successful cleaning, with high amounts of artifact power removed and clean periods undistorted. Low SER values and high ARR values are likely to indicate effective artifact removal but distortion of the clean signal, and low ARR values and high SER values are likely to indicate ineffective artifact removal [6, 29, 30].

It is worth noting that ICA approaches may detect and remove artifacts other than the most common artifacts captured by our MWF cleaning templates. As such, ICA approaches may seem to "distort" clean periods using the SER metric. As such, it is more appropriate to compare across pipelines that apply cleaning to all periods (such as those that implement ICA, ASR, or CCA), and perhaps not appropriate to compare those pipelines to the MWF_only approach (which does not detect artifacts in the clean periods at all). For this reason, we have used a number of metrics additional to the SER and ARR, in order to fully characterize artifact reduction (with the blink amplitude ratio, artifacts remaining showing muscle activity, and variance explained by brain activity detected by ICLabel after cleaning) and preservation of signal (with the measures of variance explained by the experimental manipulation).

Additionally, because the SER and ARR metrics are based on the variance in the clean and artifact periods, it is possible for the metrics to be biased by co-occurrence of low-powered brain signals during the artifact periods more commonly than the clean periods. For example, at times the clean periods may have showed more alpha activity for example (and thus high variance), while the muscle affected periods show less alpha activity and only low powered muscle activity. In this case, sometimes the variance of the artifact periods may have been less than the variance of the clean periods, leading to very low ARR and very high SER values. In fact, because individual electrodes within this metric are scaled by the amount that the artifact period variance exceeds the clean period variance, with electrodes showing higher clean variance than artifact variance set to zero before the weighting based on amount of variance in each electrode (by dividing electrodes variance by the total of all electrodes), it is possible for all electrodes to be set to zero, and the SER and ARR to produce NaN values. As such, this metric is perhaps less ideal for evaluating files where only small amplitude muscle activity is present (but is well suited to evaluating blink activity or high-power muscle activity, which is almost always higher in amplitude than the non-blink/non-muscle periods). In order to address this issue, we have also used muscle activity artifact specific metrics (described in the following sections).

##### *The Blink Amplitude Ratio*

The third metric we used was the blink amplitude ratio (BAR) [31]. This metric provides a ratio of the absolute amplitude within periods marked as blinks to the periods on either side of the blink. When applied to cleaned data, the measure provides a good indication of whether the cleaning pipeline has effectively cleaned the blink (leading to values ~=1). Alternatively, the metric indicates if the blink has been under-cleaned (leading to values of >1), or the subtraction of a blink artifact component has included the influence of brain activity as well as blink related activity, so that the subtraction creates a negative deflection where the blink was previously (also leading to values >1 due to the absolute transform). The metric also indicates if blinks are over-cleaned so that both positive and negative signals have been reduced towards zero (leading to values <1). To compute BAR, we epoched data for 4 seconds centered on the blink maximum and excluded epochs that included more than one blink within this 4 second epoch (to prevent these additional blinks from influencing the baseline period). We baseline corrected the epochs by subtracting the average of the first 500ms and last 500ms of the epoch. We then performed an absolute transform on all data in the epoch, then divided the mean of the 1 second centred on the blink maximum by the mean of the first 500ms and last 500ms of the epoch. For analysis, we examined both the frontal BAR (fBAR), which was the average BAR across electrodes FP1, FPz, FP2, AF3 and AF4, and the average BAR over all electrodes (allBAR).

##### *Log-frequency Log-power Slopes Indicating Muscle Activity*

Next, we examined the proportion of 1 second periods that contained any electrode with likely muscle activity after the cleaning pipeline, using the log-power log-frequency slope threshold of -0.59 [8]. We also examined the severity by which muscle activity slopes in these epochs exceeded the threshold by subtracting -0.59 from the log-power log-frequency slope values from all epoch / electrode datapoints that exceeded this threshold, then averaging the remaining values. This provided a value reflecting the average amount the slopes exceeded the log-power log-frequency slope threshold of -0.59 in the epochs and electrodes that were not completely cleaned of muscle activity. It is worth noting that this measure could be a misleading metric of the impact of muscle related artifacts for a minority of cleaned files. The metric is calculated only from epochs that show muscle slopes above the threshold. As such, if only a single epoch is still affected, but that one epoch shows a very severe muscle artifact, the metric will provide a very high score for that file, but the impact of the artifact on experimental measures may be very low. However, across the large number of files included in our analysis, the effect of such outliers is minimal (particularly when using the robust statistics, which exclude these outliers when calculating statistical effects).

##### *ICA Variance Categorized by ICLabel*

We examined the amount of ICA variance attributed to components categorized as brain activity by ICLabel, computed by summing the amount of variance in the EEG data explained by components categorized as brain by ICLabel (after cudaICA, defined as components that were deemed by ICLabel to be more likely to be any artifact category than to be brain activity). Variance was calculated for each component individually using `compvar` (EEGLAB). An absolute transform was performed on the value of variance for each component to ensure all components provided a positive value for the amount of variance the component contributed to the data. This was done because compvar provides negative values if a component influences the data in the opposite direction to the overall trend. However, for our purposes we were only interested in the percentage of total variance of the data that was influenced by brain activity. As such, negative variance values were made positive with this absolute transform so that their influence would not reduce the sum of brain activity component variance or artifact activity component variance, and the total values from all components would be equivalent to 100%. Following this, the variance from all brain components was summed and the variance from all artifact components was summed. The summed variance for brain activity was divided by the sum of the total brain variance and total artifact variance and multiplied by 100 to obtain a percentage of the variance explained by brain activity (as determined by ICLabel). Note that methods that subtract components such as ICA_subtract and CCA were excluded from this measure, as component subtraction completely removed any variability from that artifactual component, so the contribution to variance from artifactual components would be 0 for these artifacts from these methods. Since this metric is only applicable for some pipelines, we only report the results of these analyses in the supplementary materials. It should be noted that infomax (with cudaICA) was used to compute the ICA artifact components for selection by ICLabel. Given the use of a common ICA method for this metric across all pipelines, this approach may have biased the metric towards the pipelines using infomax and against the fastICA / AMICA pipelines.

##### *Proportion of Epochs Rejected*

We examined the proportion of total epochs that were rejected by the cleaning pipeline, after both excluding outlying data in the initial pre-cleaning steps and rejecting outlying epochs in the final stage prior to data analysis. In order to achieve this, we epoched the resting data into 5 second intervals with a 3 second overlap (a typical approach to enable analysis of resting state oscillatory power or connectivity, with the overlap providing a buffer against edge effects so that the middle 2 seconds of the epochs that are analysed provide valid estimates of oscillatory power or connectivity). For the Sternberg task, we epoched data for 15.8 seconds around the onset of the probe stimuli (10.3 seconds prior to 5.5 seconds after), in alignment with typical analyses involving this task [32]. For the cleaned data epochs, we rejected outlying epochs using a typical approach with an automated algorithm rejected epochs that still showed potential artifacts, as defined by kurtosis or improbable data with a value higher than 5 SD from the mean at any electrode or 3 SD from the mean for all electrodes (using the relevant EEGLAB functions `pop_rejkurt` and `pop_jointprob`), or epochs showing values outside of a -60 to 60 microvolt window [32]. We obtained these epochs for both the raw data (which did not have any epochs removed) and the cleaned data (after data had been removed by the cleaning process, and by the rejection of remaining outlying epochs). We then calculated the proportion of cleaned epochs available for analysis against the total number of epochs that were present in the raw data.

##### *Variance explained by Experimental Manipulations*

Lastly, we examined the amount of variance explained by a number of different experimental manipulations to test the real-world applicability of the pre-processing pipelines. Ideally, effectively cleaned EEG data should lead to data that still contains all of the brain signal, and none of the artifact. Data cleaned this effectively should in theory produce the largest amount of variance explained by different experimental manipulations. This is because we assume that non-neural artifacts are unrelated to different experimental manipulations (so their inclusion in the data would contribute noise to a comparison between two experimental conditions, reducing the variance explained), and neural activity *is* related to the experimental manipulation (so maintaining more of the neural activity leads to increased detection of the effect of the experimental manipulation on brain activity and thus more variance explained by the experimental manipulation). As such, the best cleaning pipelines should provide the maximal between condition effects [33].

In order to test the amount of variance explained by the different experimental manipulations, we used the randomization graphical user interface (RAGU) [34, 35], which analysed all electrodes and timepoints available for ERP or oscillatory power data using randomisation statistics (while controlling for multiple comparisons in both the temporal and spatial dimensions). This toolbox additionally provides the ability to test for differences in overall neural response strength (using the global field potential (GFP) or root mean squared (RMS) test) and separately to test for differences in the distribution of neural activity (using the TANOVA, which compares global field potential dissimilarity maps between conditions after the recommended L2 normalisation for differences in GFP or RMS). More details on this toolbox can be found in [34, 35]. Given the potential computation time when including all statistical tests, 1000 permutations were used for all tests within RAGU.

We computed explained variance for data from each pipeline across a range of between condition effects. We selected experimental effect related neural activities that have been well validated by previous research, showing differences between two task-related or resting related conditions. We averaged the explained variance across time periods where the different conditions have been suggested to show the strongest or most robust differences in neural activity by previous literature. The specific conditions used were selected because of their well-established effects on brain activity and the fact that the conditions selected are commonly of interest in EEG research. As such, effective cleaning should produce larger effect sizes, with higher levels of explained variance produced by more effective cleaning pipelines. We have provided statistical comparisons of the ability of the different pipelines to differentiate experimental conditions by examining the interactions between different pairs of pipelines and the condition of interest (for example between a working memory delay period and working memory probe period in the Sternberg task). We provide heat maps depicting the variance explained by this interaction for each pair of pipelines, marking the interactions that were significant. Significance values were corrected for multiple comparisons across all interaction comparisons within each metric using the Benjamini-Hochberg [36] false discovery rate (* indicates FDR-p < 0.05). We also provide indication of which pipeline provided larger values for variance explained using – and + symbols, which can be interpreted as the pipeline listed on the left of the heatmap having shown less (-) or more (+) variance explained in the comparison between the two experimental conditions than the pipeline listed at the bottom of the heatmap.

Firstly, we examined the variance explained by the difference in upper alpha power (10 to 12.5Hz), computed by Morlet wavelet analyses as per [32] between the data time locked from 500ms to 2500ms after the onset of the working memory delay period in the Sternberg task (which is known to generate very large parietal occipital alpha) and time locked from 0 to 2000ms after the onset of the working memory probe period (which does not produce significant alpha activity). No baseline correction was implemented, because the two periods being compared were from the same epoch, so would be affected equally by any ongoing alpha activity unrelated to task demands. For each pipeline, we calculated the explained variance for the RMS difference between these conditions averaged across the 250 to 1500ms period following stimulus onset, and for the TANOVA from 0 to 750ms and 750 to 2000ms, as all pipelines showed significant differences during these windows, with separate peaks in the TANOVA explained variance corresponding to these windows. Secondly, we examined the amount of variance explained within the averaged RMS and TANOVA by the difference between eyes closed resting (which produces high levels of alpha activity, particularly at parieto-occipital regions) and eyes open resting (which produces less alpha activity). Because resting alpha oscillations are not time locked to external stimuli, we used fast Fourier transforms to calculate these values (and as such, the values are not comparable to the Sternberg alpha data which was computed using Morlet wavelet analyses).

#### SECTION FOUR

##### Results - Combined Sternberg, EO and EC resting data

In this section, we provide figures depicting the results our comparisons, including post-hoc tests and raincloud plots in alternative formats (with either all outliers depicted, outliers removed for easier visualization of the data, or all pipelines present / some pipelines removed for alternative visualization of the data to that provided in the main text). We also provide mean / SD tables for data inspection. We have provided a rank order (by mean) of the best performing pipelines to worst performing pipelines, interpreted from the post-hoc tests which can be visualised in heatmap figures. Significant differences are highlighted for pipelines that performed significantly better than other pipelines using the following notation for ease of understanding: better performance > worse performance (***it is important to note that we have used this better performance > worse performance approach rather than a higher values > lower values approach, as we hope that the consistency will help the reader understand each of the results in the context of all other results***). Because sometimes pipeline 1 differed from pipeline 2, but pipeline 3 did not differ from either 1 or 2, we have used the following notation: ^ = significantly higher than the pipeline marked with a ^^ within the same section (while the others in the category are not significantly different from each other). * = significantly higher than the pipeline marked with a ** in the same category, and so on for the following symbols: [+@$!+]. For each post-hoc figure, values reflect the 95% confidence intervals for the comparison between each pipeline listed on the left, and each pipeline listed along the bottom. Asterix's indicate significant results after multiple comparison controls were applied using the robust post-hoc t-test function "rmmcp", which uses Hochberg's approach to control for the FWE ($p < 0.05$). Note that because the post-hoc t-test significant values were derived from the robust statistics, and the 95% confidence intervals were calculated in the usual parametric manner, sometimes the confidence intervals overlapped with 0 while at the same time the comparison was marked as significant. We interpreted significant differences from the p-value rather than the confidence intervals, but both can be visualised in the figures if the reader would prefer to interpret significance from the confidence intervals.

###### *Signal-to-Error Ratio*

Seven files were excluded specifically from the SER and ARR metrics due to the total artifact periods detected being insufficient for valid calculation of these metrics. There was a significant difference in SER values between the pipelines, with the robust ANOVA showing a significant effect: $F(2.49, 306.76) = 303.851$, $p < 0.0001$. The rank order of significant differences between individual cleaning pipelines from post-hoc t-tests was as follows: MWF_only > MWF_CCA > wICA_ICLabel > MWF_wICA_AMICA > MWF_wICA_fastICA*, MWF_wICA_infomax, ICA_subtract, MWF_wICA_CCA** > ASR, MWF_ICA_subtract > HAPPE > wICA_all. See Figure S1 for a raincloud plot depicting the distribution of the data.

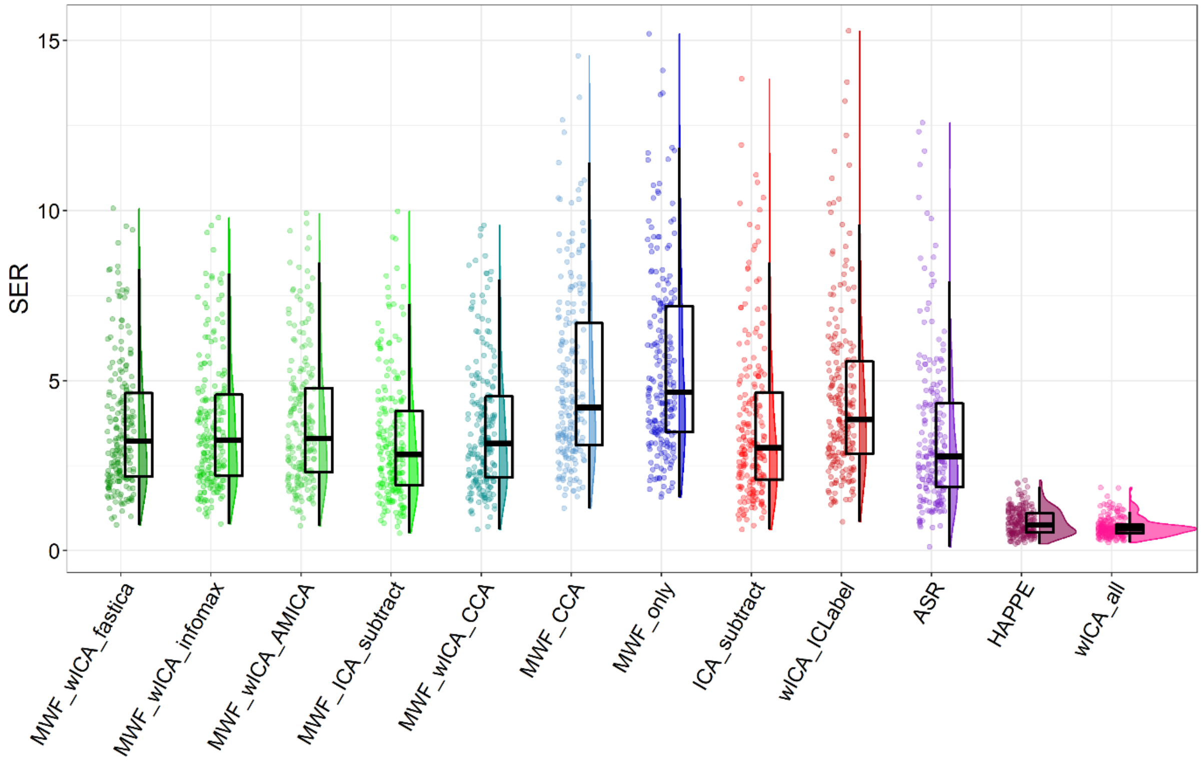

Figure S1. Raincloud plot depicting SER values from the combined EO, EC, and Sternberg data (N = 203) for each of the cleaning pipelines.

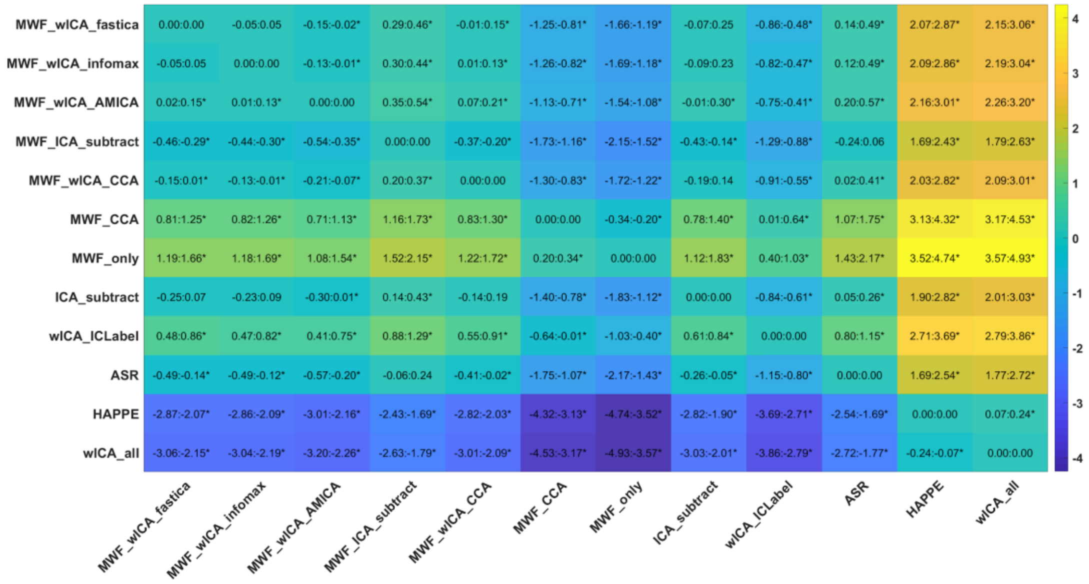

Figure S2. SER post-hoc test results for the combined Sternberg, EO, and EC resting data.

##### *Artifact-to-Residue Ratio*

There was a significant difference in ARR between the pipelines, with the robust ANOVA showing a significant effect: $F(2.65, 326.36) = 1474.71$, $p < 0.0001$. The rank order of significant differences between individual cleaning pipelines from post-hoc t-tests was as follows: HAPPE > wICA_all > MWF_ICA_subtract > MWF_wICA_fastICA,

MWF_wICA_infomax, MWF_wICA_CCA > MWF_wICA_AMICA > ASR > MWF_CCA > wICA_ICLabel > MWF_only, ICA_subtract (Figure S3).

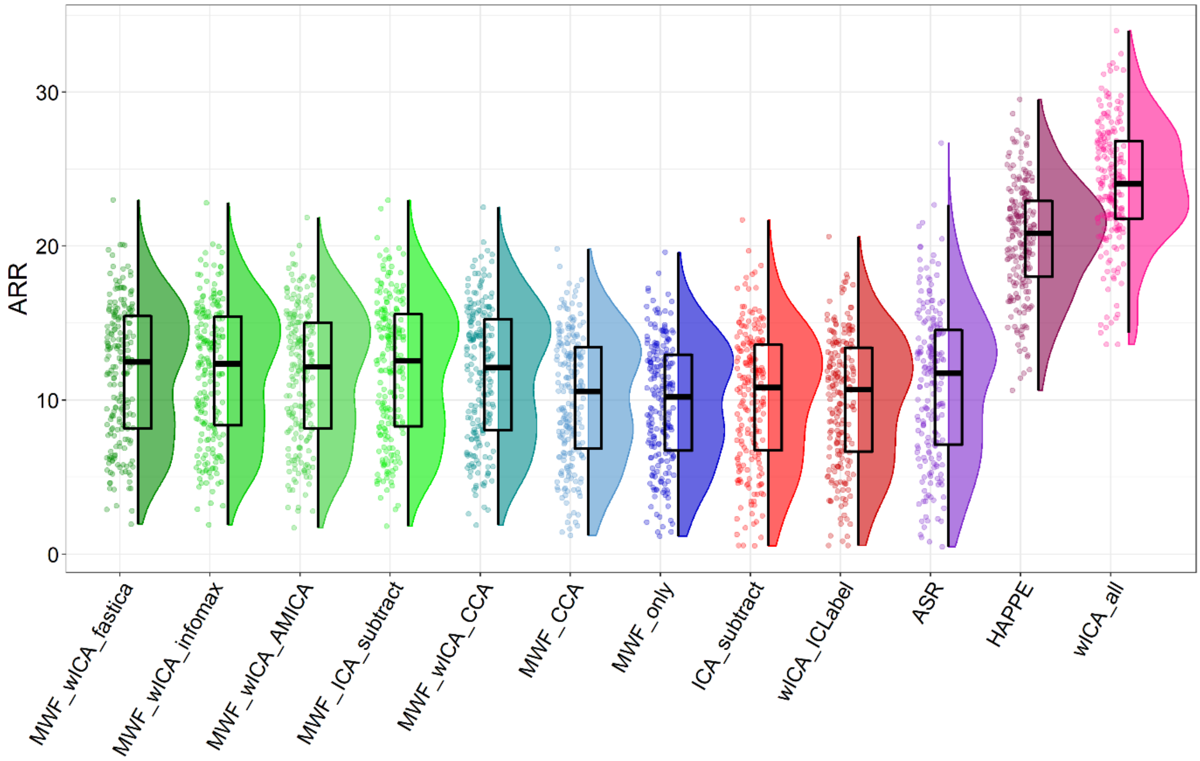

Figure S3. Raincloud plot depicting ARR values from the combined EO, EC, and Sternberg data (N = 203) for each of the cleaning pipelines.

When SER and ARR values were viewed together (Figure S4), it became apparent that the MWF_wICA methods and MWF_wICA_CCA performed better than ASR in both SER and ARR. The MWF_wICA methods and MWF_wICA_CCA also performed equally to, or higher than ICA_subtract in the SER metric, while at the same time they performed better than ICA_subtract in the ARR metric. MWF_wICA_AMICA was slightly better at preserving signal but was slightly worse at removing artifact than the other MWF_wICA methods. MWF_ICA_subtract provided higher ARR values but at the expense of lower SER values than the MWF_wICA methods, although providing similar SER and higher ARR compared to ASR. HAPPE and wICA_all performed the best at removing artifacts (with very high ARR values), but this came at the expense of very low SER values. Inversely, MWF_only, MWF_CCA, and wICA_ICLabel showed high SER values but lower ARR values (with MWF_only performing the best of these three pipelines at providing high SER while producing a similar value for ARR). Note that wICA_ICLabel outperformed ICA_subtract with both higher SER values and higher ARR values.

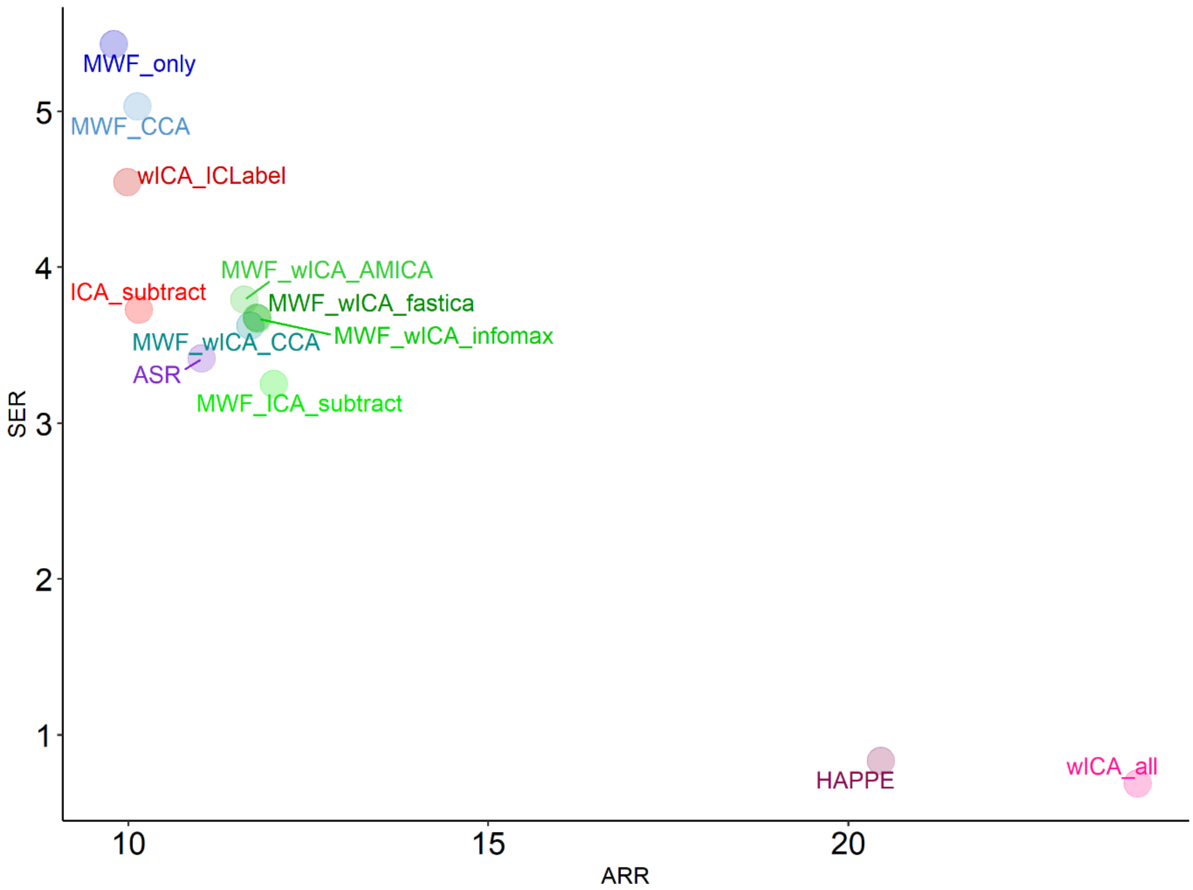

Figure S4. A scatterplot depicting both SER and ARR values for the resting EO, EC, and Sternberg dataset from each cleaning pipeline.

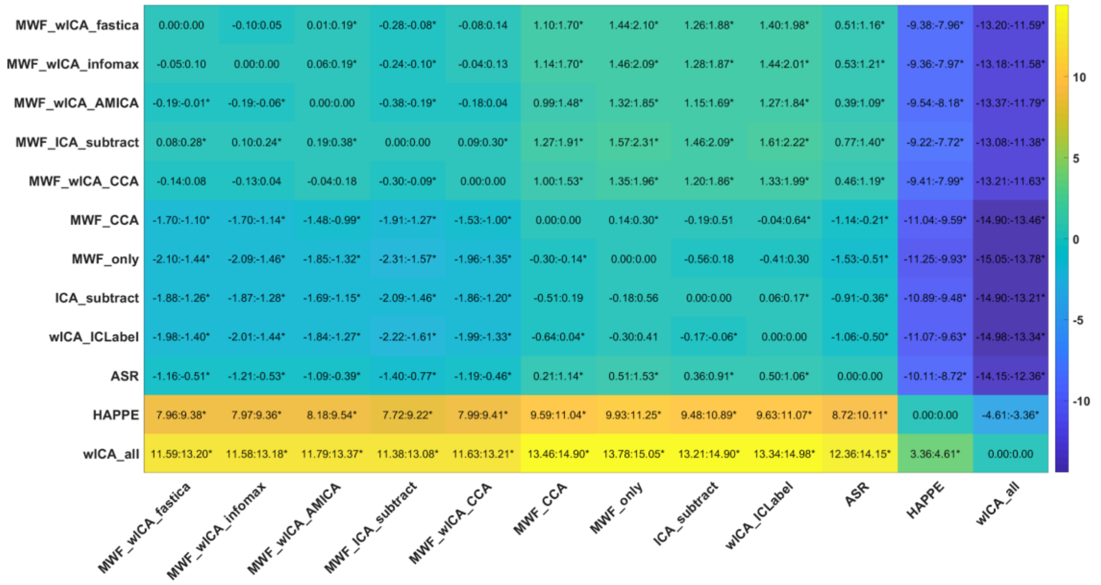

Figure S5. ARR post-hoc test results for the combined Sternberg, EO, and EC resting data.

| Pipeline | ARR Mean | ARR SD | SER Mean | SER SD |
|---|---|---|---|---|
| **MWF_wICA_fastICA** | 11.786 | 4.391 | 3.675 | 1.941 |
| **MWF_CCA** | 10.129 | 4.131 | 5.031 | 2.614 |
| **wICA_all** | 24.021 | 3.923 | 0.687 | 0.276 |
| **HAPPE** | 20.452 | 3.623 | 0.833 | 0.383 |
| **MWF_ICA_subtract** | 12.021 | 4.517 | 3.250 | 1.865 |
| **ICA_subtract** | 10.147 | 4.353 | 3.728 | 2.396 |
| **MWF_wICA_CCA** | 11.698 | 4.372 | 3.625 | 1.931 |
| **MWF_wICA_infomax** | 11.803 | 4.348 | 3.670 | 1.907 |
| **wICA_ICLabel** | 9.985 | 4.237 | 4.546 | 2.528 |
| **MWF_wICA_AMICA** | 11.614 | 4.317 | 3.791 | 1.965 |
| **ASR** | 11.019 | 4.962 | 3.414 | 2.314 |
| **MWF_only** | 9.802 | 3.987 | 5.431 | 2.722 |

Table S1. SER and ARR mean and SD table for the combined Sternberg, EO, and EC resting data.

##### *Frontal Electrode Blink Amplitude Ratio*

There was a significant difference in fBAR between the pipelines, with the robust ANOVA showing a significant effect: $F(3.78, 317.65) = 27.37$, $p < 0.0001$. The rank order of significant differences between individual cleaning pipelines from post-hoc t-tests was as follows (from the best performing pipeline to worst performing pipeline): MWF_wICA_fastICA, MWF_wICA_infomax, MWF_wICA_CCA, MWF_ICA_subtract, MWF_wICA_AMICA > MWF_CCA[@+], wICA_all*, ASR^[@], ICA_subtract^[@@], HAPPE^**, MWF_only[++], wICA_ICLabel^^[@@] (Figure S6).

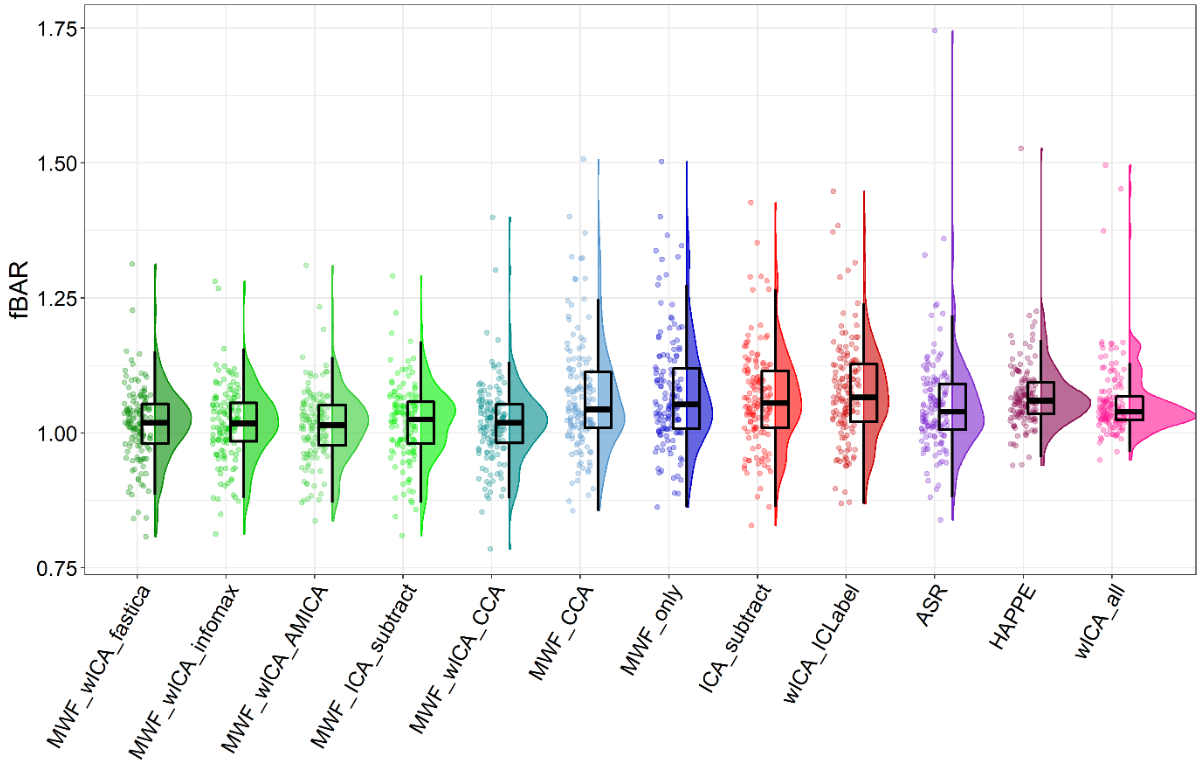

Figure S6. Raincloud plot depicting fBAR values from the combined EO, and Sternberg data (N = 140) for each of the cleaning pipelines.

| | MWF_wICA_fastica | MWF_wICA_infomax | MWF_wICA_AMICA | MWF_ICA_subtract | MWF_wICA_CCA | MWF_CCA | MWF_only | ICA_subtract | wICA_ICLabel | ASR | HAPPE | wICA_all |
|---|---|---|---|---|---|---|---|---|---|---|---|---|
| MWF_wICA_fastica | 0.000:0.000 | -0.006:0.003 | -0.003:0.006 | -0.010:0.003 | -0.005:0.006 | -0.044:-0.009* | -0.055:-0.019* | -0.058:-0.024* | -0.071:-0.037* | -0.043:-0.007* | -0.063:-0.024* | -0.059:-0.009* |
| MWF_wICA_infomax | -0.003:0.006 | 0.000:0.000 | -0.002:0.008 | -0.006:0.003 | -0.003:0.006 | -0.043:-0.009* | -0.055:-0.017* | -0.054:-0.025* | -0.069:-0.036* | -0.042:-0.009* | -0.059:-0.024* | -0.055:-0.008* |
| MWF_wICA_AMICA | -0.006:0.003 | -0.008:0.002 | 0.000:0.000 | -0.010:0.002 | -0.006:0.004 | -0.045:-0.010* | -0.056:-0.019* | -0.060:-0.028* | -0.075:-0.039* | -0.046:-0.009* | -0.065:-0.029* | -0.061:-0.014* |
| MWF_ICA_subtract | -0.003:0.010 | -0.003:0.006 | -0.002:0.010 | 0.000:0.000 | -0.003:0.007 | -0.041:-0.006* | -0.053:-0.016* | -0.053:-0.022* | -0.068:-0.033* | -0.039:-0.006* | -0.061:-0.020* | -0.053:-0.005* |
| MWF_wICA_CCA | -0.006:0.005 | -0.006:0.003 | -0.004:0.006 | -0.007:0.003 | 0.000:0.000 | -0.044:-0.009* | -0.055:-0.017* | -0.055:-0.024* | -0.068:-0.036* | -0.044:-0.008* | -0.064:-0.025* | -0.057:-0.011* |
| MWF_CCA | 0.009:0.044* | 0.009:0.043* | 0.010:0.045* | 0.006:0.041* | 0.009:0.044* | 0.000:0.000 | -0.011:-0.003* | -0.037:0.012 | -0.049:0.001* | -0.022:0.031 | -0.035:0.019 | -0.032:0.032 |
| MWF_only | 0.019:0.055* | 0.017:0.055* | 0.019:0.056* | 0.016:0.053* | 0.017:0.055* | 0.003:0.011* | 0.000:0.000 | -0.027:0.022 | -0.041:0.012 | -0.012:0.043 | -0.027:0.027 | -0.022:0.040 |
| ICA_subtract | 0.024:0.058* | 0.025:0.054* | 0.028:0.060* | 0.022:0.053* | 0.024:0.055* | -0.012:0.037 | -0.022:0.027 | 0.000:0.000 | -0.015:-0.005* | 0.006:0.029* | -0.030:0.017 | -0.023:0.031 |
| wICA_ICLabel | 0.037:0.071* | 0.036:0.069* | 0.039:0.075* | 0.033:0.068* | 0.036:0.068* | -0.001:0.049* | -0.012:0.041 | 0.005:0.015* | 0.000:0.000 | 0.016:0.042* | -0.020:0.029 | -0.012:0.040 |
| ASR | 0.007:0.043* | 0.009:0.042* | 0.009:0.046* | 0.006:0.039* | 0.008:0.044* | -0.031:0.022 | -0.043:0.012 | -0.029:-0.006* | -0.042:-0.016* | 0.000:0.000 | -0.045:0.004 | -0.033:0.016 |
| HAPPE | 0.024:0.063* | 0.024:0.059* | 0.029:0.065* | 0.020:0.061* | 0.025:0.064* | -0.019:0.035 | -0.027:0.027 | -0.017:0.030 | -0.029:0.020 | -0.004:0.045 | 0.000:0.000 | 0.004:0.026* |
| wICA_all | 0.009:0.059* | 0.008:0.055* | 0.014:0.061* | 0.005:0.053* | 0.011:0.057* | -0.032:0.032 | -0.040:0.022 | -0.031:0.023 | -0.040:0.012 | -0.016:0.033 | -0.026:-0.004* | 0.000:0.000 |

Figure S7. Blink amplitude ratio, frontal electrodes post-hoc test for the combined Sternberg, EO, and EC resting data.

##### ***Blink Amplitude Ratio for All Electrodes***

There was a significant difference in blink amplitude ratio in all electrodes between the pipelines, with the robust ANOVA showing a significant effect: $F(3.19, 268.14) = 40.32$, $p < 0.0001$. The rank order of significant differences between individual cleaning pipelines from post-hoc t-tests was as follows (from the best performing pipeline to worst performing pipeline): MWF_wICA_infomax, MWF_wICA_fastICA, MWF_wICA_AMICA, MWF_wICA_CCA > MWF_ICA_subtract^, wICA_all*, MWF_CCA^^, HAPPE**, MWF_only^^**, ASR^^** > ICA_subtract, wICA_ICLabel. See Figure S8 for a raincloud plot depicting the distribution of the data.

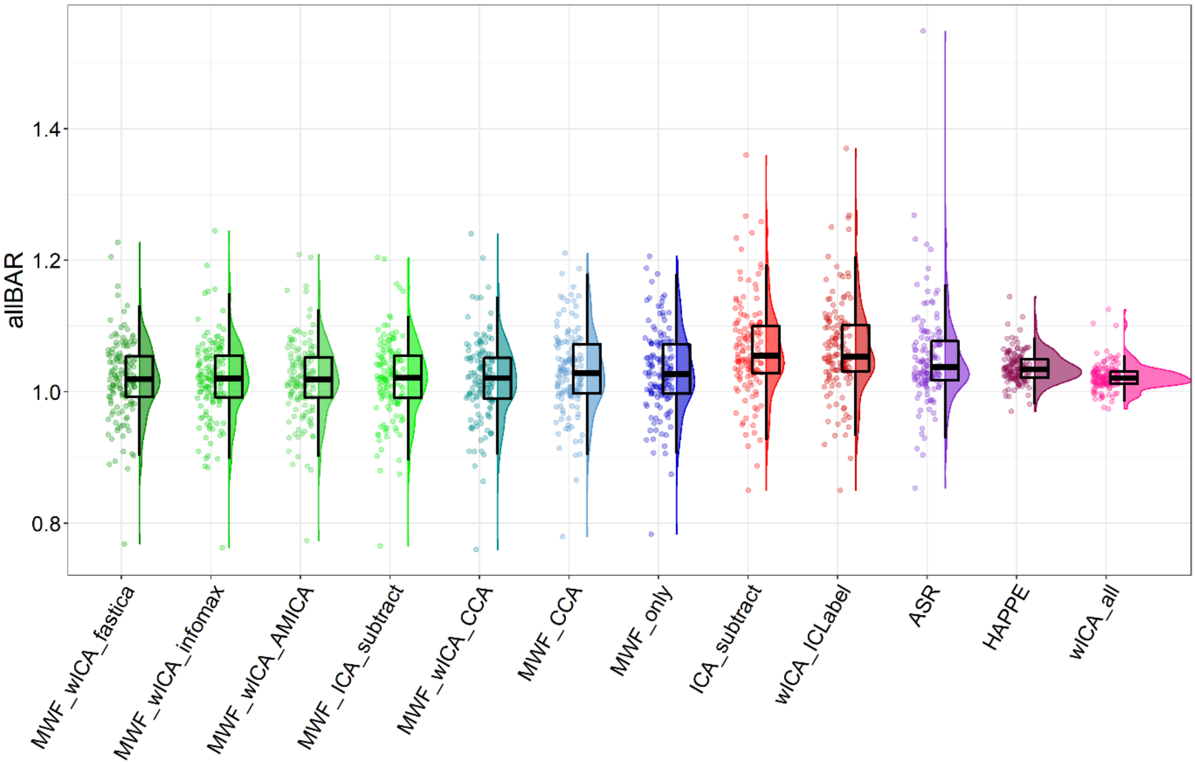

Figure S8. Raincloud plot depicting allBAR values from the combined EO, EC, and Sternberg data (N = 140) for each of the cleaning pipelines.

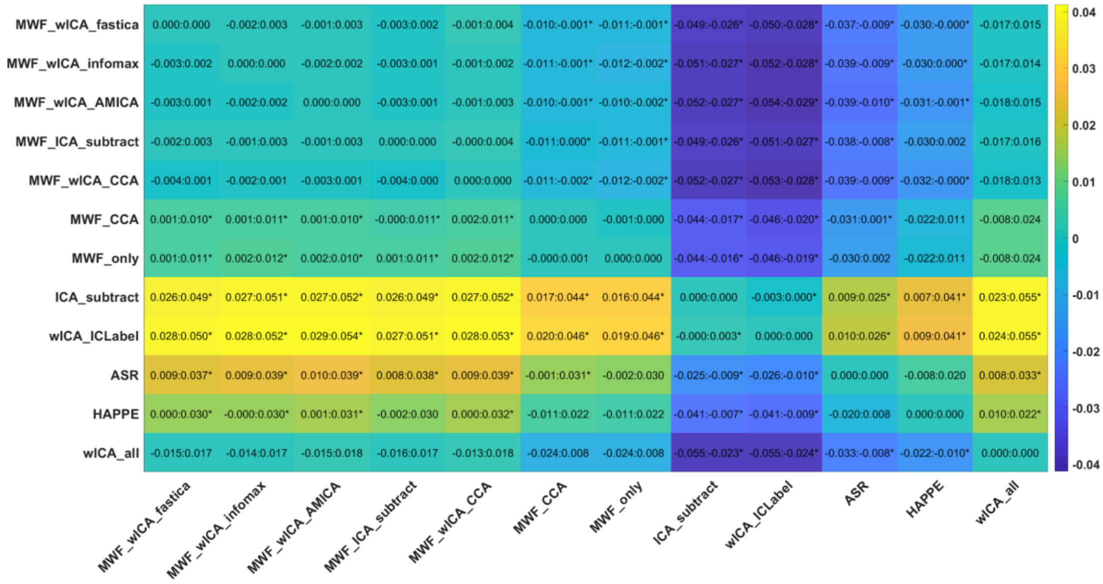

Figure S9. Blink amplitude ratio, all electrodes post-hoc test for the combined Sternberg, EO, and EC resting data.

| | fBAR | | allBAR | |
|---|---|---|---|---|
| **Pipeline** | **Mean** | **SD** | **Mean** | **SD** |
| **MWF_CCA** | 1.068 | 0.107 | 1.031 | 0.064 |
| **ICA_subtract** | 1.064 | 0.092 | 1.064 | 0.073 |
| **HAPPE** | 1.069 | 0.065 | 1.037 | 0.024 |
| **wICA_ICLabel** | 1.076 | 0.095 | 1.066 | 0.073 |
| **MWF_only** | 1.077 | 0.107 | 1.031 | 0.063 |
| **MWF_ICA_subtract** | 1.020 | 0.072 | 1.021 | 0.061 |
| **MWF_wICA_fastICA** | 1.018 | 0.073 | 1.020 | 0.060 |
| **MWF_wICA_AMICA** | 1.015 | 0.069 | 1.019 | 0.060 |
| **MWF_wICA_infomax** | 1.018 | 0.070 | 1.020 | 0.061 |
| **wICA_all** | 1.059 | 0.073 | 1.022 | 0.021 |
| **MWF_wICA_CCA** | 1.018 | 0.076 | 1.019 | 0.062 |
| **ASR** | 1.052 | 0.097 | 1.051 | 0.074 |

Table S2. Blink amplitude ratio mean and SD table for the combined Sternberg, EO, and EC resting data.

***Proportion of Epochs Showing Muscle Activity After Cleaning***

There was a significant difference between the pipelines in number of epochs with log-power log-frequency slopes indicating that muscle activity was remaining after cleaning, with the robust ANOVA showing a significant effect: $F(1.05, 133.91) = 3931.25$, $p < 0.0001$. The rank order of significant differences between individual cleaning pipelines from post-hoc t-tests was as follows (from the best performing pipeline to worst performing pipeline): MWF_ICA_subtract > MWF_wICA_infomax > MWF_wICA_fastICA > MWF_wICA_AMICA > MWF_wICA_CCA > MWF_CCA > ICA_subtract > ASR, wICA_ICLabel > MWF_only > HAPPE > wICA_all (Figure S10). It is worth noting that HAPPE and wICA produced data that showed slopes exceeding the muscle threshold for almost all epochs (and as such, were not depicted in Figure S10 so other pipelines could be compared, but can be viewed in Figure

S11). We suspect this is the result of removal of considerable low frequency data from the signal, such that slopes became flatter (and more similar to muscle affected log-power log-frequency slopes). However, we note that these pipelines did show a bump in the beta frequency after cleaning (which the other pipelines did not show), so it is possible this reflects muscle activity remaining.

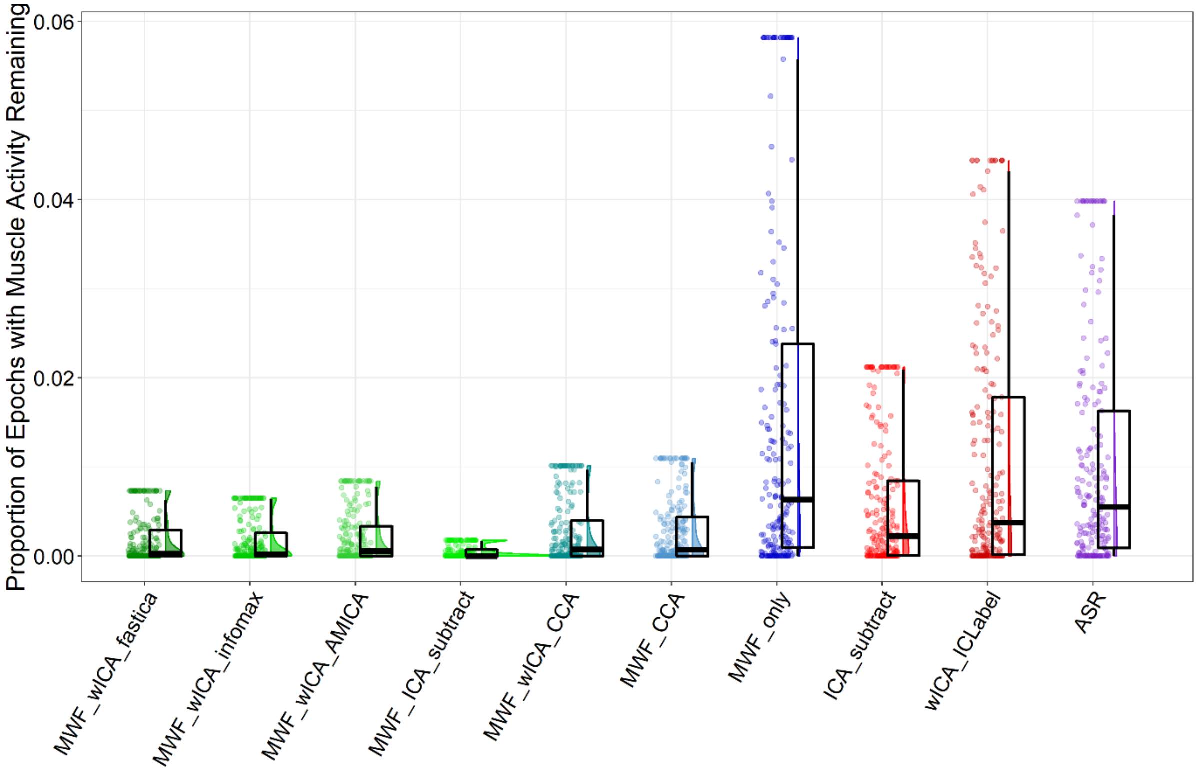

Figure S10. Raincloud plot depicting the proportion of epochs showing log-power log-frequency values above the -0.59 threshold from the combined EO, EC, and Sternberg data (N = 213) for each of the cleaning pipelines. Note that this figure excludes HAPPE and wICA_all, as these pipelines showed median values > 0.75 and made the scale of the graph such that it was difficult to visualise differences in the other pipelines. We suspect that applying wICA to all components (as both wICA_all and HAPPE do) reduces the activity contributed by the lower frequencies to the extent that the log-power log-frequency slope threshold used identifies most epochs as contaminated by muscle activity, when the shallow slope is actually because the low frequencies have been removed from the data. Note also that we have winsorized the data in the figure, as the outliers also made the scale such that it was difficult to visualise differences in the other pipelines. The full data can be viewed in Figure S11.

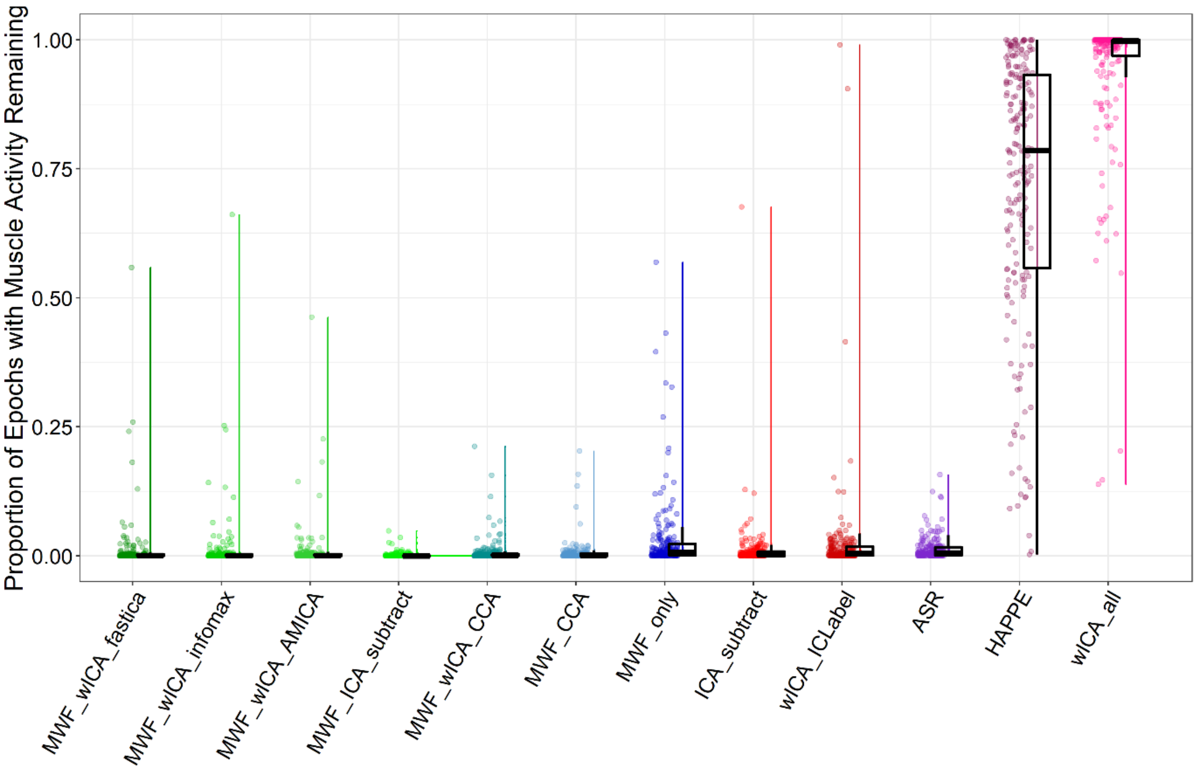

Figure S11. Raincloud plot of the proportion of epochs showing log-power log-frequency slopes that indicated muscle activity remaining after cleaning (including all pipelines and without winsorizing outliers) for the combined Sternberg, EO, and EC resting data.

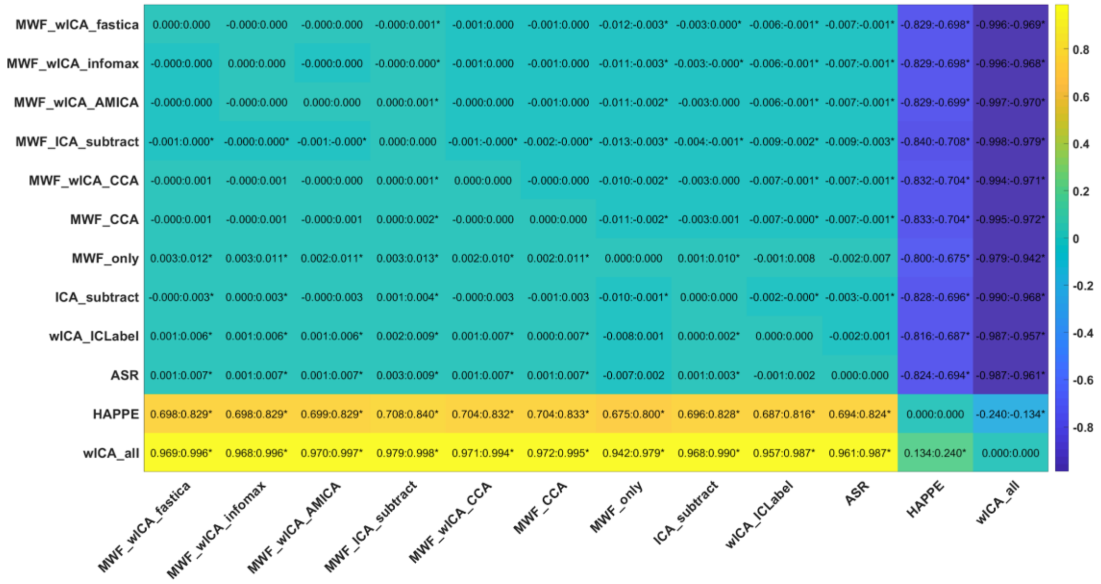

Figure S12. Post-hoc tests for the proportion of epochs showing log-power log-frequency slopes above the threshold after cleaning for the combined Sternberg, EO, and EC resting data.

***Severity by which the Log-Power Log-Frequency Slopes Exceed the Muscle Threshold***

There was a significant difference between the pipelines in the severity by which the mean slope exceeded the log-power log-frequency threshold from epochs and electrodes that showed muscle activity remaining, with the robust ANOVA showing a significant effect: $F(5.6, 716.85) = 170.4889$, $p < 0.0001$. The rank order of significant differences between individual cleaning pipelines from post-hoc t-tests was as follows (from the best performing pipeline to worst performing pipeline): MWF_ICA_subtract^*, MWF_wICA_infomax*, MWF_wICA_fastICA*^^, MWF_wICA_CCA^^, MWF_wICA_AMICA^^**, MWF_CCA^^** > wICA_ICLabel^, MWF_only, ICA_subtract, ASR^^ > HAPPE > wICA_all. See Figure S13 for a raincloud plot depicting the distribution of the data.

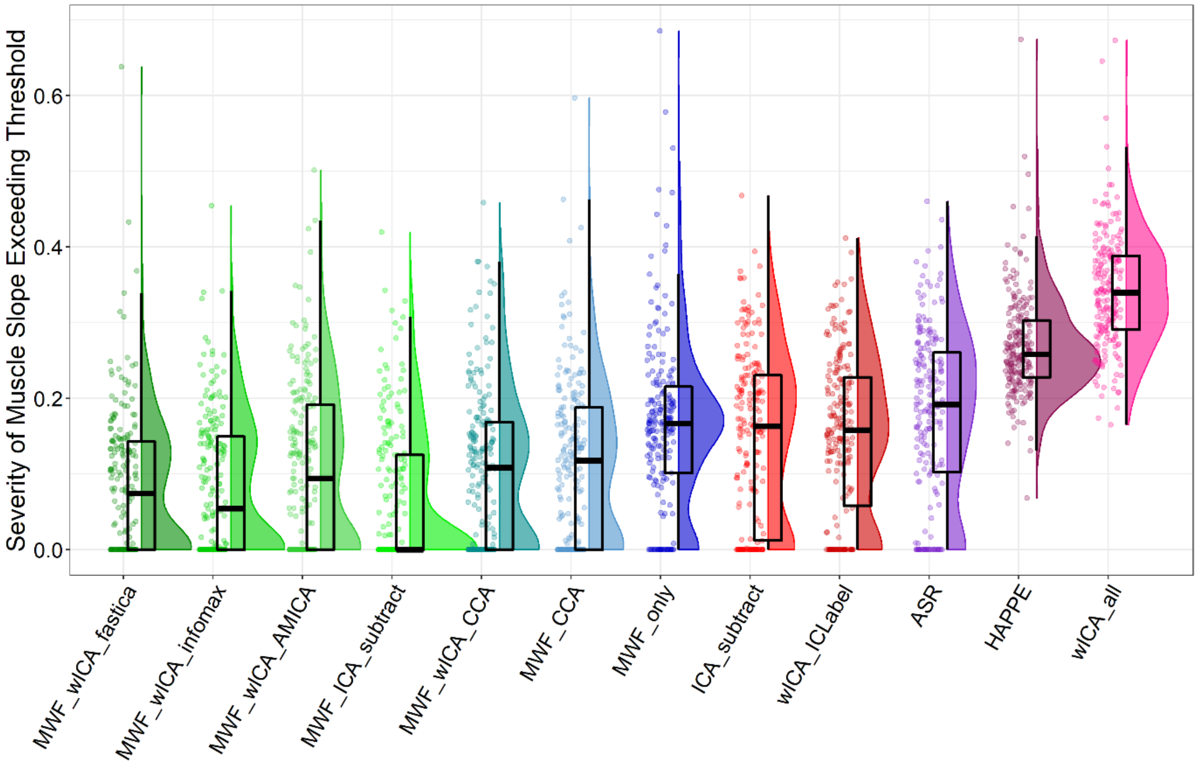

Figure S13. Raincloud plot depicting the severity by which the log-power log-frequency slopes exceeded the -0.59 threshold, when values were averaged across super-threshold epochs and electrodes from the combined EO, EC, and Sternberg data (N = 213) for each of the cleaning pipelines.

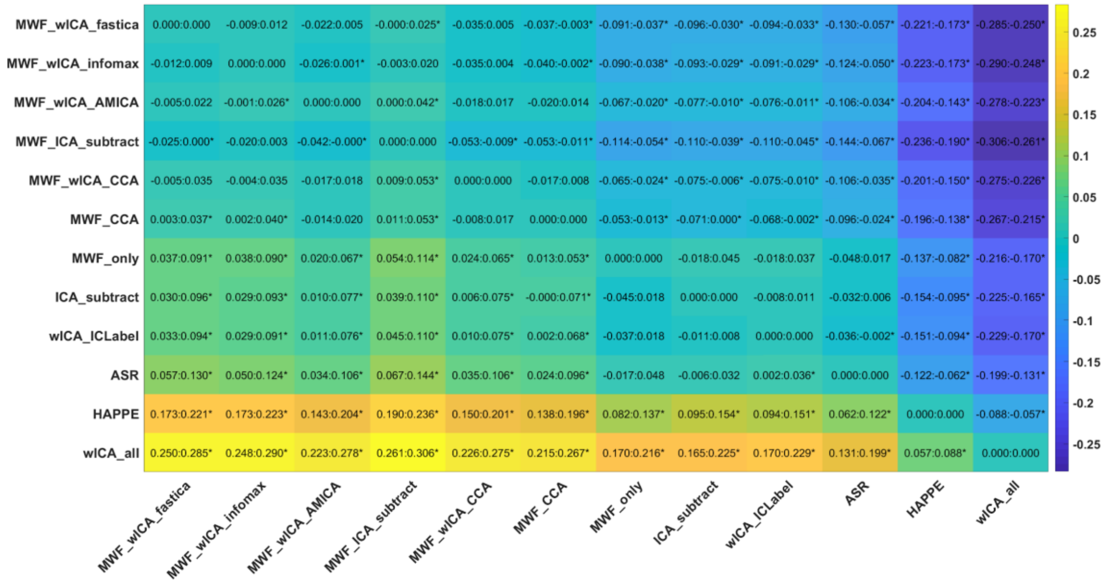

Figure S14. Post-hoc tests for the severity of the log-power log-frequency slopes exceeding the threshold in epochs that showed muscle activity remaining for the combined Sternberg, EO, and EC resting data.

| | Proportion of epochs showing muscle slope after cleaning | | Slope steepness over muscle slope threshold in epochs showing muscle slopes after cleaning | |
|---|---|---|---|---|
| **Pipeline** | **Mean** | **SD** | **Mean** | **SD** |
| **ASR** | 0.01 | 0.013 | 0.179 | 0.116 |
| **HAPPE** | 0.715 | 0.266 | 0.269 | 0.061 |
| **ICA_subtract** | 0.006 | 0.007 | 0.15 | 0.117 |
| **MWF_CCA** | 0.003 | 0.004 | 0.115 | 0.107 |
| **MWF_wICA_CCA** | 0.003 | 0.004 | 0.107 | 0.104 |
| **MWF_wICA_AMICA** | 0.002 | 0.003 | 0.11 | 0.112 |
| **MWF_wICA_fastICA** | 0.002 | 0.003 | 0.083 | 0.089 |
| **MWF_ICA_subtract** | <0.001 | 0.001 | 0.069 | 0.093 |
| **MWF_wICA_infomax** | 0.002 | 0.002 | 0.084 | 0.093 |
| **MWF_only** | 0.016 | 0.02 | 0.163 | 0.104 |
| **wICA_all** | 0.976 | 0.035 | 0.341 | 0.074 |
| **wICA_ICLabel** | 0.011 | 0.014 | 0.151 | 0.108 |

Table S3. Means and SDs for the muscle related metrics for the combined Sternberg, EO, and EC resting data.

##### *ICA Variance Explained by Brain Components*

There was a significant difference between the pipelines in the percentage of ICA variance explained by components identified as brain activity with the robust ANOVA showing a significant effect: $F(1.49, 189.39) = 1238.15$, $p < 0.0001$. The rank order of significant differences between individual cleaning pipelines from post-hoc t-tests was as follows: MWF_wICA_infomax, wICA_ICLabel > MWF_wICA_fastICA, MWF_wICA_AMICA, MWF_only > wICA_all (note that pipelines using ICA subtraction were excluded from this metric) (Figure S15).

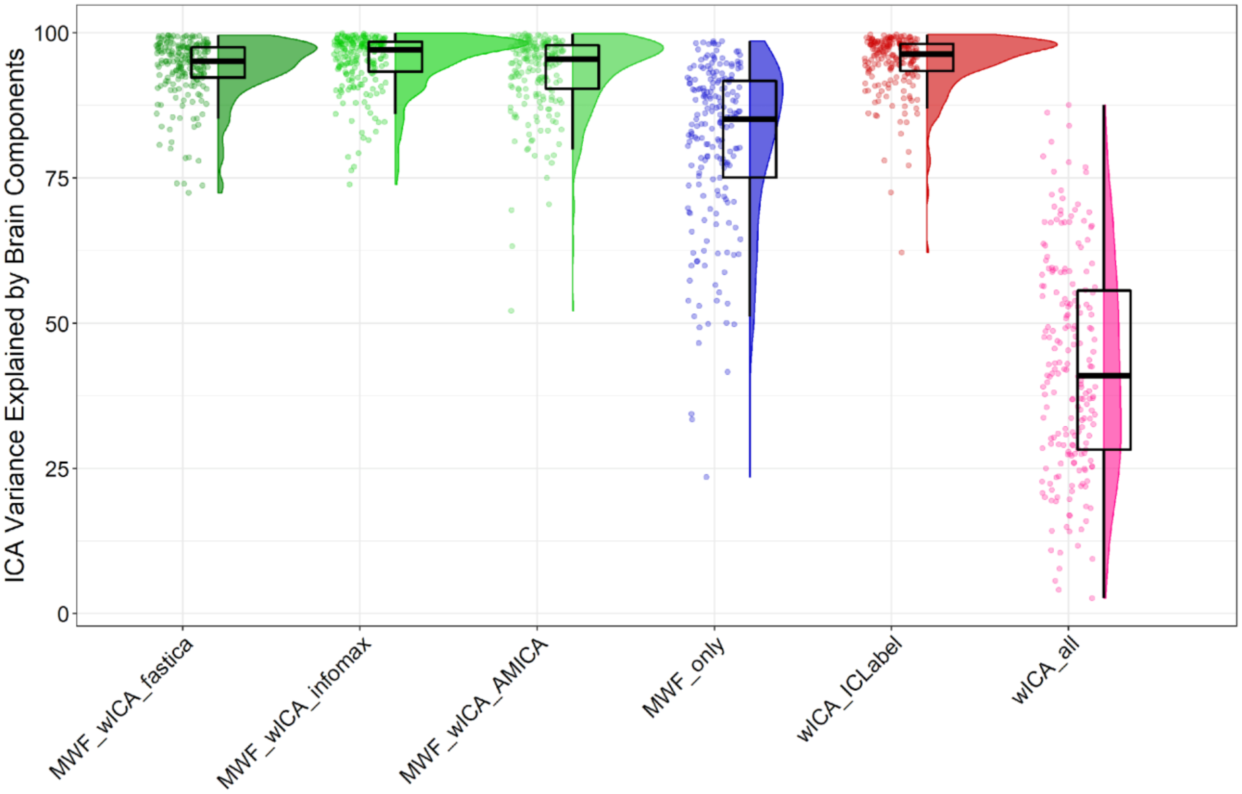

Figure S15. Raincloud plot depicting the amount of ICA variance explained by brain activity from the combined EO, EC, and Sternberg data (N = 213) for each of the cleaning pipelines available for assessment with this metric (note that pipelines using ICA subtraction were excluded from this metric).

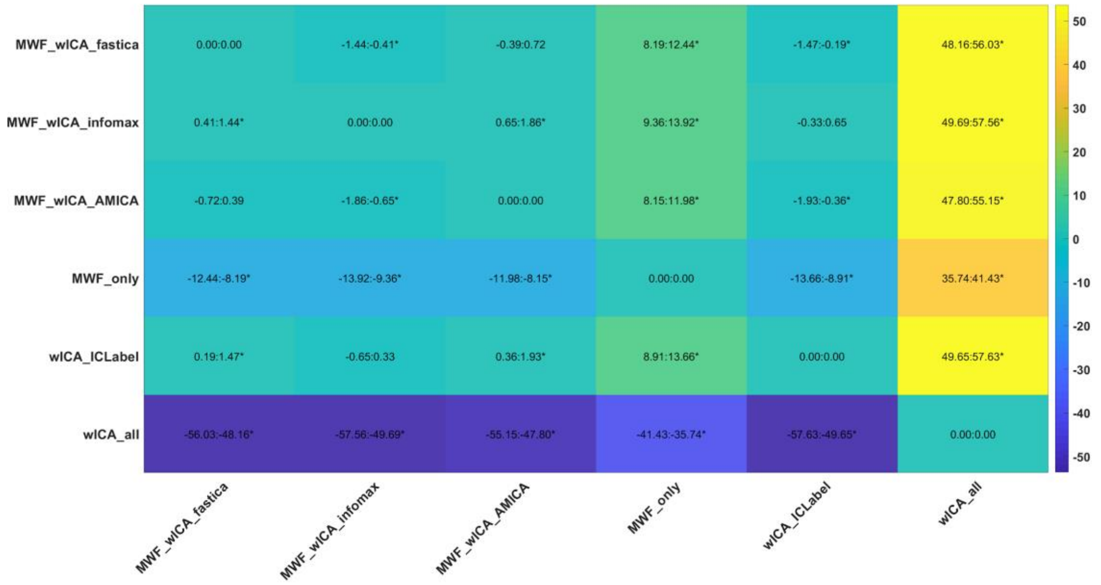

Figure S16. Variance explained by brain activity detected by ICLabel post-hoc tests for the combined Sternberg, EO, and EC resting data.

| Pipeline | Mean | SD |
|---|---:|---:|
| **MWF_only** | 81.166 | 14.150 |
| **MWF_wICA_infomax** | 95.090 | 4.896 |
| **wICA_ICLabel** | 94.852 | 5.030 |
| **MWF_wICA_AMICA** | 93.111 | 6.896 |
| **MWF_wICA_fastICA** | 93.825 | 5.316 |
| **wICA_all** | 42.401 | 18.283 |

Table S4. Means and SDs for the amount of variance explained by brain activity detected by ICLabel for the combined Sternberg, EO, and EC resting data.

***Proportion of Epochs Removed by Cleaning***

There was a significant difference between the pipelines in the proportion of total epochs in the data that were rejected by the cleaning and outlying epoch rejection steps, with the robust ANOVA showing a significant effect: $F(2.04, 258.98) = 73.9214$, $p < 0.0001$. The rank order of significant differences between individual cleaning pipelines from post-hoc t-tests was as follows (from the best performing pipeline to worst performing pipeline): HAPPE > wICA_all > MWF_wICA_fastICA^, MWF_wICA_infomax*, MWF_ICA_subtract*+^^, MWF_wICA_AMICA*, MWF_wICA_CCA^^**@, MWF_CCA^^**++, MWF_only^^**++@@, ICA_subtract^^**++, wICA_ICLabel^^**++. ASR showed the highest mean value, but only significantly differed from HAPPE. ASR also showed a very large spread of datapoints, with broad confidence intervals when compared to all other pipelines, and the rejection of 75-100% of the epochs for seven data files (whereas all other pipelines showed at most three data files with more than 75% of epochs rejected). We wondered if this suggested that the robust statistics used (rmmcp) were obscuring the pattern, so we performed post-hoc t-tests with bootstrap statistics (using pairdepb) and found that the ASR pipeline showed a higher mean proportion of the data rejected than all other pipelines (all p-bootstrap < 0.05) (Figure S17).

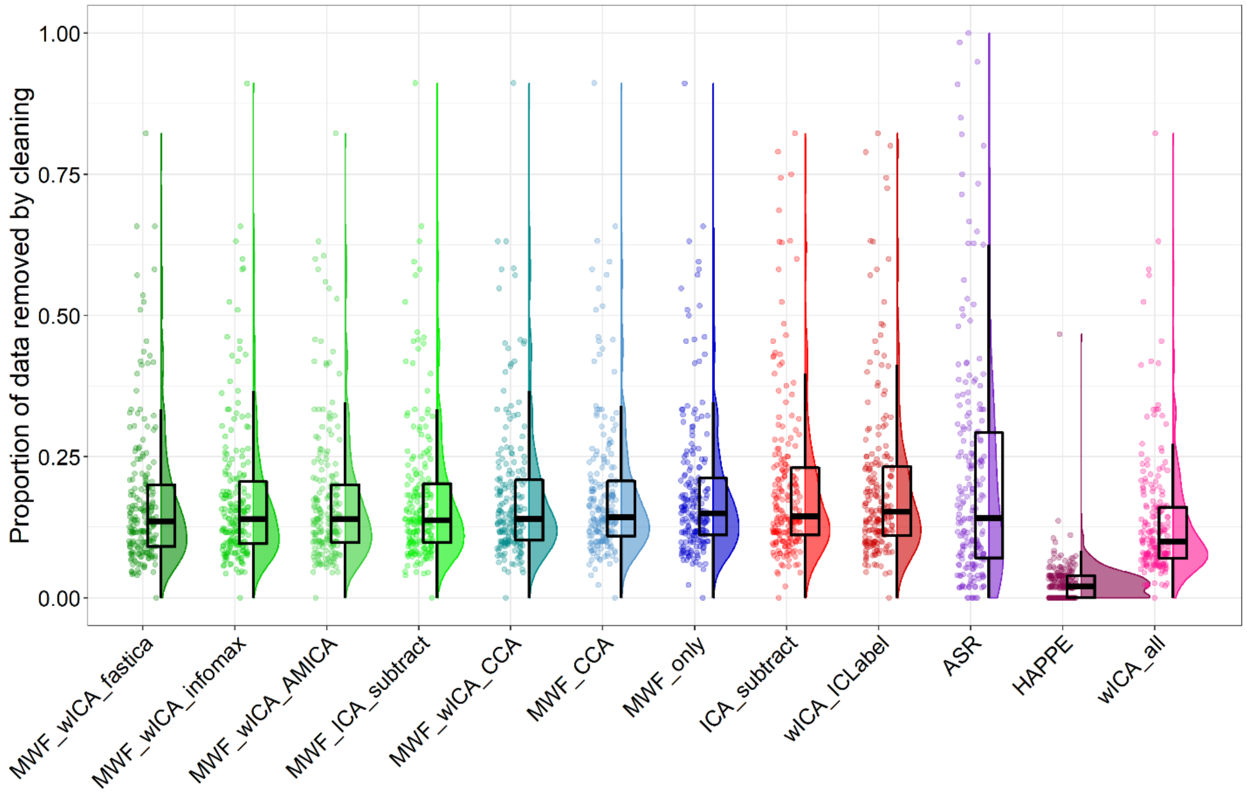

Figure S17. Raincloud plot depicting the proportion of epochs in the data rejected from the combined EO, EC, and Sternberg data (N = 213) for each of the cleaning pipelines.

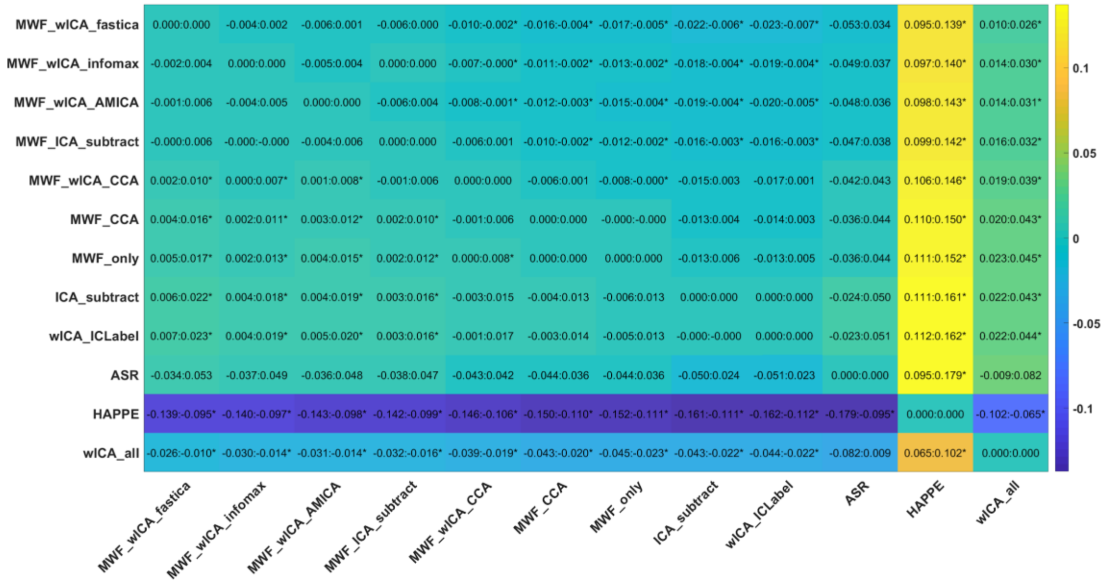

Figure S18. Proportion of epochs in the data removed by the cleaning pipeline, including epoch rejection of outliers after cleaning post-hoc test for the combined Sternberg, EO, and EC resting data.

| | Proportion of Epochs Removed | |
|---|---|---|
| **Pipeline** | **Mean** | **SD** |
| **MWF_ICA_subtract** | 0.177 | 0.128 |
| **wICA_all** | 0.136 | 0.114 |
| **MWF_wICA_infomax** | 0.175 | 0.128 |
| **MWF_only** | 0.187 | 0.128 |
| **wICA_ICLabel** | 0.200 | 0.148 |
| **MWF_wICA_CCA** | 0.181 | 0.125 |
| **ICA_subtract** | 0.198 | 0.147 |
| **ASR** | 0.212 | 0.207 |
| **MWF_wICA_AMICA** | 0.175 | 0.125 |
| **MWF_CCA** | 0.185 | 0.127 |
| **MWF_wICA_fastICA** | 0.172 | 0.126 |
| **HAPPE** | 0.028 | 0.039 |

Table S5. Means and SDs for the proportion of epochs of the data removed by the cleaning pipeline (including epoch rejection of outliers after cleaning) for the combined Sternberg, EO, and EC resting data.

***Variance Explained by the Difference Between Eyes Open and Eyes Closed Resting***

Figure S19 depicts the amount of variance explained by the difference between eyes open and eyes closed resting in alpha power RMS (overall neural response strength) and TANOVA (distribution of neural activity) tests across the different cleaning pipelines. Statistical comparisons using the RMS and TANOVA tests of the overall interaction between pipelines and condition were highly significant for both measures (both $p < 0.0001$). Post-hoc testing of the interaction between each pair of pipelines and the two conditions indicated the following rank order of the ability of the pipelines to discriminate between the experimental manipulation in alpha RMS: ASR*, MWF_wICA_fastICA*$^{@}$, MWF_wICA_infomax*$^{@}$, MWF_wICA_AMICA*$^{@}$, MWF_ICA_subtract*$^{+@@}$, MWF_wICA_CCA*, MWF_only*$^{++}$, MWF_CCA*$^{++}$, HAPPE**^, wICA_ICLabel^^, ICA_subtract^^ > wICA_all. We suspect the fact that only HAPPE and wICA_all showed significant differences to the other pipelines is a product of the low variance in alpha power across individuals within each condition in the HAPPE and wICA_all pipelines, combined with the relative high variance in alpha power across individuals within each condition in all other pipelines (see Figure S20).

Post-hoc testing of the interaction between each pair of pipelines and the two conditions indicated the following rank order of the ability of the pipelines to discriminate between the experimental manipulation in the distribution of alpha power: wICA_all > MWF_CCA > MWF_only > MWF_wICA_infomax*, MWF_wICA_AMICA*^, MWF_wICA_fastICA*$^{+}$, MWF_wICA_CCA*$^{+}$, MWF_ICA_subtract*^^, wICA_ICLabel**^^, ASR, ICA_subtract**. HAPPE provided the lowest variance explained, but significantly differed from all pipelines *except* MWF_CCA and MWF_only (which showed the 2$^{nd}$ and 3$^{rd}$ largest np$^2$ values). This odd result may be due to the multidimensional nature of interactions between pipeline and EO/EC conditions in the TANOVA, combined with the fact the TANOVA compares the distribution of activity (rather than a single value). As such, the result may be due to a more similar match in topographical difference between EO and EC conditions for HAPPE, MWF_CCA, and MWF_only compared to the other pipelines. The results also provide further suggestion of high variability in the data from the ASR pipeline, which despite showing the third lowest variance explained, did not significantly differ in the interaction between cleaning pipeline and condition for the majority of the other pipelines.

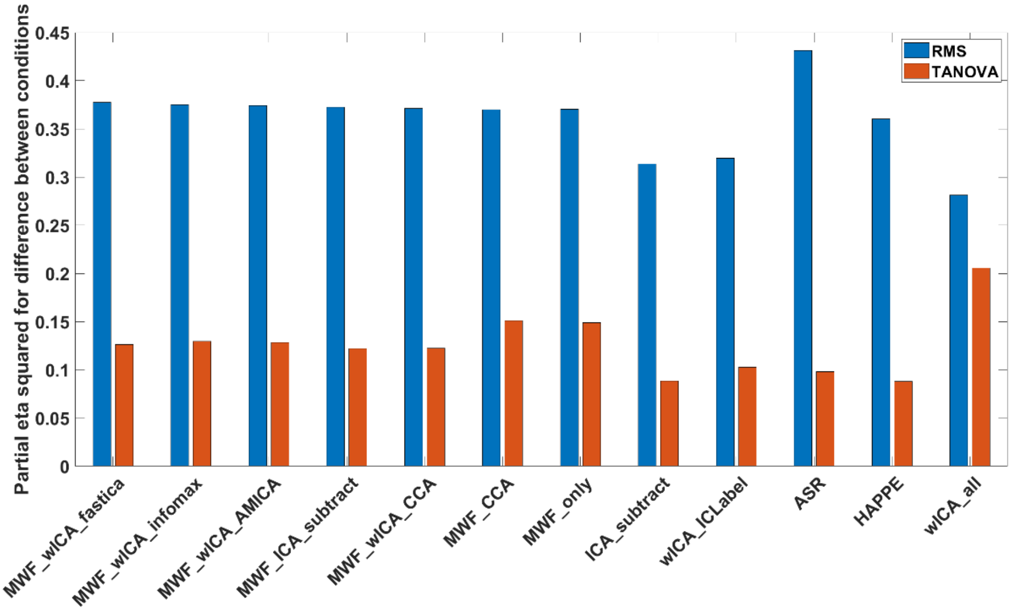

Figure S19. The variance explained (np$^2$) by differences in averaged alpha power between eyes open and eyes closed for RMS and TANOVA tests for all of the pipelines.

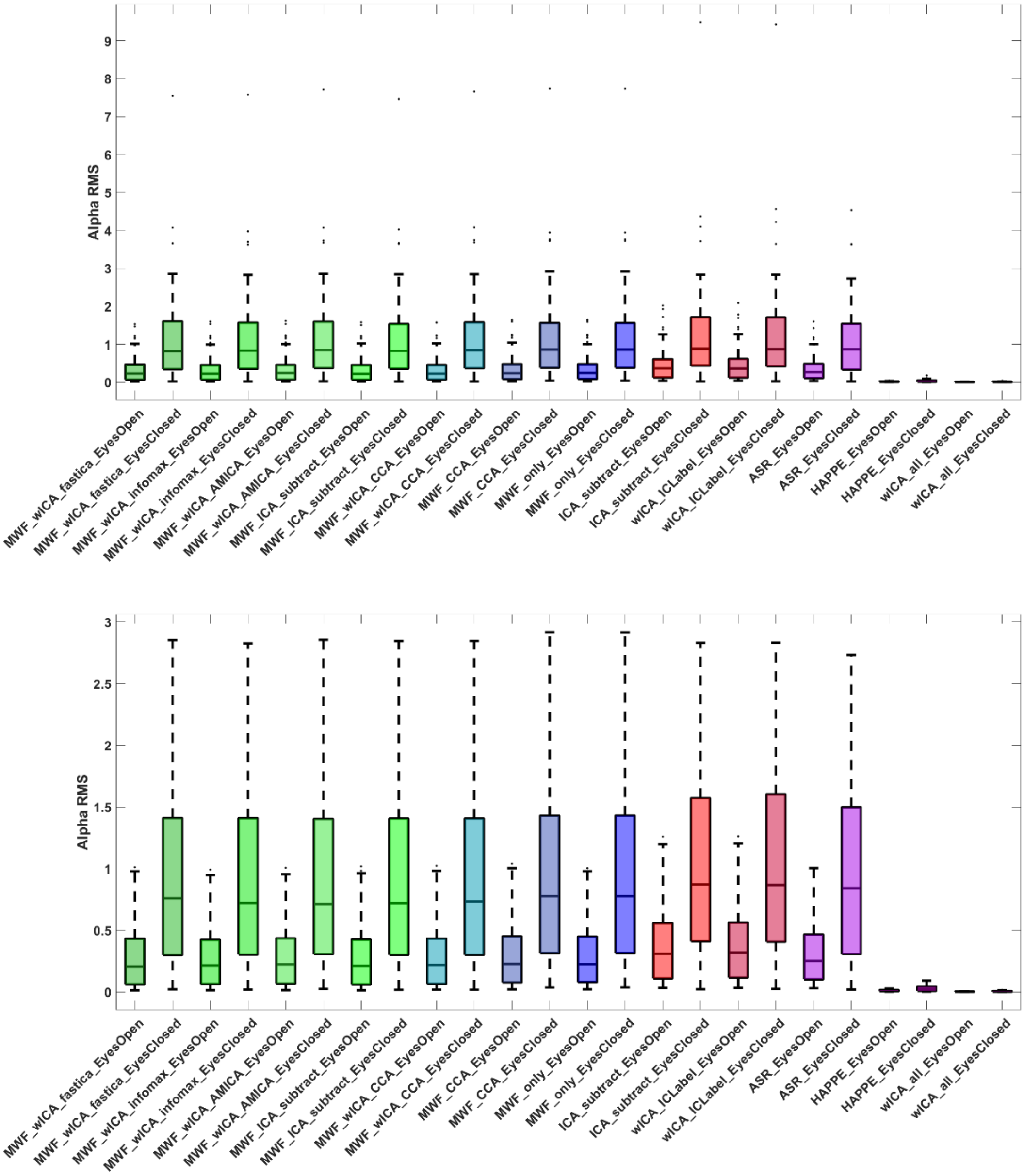

Figure S20. Box plots for the alpha RMS values from eyes open and eyes closed resting recordings after cleaning by each pipeline. Above, with outliers included, and below, with outliers removed for better visualization of the difference between the cleaning pipelines.

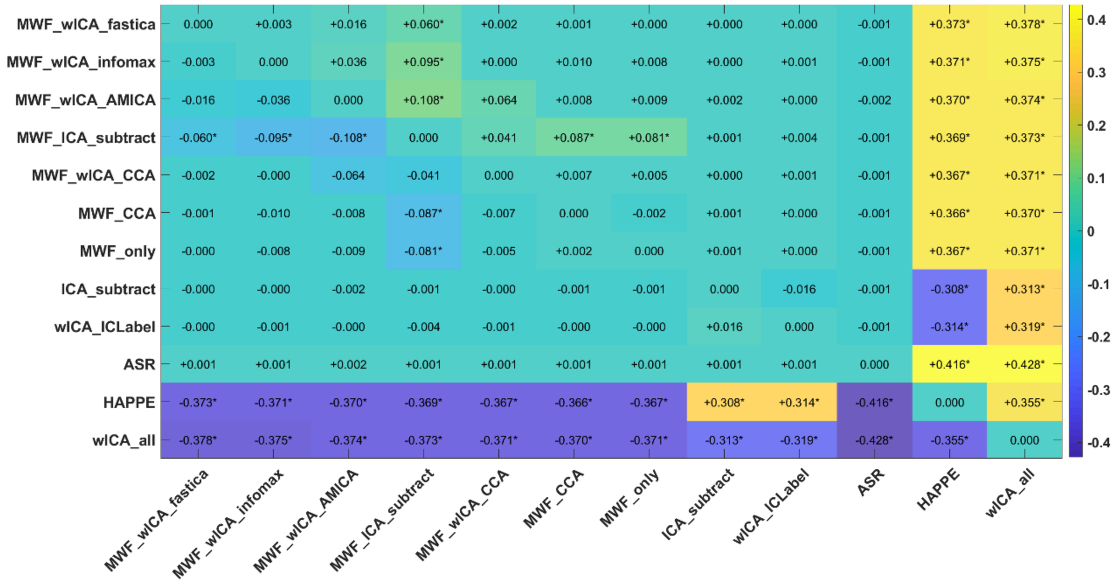

Figure S21. Heat map of the variance explained ($np^2$) by the interaction between each pair of pipelines and alpha power RMS during eyes open and eyes closed resting. Interactions that were significant (FDR-$p < 0.05$) are indicated with an *. We have also provided an indication of which pipeline of each pair provided larger values for variance explained using – and + symbols, which can be interpreted as the pipeline listed on the left of the heatmap having shown less (-) or more (+) variance explained in the comparison between the two experimental conditions than the pipeline listed at the bottom of the heatmap.

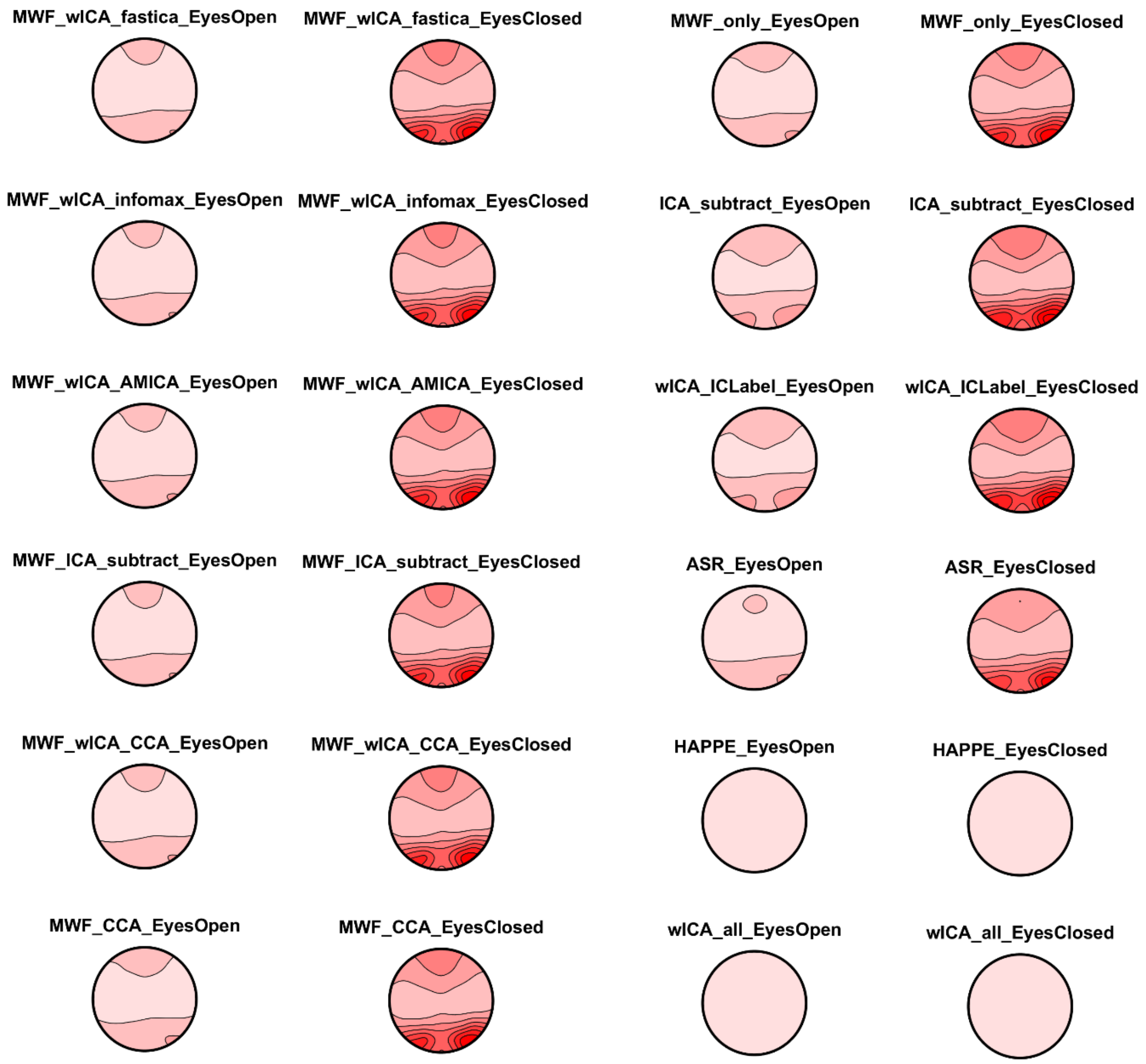

Figure S22. Resting alpha activity distributions from eyes open and eyes closed recordings after cleaning by each pipeline. All plots are on the same scale so they can all be compared to all other pipelines. Note that wICA_all and HAPPE have removed almost all alpha activity, and that ASR has removed more of the activity in the eyes open resting condition than the other pipelines.

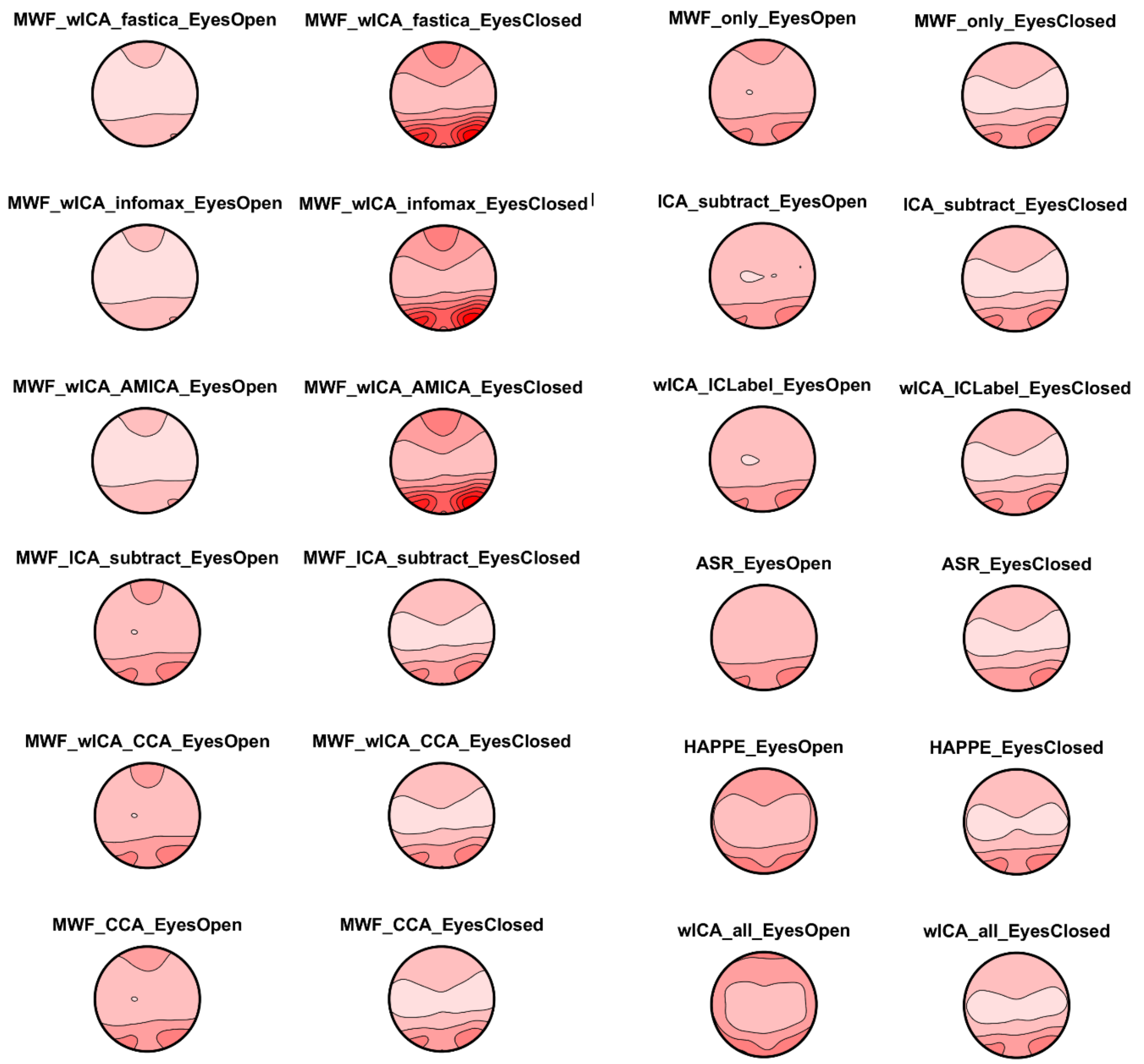

Figure S23. Resting alpha activity distributions from eyes open and eyes closed recordings after cleaning by each pipeline. All plots are on their own individual scale so the pattern of alpha activity distributions can be viewed within each plot, but comparisons between pipelines / conditions are not possible from these plots.

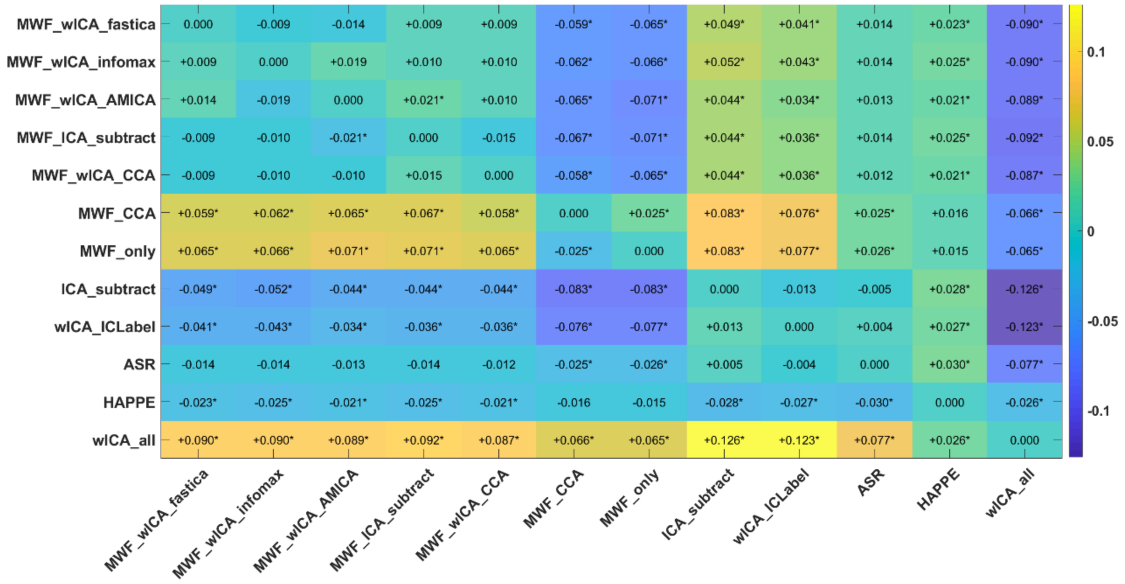

Figure S24. Heat map of the variance explained (np$^2$) by the interaction between each pair of pipelines and the distribution of alpha power (TANOVA) during eyes open and eyes closed resting. Interactions that were significant (FDR-$p < 0.05$) are indicated with an *. We have also provided an indication of which pipeline of each pair provided larger values for variance explained using – and + symbols, which can be interpreted as the pipeline listed on the left of the heatmap having shown less (-) or more (+) variance explained in the comparison between the two experimental conditions than the pipeline listed at the bottom of the heatmap.

##### *Variance Explained by the Difference in Alpha Power Between WM Periods*

We have provided below statistical comparisons of the ability of the different pipelines to differentiate experimental conditions by examining the interactions between different pairs of pipelines and a condition of interest – the Sternberg working memory retention period vs Sternberg working memory probe period. We have provided heat maps depicting the variance explained (np$^2$) by this interaction for each pair of pipelines, marking the interactions that were significant (*). We have also provided an indication of which pipeline of each pair provided larger values for variance explained using – and + symbols, which can be interpreted as the pipeline listed on the left of the heatmap having shown less (-) or more (+) variance explained in the comparison between the two experimental conditions than the pipeline listed at the bottom of the heatmap.

Most pipelines showed alpha RMS values of 800 to 1100 for the working memory probe period and 2000 to 3000 for the working memory delay period, while HAPPE and wICA_all showed values more than two orders of magnitude lower, with values of 39.38 / 56.78 for HAPPE and 9.73/12.33 wICA_all respectively (see Figure S25). Given HAPPE performs wICA_all as a first step, then applies ICA_subtract, it struck us as odd to observe that HAPPE showed higher values than wICA_all (we would have assumed two iterations of artifact removal would lead to smaller values than one). The only potential explanation we could think of is that the ICA artifact identification algorithm used on the wICA_all cleaned data in HAPPE "expects" alpha activity, and as such marked components for subtraction that end up reconstructing the alpha activity. However, this is only conjecture and would require testing.

Figure S25 depicts the amount of variance explained by the difference between the working memory delay period and working memory probe period in alpha power RMS test across the different cleaning pipelines (which compared overall neural response strength from 250-1500ms after the stimuli) and the TANOVA from 0 to 750ms and 750 to 2000ms after the stimuli. Statistical comparisons of the overall interaction between pipelines and condition were highly significant for all three measures (all $p < 0.0001$). Post-hoc testing of the interaction between each pair of pipelines and the two conditions indicated the following rank order of the ability of the pipelines to discriminate between the experimental manipulation in alpha RMS: HAPPE > wICA_all > ASR[+], ICA_subtract*, wICA_ICLabel*, MWF_wICA_infomax[++**^], MWF_ICA_subtract[@**++], MWF_only[!@@**^^], MWF_CCA[!@@**++^^], MWF_wICA_AMICA [**++], MWF_wICA_CCA[@@**], MWF_wICA_fastICA[++**!!]. While the HAPPE and wICA methods explained considerable variance, the previous metric section indicated they also showed the lowest SER values (suggesting much of the signal was eliminated by these methods) and they reduced the alpha power in the signal by >2 orders of magnitude compared to the other pipelines (Figure S26). As such, we suspect these methods may enhance the differences between the WM delay period and probe periods by considerably reducing the alpha signal, and the high values of explained variance may be the result of low variability.

With regards to the TANOVA comparisons, two separate time periods showed a difference in the distribution of alpha activity between the WM delay and probe period in the Sternberg – first a large difference from 0-750ms, then a smaller difference from 750 to 2000ms. For the first time period (0 to 750ms), post-hoc testing of the interaction between each pair of pipelines and the two conditions indicated the following rank order of the ability of the pipelines to discriminate between the experimental manipulation in the distribution of alpha power: HAPPE*, wICA_ICLabel^, ICA_subtract^, wICA_all[+^^], MWF_only[@**^^++], MWF_CCA[@**^^++], ASR[!**^^++@@], MWF_wICA_AMICA[!**^^++@@], MWF_wICA_fastICA[**^^++@@!!], MWF_wICA_CCA[**^^++@@], MWF_wICA_infomax[**^^++@@], MWF_ICA_subtract[**^^++@@].

For the second time period examined with the TANOVA test (750 to 2000ms), a different pattern was apparent: ICA_subtract, wICA_ICLabel > wICA_all > MWF_only > MWF_CCA > MWF_wICA_CCA, MWF_wICA_infomax, MWF_wICA_fastICA, MWF_wICA_AMICA, MWF_ICA_subtract > HAPPE. We did not list ASR in this ranking, because although ASR ranked highly in terms of its variance explained (between wICA_all and MWF_only in terms of absolute value of variance explained for the comparison of the distribution of activity between probe and retention period alpha power), it only showed a significant difference compared to wICA_all and HAPPE, perhaps reflecting the high variability in results from this pipeline, similar to the result seen with the number of epochs marked for rejection. Unfortunately, we are not able to discern whether this variability reflects ground truth differences in individual alpha power (in which case the variability is valuable), or variability produced by the ASR cleaning (in which case the variability is artifactual). Additionally, while most pipelines showed a very similar distribution of alpha activity, HAPPE and wICA_all showed a different distribution of activity to all other pipelines, suggesting (alongside the alpha RMS, SER, and ARR values) that these two pipelines may be overcleaning the data, removing alpha power from specific electrodes such that the distribution of activity after cleaning is significantly altered (see Figure S29-32).

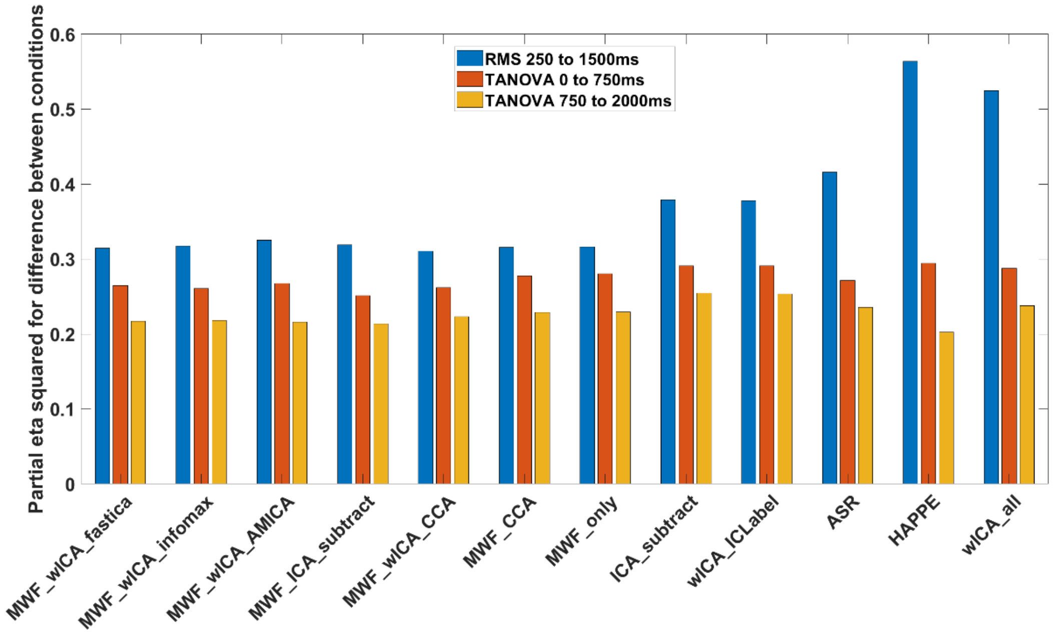

Figure S25. The variance explained (np²) by differences between alpha activity during the working memory delay and probe periods of the Sternberg task from each pipeline.

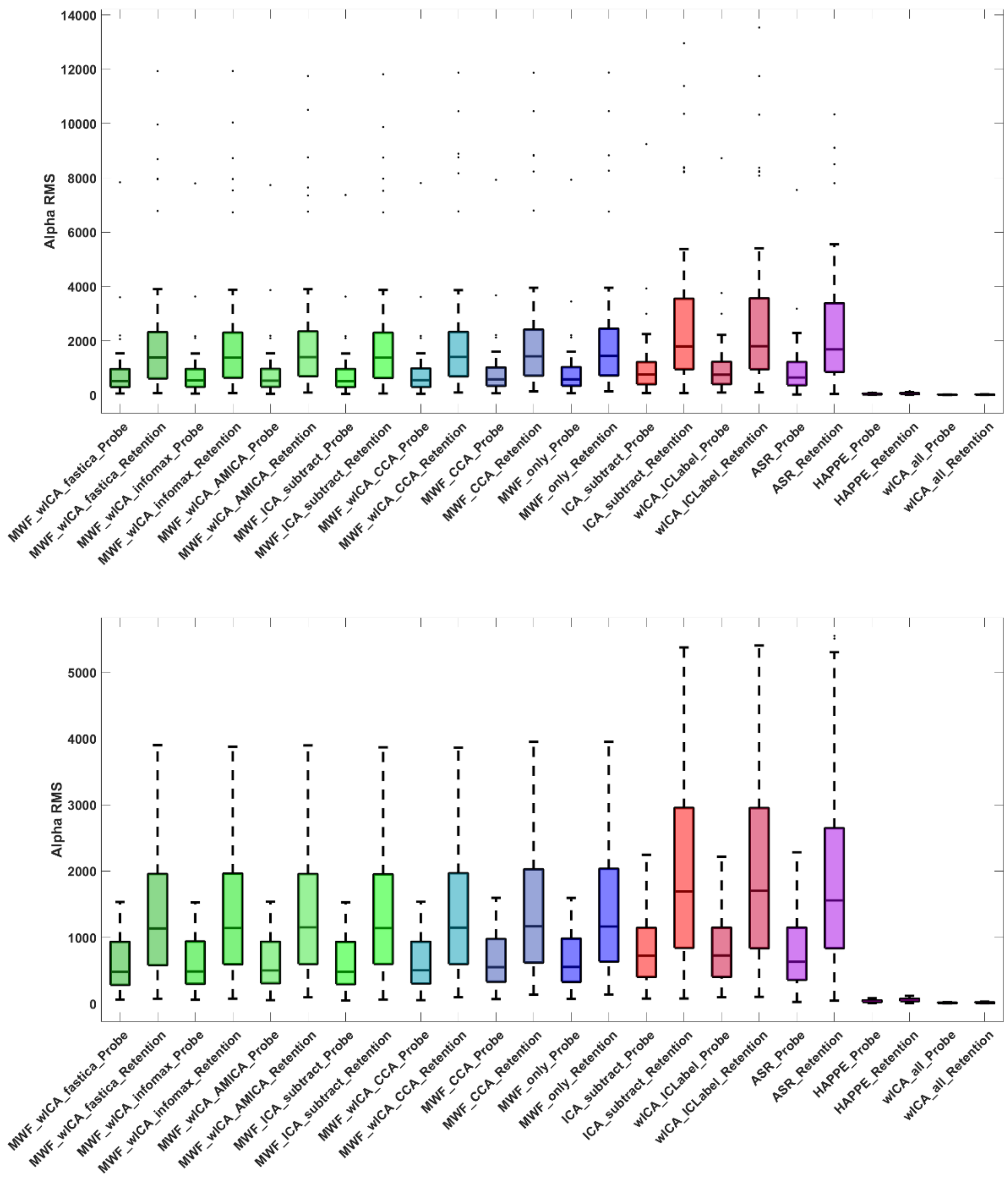

Figure S26. Alpha RMS from the working memory delay period (retention) and working memory probe period (probe) across each of the cleaning pipelines. Above, with outliers included, and below with outliers removed for better ability to distinguish the pipelines.

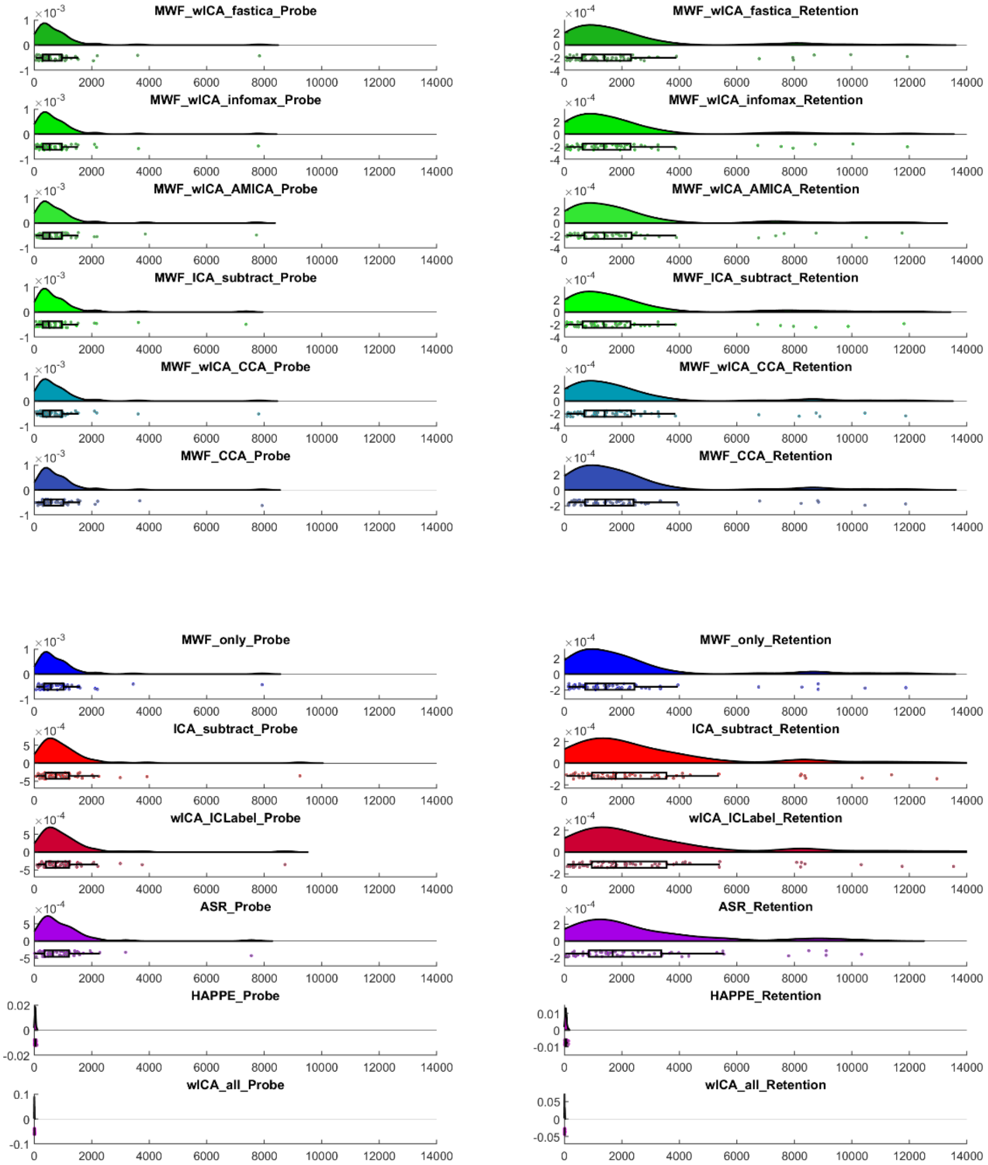

Figures S27. Raincloud plots for the alpha RMS working memory retention and probe periods for each pipeline.

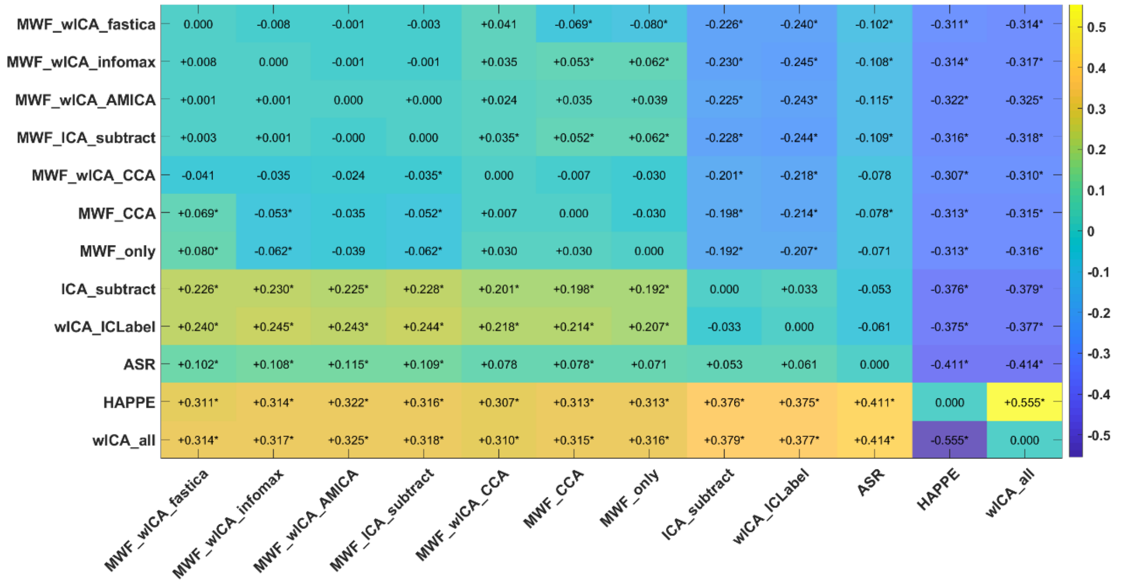

Figure S28. Heat map of the variance explained (np$^2$) by the interaction between each pair of pipelines and alpha power RMS during the Sternberg retention period vs the Sternberg probe period (from 250 to 1500ms after stimuli presentation). Interactions that were significant (FDR-p < 0.05) are indicated with an *. We have also provided an indication of which pipeline of each pair provided larger values for variance explained using – and + symbols, which can be interpreted as the pipeline listed on the left of the heatmap having shown less (-) or more (+) variance explained in the comparison between the two experimental conditions than the pipeline listed at the bottom of the heatmap.

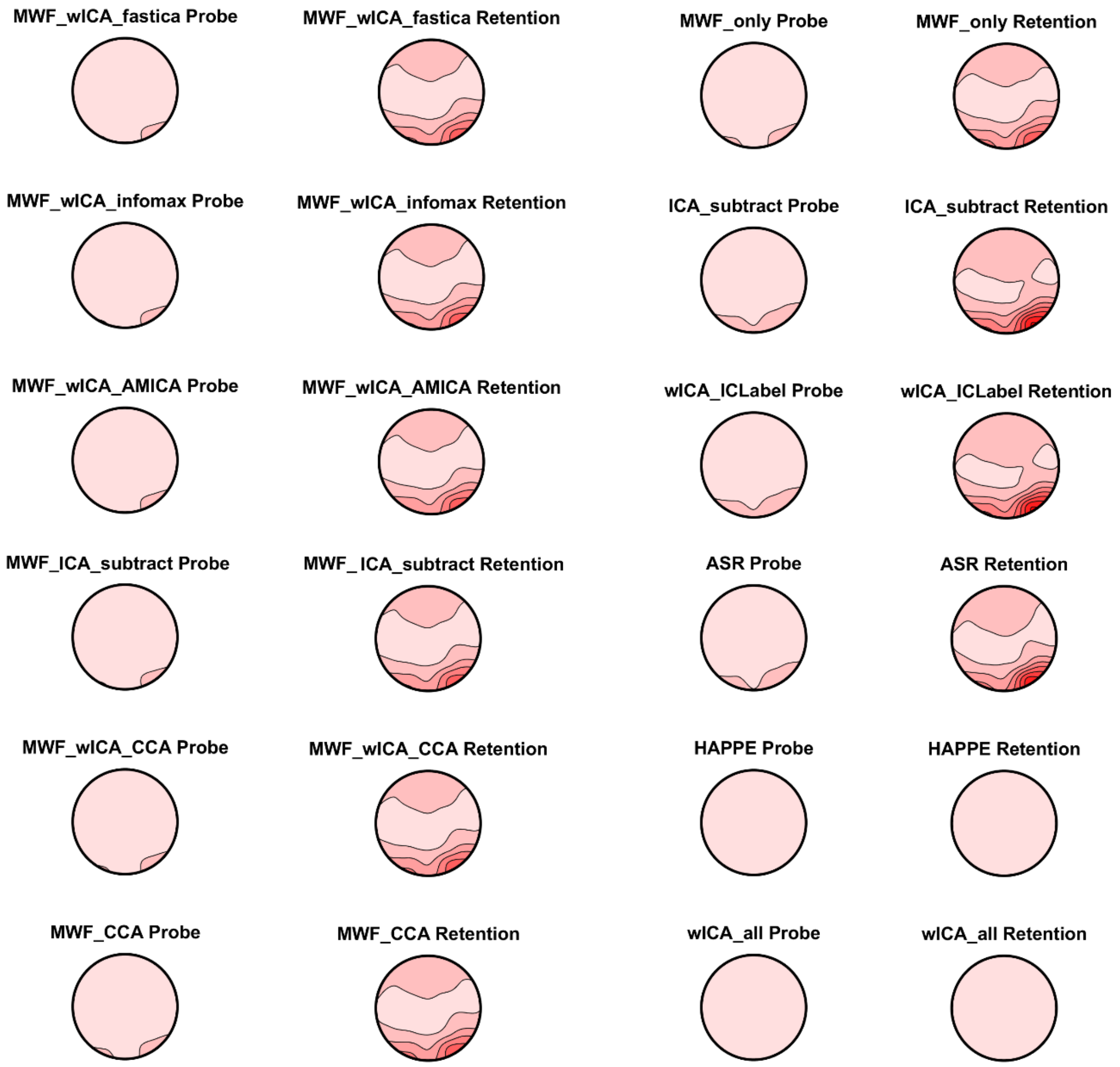

Figure S29. Alpha power distribution during the early period (0 to 750ms) after the stimuli of the working memory delay (retention) and working probe periods from each of the cleaning pipelines. All plots are on the same scale so they can all be compared to all other pipelines. Note that wICA_all and HAPPE have removed almost all alpha activity compared to the other pipelines.

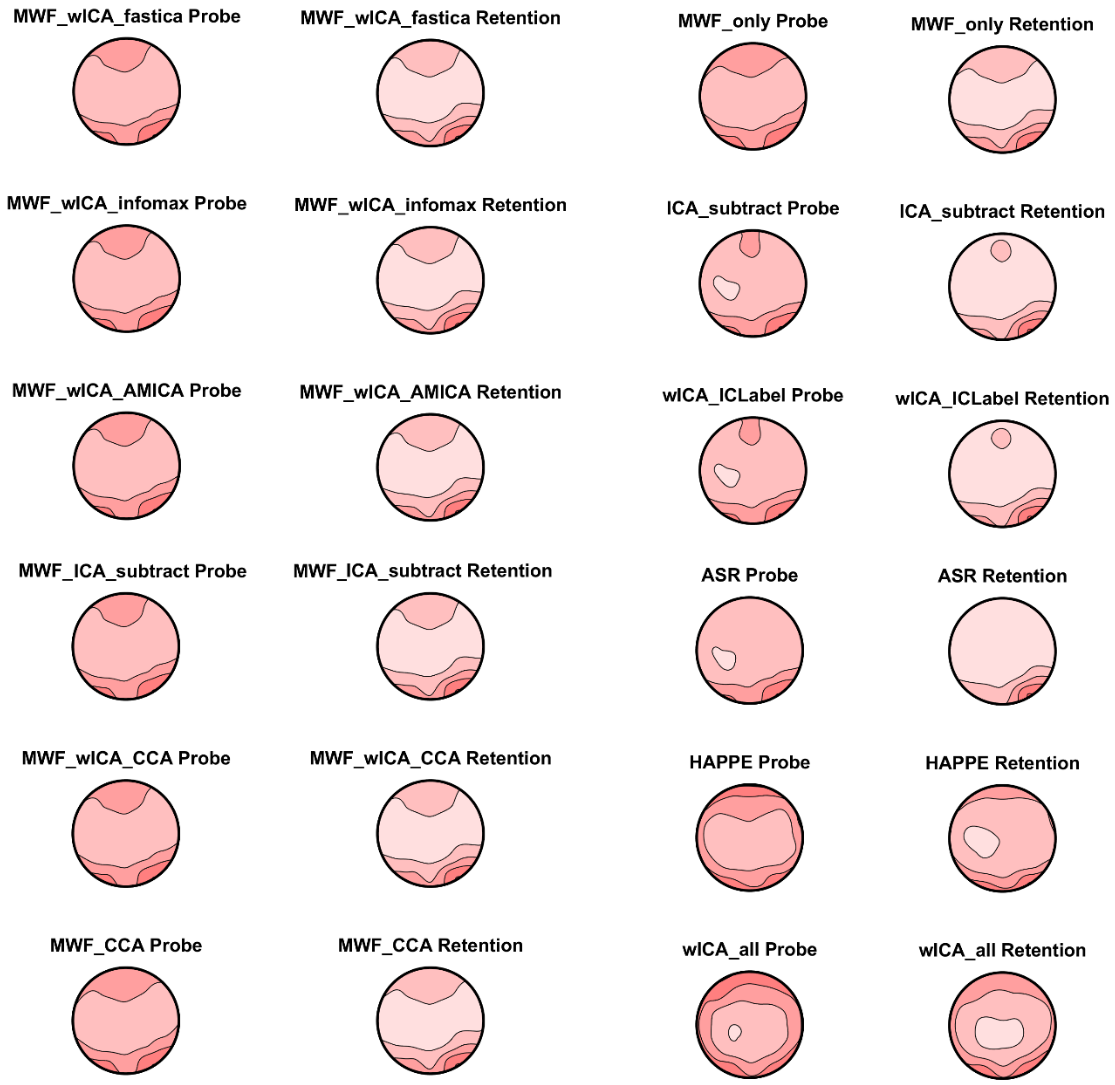

Figure S30. Alpha power distribution during the early period (0 to 750ms) after the stimuli of the working memory delay (retention) and working probe periods from each of the cleaning pipelines. All plots are on their own individual scale so the pattern of alpha activity distributions can be viewed within each plot. As expected, the retention period showed more occipital / parietal alpha maximums, while the probe period showed more widespread alpha. Note the similarity in pattern across most pipelines, including ICA only and MWF only methods, implying different cleaning approaches still lead to similar patterns. ASR, HAPPE, and wICA_all were the most different to the other cleaning pipelines, with ASR removing most of the frontal alpha, and HAPPE and wICA showing less qualitative differentiation between the probe and retention period alpha, despite these pipelines showing the highest variance explained for the comparison between the two conditions. We suspect this may be the result of removal of significant amount of the variance by these pipelines, leading to highly precise estimates of the retention and probe alpha activity, which allow strong inferences about the differences in these periods (which no longer overlap after the removal of so much variance). However, we suspect this comes at the cost of distorting the distribution/characterization of the neural activity.

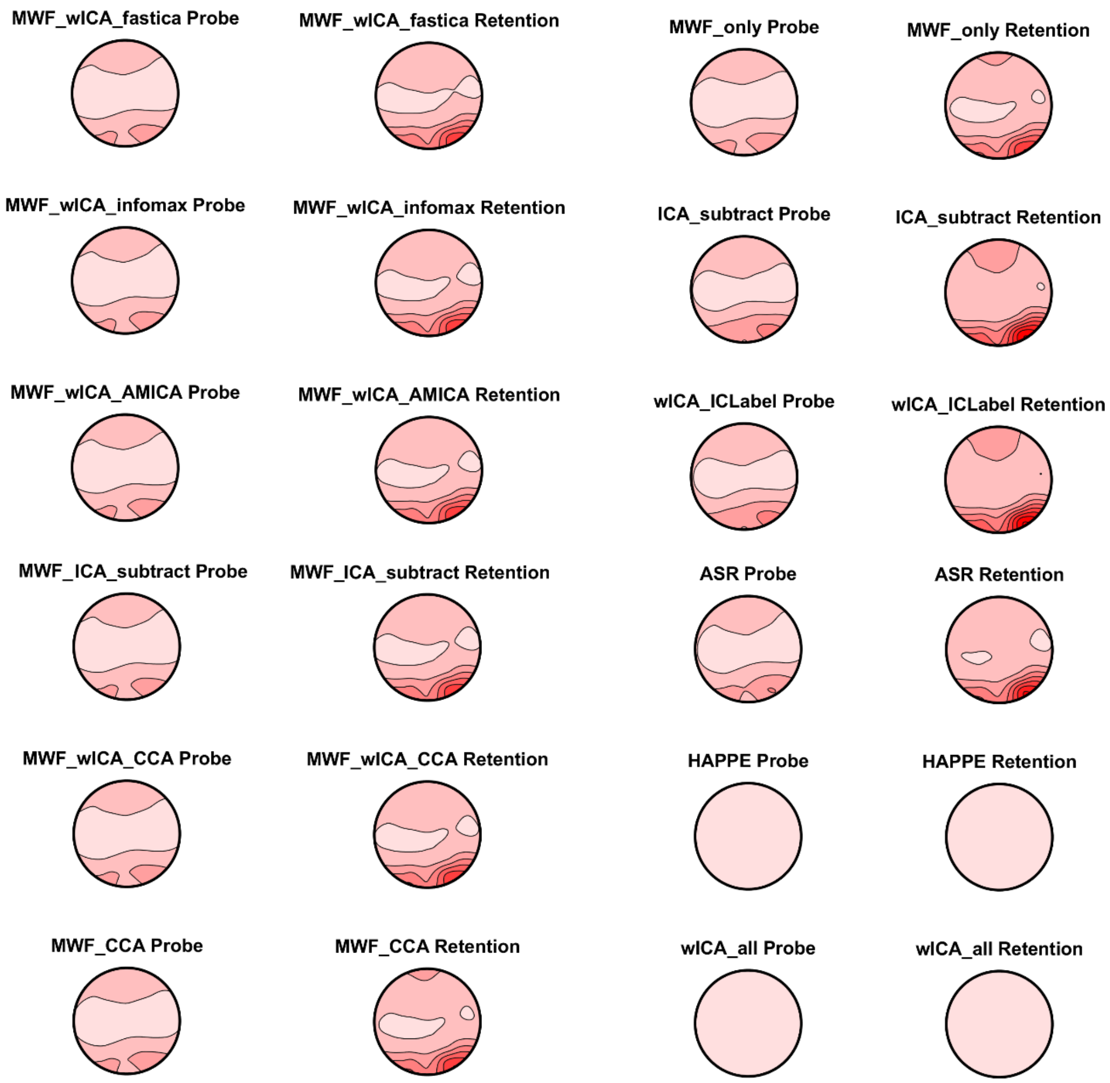

Figure S31. Alpha power distribution during the later period (750 to 2000ms) after the stimuli of the working memory delay (retention) and working probe periods from each of the cleaning pipelines. All plots are on the same scale so they can all be compared to all other pipelines. Note that wICA_all and HAPPE have removed almost all alpha activity compared to the other pipelines.

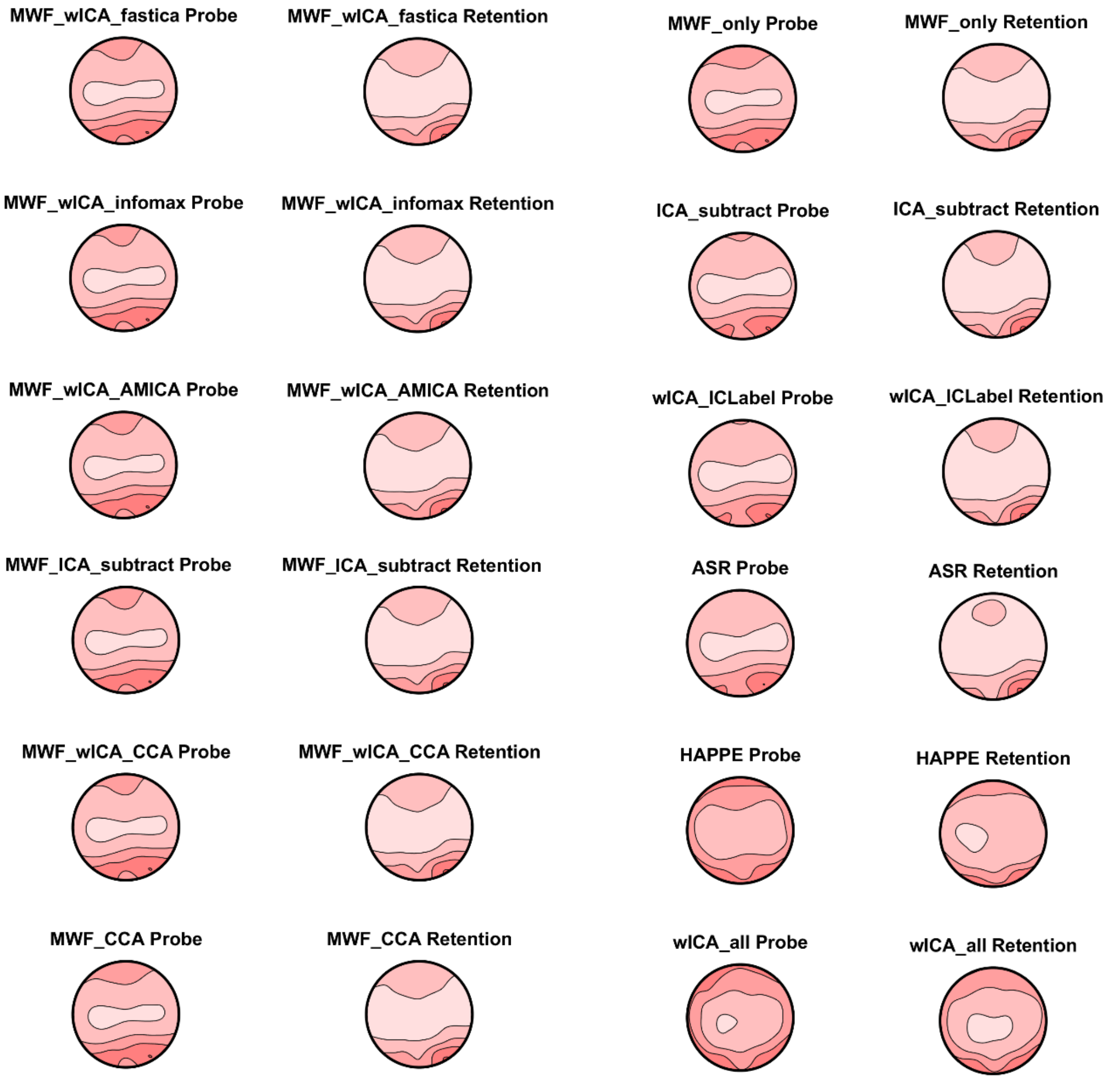

Figure S32. Alpha power distribution during the later period (750 to 2000ms) after the stimuli of the working memory delay (retention) and working probe periods from each of the cleaning pipelines. All plots are on their own individual scale so the pattern of alpha activity distributions can be viewed within each plot. As expected, the retention period showed more occipital / parietal alpha maximums, while the probe period showed more widespread alpha. As with the early period alpha, note the similarity in pattern across most pipelines, including ICA only and MWF only methods, implying different cleaning approaches still lead to similar patterns. Again, ASR, HAPPE, and wICA_all were the most different to the other cleaning pipelines, with ASR removing most of the frontal alpha, and HAPPE and wICA showing less qualitative differentiation between the probe and retention period alpha.

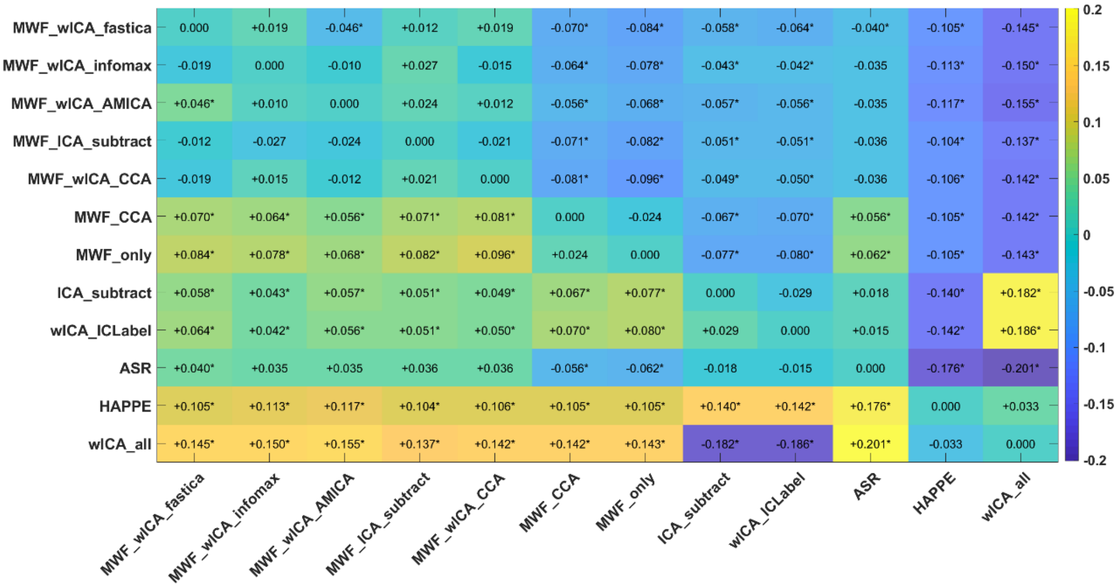

Figure S33. Heat map of the variance explained (np²) by the interaction between each pair of pipelines and alpha power TANOVA during the Sternberg retention period vs the Sternberg probe period (from 0 to 750ms after stimuli presentation). Interactions that were significant (FDR-p < 0.05) are indicated with an *. We have also provided an indication of which pipeline of each pair provided larger values for variance explained using – and + symbols, which can be interpreted as the pipeline listed on the left of the heatmap having shown less (-) or more (+) variance explained in the comparison between the two experimental conditions than the pipeline listed at the bottom of the heatmap.

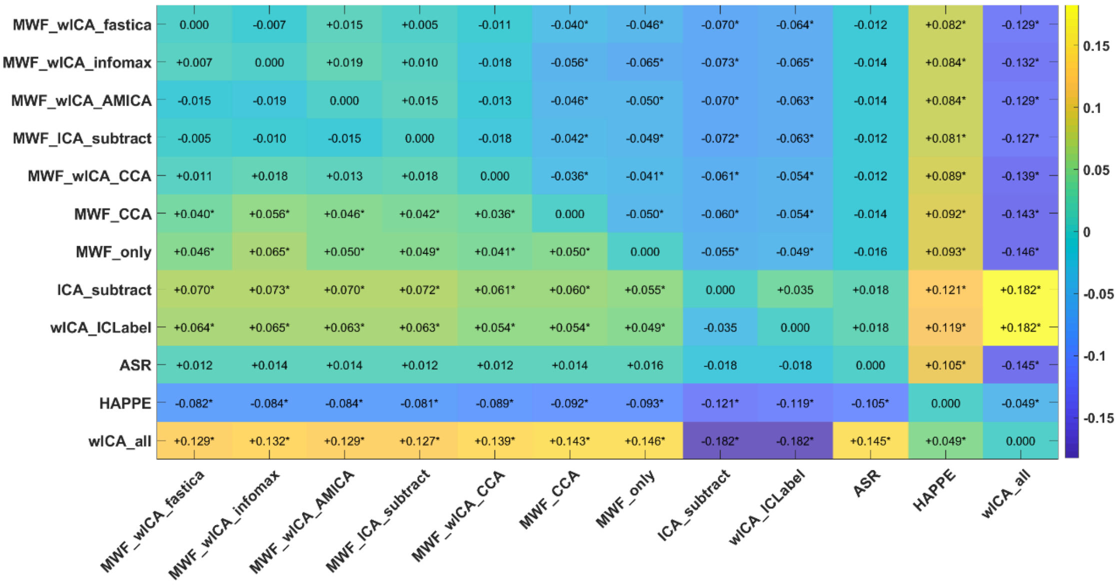

Figure S34. Heat map of the variance explained (np²) by the interaction between each pair of pipelines and alpha power TANOVA during the Sternberg retention period vs the Sternberg probe period (from 750 to 2000ms after stimuli presentation). Interactions that were significant (FDR-p < 0.05) are indicated with an *. We have also provided an indication of which pipeline of each pair provided larger values for variance explained using – and +

symbols, which can be interpreted as the pipeline listed on the left of the heatmap having shown less (-) or more (+) variance explained in the comparison between the two experimental conditions than the pipeline listed at the bottom of the heatmap.

#### SECTION FIVE

##### Analysis of a Combined EO, EC resting and 2back dataset

In addition to the analyses of datasets reported in our main manuscript, we submitted a smaller dataset which included 60 files from 20 participants, each providing an EO and EC resting recording, and a 2back task recording. This dataset was analysed to demonstrate the results from our larger datasets generalised to datasets with different recording parameters. In this dataset, data were recorded from 44 Ag/AgCl electrodes embedded within an EasyCap (Herrsching, Germany) which was connected to a Synamps2 amplifier running through the SCAN 4.3 software interface (Compumedics, Melbourne, Australia). The ground electrode was placed at AFz, and the reference was placed at CPz. A sampling rate of 1000 Hz was used for all recordings, with an online bandpass filter between 0.1 to 200 Hz.

##### *Signal-to-Error-Ratio*

There was a significant difference in SER between the pipelines: $F(1.88, 101.79) = 85.74$, $p < 0.001$. The rank order of significant differences between individual cleaning pipelines from post-hoc t-tests was as follows: MWF_only > wICA_ICLabel > MWF_wICA_AMICA^, MWF_wICA_infomax^, MWF_wICA_fastICA, ASR, ICA_subtract^^ > wICA_all, HAPPE. See Figure S35 for a raincloud plot depicting the distribution of the data. See Table S6 for means and SDs, as well as Figure S36 for a heatmap with confidence intervals for the post-hoc specification of which pipelines differed from which other pipelines, with significant differences highlighted.

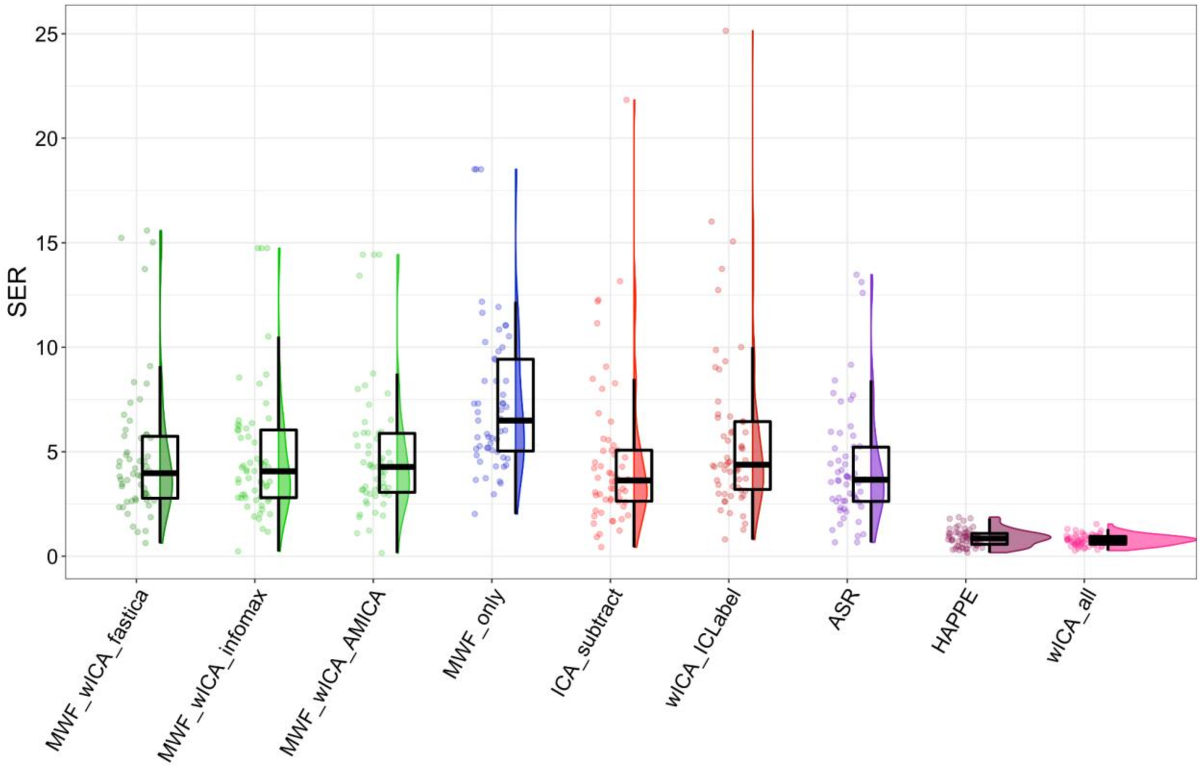

Figure S35. Raincloud plot depicting SER values from the combined EO and EC resting and 2back data (N = 20, file N = 60) for each of the cleaning pipelines.

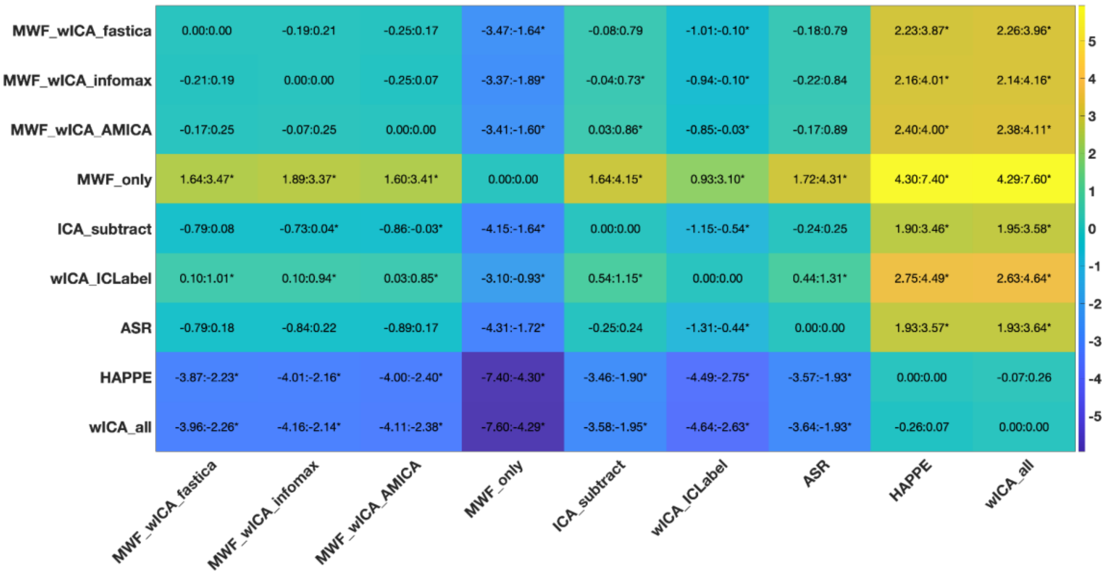

Figure S36. SER post-hoc tests for the combined 2back EO and EC resting dataset.

##### ***Artifact-to-Residue-Ratio***

There was a significant difference in ARR between the pipelines, with the robust ANOVA showing a significant effect: F(2.94, 102.8) = 449.203, $p < 0.0001$. The rank order of significant differences between individual cleaning pipelines from post-hoc t-tests was as follows: wICA_all > HAPPE > MWF_wICA_fastICA, MWF_wICA_infomax > MWF_wICA_AMICA > ASR > wICA_ICLabel, ICA_subtract > MWF_only. See Figure S37 for a raincloud plot depicting the distribution of the data. See Table S6 for means and SDs, as well as Figure S38 for a heatmap with confidence intervals for the post-hoc specification of which pipelines differed from which other pipelines, with significant differences highlighted. The combined resting and 2back datasets with reduced electrode montages showed an identical pattern to the combined Sternberg and resting dataset and combined Go Nogo datasets when SER and ARR values were viewed together, with ASR showing less SER and ARR, and ICA_subtract showing less ARR than the MWF_wICA methods.

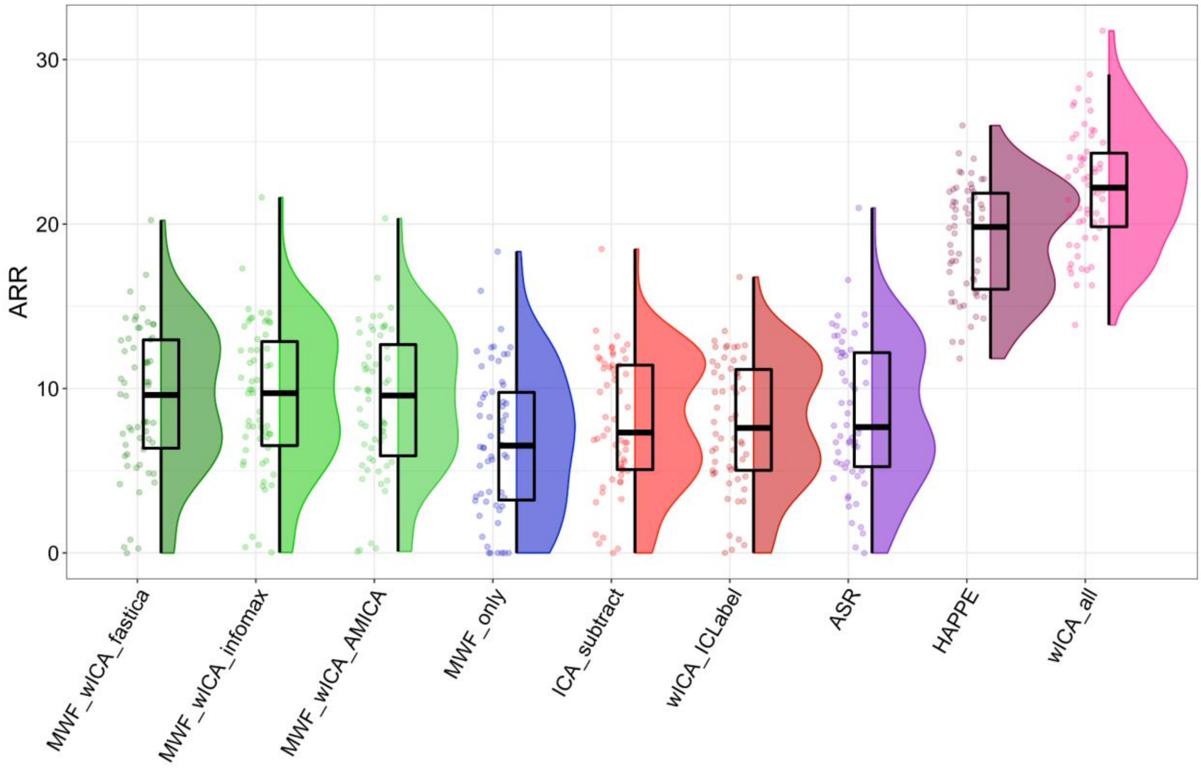

Figure S37. Raincloud plot depicting ARR values for the combined 2back EO and EC resting dataset.

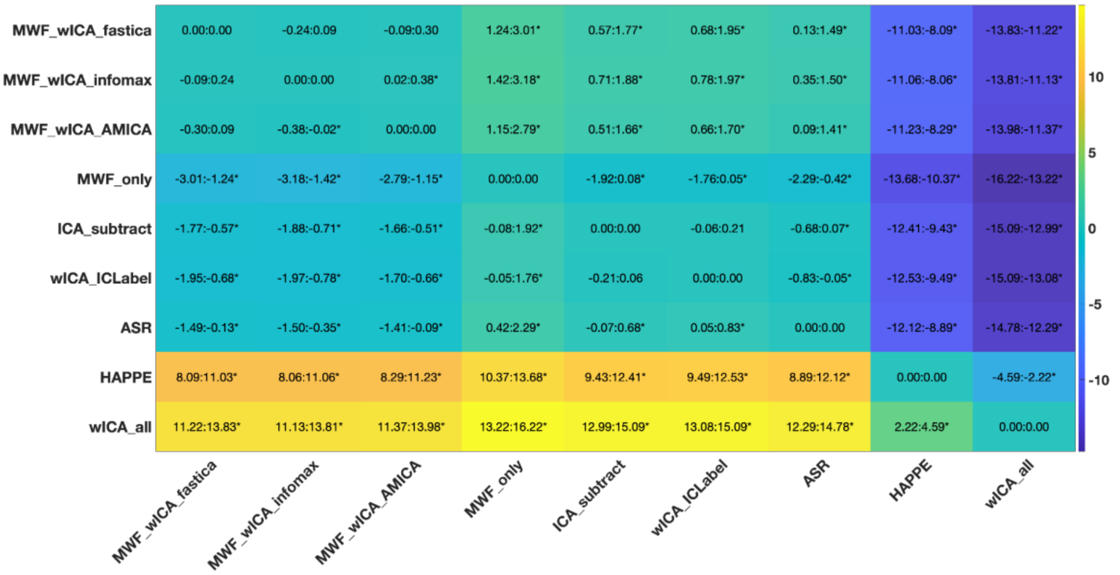

Figure S38. ARR post-hoc tests for the combined 2back EO and EC dataset.

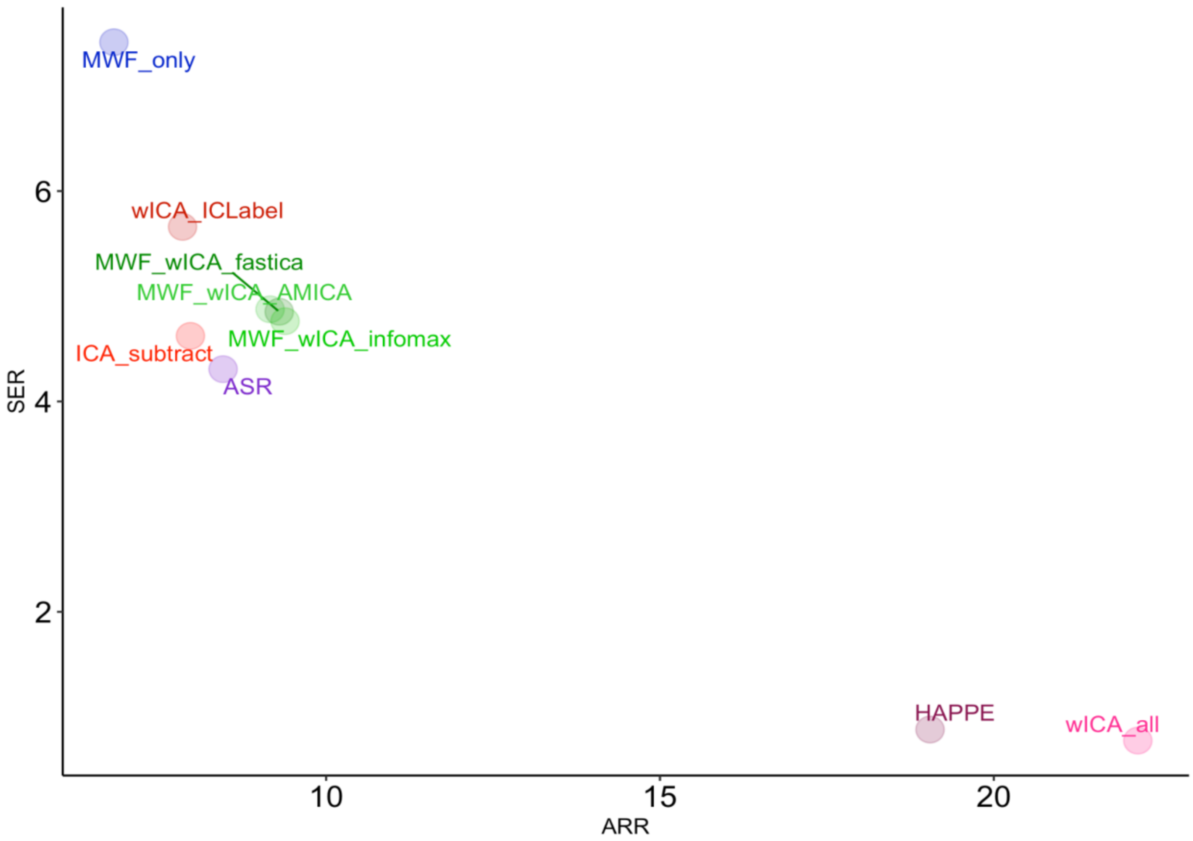

Figure S39**:** Scatterplot depicting the relationship between SER and ARR after cleaning for the combined 2back, EO and EC dataset. The HAPPE and wICA_all pipelines showed the strongest attenuation of activity within artifact periods. However, these pipelines also greatly reduced signal within the non-artifact periods. MWF_only provided the highest SER values, but also the lowest level of artifact reduction. The MWF_wICA methods appear to show a good trade-off between the amount of signal remaining in the data and the amount of artifact removed.

| | **SER** | | **ARR** | |
|---|---|---|---|---|
| **Pipeline** | **Mean** | **SD** | **Mean** | **SD** |
| **ASR** | 4.307 | 2.835 | 8.457 | 4.485 |
| **HAPPE** | 0.881 | 0.403 | 19.045 | 3.372 |
| **ICA_subtract** | 4.623 | 3.671 | 7.968 | 4.03 |
| **MWF_wICA_AMICA** | 4.876 | 3.121 | 9.161 | 4.355 |
| **MWF_wICA_fastICA** | 4.853 | 3.311 | 9.298 | 4.408 |
| **MWF_wICA_infomax** | 4.763 | 3.101 | 9.385 | 4.47 |
| **MWF_only** | 7.415 | 3.68 | 6.822 | 4.452 |
| **wICA_all** | 0.776 | 0.262 | 22.157 | 3.656 |
| **wICA_ICLabel** | 5.66 | 4.168 | 7.85 | 3.933 |

Table S6. Mean and SD values for the SER and ARR metrics for each cleaning pipeline from the combined 2back EO and EC dataset.

##### ***Blink Amplitude Ratio in Frontal Electrodes for the Combined 2back EO and EC Dataset***

There was a significant difference in blink amplitude ratio in frontal electrodes between the pipelines, with the robust ANOVA showing a significant effect: $F(3.33, 69.96) = 5.2241$, $p = 0.0019$. The rank order of significant differences between individual cleaning pipelines from post-hoc t-tests was as follows: MWF_wICA_fastICA^, MWF_wICA_infomax^, MWF_wICA_AMICA, ASR, MWF_only^^ > wICA_ICLabel, ICA_subtract. wICA_all did not significantly differ from any pipeline (despite showing the lowest values in the raincloud plot). See Figure S40 for a raincloud plot depicting the distribution of the data. See Table S7 for means and SDs, as well as Figure S41 for a heatmap with confidence intervals for the post-hoc specification of which pipelines differed from which other pipelines, with significant differences highlighted.

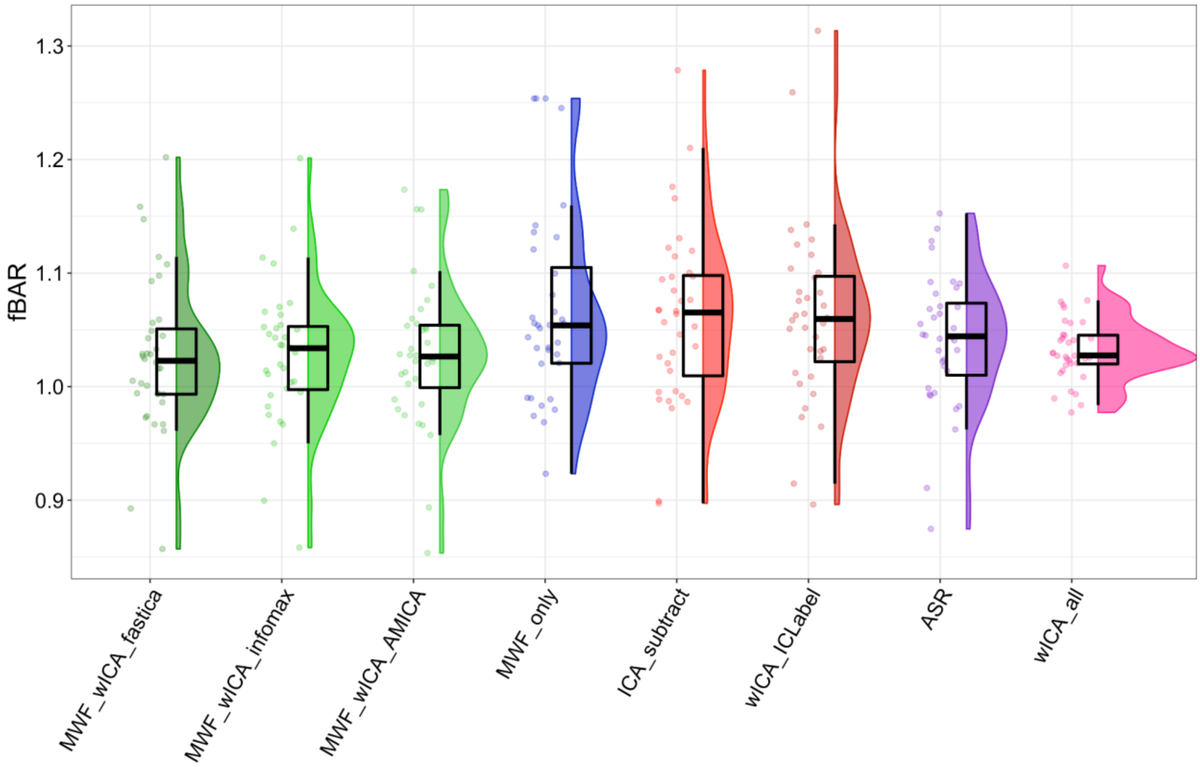

Figure S40. Raincloud plot depicting fBAR values from the combined EO and EC resting and 2back dataset.

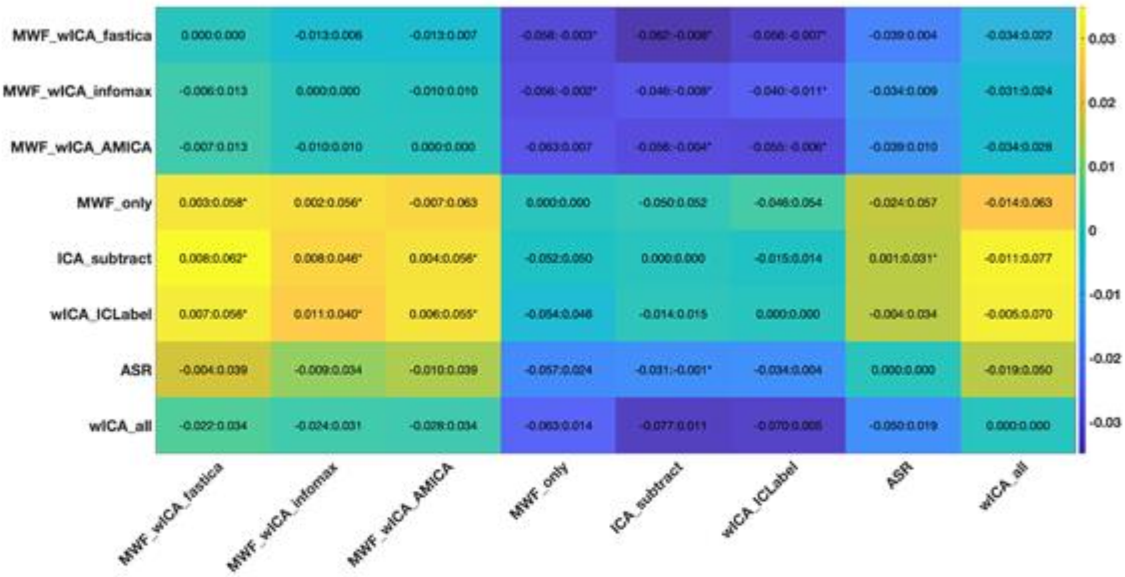

Figure S41. fBAR post-hoc tests from the combined EO and EC resting and 2back dataset.

***Blink Amplitude Ratio Across All Electrodes for the Combined 2back EO and EC Dataset***

There was a significant difference in blink amplitude ratio in all electrodes between the pipelines, with the robust ANOVA showing a significant effect: F(3.24, 68.02) = 9.193, $p < 0.0001$. The rank order of significant differences between individual cleaning pipelines from post-hoc t-tests was as follows: MWF_wICA_fastICA^, MWF_wICA_AMICA, MWF_wICA_infomax, ASR, MWF_only^^ > ICA_subtract, wICA_ICLabel. wICA_all did not significantly differ from any pipeline (despite showing the lowest values in the raincloud plot). See Figure S42 for a raincloud plot depicting the distribution of the data. See Table S7 for means and SDs, as well as Figure S43 for a heatmap with confidence intervals for the post-hoc specification of which pipelines differed from which other pipelines, with significant differences highlighted.

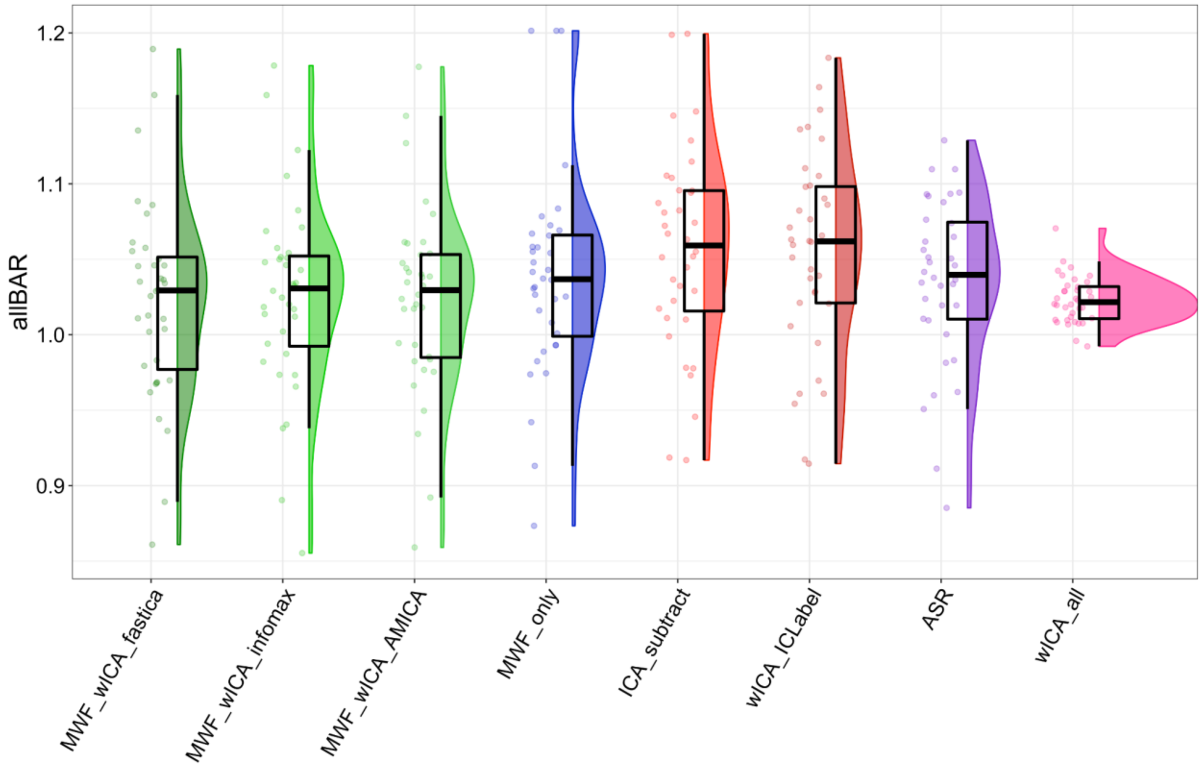

Figure S42. Raincloud plot depicting allBAR values from the combined EO and EC resting and 2back dataset.

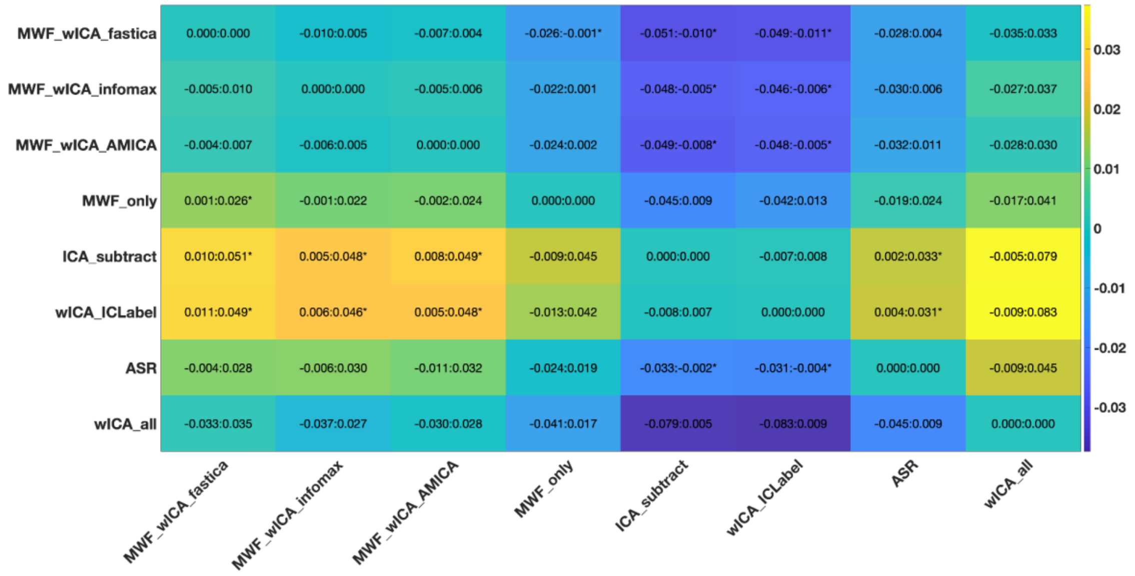

Figure S43. allBAR post-hoc tests for the combined 2back, EO and EC dataset.

|  | fBAR |  | allBAR |  |
|---|---|---|---|---|
| **Pipeline** | **Mean** | **SD** | **Mean** | **SD** |
| **ASR** | 1.041 | 0.059 | 1.035 | 0.055 |
| **ICA_subtract** | 1.061 | 0.078 | 1.056 | 0.068 |
| **MWF_wICA_AMICA** | 1.028 | 0.064 | 1.023 | 0.063 |
| **MWF_wICA_fastICA** | 1.027 | 0.068 | 1.022 | 0.067 |
| **MWF_wICA_infomax** | 1.028 | 0.062 | 1.026 | 0.065 |
| **MWF_only** | 1.068 | 0.084 | 1.038 | 0.069 |
| **wICA_all** | 1.032 | 0.028 | 1.023 | 0.016 |
| **wICA_ICLabel** | 1.062 | 0.08 | 1.055 | 0.066 |

Table S7. Blink Amplitude Ratio means and standard deviations for the combined 2back, EO and EC dataset.

***Proportion of Epochs Showing Muscle Activity Remaining After Cleaning from the Combined 2back, EO and EC Dataset***

There was a significant difference in number of epochs with muscle activity remaining between the pipelines: F(4.85, 121.26) = 36.96 , $p < 0.001$. The rank order of significant differences between individual cleaning pipelines from post-hoc t-tests was as follows: MWF_wICA_AMICA^, MWF_wICA_fastICA, MWF_wICA_infomax, ICA_subtract, wICA_ICLabel, MWF_only^^, ASR > wICA_all. See Figure S44 for a raincloud plot depicting the distribution of the data. See Table S8 for means and SDs, as well as Figure S45 for a heatmap with confidence intervals for the post-hoc specification of which pipelines differed from which other pipelines, with significant differences highlighted.

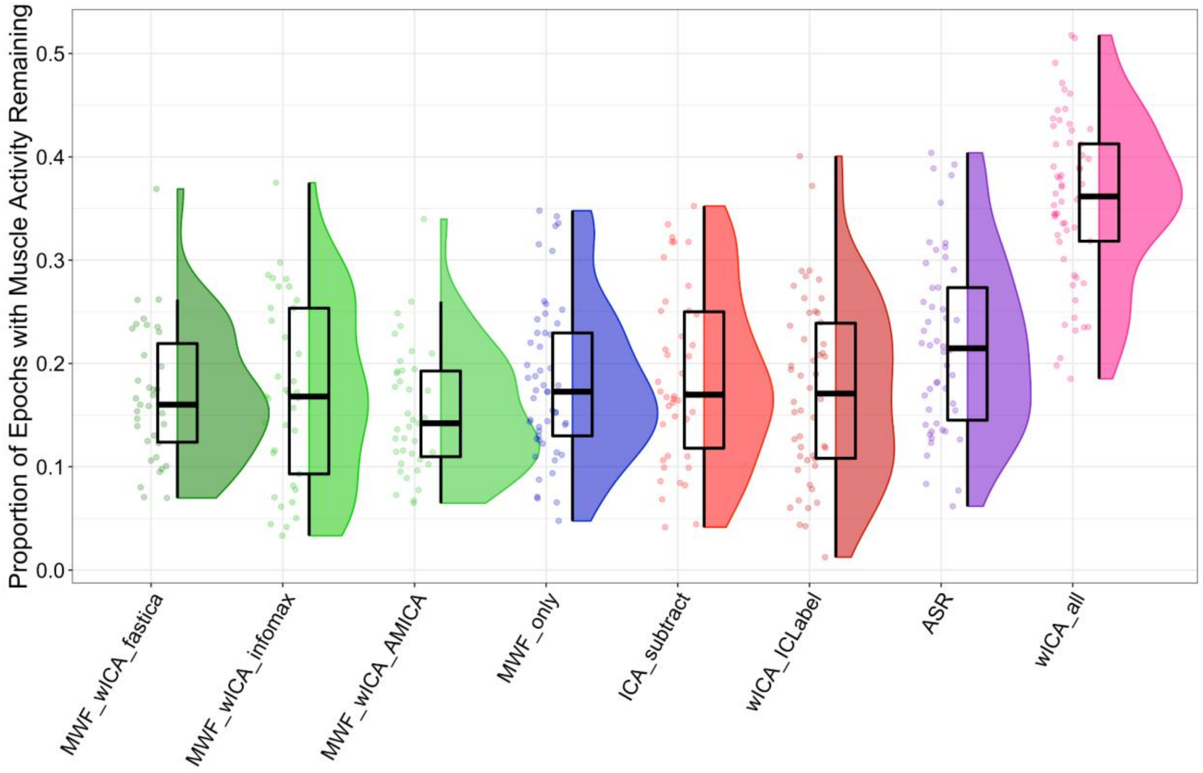

Figure S44. Raincloud plot depicting the proportion of epochs showing log-power log-frequency values above the -0.59 threshold from the combined 2back, EO and EC dataset.

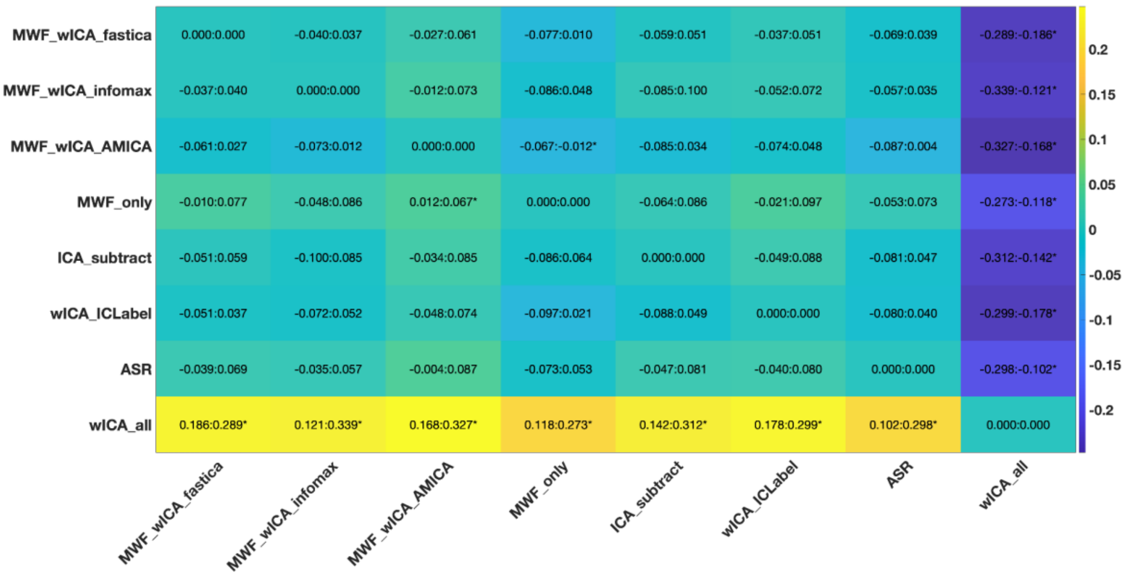

Figure S45. Proportion of epochs showing muscle activity after cleaning post-hoc tests from the combined 2back, EO and EC dataset.

***Severity by which Log-Power Log-Frequency Slopes Exceeded the Threshold from the combined 2back, EO and EC dataset***

There was a significant difference in the mean severity by which the slope exceeded the log-power log-frequency threshold from epochs and electrodes that showed muscle activity remaining between the pipelines, with the robust ANOVA showing a significant effect: F(1, 35) = 144.948, $p < 0.0001$. wICA_all performed worse than all other pipelines, but otherwise there were no differences across the pipelines. See Figure S46 for a raincloud plot depicting the distribution of the data. See Table S8 for means and SDs, as well as Figure S47 for a heatmap with confidence intervals for the post-hoc specification of which pipelines differed from which other pipelines, with significant differences highlighted.

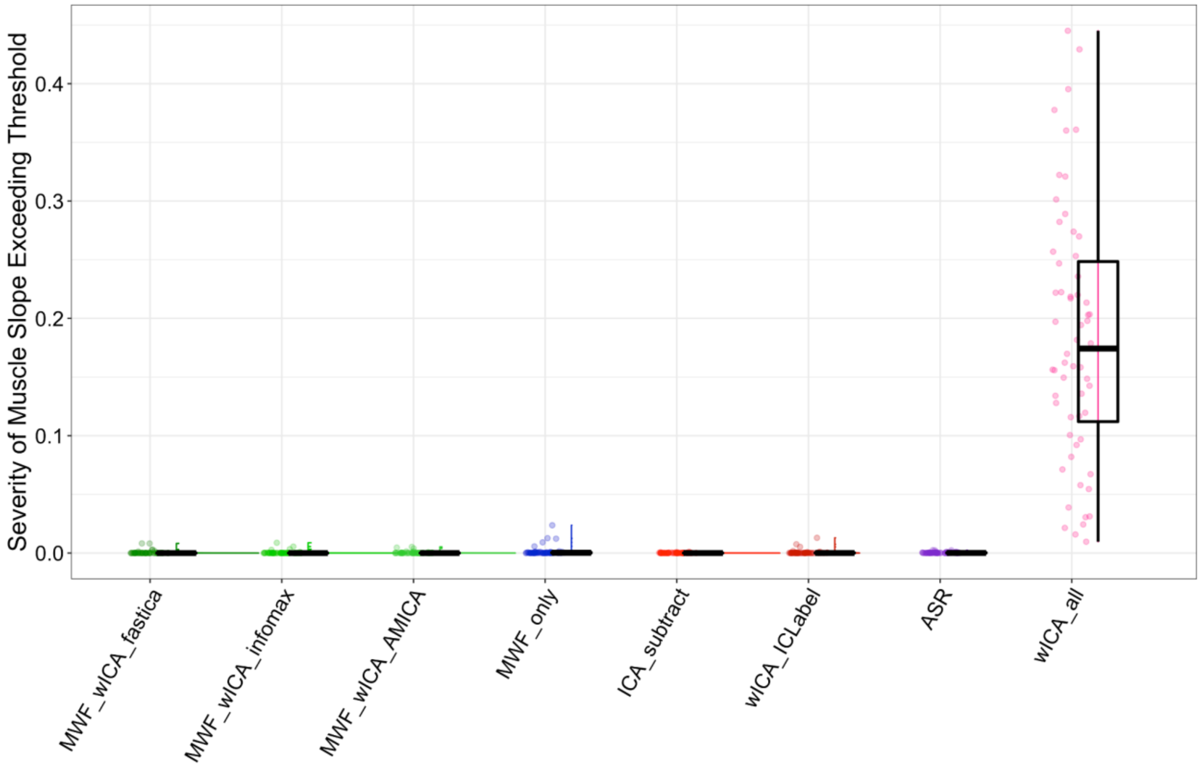

Figure S46. Raincloud plot for the slope above threshold in epochs that show slopes indicative of muscle activity after cleaning from the combined 2back, EO and EC dataset.

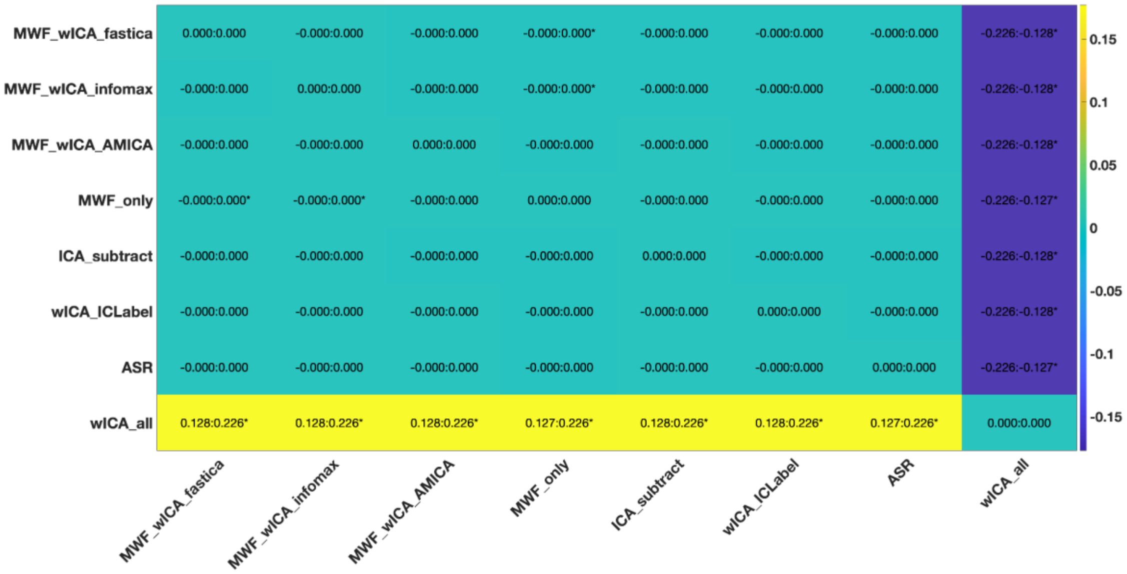

Figure S47. Post-hoc tests for the slope above threshold in epochs that show slopes indicative of muscle activity after cleaning from the combined 2back, EO and EC dataset.

| | Proportion of epochs showing slopes that indicate muscle activity remaining after cleaning | | Severity of slope that exceeded muscle slope threshold in epochs showing muscle slopes after cleaning | |
|---|---|---|---|---|
| **Pipeline** | **Mean** | **SD** | **Mean** | **SD** |
| **ASR** | 0.218 | 0.087 | 0 | 0.001 |
| **ICA_subtract** | 0.187 | 0.084 | 0 | 0 |
| **MWF_wICA_AMICA** | 0.154 | 0.061 | 0 | 0.001 |
| **MWF_wICA_fastICA** | 0.169 | 0.065 | 0 | 0.002 |
| **MWF_wICA_infomax** | 0.171 | 0.088 | 0 | 0.001 |
| **MWF_only** | 0.181 | 0.075 | 0.001 | 0.004 |
| **wICA_all** | 0.358 | 0.079 | 0.184 | 0.108 |
| **wICA_ICLabel** | 0.174 | 0.087 | 0.001 | 0.002 |

Table S8. Means and SDs for muscle activity remaining after cleaning from the combined 2back, EO and EC dataset.

##### Analysis of a Colour-Wheel Recall task dataset

Finally, one further dataset of Colour-Wheel Recall task related data was tested, which used different recording parameters to the datasets reported in the main manuscript (datasets were recorded using a Neuroscan amplifier [Compumedics, Melbourne, Australia] with a 62-electrode EASYCAP and a sampling rate of 10kHz, downsampled to 1000Hz). To reduce computation time, MWF_wICA_45Hz, MWF_ICA_subtract, MWF_CCA and MWF_wICA_CCA were not tested on this data.

###### *Signal-to-Error-Ratio in the Colour Wheel Task dataset*

There was a significant difference in SER between the pipelines; F(1.56, 32.82) = 38.91, $p < 0.0001$. The rank order of significant differences between individual cleaning pipelines from post-hoc t-tests was as follows: MWF_only > MWF_CCA > wICA_ICLabel > MWF_wICA_AMICA*, MWF_wICA_infomax*, MWF_wICA_fastICA, ICA_subtract, ASR** > HAPPE, wICA_all. See Figure S48 for the raincloud plot and S49 for the post-hoc tests.

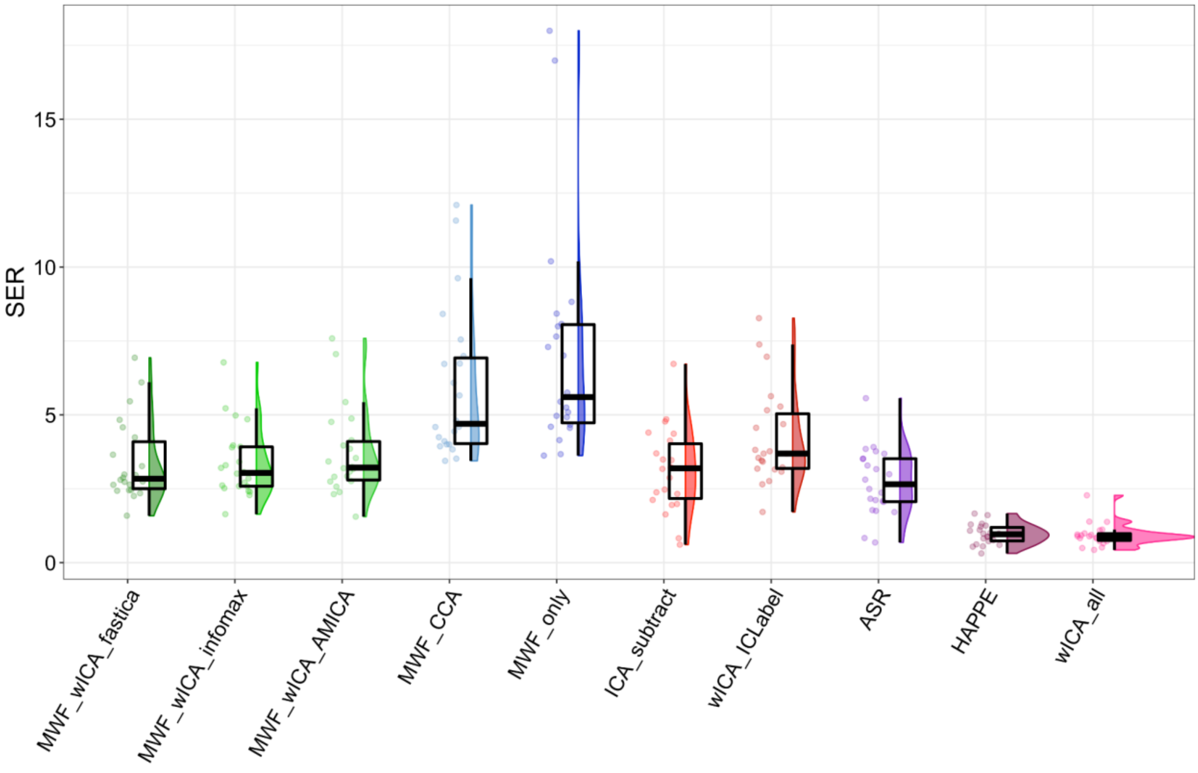

Figure S48. Raincloud plot depicting SER values for the Colour Wheel Task dataset.

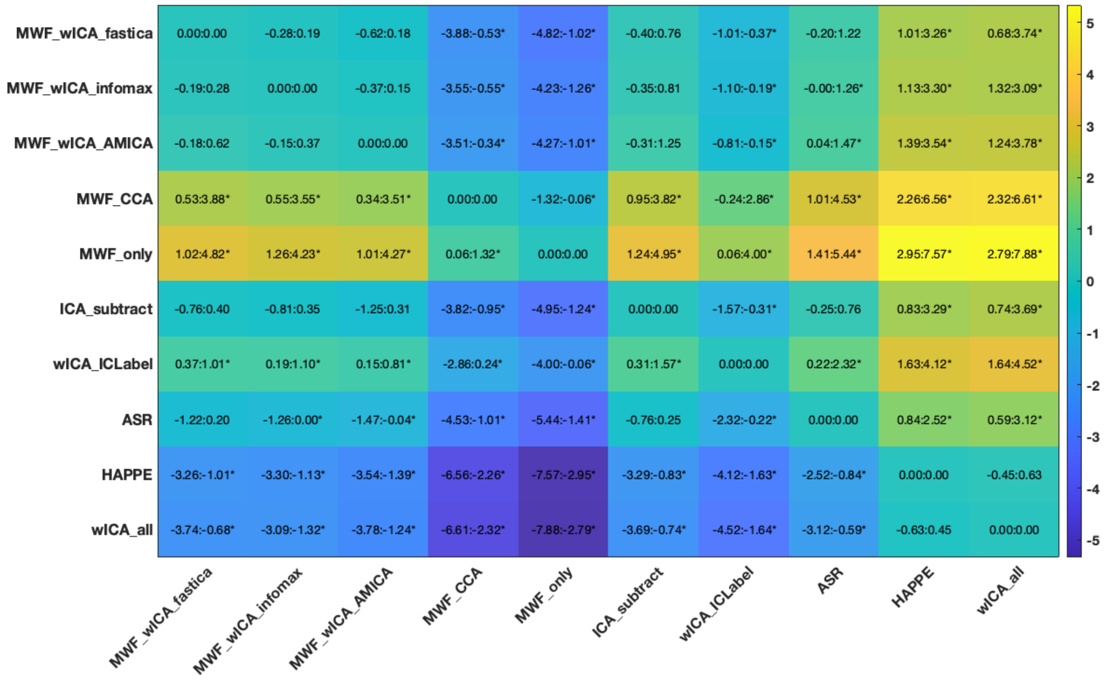

Figure S49. Post-hoc tests for the SER values for the Colour Wheel Task dataset.

##### ***Artifact-to-Residue-Ratio in the Colour Wheel Task dataset***

There was also a significant difference in ARR between the pipelines; F(3, 63.1) = 174.004, *p* < 0.0001. The rank order of significant differences between individual cleaning pipelines from post-hoc t-tests was as follows: wICA_all > HAPPE > MWF_wICA_fastICA*^$^{+}$, MWF_wICA_AMICA^$^{+}$, MWF_wICA_infomax^$^{+}$, ASR**$^{+}$, ICA_subtract^^, wICA_ICLabel$^{++}$, MWF_CCA^^$^{@}$, MWF_only^^$^{@@}$.

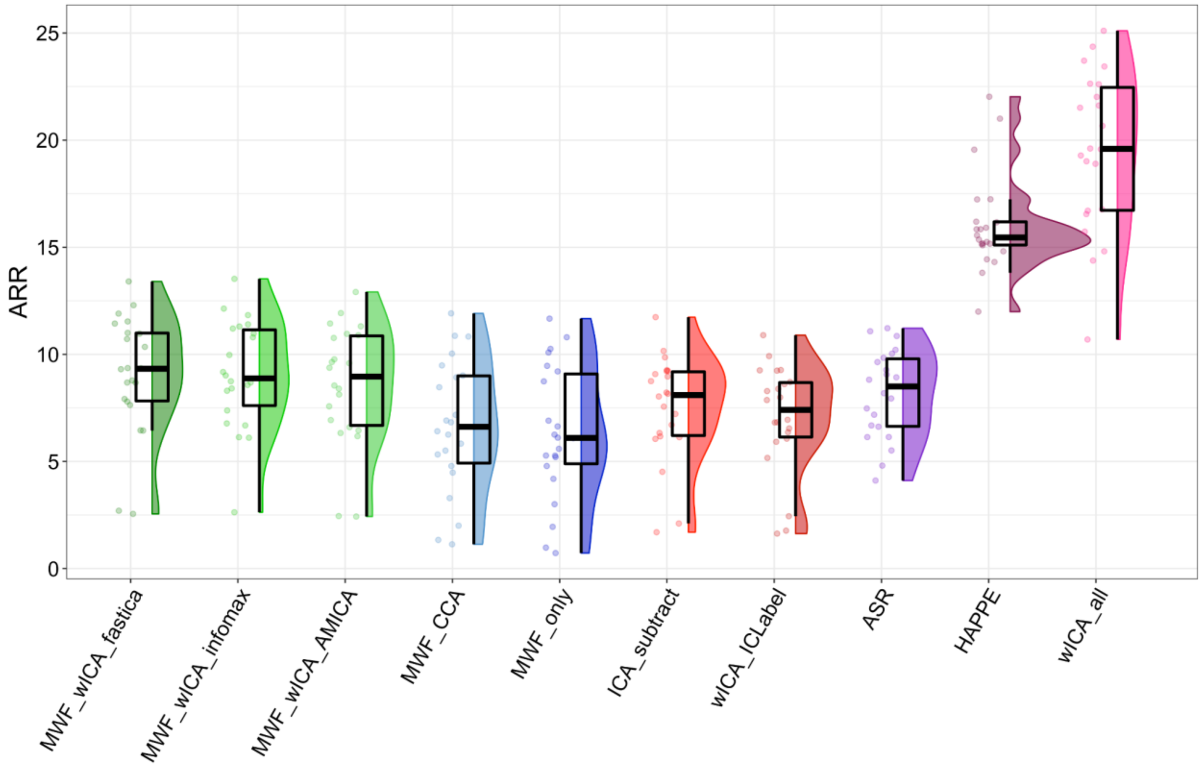

Figure S50. Raincloud plot depicting ARR values for the Colour Wheel Task dataset.

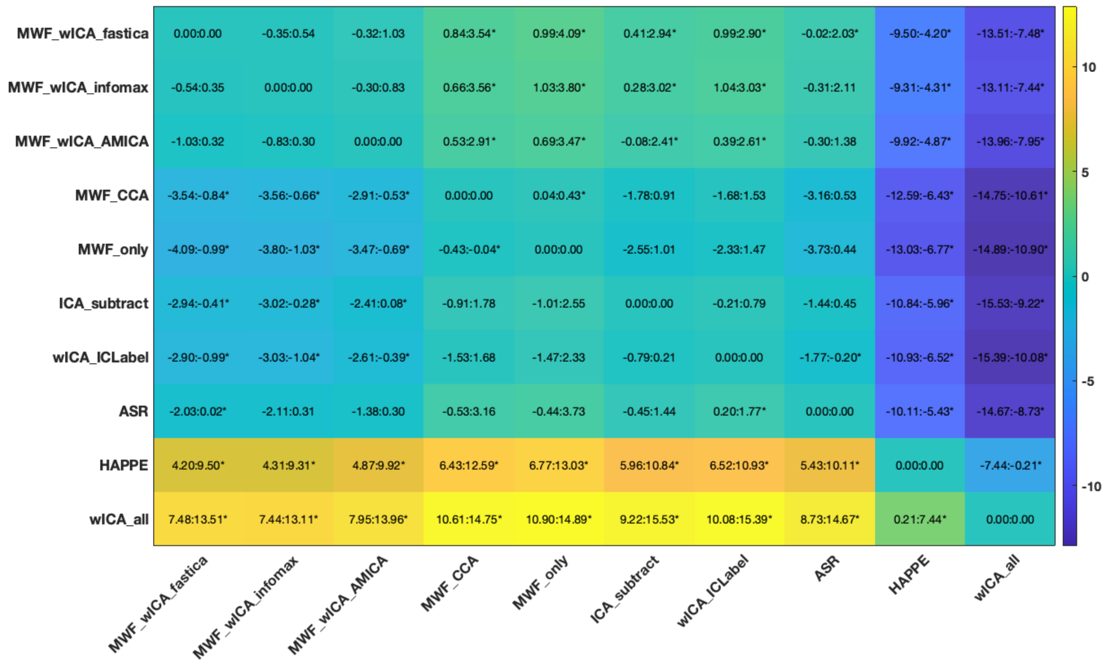

Figure S51. Post-hoc tests for the ARR values for the Colour Wheel Task dataset.

| | ARR | | SER | |
|---|---|---|---|---|
| **Pipeline** | **Mean** | **SD** | **Mean** | **SD** |
| **ASR** | 8.17 | 2.068 | 2.732 | 1.132 |
| **HAPPE** | 16.05 | 2.276 | 0.961 | 0.343 |
| **ICA_subtract** | 7.507 | 2.455 | 3.122 | 1.424 |
| **MWF_CCA** | 6.673 | 3.09 | 5.952 | 2.543 |
| **MWF_wICA_AMICA** | 8.572 | 2.797 | 3.704 | 1.479 |
| **MWF_wICA_fastICA** | 9.057 | 2.786 | 3.408 | 1.374 |
| **MWF_wICA_infomax** | 9.079 | 2.522 | 3.374 | 1.2 |
| **MWF_only** | 6.326 | 3.115 | 7.142 | 3.812 |
| **wICA_all** | 19.53 | 3.708 | 0.935 | 0.385 |
| **wICA_ICLabel** | 7.042 | 2.514 | 4.231 | 1.637 |

Table S9. Means and SDs for the SER and ARR from the Colour Wheel task.

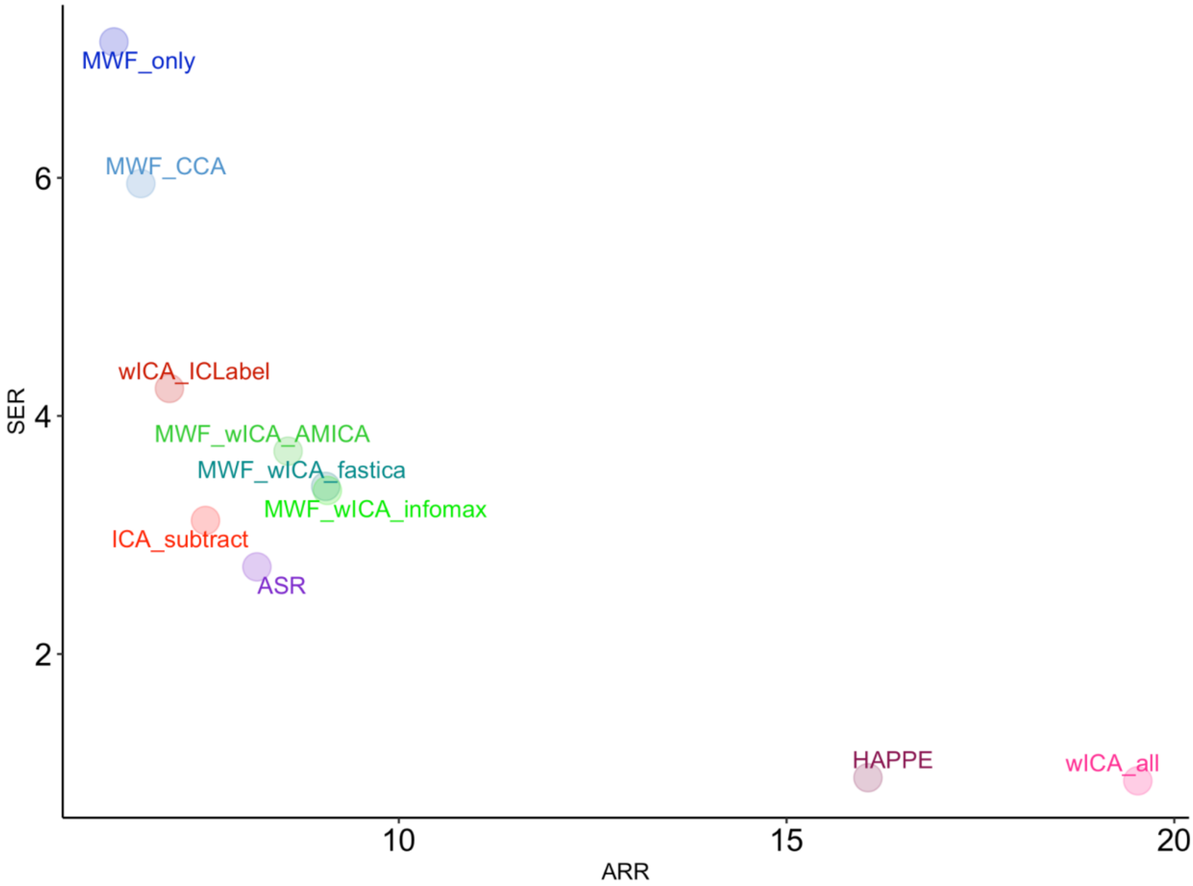

Figure S52. Scatter plot depicting the relationship between SER and ARR values for the Colour Wheel task dataset from each pipeline.

##### ***Blink Amplitude Ratio in the Colour Wheel Task dataset***

A significant difference was found in fBAR between the pipelines; F(2.09, 41.79) = 3.923, $p$ = 0.026. However, after controlling for multiple comparisons, no single pipeline showed higher values than any other pipeline in the post-hoc tests.

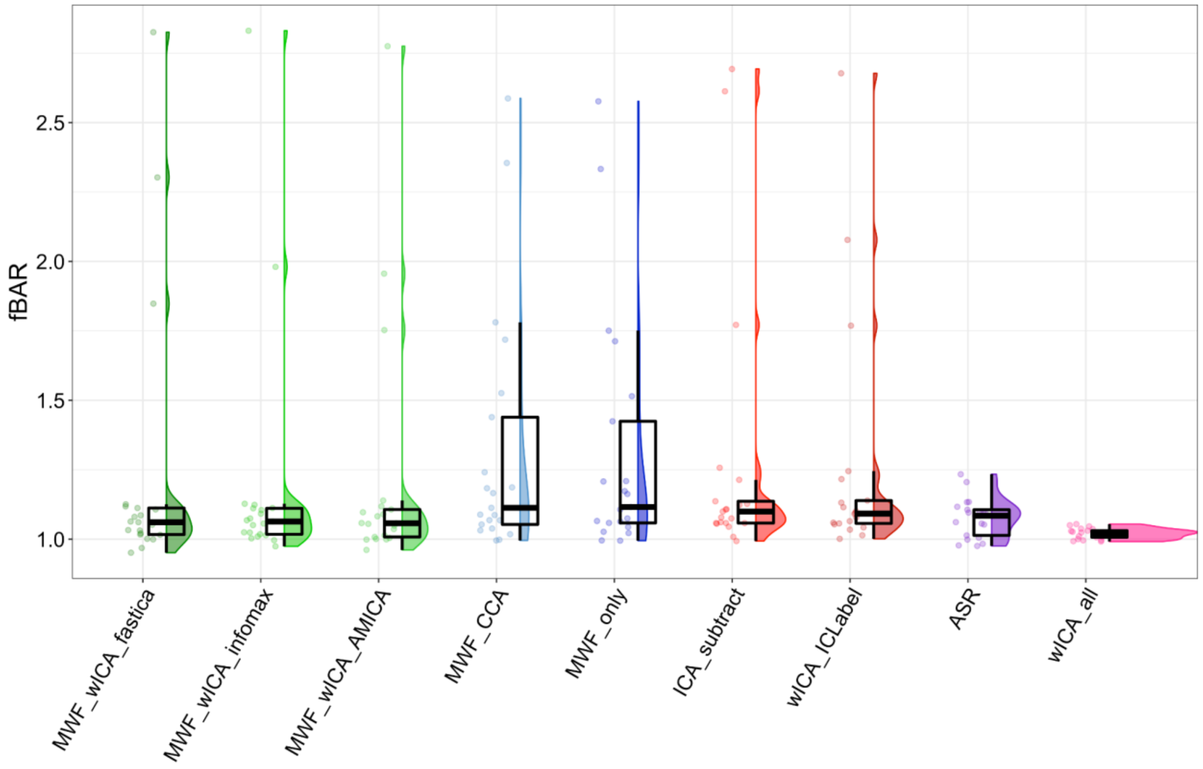

Figure S53. Raincloud plot depicting fBAR values for the Colour Wheel Task dataset.

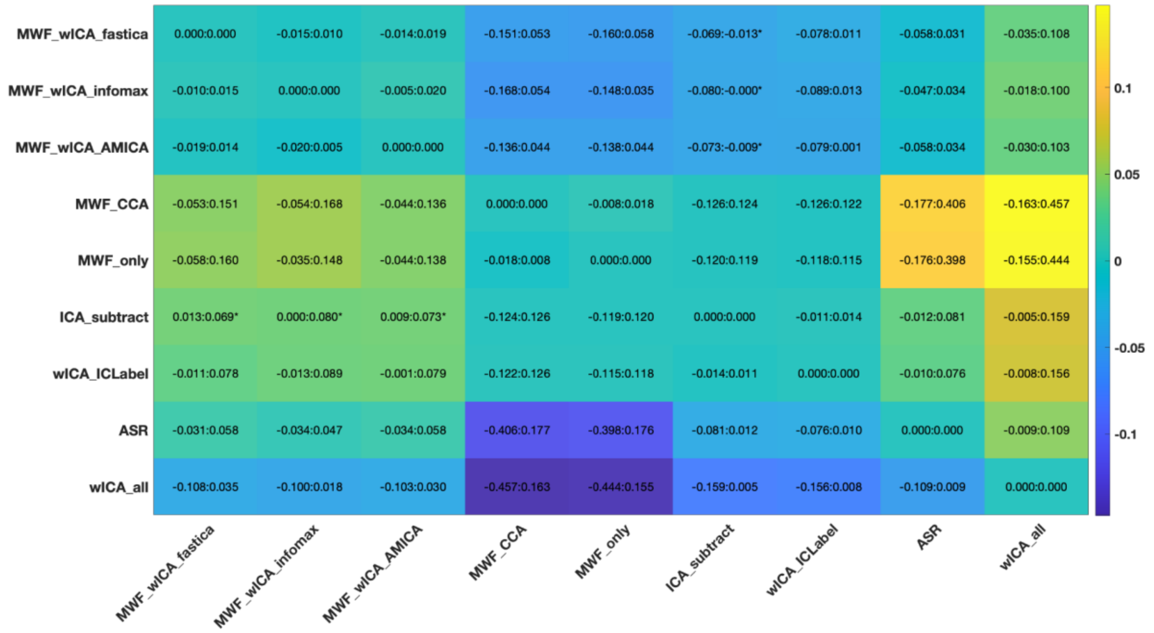

Figure S54. Post-hoc tests for the fBAR values for the Colour Wheel Task dataset.

Similarly, a significant difference was found in allBAR between the pipelines; $F(2.28, 45.53) = 4.184$, $p = 0.018$. However, very few differences were present in the post-hoc tests, with wICA_all performing better than ICA_subtract, wICA_ICLabel, and ASR, and MWF_wICA_infomax and MWF_wICA_AMICA, performing better than ICA_subtract, wICA_ICLabel, and finally, and MWF_wICA_fastICA performing better than ICA_subtract.

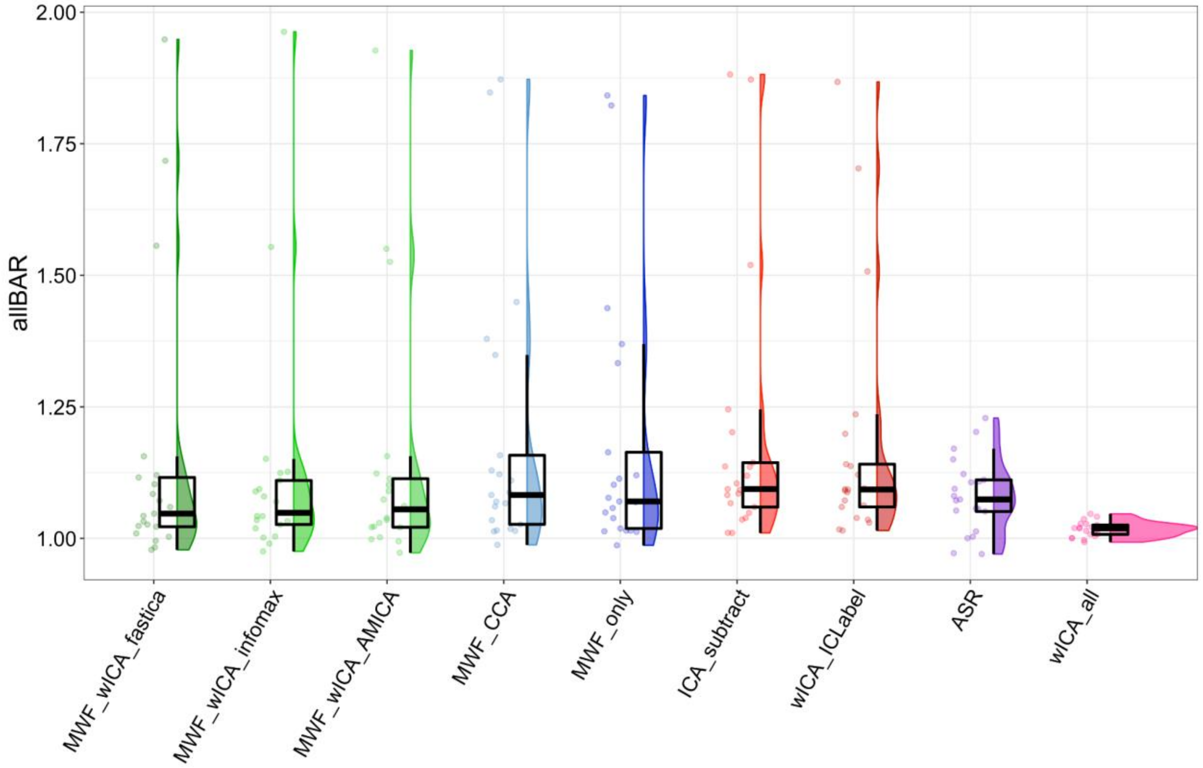

Figure S55. Raincloud plot depicting allBAR values for the Colour Wheel Task dataset.

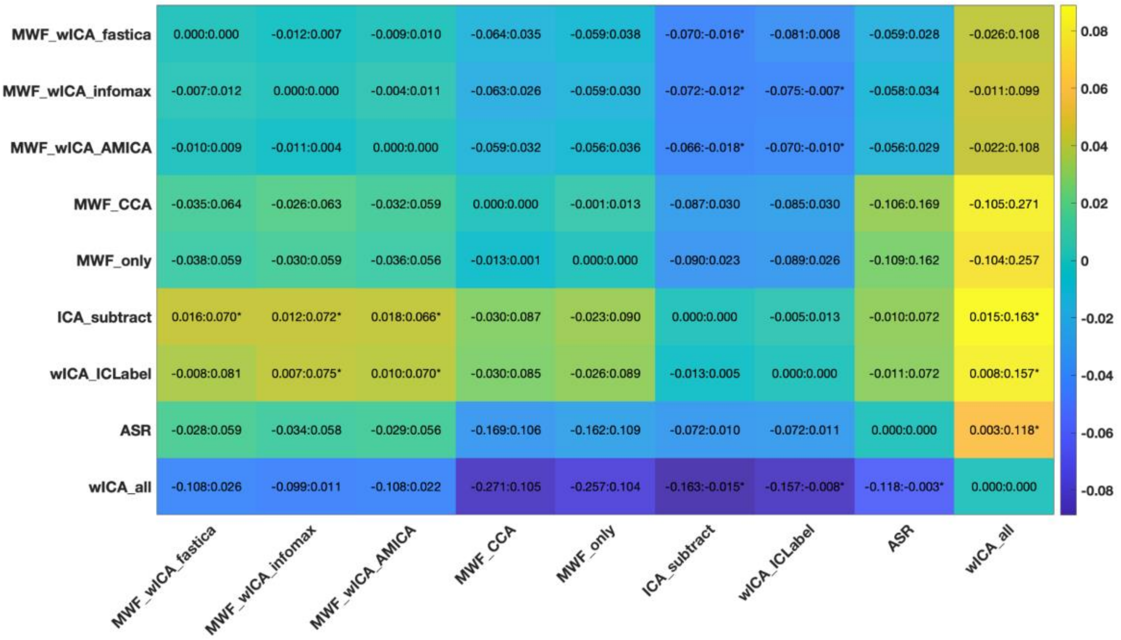

Figure S56. Post-hoc tests for allBAR values for the Colour Wheel Task dataset.

| | Frontal Blink Amplitude Ratio | | All electrode Blink Amplitude Ratio | |
|---|---|---|---|---|
| **Pipeline** | **Mean** | **SD** | **Mean** | **SD** |
| ASR | 1.077 | 0.07 | 1.081 | 0.07 |
| ICA_subtract | 1.273 | 0.486 | 1.19 | 0.253 |
| MWF_CCA | 1.321 | 0.446 | 1.185 | 0.257 |
| MWF_wICA_AMICA | 1.207 | 0.437 | 1.139 | 0.237 |
| MWF_wICA_fastICA | 1.229 | 0.486 | 1.147 | 0.26 |
| MWF_wICA_infomax | 1.184 | 0.43 | 1.123 | 0.226 |
| MWF_only | 1.316 | 0.44 | 1.177 | 0.25 |
| wICA_all | 1.02 | 0.019 | 1.017 | 0.014 |
| wICA_ICLabel | 1.245 | 0.419 | 1.176 | 0.23 |

Table S10. BAR value mean and SDs from the Colour Wheel Task dataset.

##### ***Muscle Activity Statistics for the Colour Wheel Task dataset***

There was a significant difference between the pipelines in the proportion of epochs showing muscle above the threshold after cleaning; $F(1.64, 36.01) = 566.588$, $p < 0.0001$. Post-hoc tests indicated that wICA_all and HAPPE performed more poorly than all other pipelines, but no other pipelines differed from any other pipeline.

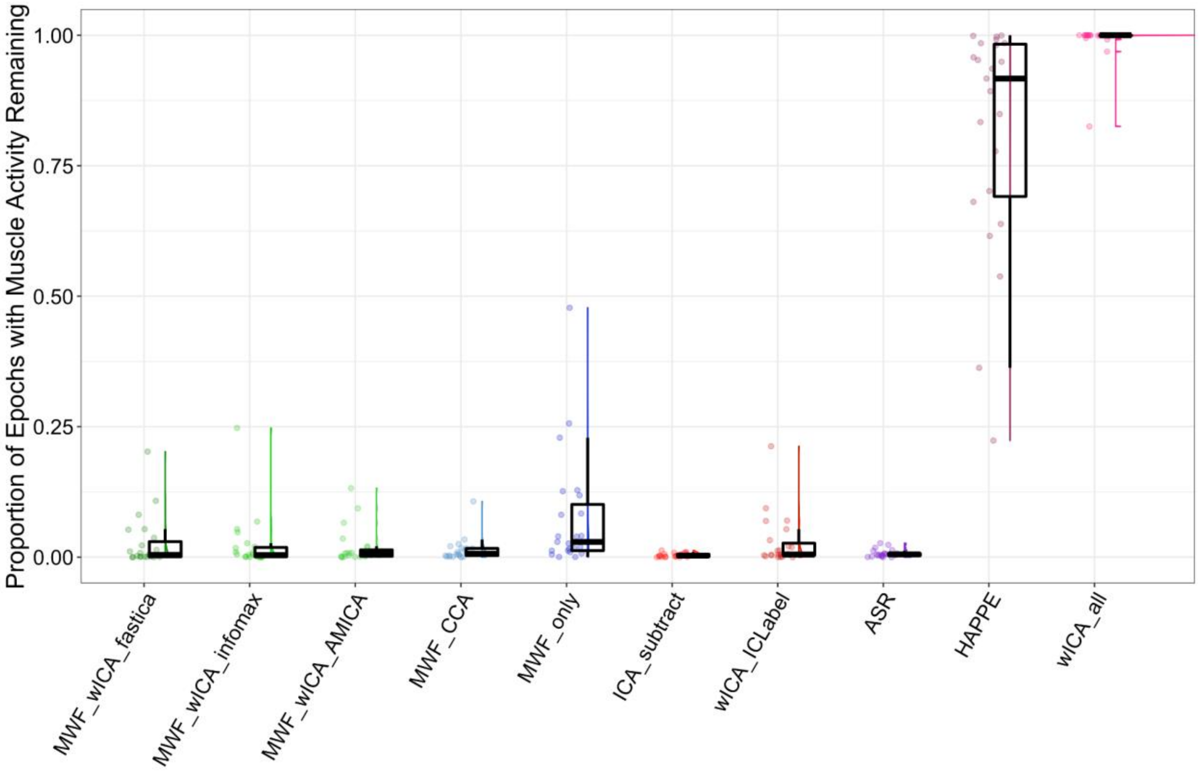

Figure S57. Raincloud plot depicting the proportion of epochs showing muscle activity after cleaning for the Colour Wheel Task dataset.

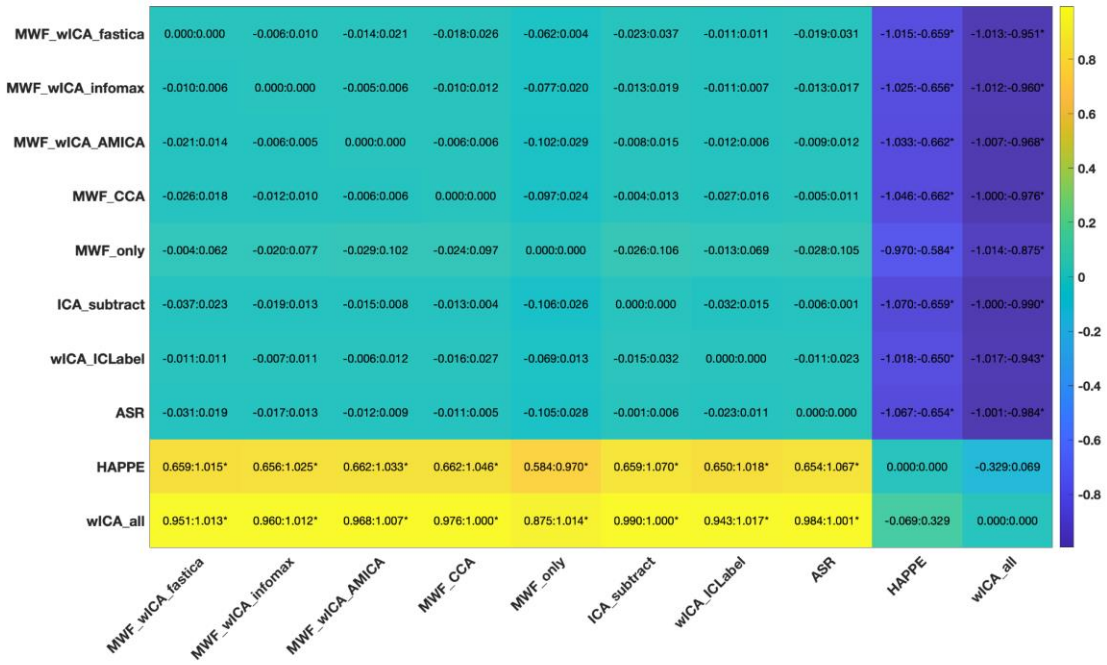

Figure S58. Post-hoc tests for the proportion of epochs showing muscle activity after cleaning for the Colour Wheel Task dataset.

There was a significant difference between the pipelines in the severity by which the log-power log-frequency slope exceeded the muscle threshold within epochs that exceeded the threshold; $F(2.26, 42.86) = 15.214$, $p < 0.0001$. For this metric, post-hoc tests indicated that wICA_all performed more poorly than all other pipelines, followed by HAPPE which performed more poorly than all pipelines except wICA_all. No other pipeline differed from any other pipeline.

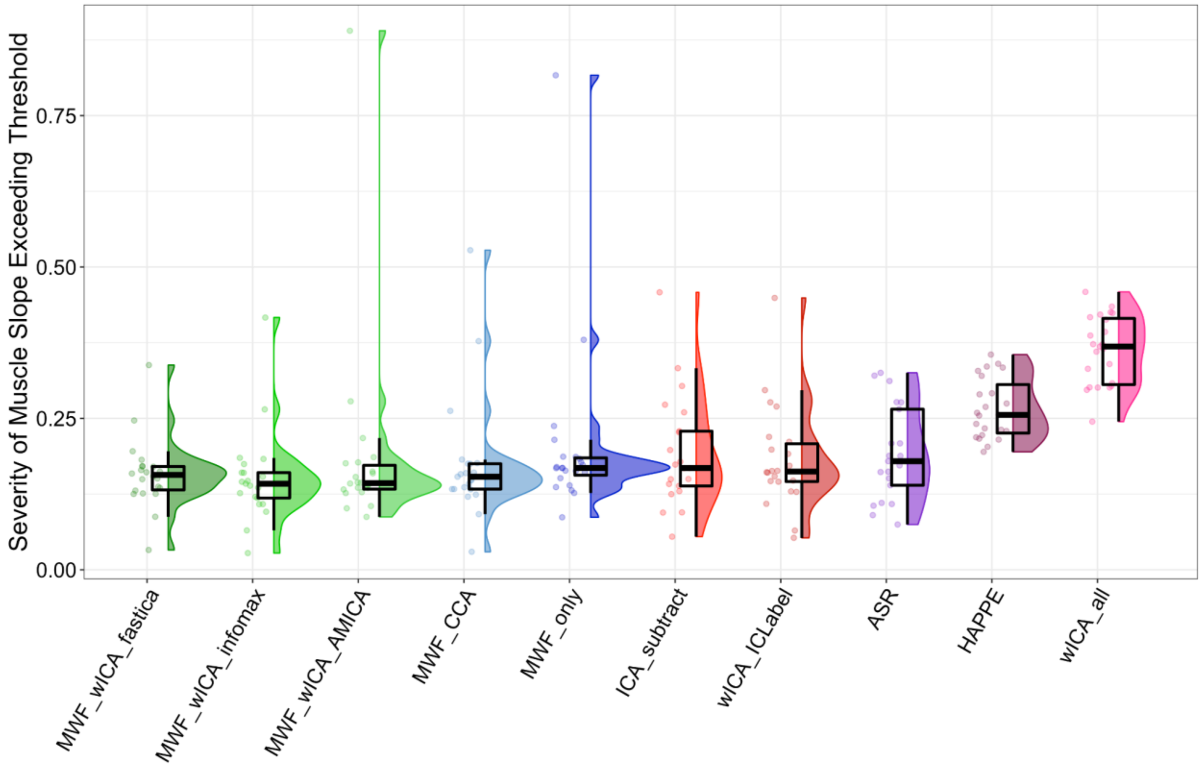

Figure S59. Raincloud plot for the severity by which the log-power log-frequency slopes exceed the threshold in epochs that show slopes indicative of muscle activity after cleaning for the Colour Wheel Task dataset.

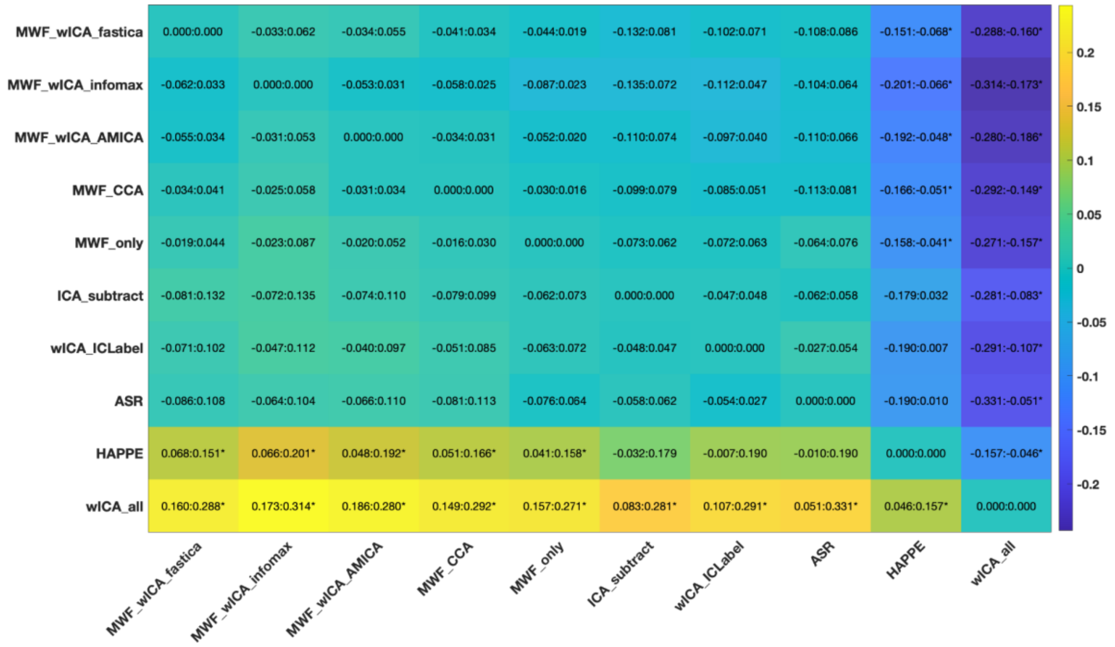

Figure S60. Post-hoc tests for the severity by which the log-power log-frequency slopes exceeded the threshold in epochs showing muscle activity after cleaning.

| | Proportion of epochs showing slopes that indicate muscle activity after cleaning | | Severity of slope that exceeded muscle slope threshold in epochs showing muscle slopes after cleaning | |
|---|---|---|---|---|
| **Pipeline** | **Mean** | **SD** | **Mean** | **SD** |
| **ASR** | 0.007 | 0.007 | 0.192 | 0.078 |
| **HAPPE** | 0.816 | 0.218 | 0.266 | 0.049 |
| **ICA_subtract** | 0.004 | 0.004 | 0.193 | 0.093 |
| **MWF_CCA** | 0.014 | 0.022 | 0.174 | 0.099 |
| **MWF_wICA_AMICA** | 0.019 | 0.034 | 0.185 | 0.163 |
| **MWF_wICA_fastICA** | 0.027 | 0.048 | 0.158 | 0.057 |
| **MWF_wICA_infomax** | 0.023 | 0.053 | 0.151 | 0.077 |
| **MWF_only** | 0.077 | 0.112 | 0.202 | 0.144 |
| **wICA_all** | 0.99 | 0.037 | 0.363 | 0.057 |
| **wICA_ICLabel** | 0.028 | 0.048 | 0.182 | 0.085 |

Table S11. Means and SDs for muscle metrics from the Colour Wheel Task dataset.

#### SECTION SEVEN

##### Supplementary discussion points

When considering the reduction of the influence of blinks, across all the pipelines, it is worth noting that the MWF_wICA methods perform the best across both frontal electrodes and all electrodes. wICA_ICLabel and ICA_subtract performed the worst. Our perspective is that this highlights the advantage of using a two-step process to address blink artifacts, providing redundancy (in case one step inadequately addresses the artifact, the other step is still likely to address it), while each step still provides discrimination between the influence of the artifact and brain activity not influenced by the artifact. As such, if the MWF cleaning completely addresses the blink artifact, ICLabel will not detect any blink artifact and thus will not attempt to clean periods that were affected by blinks in the raw data, preventing overcleaning. However, it seems that not all two-step cleaning approaches were as effective. In contrast to the results for MWF_wICA, the ASR method (which also applied ICA_subtract) appeared to inadequately correct for blinks.

In terms of the amount of variance detected by ICLabel to be explained by brain activity after cleaning, the MWF_wICA methods and wICA_ICLabel performed the best, better than MWF_only and wICA_all. Additionally, while our data also indicated that MWF_wICA_infomax may perform better than MWF_wICA_fastICA and MWF_wICA_AMICA on this metric, this may be a product of the infomax algorithm being used both for cleaning and for detection of brain variance in the measure, the effect of which we are not sure, but which may have biased results towards that cleaning pipeline.

MWF_only provided slightly lower ARR values (similar to MWF_CCA and wICA_ICLabel), but higher SER values than the MWF_wICA methods (and MWF_CCA and wICA_ICLabel). As such, if the aim of a study were to maximise SER (indicating minimal adjustment of EEG periods deemed to be clean by our initial artifact marking template), MWF_only could be recommended over MWF_wICA, MWF_CCA and wICA_ICLabel methods. However, MWF_only led to less variance explained by the experimental manipulation for working memory metrics and resting alpha RMS, a lower proportion of variance explained by brain activity after cleaning when measured by ICLabel, and MWF_only left more blink and muscle artifact than other methods, so may not eliminate artifacts as effectively, producing less reliable results (with less power to detect differences in experimental designs). Part of this is likely to be due to the fact that the MWF cleaning was only set up to clean blinks, muscle activity, horizontal eye movements and drift (and not other atypical artifacts), so may have left noise behind that reduced the ability to detect the experimental effects compared to the more effective cleaning pipelines. Part of the explanation is also likely to be due to the benefits of the double artifact reduction approach used by the methods that applied both MWF cleaning and then wICA or CCA, increasing the cleaning efficacy in these pipelines compared to MWF_only. We think it is beneficial to apply wICA after MWF, both to catch atypical artifacts, and to address artifacts that MWF cleaning alone might have not cleaned completely (muscle activity seems to be the most significant example of activity that MWF cleaning missed in many cases).

Conversely, wICA_all and HAPPE provided very high ARR values, but at the expense of also providing very low SER values. These methods lead to power-frequency slopes that were very shallow, which would typically suggest muscle activity remaining after cleaning due to a high proportion of high frequency power. We suspect the shallow slopes may be driven more by the removal of low brain activity by these methods, as a result of applying wICA to all components (leading to the removal of considerable brain activity from the data as many components are primarily brain activity rather than artifact). Consistent with this interpretation, wICA_all and HAPPE provided much smaller values for many measures of variance explained by the experimental manipulation (with a few specific counter-examples to this trend that may have been produced by the almost complete elimination of alpha brain activity in the low strength activity condition). However, it is also worth noting that wICA_all and HAPPE both showed more power in the beta frequencies relative to the total spectrum than other pipelines, so it may also be that wICA_all and HAPPE are inadequately cleaning muscle activity, despite over-cleaning the signal.

For researchers interested in connectivity measures, it may be useful to note that cleaning EEG data using wICA to reduce artifact components (and only the artefactual contribution of the artifact component, in theory) does not reduce the rank of the data, so might allow for higher resolution of nodes when using connectivity analysis in the source space (in contrast to subtracting ICA components, which does reduce the rank) [5]. We think that it is also worth noting that although the difference was small, our results seemed to suggest that applying ICA or wICA in a pipeline reduced the difference in the distribution of alpha activity between eyes open and eyes closed resting compared to the analogous method that did not apply ICA or wICA (MWF_only > all MWF_wICA methods and MWF_CCA > MWF_wICA_CCA). However, we could see no obvious reason for this, and note that inspection of the power-frequency plots from parieto-ocipital electrodes indicates that the ICA_subtract and wICA_ICLabel pipelines led to the largest values for alpha power, suggesting these methods were the best at preserving the alpha oscillations in these electrodes where the alpha signal is most prominent.

##### *Limitations and Potential improvements*

It is worth noting that the SER and ARR metrics may be biased towards pipelines that cleaned the same time periods as those used to calculate ARR (for example, those that used MWF). The periods used to calculate SER and ARR were contained blinks, horizontal eye movement, muscle activity and voltage drift. However, atypical artifacts (that do not belong to these categories) were not included in the artifact templates for computation of the SER and ARR cleaning efficacy metrics, potentially biasing these metrics against cleaning methods that address atypical artifacts (such as ICA_subtract and ASR), and towards methods that do not clean atypical artifacts (such as MWF_only, which may explain the high SER values for this pipeline). As such, to provide a fair assessment of each of the pipelines, we have also included other metrics such as remaining muscle, blink amplitude ratio, epochs remaining after cleaning and epoch rejection, the variance explained by neural activity after ICA component selection by ICLabel, the reliability of extracted ERPs, and the variance explained by the experimental manipulation (as a pragmatic measure of high practical importance). These other methods aligned with the SER and ARR metrics in suggesting MWF_wICA was amongst the best performing pipeline at cleaning artifacts.

Our data was all sampled at 1000Hz. Higher sampling rates may require more of a computer's resources and slow down the cleaning pipeline. However, 1000Hz is a sufficiently high sampling rate for the study of essentially all ERP or oscillation analyses. Sampling rates as low as 250Hz are still cleaned effectively be our pipeline (however, we have not tested the pipeline on sampling rates lower than this). Additionally, the MWF cleaning of large files can use all a computer's RAM if the computer has <8GB of RAM. We recommend using a computer with more RAM (much of our data was processed using 32GB of RAM) or reducing the sampling rate of the data if "out of memory" errors occur.

There are a number of potential improvements we think are worth exploring in future artifact cleaning pipeline development. Firstly, taking into account temporal data when performing the ICA strikes us as potentially valuable. This can be achieved by using independent vector analysis [37] instead of more traditional ICA methods. Previous research has suggested this indeed does lead to improved artifact reduction [12]. However, in our preliminary tests of an independent vector analysis approach, we were unable to work out how to obtain sensible decompositions or cleaned data using these methods, so we suggest that more explanatory or tutorial publications on these methods would be helpful.

Another potential improvement is that using adaptive wICA thresholds rather than a fixed threshold may improve the separation of signal and artifact, leading to improved cleaning with higher levels of signal left after cleaning. Currently, RELAX implements wICA with a fixed threshold (mult = 1). Lower thresholds may reduce the potential neural activity in a component (the low amplitude activity within a component) by a larger amount, whereas higher thresholds reduce this low amplitude activity within the component by less. Adaptive wICA thresholds have been implemented in previous research and researchers have suggested the default threshold removes too much of the signal [38]. We were unable to find an evidence-base for an optimal threshold, so we used the default approach. We also experimented with using a data driven approach to set the threshold, with level dependent Bayes as suggested in the updated version of HAPPE (HAPPILEE) [39]. However, we found this approach to perform more poorly than the standard threshold (generally not cleaning artifacts as well as standard wICA applied to artifact components). Ideally, we think the threshold should be based on the type of artifact, and perhaps even the confidence that ICLabel provides that the component is an artifact, with components that are more likely to be artifact being more heavily reduced by the wICA approach. wICA could even be adapted to specifically focus on the frequencies that should be minimized in a specific artifact component, for example high frequencies for muscle activity components.

Additionally, although we used 60 microvolts as our epoch rejection criteria after cleaning, we noticed that for some participants with very large alpha activity, some epochs were rejected that did not appear from visual inspection to be contaminated with artifacts, but rather just showed very high alpha power. We would suggest perhaps using 100 microvolts as the criteria for remaining artifacts. We also noticed that all data cleaning methods seemed to reduce the amplitude of the higher-powered alpha oscillation periods. We suspect this might be due to a mixing of alpha activity into the MWF artifact cleaning templates and the ICA artifact components. It may be that adjusting the wICA threshold or having fewer MWF cleaning steps could address this at least in part, but we suggest future research explore more sophisticated methods to address the discrimination of signal and artifact to avoid cleaning signal from the data.

Finally, we note that for future cleaning pipeline development, in order to optimise a cleaning pipeline, some measures need to be considered together. For example, a pipeline is only superior to previous pipelines if SER and ARR measures are both higher concurrently, or if SER is higher while ARR remains the same (or vice versa). Blink amplitude ratio should be as close to 1 as possible and as few epochs showing muscle slope and as small muscle slopes as possible in epochs that do show muscle activity remaining. However, if these measures are reduced and at the same time the variance explained by the experimental manipulation is reduced, the pipeline is not an improvement over previous pipelines (unless it can be demonstrated that the variance explained by the experimental manipulation in a previous pipeline was due to an artifact alone, and not brain activity).

#### Cleaned dataset examples

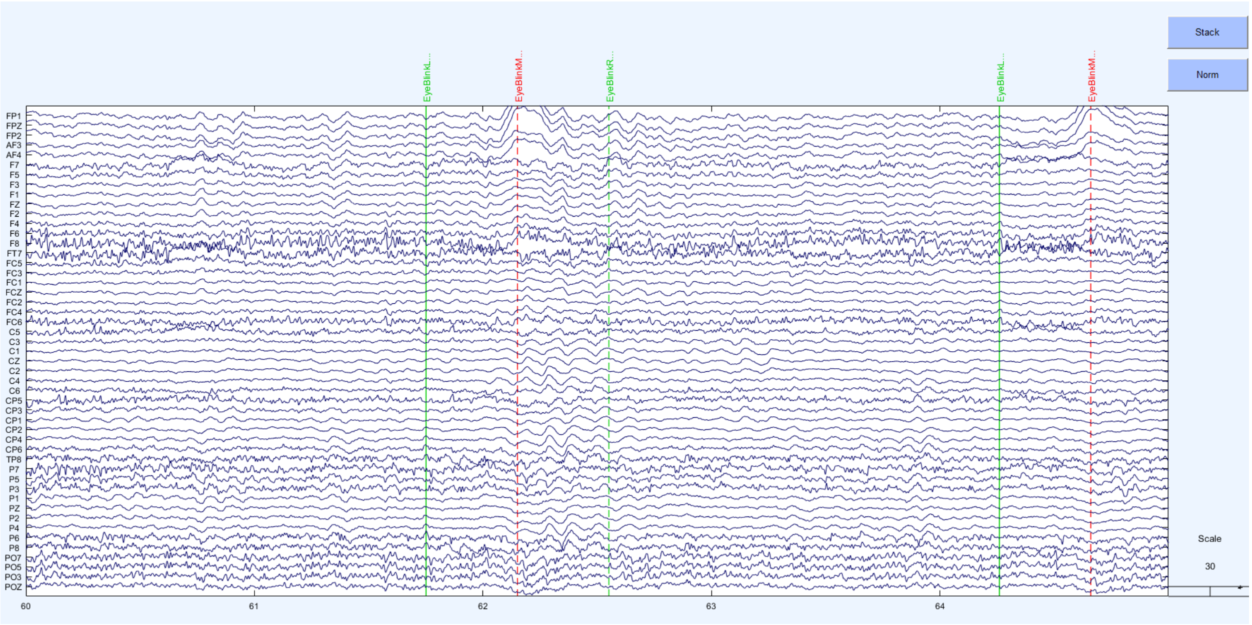

Figure S61. Raw data example

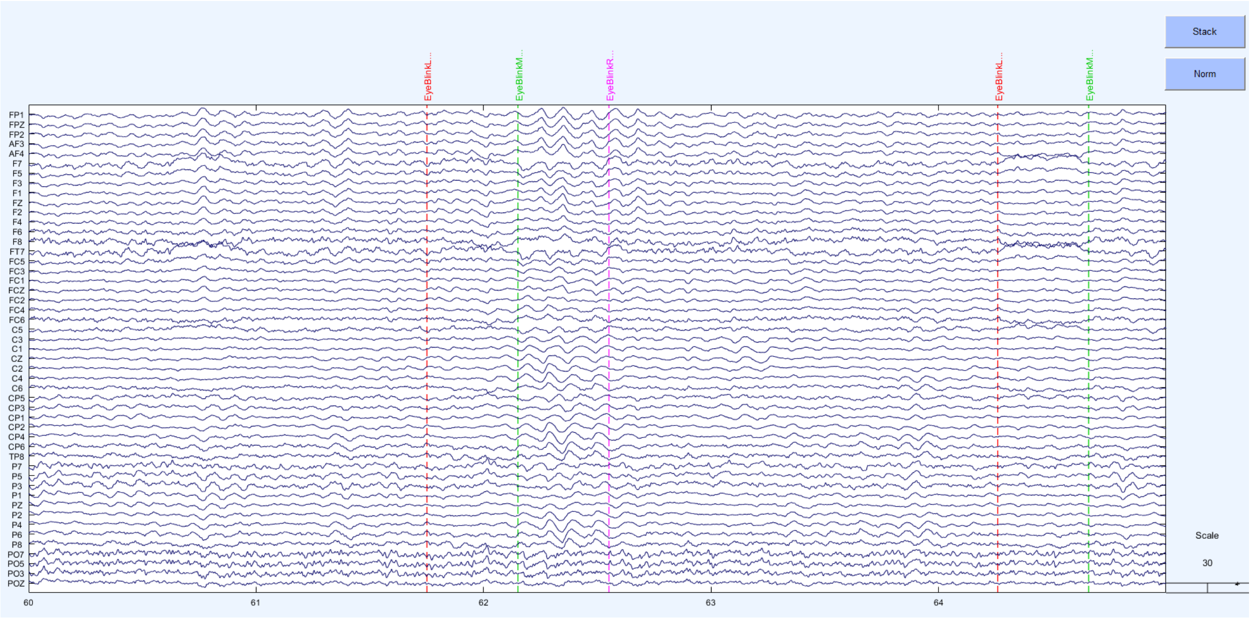

Figure S62. MWF_wICA_infomax cleaned example

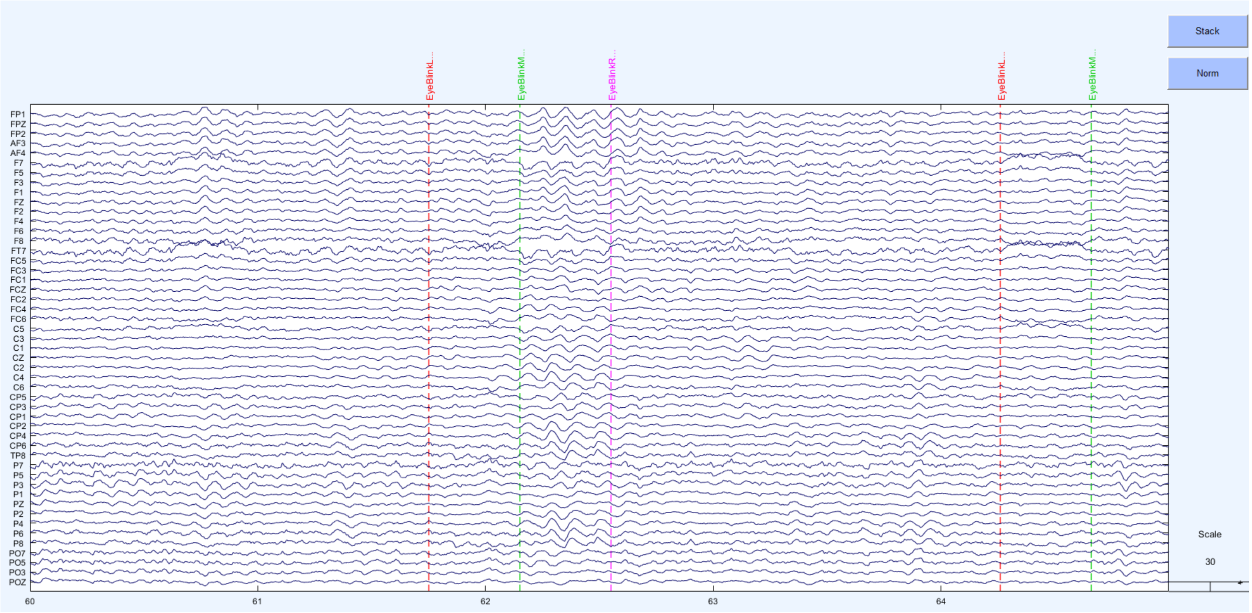

Figure S63. MWF_ICA_subtract cleaned example

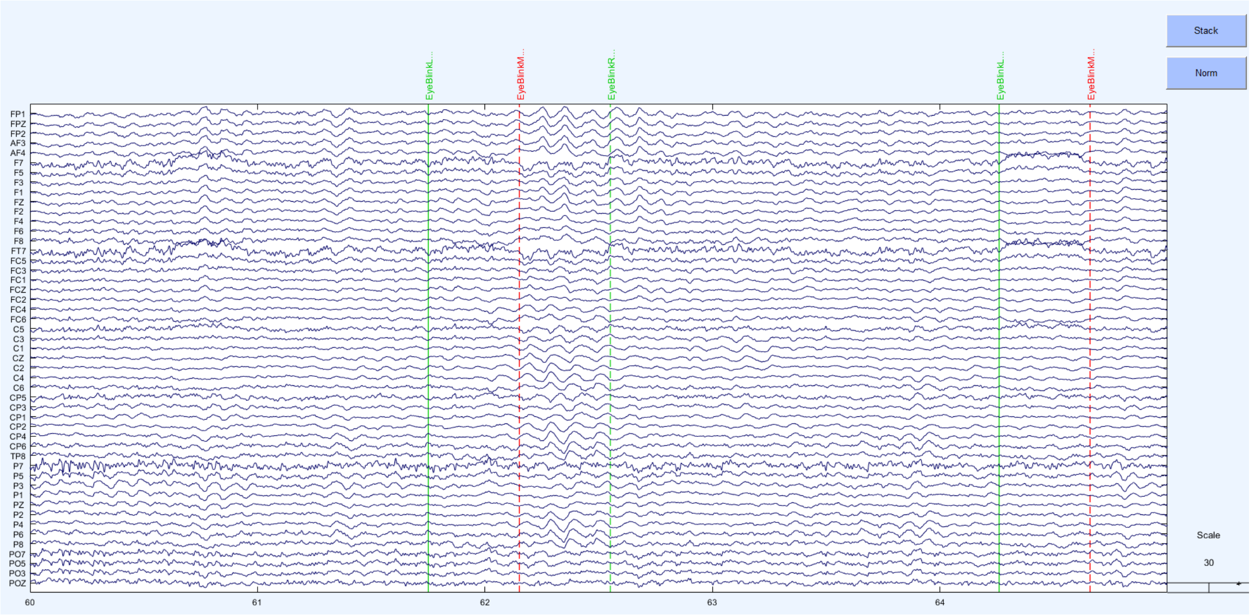

Figure S64. ICA_subtract cleaned example

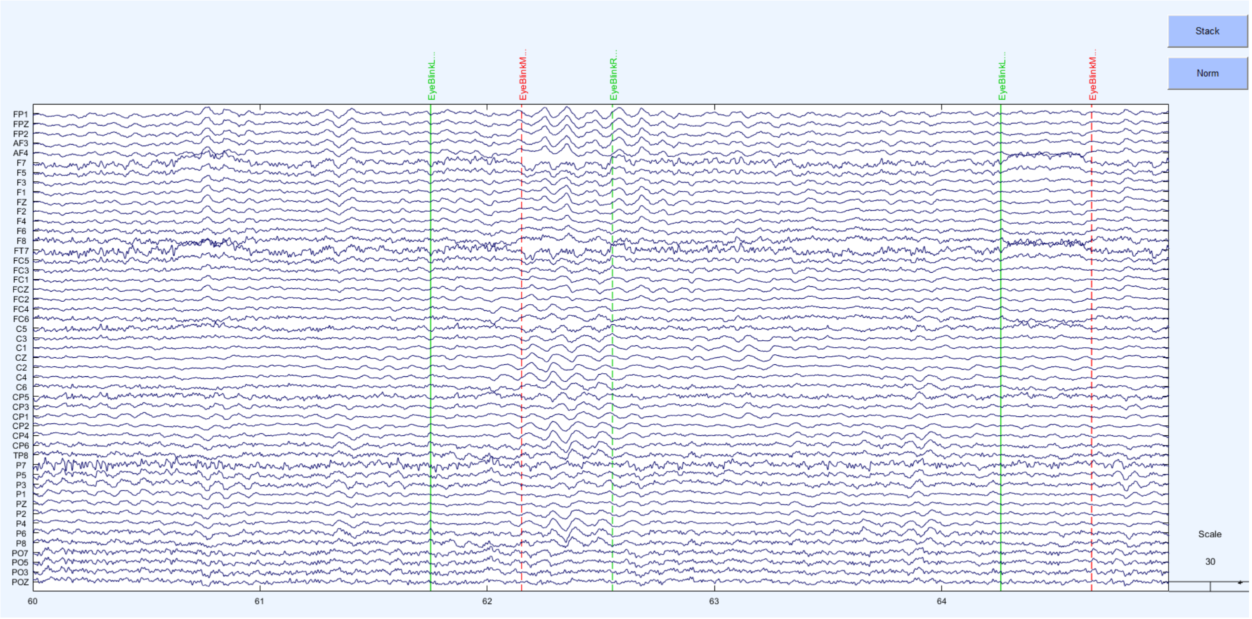

Figure S65. wICA_ICLabel cleaned example

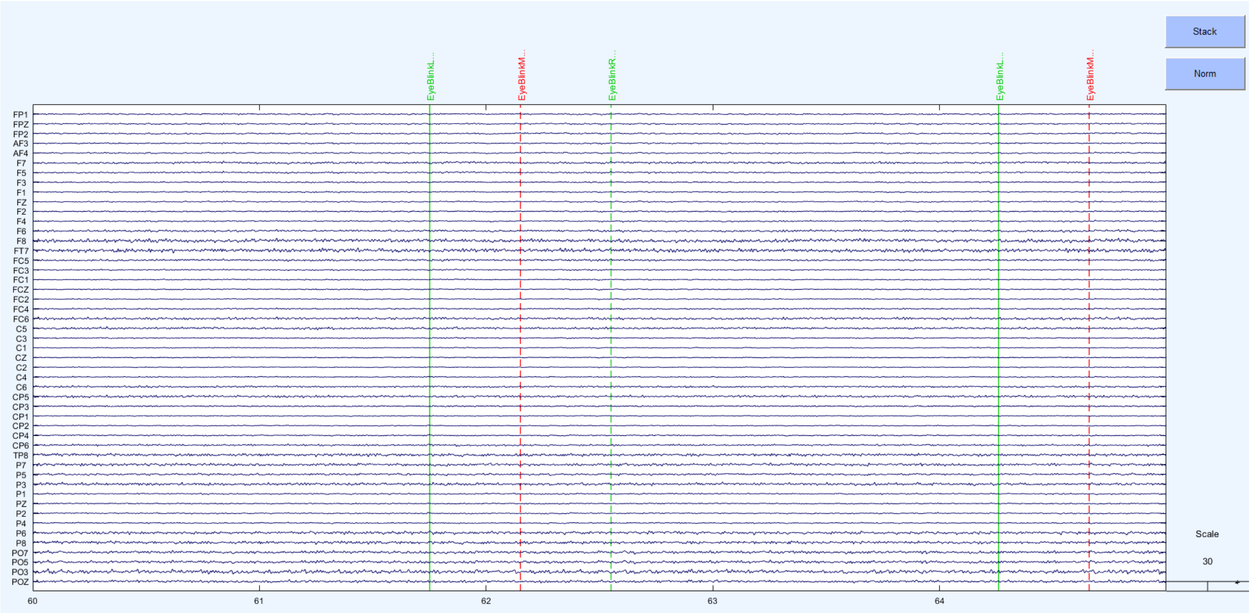

Figure S66. wICA_all cleaned dataset

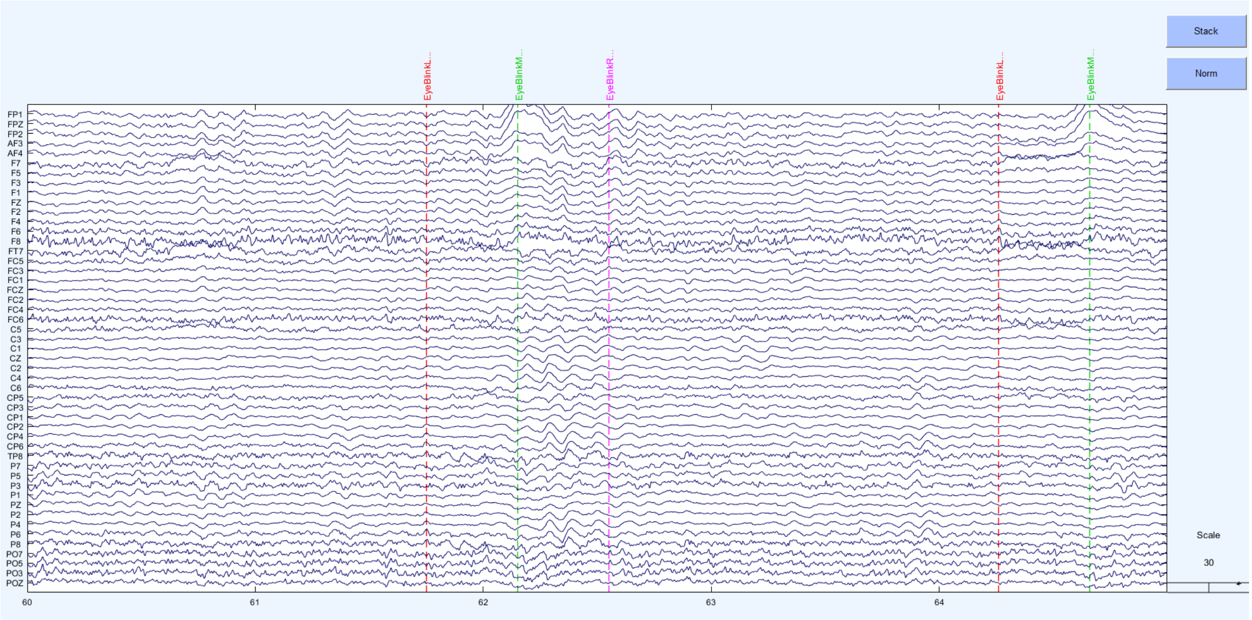

Figure S67. MWF_only cleaned example

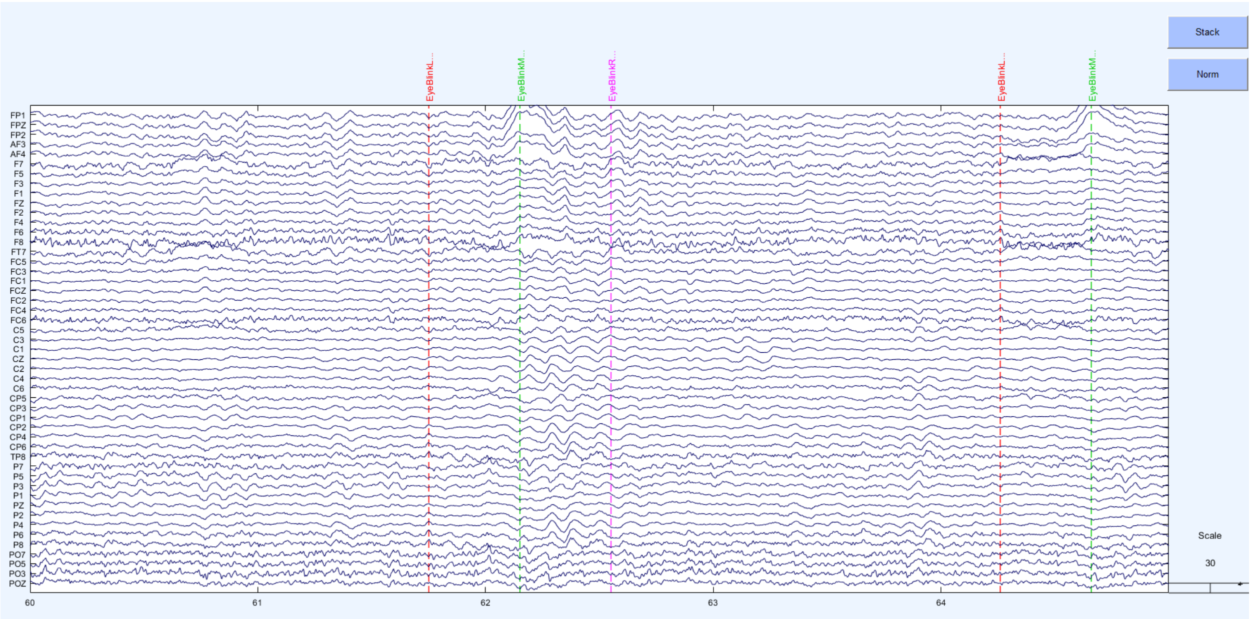

Figure S68. MWF_CCA cleaned example dataset

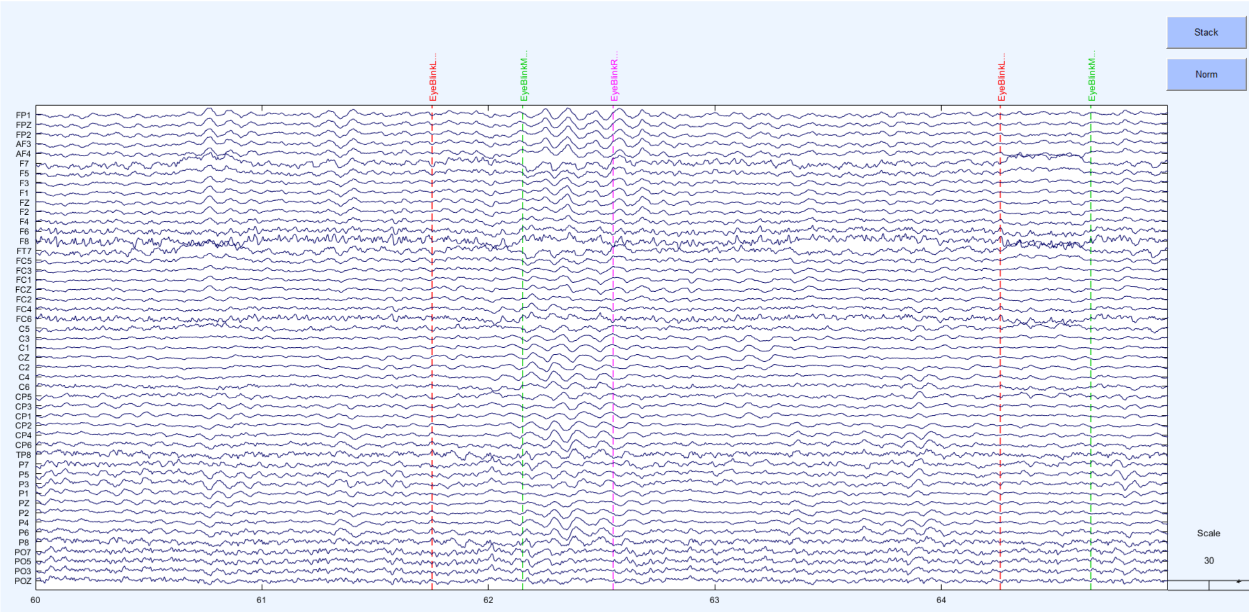

Figure S69. MWF_wICA_CCA cleaned dataset

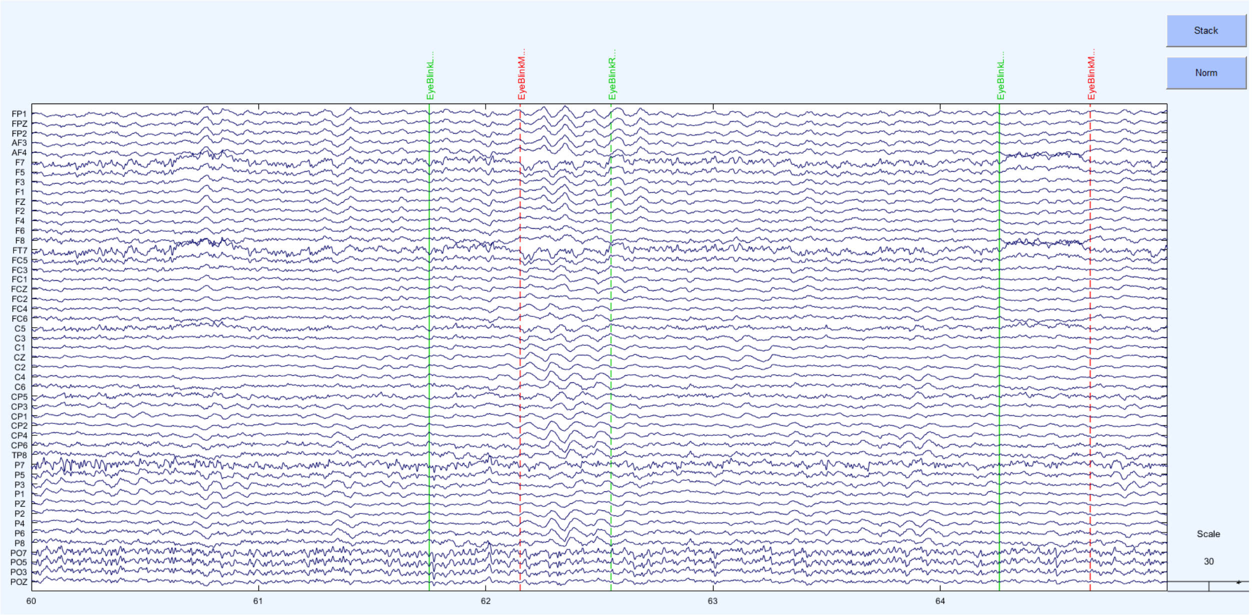

Figure S70. ASR cleaned dataset

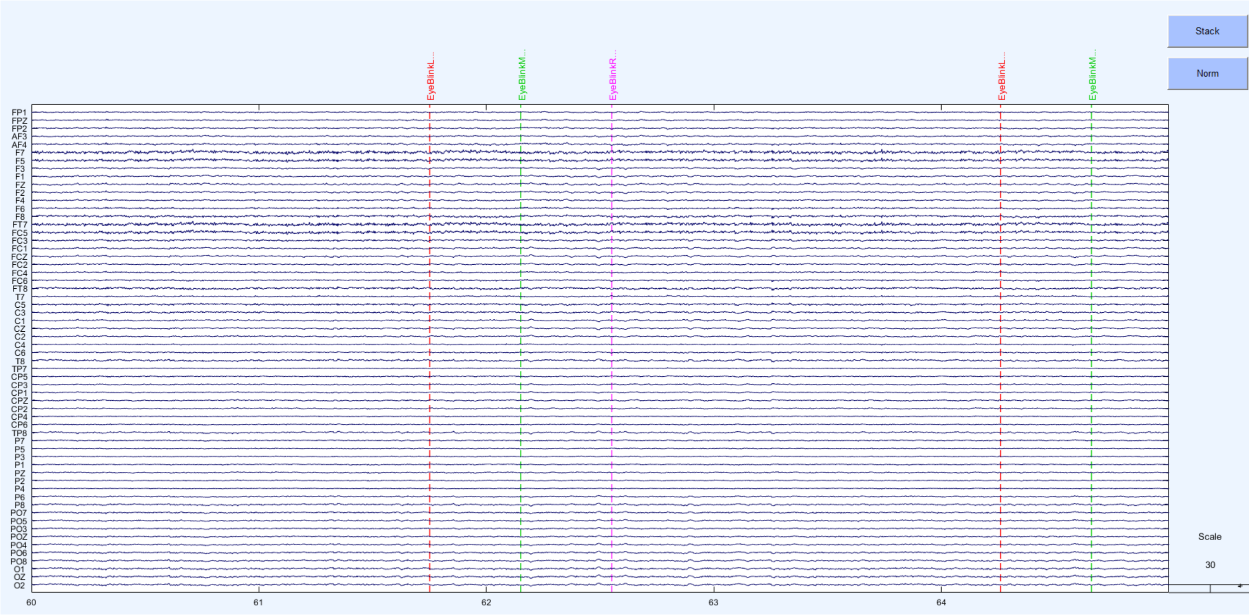

Figure S71. HAPPE cleaned dataset

#### Supplementary Materials References

1. Gerla V, Kremen V, Covassin N, Lhotska L, Saifutdinova E, Bukartyk J, et al. Automatic identification of artifacts and unwanted physiologic signals in EEG and EOG during wakefulness. Biomedical Signal Processing and Control. 2017;31:381-90.
2. Muthukumaraswamy S. High-frequency brain activity and muscle artifacts in MEG/EEG: a review and recommendations. Frontiers in human neuroscience. 2013;7:138.
3. Schlögl A, Keinrath C, Zimmermann D, Scherer R, Leeb R, Pfurtscheller G. A fully automated correction method of EOG artifacts in EEG recordings. Clinical neurophysiology. 2007;118(1):98-104.
4. Zeng H, Song A. Removal of EOG artifacts from EEG recordings using stationary subspace analysis. The Scientific World Journal. 2014;2014.
5. Castellanos NP, Makarov VA. Recovering EEG brain signals: Artifact suppression with wavelet enhanced independent component analysis. Journal of neuroscience methods. 2006;158(2):300-12.
6. Somers B, Francart T, Bertrand A. A generic EEG artifact removal algorithm based on the multi-channel Wiener filter. Journal of neural engineering. 2018;15(3):036007.
7. Bigdely-Shamlo N, Mullen T, Kothe C, Su K-M, Robbins KA. The PREP pipeline: standardized preprocessing for large-scale EEG analysis. Frontiers in neuroinformatics. 2015;9:16.
8. Fitzgibbon S, DeLosAngeles D, Lewis T, Powers D, Grummett T, Whitham E, et al. Automatic determination of EMG-contaminated components and validation of independent component analysis using EEG during pharmacologic paralysis. Clinical Neurophysiology. 2016;127(3):1781-93.
9. de Cheveigné A, Arzounian D. Robust detrending, rereferencing, outlier detection, and inpainting for multichannel data. NeuroImage. 2018;172:903-12.
10. Nolan H, Whelan R, Reilly RB. FASTER: fully automated statistical thresholding for EEG artifact rejection. Journal of neuroscience methods. 2010;192(1):152-62.
11. Rogasch NC, Sullivan C, Thomson RH, Rose NS, Bailey NW, Fitzgerald PB, et al. Analysing concurrent transcranial magnetic stimulation and electroencephalographic data: A review and introduction to the open-source TESA software. Neuroimage. 2017;147:934-51.
12. Barban F, Chiappalone M, Bonassi G, Mantini D, Semprini M. Yet another artefact rejection study: an exploration of cleaning methods for biological and neuromodulatory noise. Journal of Neural Engineering. 2021.
13. van Driel J, Olivers CN, Fahrenfort JJ. High-pass filtering artifacts in multivariate classification of neural time series data. Journal of Neuroscience Methods. 2021;352:109080.
14. Kleifges K, Bigdely-Shamlo N, Kerick SE, Robbins KA. BLINKER: Automated extraction of ocular indices from EEG enabling large-scale analysis. Frontiers in neuroscience. 2017;11:12.
15. Gabard-Durnam LJ, Mendez Leal AS, Wilkinson CL, Levin AR. The Harvard Automated Processing Pipeline for Electroencephalography (HAPPE): standardized processing software for developmental and high-artifact data. Frontiers in neuroscience. 2018;12:97.
16. Mullen T. CleanLine EEGLAB plugin. San Diego, CA: Neuroimaging Informatics Toolsand Resources Clearinghouse (NITRC). 2012.
17. Winkler I, Haufe S, Tangermann M. Automatic classification of artifactual ICA-components for artifact removal in EEG signals. Behavioral and Brain Functions. 2011;7(1):1-15.
18. Chang C-Y, Hsu S-H, Pion-Tonachini L, Jung T-P. Evaluation of artifact subspace reconstruction for automatic artifact components removal in multi-channel EEG recordings. IEEE Transactions on Biomedical Engineering. 2019;67(4):1114-21.

19. Pion-Tonachini L, Kreutz-Delgado K, Makeig S. ICLabel: An automated electroencephalographic independent component classifier, dataset, and website. NeuroImage. 2019;198:181-97.
20. Issa MF, Juhasz Z. Improved EOG artifact removal using wavelet enhanced independent component analysis. Brain sciences. 2019;9(12):355.
21. Mammone N, La Foresta F, Morabito FC. Automatic artifact rejection from multichannel scalp EEG by wavelet ICA. IEEE Sensors Journal. 2011;12(3):533-42.
22. Janani AS, Grummett TS, Lewis TW, Fitzgibbon SP, Whitham EM, DelosAngeles D, et al. Improved artefact removal from EEG using Canonical Correlation Analysis and spectral slope. Journal of neuroscience methods. 2018;298:1-15.
23. De Clercq W, Vergult A, Vanrumste B, Van Paesschen W, Van Huffel S. Canonical correlation analysis applied to remove muscle artifacts from the electroencephalogram. IEEE transactions on Biomedical Engineering. 2006;53(12):2583-7.
24. Gao J, Zheng C, Wang P. Online removal of muscle artifact from electroencephalogram signals based on canonical correlation analysis. Clinical EEG and neuroscience. 2010;41(1):53-9.
25. Raimondo F, Kamienkowski JE, Sigman M, Fernandez Slezak D. CUDAICA: GPU optimization of infomax-ICA EEG analysis. Computational intelligence and neuroscience. 2012;2012.
26. Hyvarinen A, editor Fast ICA for noisy data using Gaussian moments. 1999 IEEE international symposium on circuits and systems (ISCAS); 1999: IEEE.
27. Palmer JA, Kreutz-Delgado K, Makeig S. AMICA: An adaptive mixture of independent component analyzers with shared components. Swartz Center for Computatonal Neursoscience, University of California San Diego, Tech Rep. 2012.
28. Zakeri Z. Optimised use of independent component analysis for EEG signal processing: University of Birmingham; 2017.
29. Bertrand A. Distributed signal processing for wireless EEG sensor networks. IEEE Transactions on Neural Systems and Rehabilitation Engineering. 2015;23(6):923-35.
30. Somers B, Bertrand A. Removal of eye blink artifacts in wireless EEG sensor networks using reduced-bandwidth canonical correlation analysis. Journal of neural engineering. 2016;13(6):066008.
31. Robbins KA, Touryan J, Mullen T, Kothe C, Bigdely-Shamlo N. How sensitive are EEG results to preprocessing methods: a benchmarking study. IEEE transactions on neural systems and rehabilitation engineering. 2020;28(5):1081-90.
32. Bailey NW, Freedman G, Raj K, Spierings KN, Piccoli LR, Sullivan CM, et al. Mindfulness meditators show enhanced accuracy and different neural activity during working memory. Mindfulness. 2020;11:1762-81.
33. Clayson PE, Baldwin S, Rocha H, Larson MJ. The Data-Processing Multiverse of Event-Related Potentials (ERPs): A Roadmap for the Optimization and Standardization of ERP Processing and Reduction Pipelines. 2021.
34. Habermann M, Weusmann D, Stein M, Koenig T. A student's guide to randomization statistics for multichannel event-related potentials using ragu. Frontiers in neuroscience. 2018;12:355.
35. Koenig T, Kottlow M, Stein M, Melie-García L. Ragu: a free tool for the analysis of EEG and MEG event-related scalp field data using global randomization statistics. Computational intelligence and neuroscience. 2011;2011.
36. Benjamini Y, Hochberg Y. Controlling the false discovery rate: a practical and powerful approach to multiple testing. Journal of the Royal statistical society: series B (Methodological). 1995;57(1):289-300.
37. Lee I, Kim T, Lee T-W. Fast fixed-point independent vector analysis algorithms for convolutive blind source separation. Signal Processing. 2007;87(8):1859-71.
38. Zima M, Tichavský P, Paul K, Krajča V. Robust removal of short-duration artifacts in long neonatal EEG recordings using wavelet-enhanced ICA and adaptive combining of tentative reconstructions. Physiological measurement. 2012;33(8):N39.

39. Lopez K, Monachino A, Morales S, Leach S, Bowers M, Gabard-Durnam L. HAPPILEE: The Harvard Automated Processing Pipeline In Low Electrode Electroencephalography, a standardized software for low density EEG and ERP data. bioRxiv. 2021.
